# SURF2 is a MDM2 antagonist in triggering the nucleolar stress response

**DOI:** 10.1101/2024.01.09.574617

**Authors:** Sophie Tagnères, Paulo Espirito Santo, Julie Radermecker, Dana Rinaldi, Carine Froment, Quentin Provost, Solemne Capeille, Nick Watkins, Julien Marcoux, Pierre-Emmanuel Gleizes, Virginie Marcel, Célia Plisson-Chastang, Simon Lebaron

## Abstract

Cancer cells are addicted to strong ribosome production to sustain their proliferation rate. Many chemotherapies impede ribosome production which is perceived by cells as “nucleolar stress” (NS), triggering p53-dependent and independent response pathways leading to cell cycle arrest and/or apoptosis. The 5S RNP particle, a sub-ribosomal particle, is instrumental to NS response. Upon ribosome assembly defects, the 5S RNP accumulate as free form. This free form is able to sequester and inhibit MDM2, thus promoting p53 stabilization. To investigate how cancer cells can resist to NS, we purified free-5S RNP and uncovered a new interaction partner, SURF2. Functional characterization of SURF2 shows that its depletion increases cellular sensitivity to NS, while its overexpression promotes their resistance to it. Consistently, SURF2 expression level negatively correlates with the overall survival in adrenocortical and head and neck squamous cell carcinomas. Our data demonstrate that SURF2 buffers free-5S RNP particles, and can modulate their activity. SURF2 regulates NS responses, and is a key player in both ribosomopathies and oncogenic mechanisms.

## Introduction

Cancers originate from accumulation of genetic events that allow uncontrolled cell proliferation. Thanks to large-scale sequencing of the cancer genome, it is now possible to identify cancer-specific alterations. This information can then be used for diagnosis but also allows the development of new therapeutic approaches for specific targets, i.e. targeted therapies (*1*). Although these targeted therapies have proven to be effective, they are also associated with side effects that hinder their therapeutic advantages. Indeed, drug resistance, induced either by mutation of the specific target or the activation of alternative survival pathways is often observed (*2*). Combination therapies are therefore needed to modulate the metabolic circuits that support proliferation in cancer cells. A key metabolic pathway cancer cells is the abnormal boost in ribosome synthesis to meet the protein requirements associated with uncontrolled cell growth and proliferation (*3*). The nucleolus is the nuclear sub-compartment where ribosome synthesis begins, and the boost in ribosome synthesis is associated with increased nucleolus size. This phenotype has long been used as a marker of tumor cells aggressiveness (*3*, *4*). Cancer cells are thus addicted to ribosome synthesis.

Nucleolar stress (NS) can be defined as any stress leading to the inhibition of ribosome assembly, a complex and energy-consuming process that begins with transcription of rDNA by the specific RNA polymerase I (Pol I) in the nucleolus (*5*). Under stress, cells reduce ribosome production to conserve energy and stop cell proliferation (*6*). These cell-specific regulations of ribosome synthesis are thus essential to the control of cell growth in all living cells. Nevertheless, mature ribosomal subunits are stable entities, so the inhibition of their synthesis will only affect cellular protein homeostasis after long-term exposure to stress. In addition to these long-term regulations, ribosome synthesis shutdown upon stress signaling is directly linked to cell cycle arrest (*7*, *8*). In humans, this is induced by stabilization and activation of the tumor suppressor p53 (*7*, *9–13*). This pathway is prevalent in the regulation of p53 under normal and pathological conditions (*14–16*). Several ribosomal proteins have been described as being able to mediate such p53 regulation in response to stress. The general idea is that ribosomal proteins that are released from or not integrated to ribosomes can accumulate as free form in the nucleoplasm. There, they can directly bind MDM2 and inhibit its ubiquitinylase activity (*17–21*). Such role for ribosomal proteins has been generally termed as extra-ribosomal function.

However, among the different ribosomal proteins that support such regulation of p53 activation only RPL5 and RPL11 are instrumental to this response. Interestingly, both proteins are components of the same complex, the 5S RNP particle. (*14*, *22–24*). 5S RNPs consist of the association of 5S ribosomal RNA (rRNA) with the ribosomal proteins RPL5 and RPL11. These particles are largely incorporated into nascent large/60S ribosomal subunits. Disruption of ribosome synthesis results in the accumulation of so-called free-5S RNPs in the nucleoplasm. Free 5S RNPs interact with the E3-ubiquitin ligase MDM2 and inhibits its activity. MDM2 normally targets p53 to the proteasome for degradation, so free 5S RNP promotes stabilization and activation of p53 (*14*, *22*). Importantly, NS does not result in p53 stabilization and activation in the absence of 5S RNPs components, demonstrating that 5S RNPs play a critical role in the NS response (*14*, *22*). It appears that altering the balance in favor of 5S RNP integration into ribosomes will contribute to cancer development and therapeutic resistance (*25*). Thus, promoting the extra-ribosomal activity of free 5S RNPs in cancers with wild-type TP53 may improve the p53-dependent anticancer effects of the chemotherapy used in most poor-prognosis cancers. On the other hand, little is known about the putative existence of a free 5S RNP pool under basal conditions. Given that 5S rRNA is independently transcribed by RNA polymerase III instead of RNA polymerase I for the other rRNAs, a logical hypothesis would be that 5S rRNA production is not in complete stoichiometry with other ribosomal RNAs, even under basal conditions.

Many cancer treatments that are not designed to target ribosome synthesis induce a NS response, including chemotherapies (*14*, *16*, *26*). Even though ribosome synthesis is essential in all living cells, cancer cells are more sensitive to its inhibition. Indeed, ribosome synthesis is strongly increased in cancer cells, which require a greater synthesis of all its components including 5S RNPs. When this process is altered, a greater amount of free 5S accumulates, thus mediating a stronger p53 activation (*27*). This phenomenon partly explains why cancer cells are more sensitive to nucleolar stress than normal (*28*, *29*). Therefore, activation of the NS response by anticancer drugs such as 5-fluorouracil (5-FU) or doxorubicin contributes to their therapeutic benefit (*26*). However, as frequently observed for many cancer treatments, in clinical trials using new class of Pol I inhibitors, half of the patients showed progressive disease, indicating that resistance mechanisms may be occurring (*30*). But how cancer cells resist to NS in wild-type TP53 tumors patients remains to be understood.

In order to better understand how free 5S RNP can influence cell fate (ie, survival or apoptosis) under nucleolar stress conditions, we purified these particles to find partners that could regulate their activity. We identified the protein SURF2 as a faithful interactant of free 5S RNP particles. Our results demonstrate that SURF2 buffers the activity of for free-5S RNP particles in control cells to avoid unnecessary activation of p53. However, SURF2 counteracts p53 activation after NS, hence its expression is upregulated in some cancers and negatively correlates with overall survival. Our comprehensive dataset supports a model in which SURF2 competes with MDM2 for their association to free-5S RNPs, and their alternative binding thus modulates the response to nucleolar stress. Inhibition of SURF2 may be a promising therapeutic avenue to promote p53 activation in conjunction with treatments that activate nucleolar stress.

## Results

### SURF2 is a new binding partner of free 5S RNP particles

To identify potential regulators of free 5S RNPs, we set out to purify these particles outside the ribosomes. 5S particles are formed by the association of the 5S rRNA, with the ribosomal proteins RPL5 and RPL11, whether they are integrated into the ribosome or accumulate as independent particles. We therefore developed a novel U2OS cell line, known to respond well to NS, that inducibly overexpresses RPL5-Flag using Flip-In T-REx constructs (Fig S1). We verified the expression of RPL5-Flag, and used sucrose gradient fractionation to confirm that the fusion protein accumulates in both ribosomal and free fractions without altering ribosomal subunits production (Fig S1A and S1B). The process of ribosome assembly consists of the maturation of a large primary transcript that contains mature rRNA sequences (18S, 5.8S, and 28S) separated by spacers that are removed after exo and endo-nucleolytic steps. rRNA processing occurs inside pre-ribosomal particles constituted of pre-rRNAs associated to both ribosomal proteins and transiently associated factors, the latter being absent from mature ribosomal subunits (Fig S1C) (*5*, *31*). Disruption of ribosome assembly will lead to abnormal accumulation of rRNA precursors reduced rRNAs production (*32–34*). To confirm that ectopic expression of RPL5-Flag did not affect ribosome synthesis, we assessed pre-rRNA processing using northern blots (Fig S1D). No changes in accumulation of rRNA precursors nor mature forms were observed, indicating that RPL5-Flag expression does not perturb ribosome maturation.

To enrich partners of the free 5S RNP, ribosomes and pre-ribosomes were pelleted through a sucrose cushion. Free 5S RNP particles were purified from the supernatant, using RPL5-Flag and beads coupled with anti-Flag antibodies (Fig S1E). Interacting proteins were recovered, digested with trypsin and analyzed by nano-liquid chromatography-tandem mass spectrometry (nanoLC-MS/MS) using differential label-free quantitative proteomics (Fig 1). To evaluate interaction changes, pairwise comparison based on MS intensity values were performed for each quantified protein, first, between RPL5-Flag-expressing cells and control U2OS cells only expressing the Flag-tag (Fig 1A). Enriched proteins were selected based on their significant protein abundance variations between the two compared conditions (fold-change (FC) > 2 and < 0.5, and Student t-test p-value < 0.05) (see Mat&Meth for details). The volcano plot presented in Figure 1A shows that several proteins were significantly co-purified with RPL5-Flag, indicating them as potential partners of free-5S (Fig 1A and Table S1). RPL11, the other component of the 5S particles was found with a similar fold-change (FC = 33.34) compared to the RPL5-Flag bait (FC = 59.3), confirming the efficient purification of the 5S particles and that RPL5-Flag is mainly in complex with RPL11. No other ribosomal proteins were found specifically enriched in this purification, attesting the efficiency of the ribosomal fraction elimination step.

**Figure 1:**
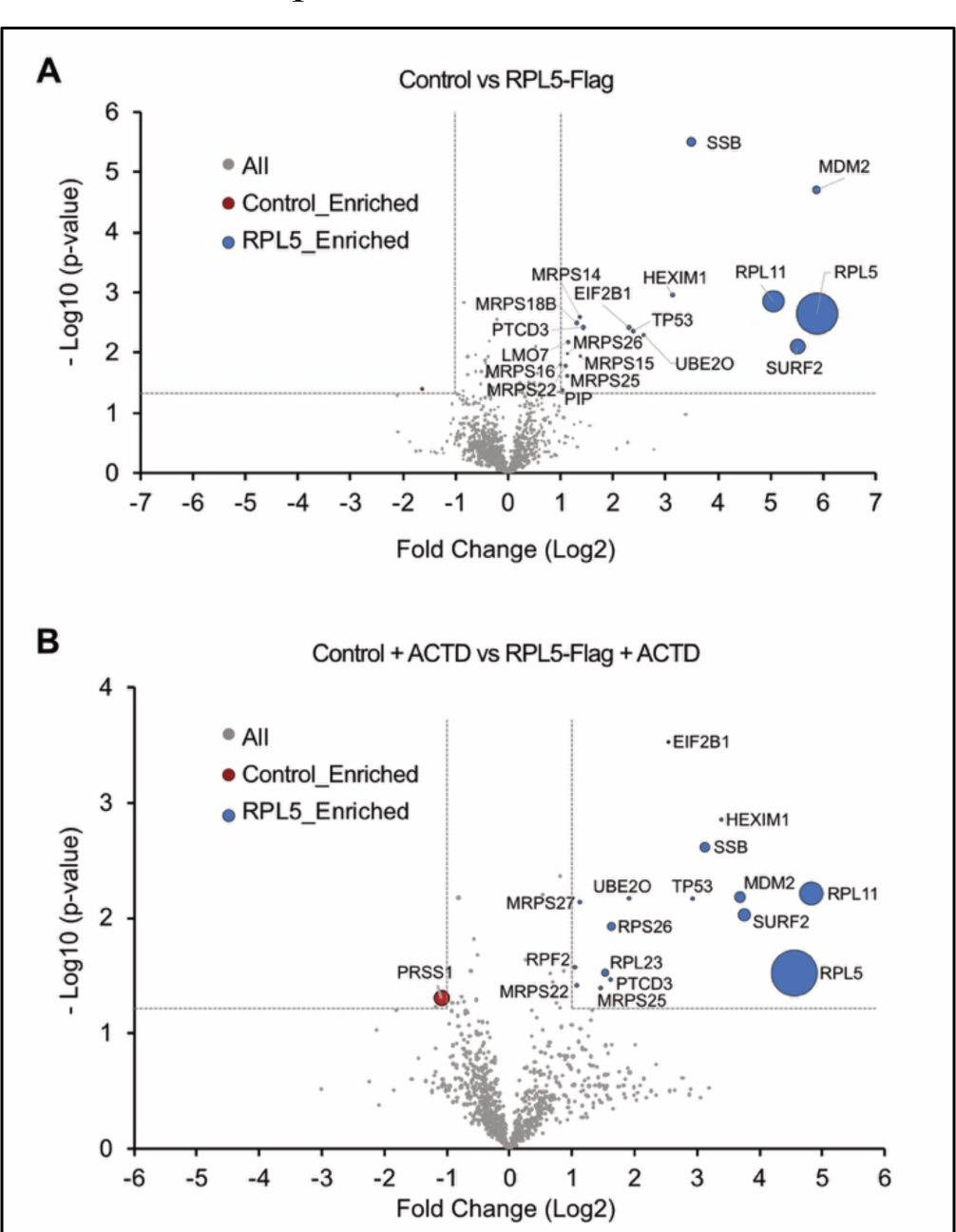
Label-free quantitative proteomics analysis of RPL5-FLAG co-purified proteins. Nano-liquid chromatography-tandem mass spectrometry (nanoLC-MS/MS) analysis of trypsin digested proteins retained on flag coated beads issued from U2OS control cells (LEFT quadrant) or cell expressing RPL5-FLAG (RIGHT quadrant). At least three independent experimental replicates were performed. Volcano plots showing proteins significantly enriched for control cells (red) versus cells expressing RPL5-FLAG (A) in untreated cells (B) in cells treated with actinomycin D (ACTD, 10 ng/ml) for 24 h. An unpaired bilateral Student t-test with equal variance was used. Enrichment significance thresholds are represented by an absolute log2-transformed fold-change (FC) greater than 1 and a -log10-transformed (p-value) greater than 1.3. The iBAQ (intensity-Based Absolute Quantification) values which are relevant to rank the absolute abundance of different proteins within a single sample (*35*) are represented by the diameter of each dot.

We also found MDM2 (FC = 58.01), already described as a major free-5S partner (*22*, *36–38*). La-protein, a known chaperone of newly synthesized 5S rRNA, is also enriched as well as HEATR3, a chaperone of RPL5 (*14*, *39–43*). We also found p53, which was not anticipated but might correspond to an indirect interaction via MDM2. We tested this hypothesis, by using Nutlin-3a, a p53 interaction inhibitor that targets MDM2 binding pocket (*44*)(Fig S2). During these experiments, we could reproduce our previous results showing an interaction between p53 and free-5S using IPs followed by western-blot analysis (Fig S2). However, this association is lost in presence of Nutlin-3a. This observation demonstrates that binding of MDM2 to free-5S and p53 are not mutually exclusive in U2OS cells.

HEXIM1 was also significantly enriched (FC= 8.75) during this experiment, but we could not reproduce this interaction using HEXIM1 as a bait during reverse IP experiment (data not shown). Of interest, SURF2, a previously uncharacterized protein, was significantly and highly enriched (FC= 45.69) during RPL5-Flag purification (Fig 1A). Other proteins such as mitochondrial ribosomal proteins (MRPS14, MRPS15, MRPS16, MRPS18B, MRPS22, MRPS25 and MRPS26); UBE2O (a ubiquitin-ligase proteins) and EIF2B1 (translation initiation factor) were also specifically found albeit with reduced fold-changes compared to other partners. To investigate how the panel of free 5S RNP binding partners changes during stress, we used the same purification strategy but this time after inducing NS with a low dose of actinomycin D (ACTD) (10 ng/mL for 24 h) (Fig 1B). Unexpectedly, the repertoire of potential free RPL5-flag partners showed few changes, with only increased enrichment of MDM2 (Fig 1B and Table S2).

Among RPL5-Flag partners, SURF2 stood out, both as an abundant (areas of circles is proportional to the IBAQ abundance of each protein) and strongly enriched (FC = 45.69 and 14.42, before or following NS induction respectively). Indeed, it is the second most enriched partner of RPL5-Flag in both normal and NS conditions. SURF2 (Surfeit Locus 2) shares a c-Myc responsive promoter with SURF1, located in a crowded and conserved region (*45–47*). Surprisingly, we found almost no functional data for SURF2. However, the genomic region contains other genes encoding for factors involved in ribosome production (SURF3/RPL7A and SURF6/RRP14), a transmembrane receptor involved in endoplasmic reticulum export (SURF4), a component of the mediator complex (SURF5/MED22) and a factor involved in cytochrome-c synthesis (SURF1/SHY1), all of which promote cell growth and proliferation. To understand what role SURF2 might play in relation to free 5S particles, we undertook its functional characterization in U2OS cells.

### SURF2 binds 5S RNP particles but is not involved in ribosome assembly

Free 5S RNP particles are normally incorporated into ribosomes, so we tested whether SURF2 was involved in this process. SURF2 is described in the human protein atlas as a nucleoplasmic and nucleolar component suggesting a potential role in ribosome assembly (https://www.proteinatlas.org/ENSG00000148291-SURF2). SURF2 was purified from U2OS whole cell extract by immunoprecipitation using anti-SURF2 antibodies, with anti-GAPDH as control. Protein partners were identified by a differential label-free quantitative proteomics approach and shown in a volcano plot (Fig 2A). As expected, RPL5 and RPL11 were significantly enriched with SURF2 (Fig 2A and 2B). Among other factors identified, TRIM21 and PRDX1 are common contaminants from U2OS extracts (see CRAPome database) (*48*), while DLST (dihydrolipoamide S-succinyltransferase) is mitochondrial, and these were not considered further. RPF2 and RRS1, which mediate proper integration of 5S RNP particles to pre-60S pre-ribosomal subunits, were identified (*43*, *49–51*). Their low abundance could reflect a transient interaction of these proteins with free 5S RNP particles before being integrated in ribosomes (Fig 2B).

**Figure 2:**
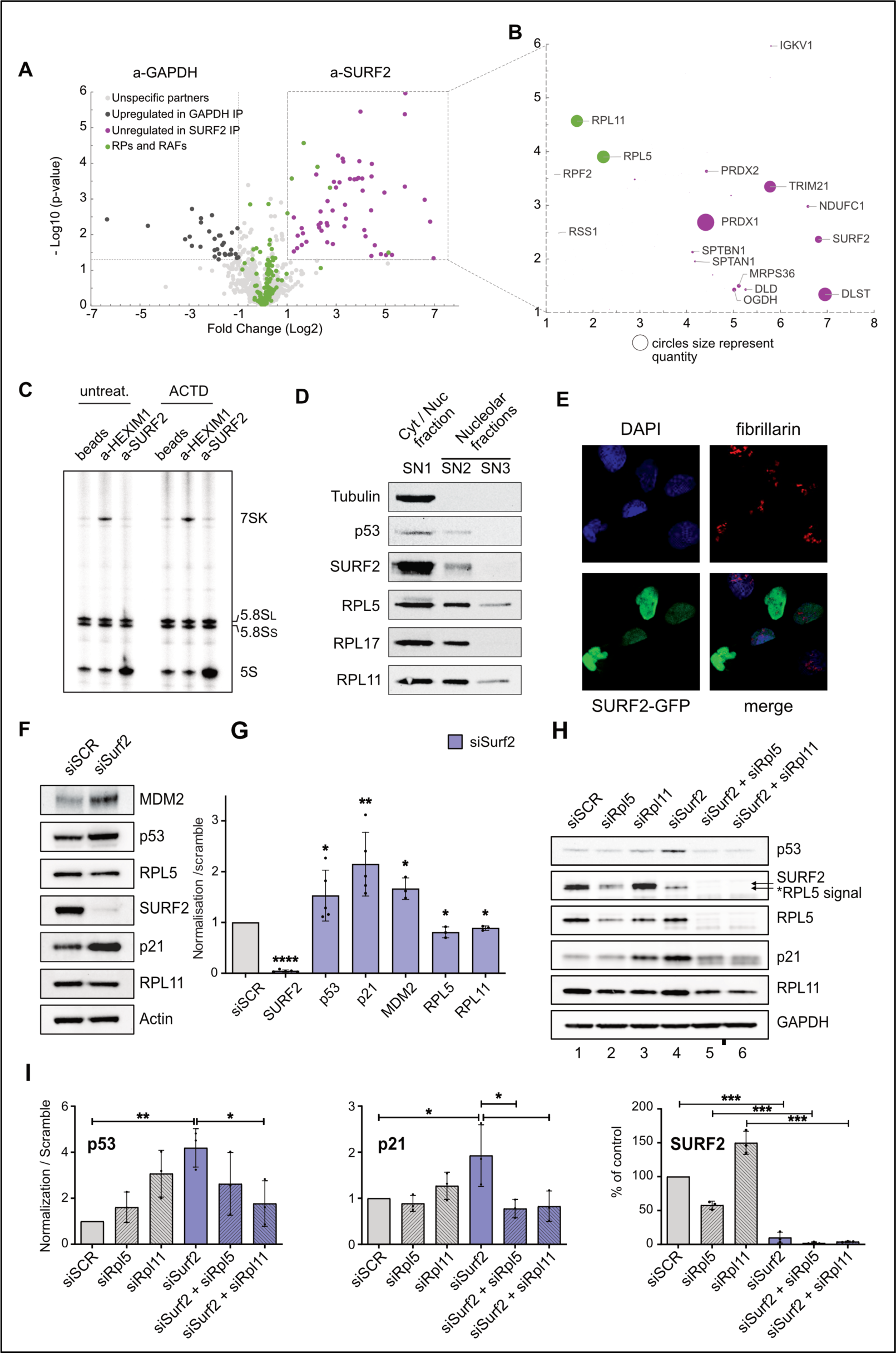
SURF2 is a new partner of free-5S particles and is involved in p53 regulation. ((A) Label-free quantitative proteomics analysis of trypsin digested proteins retained on beads coated by anti-GAPDH or anti-SURF2 antibodies. At least three independent experimental replicates were performed. (A) Volcano plot showing proteins significantly enriched in the GAPDH IP (brown) versus the SURF2 IP (green). An unpaired bilateral Student t-test with equal variance was used. Enrichment significance thresholds are represented by an absolute log2-transformed fold-change (FC) greater than 1 and a -log10-transformed (p-value) greater than 1.3. Proteins involved in ribosome biogenesis and function are indicated in purple. (B) Focus on the iBAQ values, represented by the diameter of each dot, for the proteins significantly enriched in the SURF2 IP. (C) Detection of RNAs associated with HEXIM1 and SURF2. U2OS RNAs co-immunoprecipitated with endogenous HEXIM1 (a-HEXIM1) or endogenous SURF2 (a-SURF2) were 3’-labelled and separated on a 6% acrylamide gel. Control IP reaction performed without antibodies (beads) is shown. (D) U2OS cell extracts were fractionated using the PSE method. Western-blot analyses showing the contents of different factors (indicated at the left of the gel) in the SN1, SN2 and SN3 fractions obtained with the PSE method. (E) Microscopy analyses of U2OS cells over-expressing SURF2-GFP from TET inducible promoter. Cells were induced with tetracycline to obtain a similar expression as the endogenous copy (tetracycline at 5 ng/mL) and SURF2-GFP signal detected (Green). The same cells were also probed for fibrillarin to stain nucleoli. DAPI coloration is used to localize nuclear compartment. (F) Western-blot analyses showing the accumulation of different proteins in 20 µg of cellular extracts produced from U2OS cells treated with scrambled siRNAs (siSCR) or with siRNAs directed against SURF2 (siSurf2). (G) Quantification of signals obtained by western-blot analysis and normalized to actin signals in each condition (f) (n=3). t-test analysis was used for statistics. Significant differences are indicated by stars (*p-value* ≤0.05*; ≤0.01** ; ≤0.001*** and ≤0.0001****).(H) Co-depletion experiment. Western-blot analyses showing the accumulation of different proteins in 20 µg of cellular extracts issued from U2OS cells treated with a combination of different siRNAs: scrambled (siSCR); RPL5 (siRPL5); RPL11 (siRPL11) or SURF2 (siSurf2). (I) Quantification of signals obtained by western-blot analysis and normalized to GAPDH signals in each condition (h) (n=3). Anova test was used for statistics. Significant differences are indicated by stars (*p-value* ≤0.05*; ≤0.01** and ≤0.001***).

To test whether SURF2 interaction with RPL5 and RPL11 occurs inside or outside of free 5S RNP particles, we tested the interaction of SURF2 with RNAs in the cell by performing IP using either beads coupled to anti-SURF2, anti-HEXIM1 (HEXIM1 being found associated with RPL5-Flag, see Fig1 above) or beads not coupled with antibodies (termed “beads” in Fig 2) as control experiment. RNAs enriched during the IPs in the different conditions were labelled with P^32^ using pCp labelling and separated on a polyacrylamide gel (Fig 2C). HEXIM1 specifically enriched its known partner 7SK, but no other RNAs in normal or NS (actinomycin D treatment) conditions. In contrast, SURF2 retained significantly more 5S rRNA than control beads, indicating that SURF2 specifically interacts with 5S rRNA in addition to RPL5 and RPL11. To confirm that SURF2 only interacts with 5S RNP particles in their free form, we performed a cell fractionation analysis using the recently developed Pre-ribosome Sequential Extraction (PSE) method (*52*) (see material and methods) (Fig 2D). Localization of SURF2 in different cell fractions (cytoplasmic/nuclear and nucleolar fractions) was tested by western-blots and compared to other proteins. SURF2 was preferentially enriched in the cytoplasmic/nuclear fraction as observed for p53, while RPL5, RPL11 and RPL17 are equivalently present in nucleolar fractions. This indicates that SURF2 preferentially localizes outside the nucleoli. To confirm this conclusion, we localized SURF2 using microscopy. Despite being efficient for western-blot and IPs, anti-SURF2 antibodies did not work for immuno-fluorescence experiments in our hands (data not shown). Therefore, we developed a new U2OS cell line expressing a SURF2-eGFP tagged version in the presence of tetracycline. When expressed in the same range as the endogenous one, SURF2-eGFP localized mainly to the nucleoplasm was also more weakly detected in the nucleolus, supporting the PSE result (Fig 2E).

To formally exclude a role of SURF2 in ribosome assembly, we assessed whether SURF2 depletion using siRNAs influences ribosome assembly in U2OS cells. We confirmed an 85% reduction in SURF2 protein accumulation by western-blot analysis after 96 hours of treatment by two successive transfections with siRNAs (Fig 2F and 2G). As a consequence, we kept this treatment through the remaining part of this study. After SURF2 depletion, we analyzed pre-rRNAs maturation process by northern-blots (Fig S3A and S3B). Data were quantified by Ratio Analysis of Multiple precursors (RAMP), which detects alterations of rRNA maturation by displaying the ratios between one pre-RNA and its immediate processing product (*34*, *53*). No obvious changes in the accumulation of mature rRNAs, nor of their precursors were observed after SURF2 depletion. The combined analysis of three northern blots showed no significant difference in the relative levels of the 32S, 30S, 21S, 12S or 18S-E precursors (Fig S3B). This absence of pre-rRNAs processing defects was confirmed by sucrose gradient analyses, showing no depletion of the small or large ribosomal subunit or other changes (Fig S3C). SURF2 was found in the lighter fractions of the gradient, supporting association with 5S RNPs but not (pre-) ribosomal particles. The association of RPL5 and RPL11 with ribosomal particles was tested by sucrose gradients and western-blot analysis, compared to RPL17 (Fig S3D and S3E). Their association was unaffected by SURF2 depletion. Altogether, our data support the conclusion that SURF2 does not intervene in ribosome assembly, and interacts only with the free form of 5S RNP particles.

### SURF2 depletion promotes p53 activation independent of NS induction

Free 5S RNP induces NS response by promoting p53 stabilization and expression of its target gene p21/CDKN1A, which in turn triggers G1 arrest and/or apoptosis. Using siRNAs and western-blots, we investigated whether SURF2 depletion promotes p53 activation in the absence of NS exposure by western blots. As a control, we used cells treated with scrambled siRNAs. After SURF2 depletion alone in U2OS cells, we observed increased protein levels for p53, p21 and MDM2 compared to control conditions (Fig 2F and 2G).

To determine whether the activation of p53 induced by SURF2 depletion was related to or independent from free 5S RNP function, we repeated SURF2 depletion in the absence of RPL5 or RPL11, conditions known to abolish p53 stabilization (*22*) (Fig 2H and 2I). Following the treatments with various combination of siRNAs, accumulation of NS-related proteins involved in p53 regulation was examined by western-blot (Fig 2H). As previously reported, depletion of either RPL5 or RPL11 reduced the level of the other (*24*) (Fig 2H, lane 5 and 6 compare to 2 and 3, Fig S3). Strikingly, this was also found for SURF2; depletion of RPL5, RPL11 or SURF2 proteins affects the stability of the two other members of this group, strongly supporting direct interactions (Fig 2H, 2I and S4). After U2OS cells treatment with siRPL5 or siRPL11, mild accumulations of p53 and p21 were observed that could reflect a bit of NS response before proteins depletion really affect free 5S RNPs accumulation (Fig 2H and 2I). A greater accumulation of p53 was observed after RPL11 depletion, but without correlated p21 induction. This confirms previous results showing that although RPL5 and RPL11 depletion hinders ribosome assembly, their depletion impedes p53 and p21 accumulation by free-5S particles (*14*, *22*). In this new set of experiments, we were able to confirm by western-blots that SURF2 depletion significantly promotes both p53 and p21 accumulation (Fig 2H and 2I). However, when SURF2 is co-depleted with RPL5 or with RPL11 we could no longer observe any p21 accumulation and a strong reduction in p53 stabilization observed when we only deplete for SURF2 (Fig 2H, lanes 5 and 6 and 2I). This result indicates that SURF2 depletion effect on p53 activation is mediated by free-5S particles, independently of NS induction.

### SURF2 depletion increases MDM2 binding to free 5S RNP particles

As previously described, free-5S RNP promote p53 activation through sequestration of MDM2, its primary negative regulator. To decipher how SURF2 might regulate free-5S RNP function, we investigated whether and how SURF2 was related to MDM2. While studying SURF2 interactome, we could not find any interaction of SURF2 with MDM2 nor with p53 (Fig 2A, Table S3). This data indicates that the regulation of free 5S RNP function might not occur through a direct interaction between SURF2 and MDM2. We thus then tested how SURF2 might affect the binding of free 5S RNP particles to MDM2. First, we performed IPs of free 5S RNP particles in the presence or absence of SURF2, using RPL5-Flag as bait. Since the interaction between free-5S RNP and MDM2 is enhanced after NS, we performed these IPs before or after NS exposure by treating U2OS cells with low dose of actinomycin D (ACTD, as for Fig 1).

We used U2OS that do not express RPL5-Flag as a control (Figure 3A). As expected, only a mild but specific retention of MDM2 and p53 is observed in cells expressing RPL5-Flag under normal conditions. Importantly, no interaction between RPL5-Flag and p21 was identified, confirming that p21 accumulation only reflects p53 transcriptional activity. Under the same conditions, depletion of SURF2 specifically increased interactions of both MDM2 and p53 with RPL5-Flag (Fig 3A and 3B). Following NS induction by actinomycin D, interactions of both MDM2 and p53 with RPL5-Flag were also increased (Fig 3A and 3B). Remarkably, an additional and statistically-significant increase of p53 retention with RPL5-Flag is observed when cells are both exposed to NS and depleted of SURF2 (Fig 3A and 3B). A similar trend was also observed for MDM2, but the variability of signal quantification between experiments was too high to reach statistical significance. These data indicate that SURF2 depletion promotes both MDM2 and p53 binding to free 5S RNP particles under normal conditions or upon nucleolar stress.

**Figure 3:**
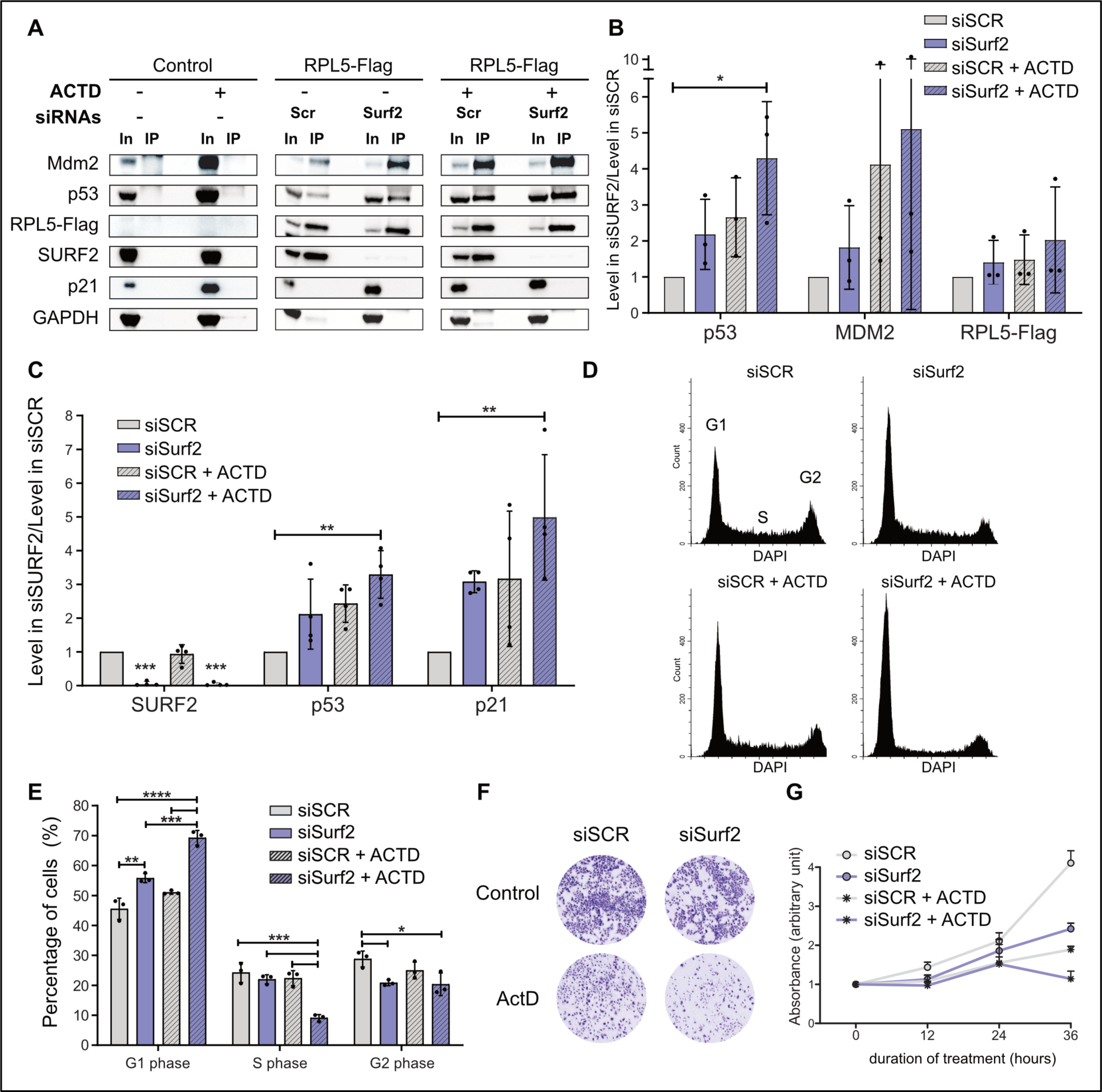
SURF2 depletion increases free-5S binding to MDM2 and increase cell sensitivity to nucleolar stress. (A) Detection of proteins associated with RPL5-Flag. Cell extracts produced from U2OS cells overexpressing RPL5-FLAG or not (control) and differentially treated by actinomycin D addition for 24 h (at 10 ng/mL) or not (ACTD + or -) and transfected with scrambled siRNAs (siSCR) or directed against SURF2 (siSurf2) were used to perform immuno-precipitation on beads coated with anti-flag. 10% of the inputs were loaded (In) aside IPs (IP) on gel to perform western-blots using antibodies directed against the indicated proteins to analyze co-purification efficiency of the different factors. (B) Quantification of the co-purification efficiency with RPL5-Flag observed (A) for the indicated proteins (n=3). Results are represented as a comparison of the enrichments observed in cells treated with siRNAs directed against SURF2 (siSurf2) normalized to the ones observed in cell treated with scrambled siRNAs (siSCR). t-test analysis was used for statistics. Significant differences are indicated by stars (*p-value* ≤0.05*; ≤0.01** and ≤0.001***). (C) Quantification of the signals corresponding to the indicated proteins in the different inputs of panel (A) n=3. t-test analysis was used for statistics. Significant differences are indicated by stars (*p-value* ≤0.05*). (d) DNA content analysis of U2OS cells treated as indicated (siSCR: scrambled siRNAs; siSurf2: siRNAs against SURF2; ACTD: actinomycin D at 10 ng/mL for 24 h) by FACS. (E) Quantification of U2OS cells repartition in the different phase of cell cycle following different treatments (n=3). Anova Test was used for statistics. Significant differences are indicated by stars (*p-value* ≤0.05*; ≤0.01** and ≤0.001***). (F and G) Proliferation analysis of differentially treated U2OS cells. Cells are treated as indicated (siSCR: scrambled siRNAs; siSurf2: siRNAs against SURF2; ACTD: actinomycin D at 10 ng/mL for 24 h). (F) Cells are platted on 6 well plate after being transfected with the indicated siRNAs, 24 h before being analyzed cells were treated with actinomycin D (10 ng/mL) or H2O. Cells were stained by crystal violet to take picture. (G) Fixed crystal violet was resolubilized and quantified by absorbance for each condition after different time exposure to treatments.

### SURF2 depletion increases cells sensitivity to NS

To test whether SURF2 depletion modulates free-5S RNP function in NS response, we reanalyzed the levels of p53, MDM2 and p21 in our previous IPs (Fig 3A). This showed that p53, p21 and MDM2 were all stabilized both after NS exposure (ACTD) or following SURF2 depletion. Notably, depleting SURF2 in cells exposed to actinomycin D promoted a statistically significant increase in the accumulation of p53 and p21 compared to each condition independently (Figure 3C). To confirm that SURF2 depletion increases the response to NS, we analyzed the effect on cell cycle progression by flow-cytometry analysis (Fig 3D and 3E). As expected, exposure of cells to actinomycin D induced an accumulation of cells in G1 phase, reflecting cell-cycle arrest. This arrest was also observed following SURF2 depletion alone (Fig 3D). Interestingly, inducing NS in cells depleted of SURF2 promoted an even stronger and more statistically significant G1 arrest (Fig 3D and 3E). We also performed cell proliferation assays on U2OS cells, either depleted of SURF2 with siRNAs, exposed to Actinomycin D or receiving both treatments (Fig 3F and 3G). Cell proliferation was assessed in triplicate, using crystal violet staining, at timepoints up to 36h. Treatment with anti-SURF2 siRNAs or actinomycin D reduced cell proliferation compared to scramble siRNAs (siSCR). Strikingly, the combination of both treatments had the strongest effect on cell proliferation, and was accompanied by a loss of cellular material between 24 and 36 h of treatment (Fig 3G). We propose that the combination of both treatments promotes apoptosis, in addition to slowing proliferation, which was not obvious for single treatments.

To confirm the generality of this conclusion, we repeated these experiments using another cancer cell line; Hepatocellular carcinoma (HCC) HepG2 cell line carrying a wild-type TP53 gene. In HCC cells, SURF2 depletion promoted p53 activation and cell cycle arrest (Figure S5A, S5B and S5C). The combination of low dose of actinomycin D treatment and SURF2 depletion further decreased cell proliferation and also induced cell death (sub-G1), which was not observed in cells exposed only to SURF2 depletion or actinomycin D (Fig S5C and S5D).

### SURF2 overexpression impedes p53 activation following NS

As SURF2 depletion increases cell response to NS, its overexpression could inversely, inhibit cell response to NS. To test this hypothesis, we developed a new U2OS Flip-In T-rex cell line with a tetracycline-induced overexpression of SURF2-Flag. To avoid misinterpretation due to side effects of SURF2 overexpression, we verified that this induction did not alter ribosome assembly (Fig S6A and S6B). We then compared by western-blot analysis the level of p53 and p21 in control cells treated or not with actinomycin D, with or without overexpression of SURF2-flag for 24 hours (Fig 4A). Quantification of these signals confirmed that cells exposure to actinomycin D promotes p53 stabilization and activity, as reflected by p21 increased levels (Fig 4A). In stark contrast, cells overexpressing SURF2 showed only a mild increase in p53 and p21 levels (Fig 4A and 4B). These data indicate that overexpression of SURF2-Flag impedes p53 activation by free 5S RNP. To confirm this observation, we performed cell cycle analysis by flow cytometry using the same cells and treatments, and quantified the percentage of cells in each phase (Fig 4C). Exposure to actinomycin D promoted G1 arrest in untreated cells, but overexpression of SURF2-Flag completely blocks this response (Fig 4C). This striking result indicates that overexpression of SURF2 inhibits NS response in U2OS cells in this time frame.

**Figure 4:**
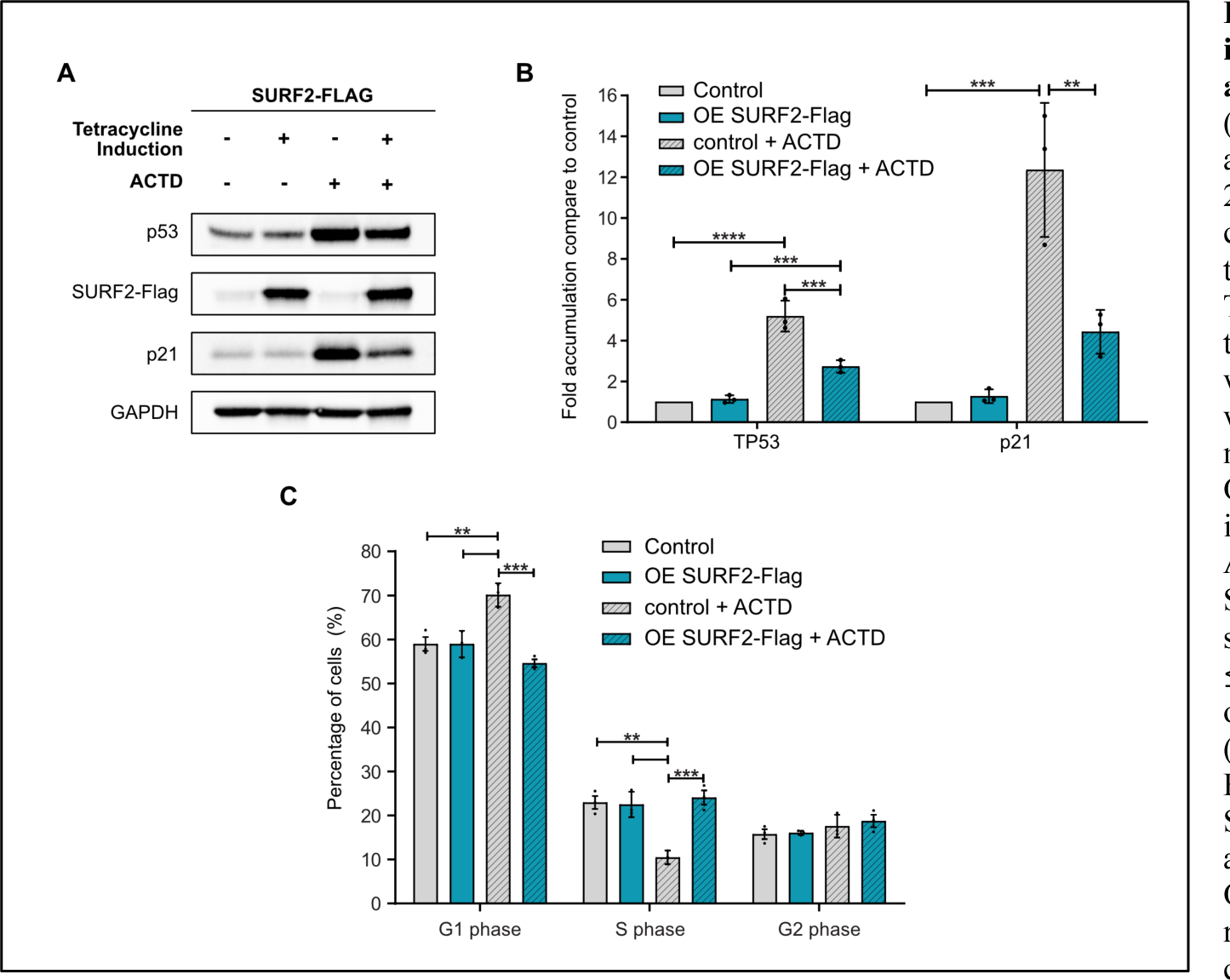
SURF2 overexpression impedes p53 activation and cell cycle arrest following nucleolar stress. (A) Western-blot analyses showing the accumulation of different proteins, in 20ug of cellular extracts produced from control U2OS cells or from U2OS cells that overexpress SURF2-Flag after Tetracycline induction (1 µg/ml) from their T-Rex locus for 24 hours. Cells were also treated differentially with or without addition of actinomycin D (10 ng/ml) for the same period. (B) Quantification of the signals observed in independent experiment (A) (n=3). Anova Tests were used for statistics. Significant differences are indicated by stars (*p-value* ≤0.05*; ≤0.01** and ≤0.001***). (C) DNA content analysis of U2OS cells treated as indicated (Ctrl: U2OS control cells; OE SURF2-Flag: U2OS cells that overexpress SURF2-Flag; ACTD: Treated with actinomycin D) by FACS. Quantification of different U2OS cells repartition in the different phase of cell cycle following different treatments is represented as histograms (n=3). Anova Tests were used for statistics. Significant differences are indicated by stars (*p-value* ≤0.05*; ≤0.01** and ≤0.001***).

### SURF2 abundance correlates with poor overall survival in some cancers

In cancer, NS stress can participate in both tumor development and drug response. We therefore sought to determine whether SURF2 expression level correlates with cancer patient outcome. We analyzed reported SURF2 expression in a PanCancer patient cohort, using the TGCA resource (https://portal.gdc.cancer.gov), comparing tumor SURF2 levels relative to their associated healthy tissues (GTex datasets) (Fig 5A).

**Figure 5:**
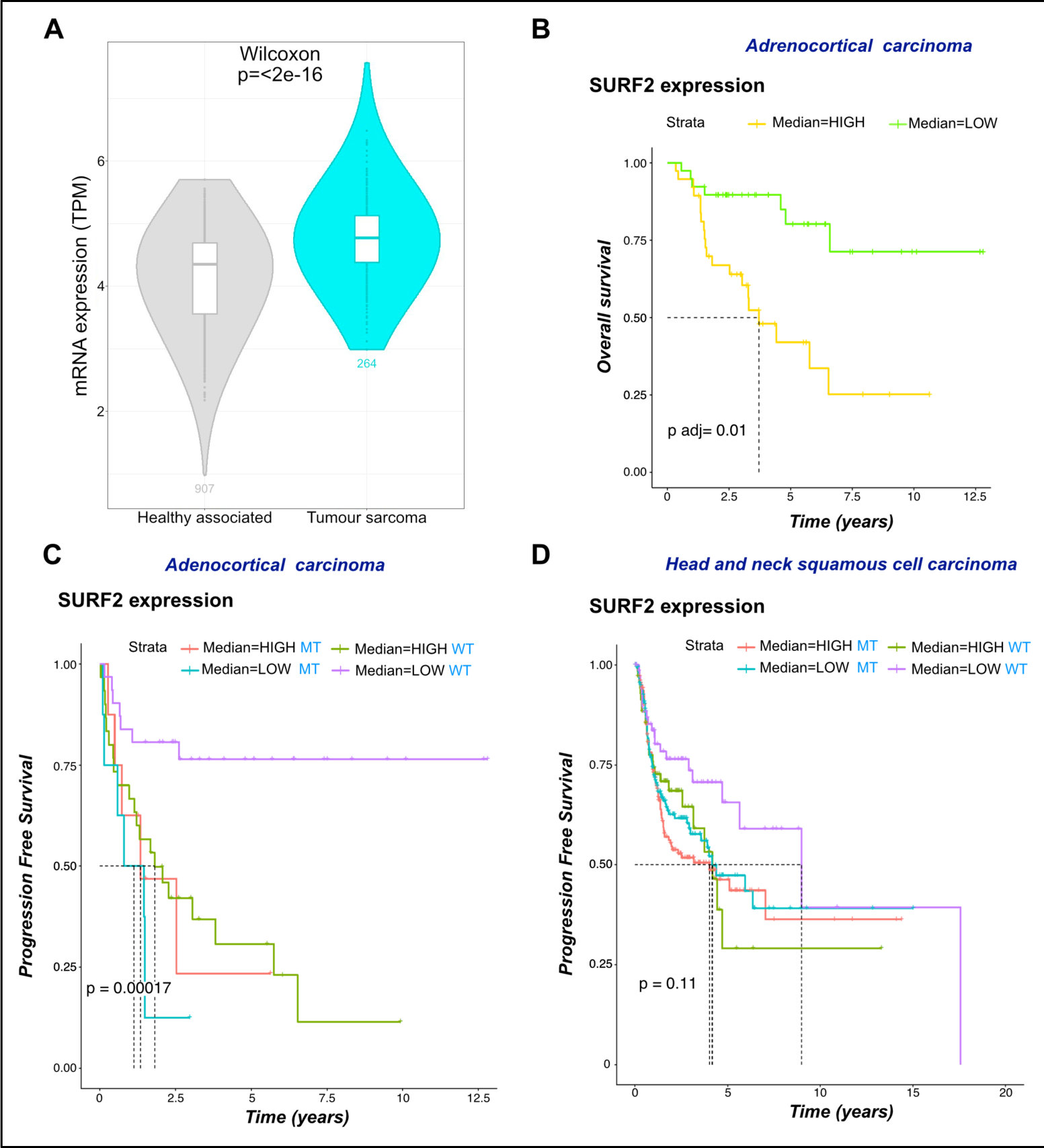
Surf2 expression associates with patient outcomes. (A) SURF2 overexpression in cancer. Surf2 mRNA levels was compared between tumoral tissues (TCGA dataset) and their relative healthy counterparts (GTex dataset) using normalized Xena data. An example is given showing that tumor tissues express higher Surf2 mRNA level compared to their relative healthy tissues with adjusted *p-value* (Wilcoxon test). (B-D) Association between SURF2 expression and cancer patient survival. Kaplan-Meier curves were performed to compare overall survival depending on Surf2 mRNA levels and examples are shown for adrenocortical carcinoma (B), head and neck carcinoma (C) and hepatocellular carcinoma (D), in the whole population (B) or depending on TP53 mutation status (C-D).

In a vast majority of evaluated cancers, we found an increased level of SURF2 mRNA compared to healthy tissue (*p-value* < to 2^e-16^), indicating that SURF2 is over-expressed in several cancer types. We then analyzed the association between SURF2 mRNA expression and the overall or progression-free survival across cancers (Fig 5B, 5C and 5D). In adrenocortical carcinoma, a pediatric cancer, high level of SURF2 expression significantly correlates with poor overall survival compared to low expression of SURF2 (25% at 6 years, p adj value= 0.01) (Fig 5B). Interestingly, this difference and trend is conserved in progression-free survival in wild-type (WT), but not in mutant (MT) TP53 tumors, suggesting that in WT TP53 adrenocortical carcinoma, SURF2 expression would contribute to patient survival (Fig 5C). A similar trend was also observed in head and neck carcinomas, in which a better progression free survival correlates with lower SURF2 expression in WT TP53 tumors (Fig 5D). This is coherent with our previous biochemical data, that show opposing levels of SURF2 and p53 accumulation, and supports the model for antagonist roles of SURF2 and p53 in response to NS.

### Over-expressed SURF2 impedes MDM2 binding to 5S RNP in vivo

We directly tested whether SURF2 regulates MDM2 binding to free-5S RNP particles. We assessed free 5S RNP particle binding, by recovery of RPL5 and RPL11 on beads coupled with either anti-MDM2 or anti-SURF2 antibodies, with beads devoid of antibodies as control (Fig 6A). 5S RNP association was tested following exposure to NS, in control cells or cells overexpressing SURF2-Flag from the U2OS-T-rex locus. In control cells (not overexpressing SURF2-Flag), we clearly observed specific retention of RPL5 and RPL11 with anti-MDM2 or anti-SURF2. This demonstrates that free-5S RNP particles can interact with these two factors. Notably, SURF2 was not retained by anti-MDM2, nor MDM2 by anti-SURF2, consistent with our interactome analysis (Figure 2A and B). In cells overexpressing SURF2-Flag, RPL5 and RPL11 were enriched in total cell extracts (see inputs Fig 6A), potentially reflecting their stabilization in a complex formed by SURF2-RPL5-RPL11-5S rRNA. Despite this, we repeatedly (n=4) failed to detect any signal for RPL5 or RPL11 in the IPs using anti-MDM2 coupled beads in cells overexpressing SURF2-Flag.

**Figure 6:**
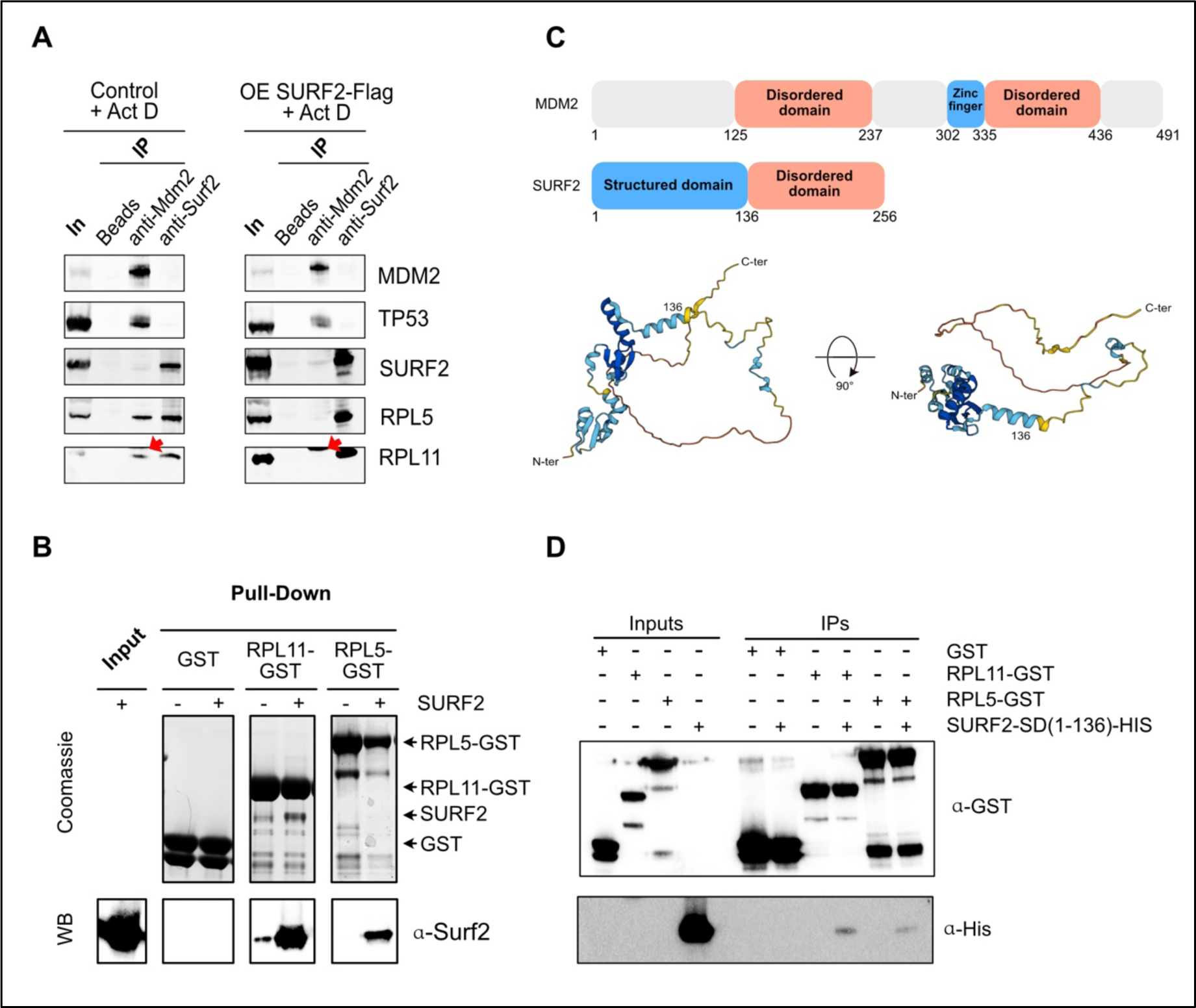
SURF2 is able to directly interact with both RPL5 and RPL11 and competes for their binding with MDM2 in vivo. (A) Extracts from U2OS cells that overexpress SURF2 (OE SURF2-Flag) or not (control) and treated by actinomycin D (ACTD) are used to perform IPs using beads only, beads coupled to anti-SURF2 or beads coupled to anti-MDM2. After washes, remaining proteins are resuspended in loading dye and analyzed by western-blots using the indicated antibodies. (B) GST-pulldown assays. Extracts of BL21 that overexpress recombinant SURF2-HIS were mixed with extracts that overexpress either GST alone, GST-RPL5 or GST-RPL11. Proteins specifically retained on glutatione-sepharose beads were analyzed both by Coomassie staining and western-blot analysis (WB). 10% of the extracts were used for inputs. (C) Secondary structure of SURF2 and MDM2 proteins. Functional domains are indicated. SURF2 3D structure modelisation by Alphafold software is represented (D) Same experiments as in (B) but replacing extract with SURF2-HIS by extracts that overexpress structural domain of SURF2 (SURF2-SD-HIS) as prey. Proteins specifically retained on glutatione-sepharose beads were analyzed both by western-blots with indicated antibodies.

We conclude that over-expression of SURF2 impedes free-5S RNP binding to MDM2 following treatment with actinomycin D.

### SURF2 interacts directly with RPL5 and RPL11 *in vitro*

Following NS, MDM2 binds to free-5S particles mainly through a direct interaction with RPL11, mediated by its zinc-finger domain (*37*, *43*). A mutation in this domain (MDM2C305F) fully abolishes its interaction with free-5S particles (*54*). The previous results suggested that SURF2 and MDM2 association with free-5S RNP might be mutually exclusive. To test this hypothesis, we performed pull-downs using recombinant RPL5-GST, RPL11-GST, or GST alone purified from *E. coli* (Figure 6B). The purified proteins were incubated with the same amount of recombinant SURF2-HIS and performed GST pull-down assays (Fig 6B). We observed weak but specific retention of recombinant SURF2 on beads coated with RPL5-GST or RPL11-GST but not with GST alone, both by Coomassie staining and western-blot. Comparison of SURF2-HIS signals indicated that RPL11-GST retains more SURF2 than does RPL5-GST, although association was seen for both proteins (Fig 6B). Like MDM2, SURF2 contains an intrinsically disordered domain (Fig 6C). To decipher by which domain SURF2 interacts with RPL5 and/or RPL11 generated truncated SURF2_1-136_-GST, corresponding to its structured domain (here after termed SURF2-SD) (Fig 6C and 6D). Due to intrinsic instability, we could not test the interaction of SURF2 disordered domains using similar experiments. SURF2-SD was bound by both RPL11-GST and RPL5-GST, with greater binding to RPL11 (Fig 6C and 6D). Preferential binding to RPL11, by both the structured domain of SURF2 and the disordered domain of MDM2, suggests a mechanism for their competitive binding to free-5S RNPs.

## Discussion

Free-5S RP particles are key in the response to nucleolar stress, a cellular mechanism that is instrumental to cancer treatment by chemotherapeutic drugs. Thus, promoting the extra-ribosomal activity of free 5S RNPs in wild-type TP53 cancers may improve the p53-dependent anticancer effects of therapeutic agents such as chemotherapy used in most poor-prognosis cancers. In order to understand how the function of these particles could be regulated at basal level and under nucleolar stress conditions, we undertook the characterization of these particles in U2OS cells. We retrieved well-known partners of free 5S RNP particles, such as SSB/La and MDM2 (Fig 1A). SSB/La, a chaperone of 5S rRNA, binds this RNA just after its transcription (*39*). The specific enrichment of SSB/La protein with RPL5-Flag suggests that during 5S particles biogenesis there is a transient intermediate containing both La and RPL5. Other known partners such as HEATR3 and BXDC1/RPF2 were also found associated to RPL5-Flag but with a statistic significance below our threshold (-log10(p-value)< 1) of 0.974 and 0.929 reciprocally (Table S1). In addition, we identified the tumor suppressor p53 as a partner of free 5S RNP particles, and were able to show that this interaction is mediated through MDM2 binding. So, how free 5S RNP particles fully inhibit MDM2 catalytic activity and promote p53 function remains elusive and will need further examination. Interestingly, the fact that we can purify free 5S RNP particles even in absence of stress demonstrates that a portion of these particles are not incorporated into ribosomes even in proliferative cell state. This over production of free 5S RNP particles observed in our experiments even in absence of stress may find its origin in the genomic and transcriptional independence of 5S rRNA compared to other rRNAs, which most likely fails to warrant a fully equilibrated production of rRNAs. Interestingly, 5S rRNA both genomic and transcriptional independence of 5S rRNA have been reinforced during evolution, going from an extra-copy of 5S rDNA in some bacteria such as *E. coli*, to a transcriptional independence in yeast *S. cerevisiae* and segregation of rDNA and 5S genes on different chromosomes in Humans. Therefore, differential production of 5S rRNA compared to the other rRNAs seems to give an advantage to the cells, but how this occurs remains an open question.

In addition to these factors, we found a potential new partner and precedingly uncharacterized factor, SURF2 (Figure 1A). SURF2 is a member of the surfeit genomic locus that contains 6 genes, all involved in promoting cell proliferation. Furthermore, it shares a bidirectional promoter with SURF1 that is positively regulated by the oncogene c-Myc, which highly supports cell proliferation (*47*, *55*). These data indicate this factor as a potential regulator of free 5S RNP particles. In the course of its characterization, we were able to show that this mainly nucleoplasmic factor is not directly involved in pre-ribosomes processing and only interacts with 5S RNP particles independently of ribosomes in their free state (Fig 2). We then showed (using both U2OS and HepG2 cell lines) that in absence of SURF2, cells accumulate p53 and p21 in a free-5S RNP dependent manner, with or without inducing nucleolar stress (Fig 2H, 2I and Fig 3A, 3C). Moreover, in response to nucleolar stress, induced by actinomycin D, cells depleted of SURF2 showed a greater accumulation of p53 and p21 and a stronger arrest in G1 cell cycle with associated cell death compared to control cells (Fig 3E, 3F and 3G). This greater cell sensitivity to nucleolar stress in absence of SURF2 is associated with an increased ratio of free 5S RNP particles associated with MDM2 and p53 (Fig 3A and 3B). Reciprocally, an over-expression of SURF2 impedes p53 activation and cell cycle arrest which are normally induced by NS, and abolishes the interaction between free 5S RNP particles and MDM2 (Fig 4 and 6A). This strong correlation between NS response and SURF2 level is also supported by PanCancer analysis performed using TCGA data (Fig 5). We found that high levels of SURF2 associate with poor overall survival in specific cancers such as head and neck squamous cell carcinomas, and adrenocortical carcinomas, and allow to distinguish the outcome of patients displaying WT TP53 tumors. This positions SURF2 both as a good anti-cancer therapeutic target and a prognostic marker, at least in these cancer types. Our results further show that the interaction of SURF2 or MDM2 with free-5S particles is mutually exclusive *in cellulo* and that SURF2 mainly interacts with free-5S particles through binding to RPL11, which involves SURF2’s structural domain (Fig 6).

All these observations support a functional model presented in Fig 7. In normal conditions (ie, without ribosomal stress), SURF2 acts as a buffer for free 5S RNP particles that could accumulate in excess compared to other ribosomal components even in absence of NS (Fig 7A). In response to NS, due to chemotherapeutic drugs for example, a greater portion of free 5S RNP particles accumulates in the nucleoplasm and cannot be buffered by SURF2 (Fig 7B). There, free 5S RNP particles can bind MDM2 to promote p53 activation and cell cycle arrest. In absence of SURF2, in normal conditions, any small amount of free-5S can induce p53 activation and affect cell cycle. Upon nucleolar stress, a greater fraction of free-5S RNP accumulates in nucleoplasm compared to normal cells, that promotes an even stronger response to NS (Fig 7C). In contrast, when SURF2 is over-expressed, free-5S particles released upon nucleolar stress are sequestered by SURF2, thus inhibiting their interaction with MDM2, which prevents p53 activation (Fig 7D). This free 5S RNP buffering capacity of SURF2 could be hijacked by cancer cells to resist to p53 activation upon nucleolar stress, hence its overexpression found in many cancers compared to healthy tissues (Fig 5A).

**Figure 7:**
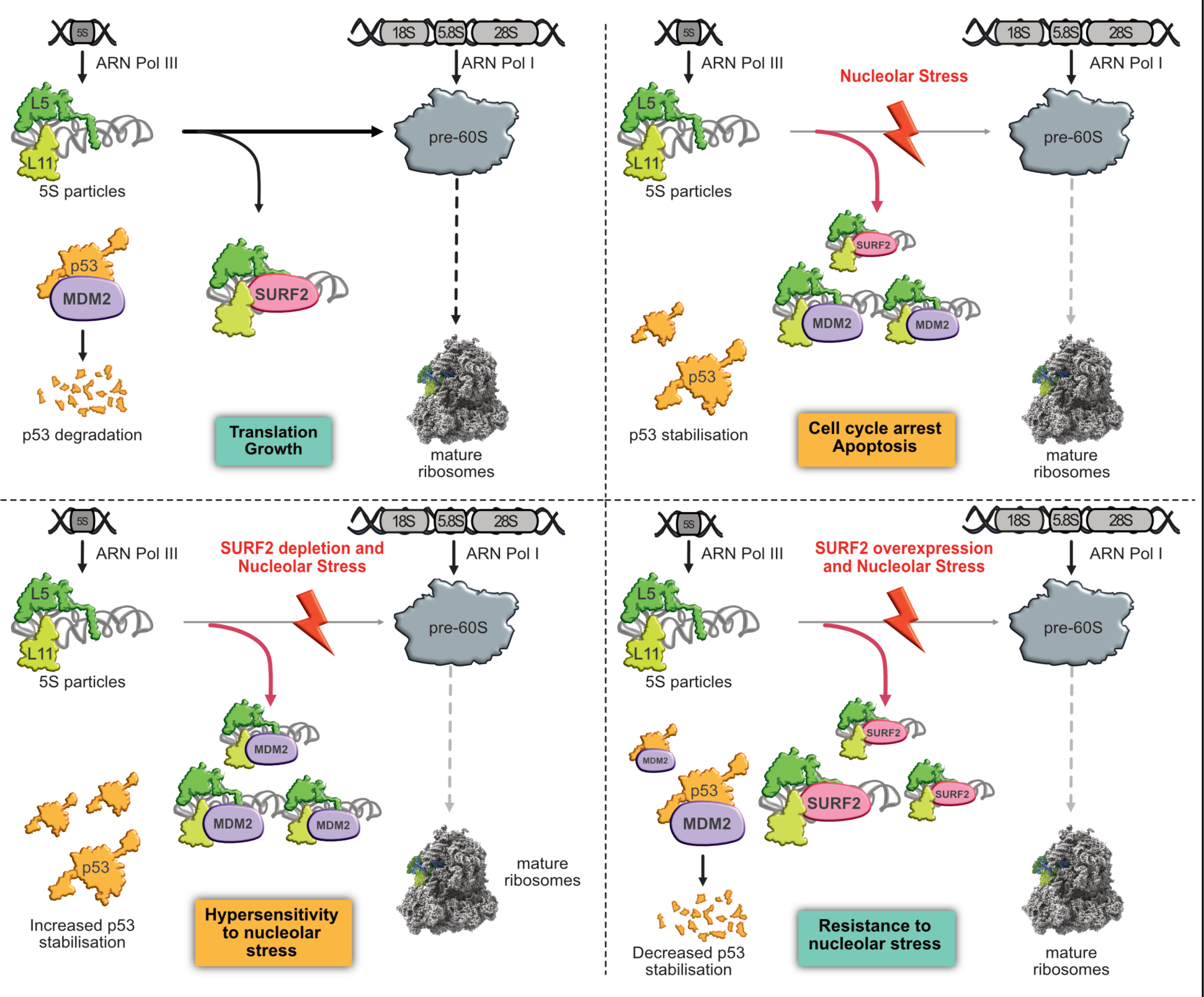
Model of SURF2 function in free 5S RNP regulation. Schematic representation of free-5S regulation by SURF2 in different conditions. In normal cells, both 5S and 47S rDNAs are transcribed by RNA polymerase III and I respectively. Ribosomes synthesis is producing pre-60S ribosomes and 5S particles constituted by the association of RPL5 and RPL11 to 5S rRNA are incorporated into these large pre-ribosomes. The remaining overproduced free 5S are bound by SURF2 to avoid MDM2-Free 5S interaction, which would induce p53 stabilization and cell cycle arrest. At the same time, p53 is constantly ubiquitinated by MDM2 to promote its degradation by the proteasome. After nucleolar stress (drug-induced or caused by genetic mutations/ribosomopathies), ribosome synthesis is impaired and a larger amount of free 5S particles accumulate in the nucleoplasm. The extra free 5S can then be recognized by MDM2, which can no longer ubiquitinylate p53, thereby stabilizing p53 and promoting cell cycle arrest. In cells lacking SURF2, nucleolar stress still impairs ribosome synthesis, but this time even more free 5S are able to bind to MDM2, inducing stronger stabilization and activation of p53, followed by more cell cycle arrest. In contrast, in cells overexpressing SURF2, nucleolar stress promotes free-5S accumulation in the nucleoplasm, all of which are recognized by SURF2, which competes with MDM2 for binding. As a result, MDM2 is free and ubiquitinylates p53, conferring to these cells a capacity to resist to nucleolar stress.

This regulation of free-5S RNPs by SURF2 opens the possibility of highly innovative therapeutic strategy in order to increase cancer cell sensitivity to chemotherapeutic drugs, since most of them rely on NS to activate p53 and induce cell death. This is supported by the negative correlation between overall survival and SURF2 expression level in some cancers. It is of particular interest to notice that co-treatment of U2OS cell by Actinomycin D and SURF2 siRNAs induce cell death whereas these treatments taken individually only promote cell cycle arrest. This suggests a synergistic effect of these two treatments. Furthermore, typical drawbacks of chemotherapies to treat cancer are the associated secondary effects. Hence, finding a way to reduce drug dosage by specifically increase cancerous cell sensitivity to stress (without affecting pathways equally important for healthy cells, such as ribosome production), could represent a means of increasing the therapeutic window and to reduce adverse effects. Of note, another study underlines the link between 5S RNPs homeostasis and HCC cancers, suggesting a particular link between some cancers and 5S RNPs metabolism, which needs to be studied in depth (*25*).

Free 5S RNP homeostasis is also key in a group of diseases originating from ribosome production defects and regrouped as ribosomopathies (*33*). In Diamond-Blackfan Anemia (DBA), a well characterized ribosomopathy, activation of p53 by free-5S particles is likely at the core of the etiology of these diseases, since some symptoms are linked to early p53 activation such as growth retardation, developmental problems and even erythropoiesis failure (*56–58*). In addition, the patients suffering from these diseases show a higher cancer incidence compared to general population. Hence, DBA is associated with a 30-fold increased risk to develop acute myeloid leukemia (AML), osteosarcoma or colon cancer (*59*). Recently, two different studies on patients suffering from Shwachman-Diamond syndrome (SBDS), another ribosomopathy due to mutations in the pre-60S subunit assembly factor SBDS, nicely demonstrated how patients first harbor hypo-proliferative patterns due to SBDS mutations, then acquire compensatory mutations in TP53 to escape the growth selective pressure, and thus develop a pre-malignant cancer state (*60–62*). Because SURF2 is able to impede p53 activation following NS, it may represent there again an interesting therapeutic target. Mimicking SURF2 or part of it using small peptides or compound drug approaches could be used to mitigate p53 activation by free 5S RNPS in these patients. It would also be interesting to investigate if there is any correlation between SURF2 expression level and the high variability of symptoms severity observed between patients from the same family.

Of note, over-expression of SURF2 in U2OS cells does not seem to alter ribosome synthesis nor cell fitness as seen by cell cycle analysis (Fig S3). First, this observation validates this strategy although testing this over-expression in more relevant cellular models will be needed in the future. Second, this implies that 5S RNP integration into ribosome is not affected by SURF2 over-expression, suggesting that SURF2 only interacts with free 5S RNPs that are released from or not integrated into pre-ribosomes. How free 5S RNP particles interact with their different partners and how this is regulated in cells remains to be elucidated.

In summary the work presented here, demonstrates how free 5S are key in cell stress response, and highlights SURF2 as a therapeutic target to either increase cancer cells sensitivity to chemotherapeutic drugs or impede free 5S RNP function to reduce p53-associated symptoms in genetic diseases such as ribosomopathies.

## Materials and Methods

### Cell lines

The U-2OS Flp-In T-Rex cell lines were produced according manufacturer’s instruction (Invitrogen/Thermo Fisher). The cDNAs of RPL5 or SURF2 were cloned into a pcDNA5-FRT-TO vector to enable expression of the protein with a C-terminal 2xFLAG-PreScission protease site-His6 (FLAG) tag. Alternatively, the cDNA of SURF2 was fused to EGFP tag and cloned into pcDNA5-FRT-TO plasmid. These plasmids or the empty plasmids (control) were co-transfected with a pOG44 plasmid into Flp-In T-Rex U-2OS or HepG2 cells and cells that had stably integrated the plasmid into their genome were selected using Hygromycin B, according to the manufacturer’s instructions. Expression of tagged proteins was induced by addition of 1µg/mL of tetracycline for 24h prior harvesting.

All the cell lines were maintained in high-glucose Dulbecco’s modified Eagle’s medium supplemented with 10% fetal bovine serum and 1mM sodium pyruvate. The cells were incubated at 37°C in a humidified incubator containing 5% CO2. Flip-In T-Rex cells were maintained using hygromycin B (30 µg/mL, InvivoGen, ant-hg-5) and blasticidin (30 µg/mL, InvivoGen, ant-bl-1: 1). The expression of proteins was induced with tetracyclin (1 µg/mL; Sigma-Aldrich, T7660). Cell were treated with actinomycin D (10 ng/mL, Sigma-Aldrich, A9415).

A pool of four siRNA duplexes from eurogentech were used to target SURF2 mRNA (CUG-CAA-GUG-AUG-ACA-GCA-U/AUG-CUG-UCA-UCA-CUU-GCA-G, GGA-GGG-AGG-ACC-AGA-UGG-A/UCC-AUC-UGG-UCC-UCC-CUC-C, AAG-CAC-AUG-CCG-UGA-AGU-U/AAC-UUC-ACG-GCA-UGU-GCU-U and CCA-GCG-AGC-UCU-GUG-UAA-A/UUU-ACA-CAG-AGC-UCG-CUG-G. On duplex for RPL11 mRNA AAG-GUG-CGG-GAG-UAU-GAG-UUA/UAA-CUC-AUA-CUC-CCG-CAC-CUU and on duplex for RPL5 mRNA GCC-ACA-AUG-UUG-CAG-AUU-A/UAA-UCU-GCA-ACA-UUG-UGG-C were used. Each siRNA solution was added at a final concentration of 5 µM to 10^6^ cells diluted in ZAP buffer (10mM sodium phosphate buffer, pH 7.25, containing 250 mM sucrose and 1 mM MgCl2). Electroporation was performed at 240 V with a Gene Pulser (Bio-Rad). After 5 min incubation at RT, cells were plated and grown at 37°C for 48h. Depletion of SURF2 was completed with a second round of siRNA treatment (96h total). Control cells were electroporated with a scramble siRNA (siRNA-negative control duplex; Eurogentec).

### Label-free quantitative proteomics

#### Protein preparation for 5S particle partners identification

For the proteome analysis, cells were prepared in quadruple biological replicates for four conditions: i) U2OS cells (control); ii) U2OS cells overexpressing RPL5-flag; iii) control cells treated with Actinomycin D; iv) U2OS cells overexpressing RPL5-flag and treated with actinomycin D (10 ng/mL, Sigma-Aldrich A9415). RPL5-Flag proteins and associated complexes were immunoprecipitated using the same protocol as “proteins immunoprecipitation after sucrose cushion” until the elution step. Trapped proteins on anti-Flag beads were eluted using Flag buffer (20 mM Tris-HCl pH 7.5, 200 mM NaCl, 5 mM MgCl2) supplemented with 0.4 mg/mL 2xflag peptide (H-MDYKDDDDKGTDYKDDDDKG-OH, Schafer), precipitated by TCA (Sigma-Aldrich, T9159) and glycogen (Thermo Scientific, R0551) and re-suspended in a buffer containing 20 mM Tris-HCl pH 7.5, 200 mM NaCl, 5 mM MgCl2, 5% glycerol, 5 % SDS. Then the protocol “*Trypsin digestion and mass spectrometry analysis” was followed*.

#### Protein preparation for SURF2 partners identification

For the proteomic analysis, U2OS control cells were prepared in quadruple biological replicates for two conditions: i) GAPDH immunoprecipitation and ii) SURF2 immunoprecipitation. Cells were harvested, washed with PBS with 1mM EDTA, resuspended in buffer E (20 mM Tris-HCl pH 7,5, 200 mM NaCl, 5 mM MgCl2, 0.5 mM EDTA, 0.2% Triton, 1mM DTT, complete protease inhibitor cocktail (Roche), RNase ribonuclease inhibitor (Promega, N261B)) and disrupted with a Bioruptor Sonicator by sonication (2 min, 5 s/5 s on/off, 20% amplitude). Cell debris were removed by centrifugation (10 min, 14,000 g, 4 °C). Protein concentrations of the extracts were determined using a Bio-Rad protein assay kit (Biorad, 5000006). The same amounts of proteins were incubated with antibodies (anti-SURF2 or anti-GAPDH antibodies) coupled to protein G sepharose beads (Cytiva, 17061801) for 2 h at 4 °C. After immunoprecipitation, beads were washed tree times with buffer E (20 mM Tris-HCl pH 7.5, 200 mM NaCl, 5 mM MgCl2, 0.5 mM EDTA, 0.2% Triton, 1 mM DTT) and the associated proteins were eluted with 2X buffer (100 mM Tris-HCl pH 7.5, 10% glycerol, 10 % SDS). Then the protocol “*Trypsin digestion and mass spectrometry analysis” was followed*.

#### Trypsin digestion and mass spectrometry analysis

Disulfide bonds were reduced with 25 mM DTT for 5 min at 95°C under agitation followed by an alkylation of cysteine residues in 60 mM iodoacetamide for 30 min in the dark at room temperature. Each reduced/alkylated protein sample was then digested using the S-Trap™ Mini spin column protocol (*63*). Briefly, undissolved matter was removed by centrifugation for 8 min at 13,000g. 12% aqueous phosphoric acid was added at 1:10 to the protein sample for a final concentration of ∼1.2% phosphoric acid followed by seven volumes of S-Trap binding buffer (90% methanol, 100 mM TEAB, pH 7.1). After gentle mixing, the protein solution was loaded onto an S-Trap filter by centrifugation at 4,000 g. Afterwards, the captured proteins were washed six times with 400 µL S-Trap binding buffer. Digestion was performed over-night at 37°C by addition of 20 µL of trypsin (Sequencing Grade Modified Trypsin, Promega) at 37.5 ng/µL in 50 mM ammonium bicarbonate. The digested peptides were eluted by addition of 40 µL of 50 mM ammonium bicarbonate, followed by 40 µL of 0.2% formic acid (FA), and finally 35 µL of 50% aqueous acetonitrile containing (ACN) 0.2 % FA. Each elution was performed at 4,000 g during 1 min. The different eluates were pooled together, dried down, resuspended in 20 µl of 0.05% trifluoroacetic acid (TFA) in 2% ACN, and sonicated for 10 min before analysing by online nanoLC using an UltiMate® 3000 RSLCnano LC system (ThermoScientific, Dionex) coupled to an Orbitrap Fusion™ Tribrid™ mass spectrometer (Thermo Scientific, Bremen, Germany) operating in positive mode. 200 ng of each sample were loaded onto a 300 µm ID × 5mm PepMap C18 pre-column (Thermo Scientific, Dionex) at 20 µL/min in 2% ACN, 0.05% TFA. After 3 min of desalting, peptides were on-line separated on a 75 µm ID × 50 cm C18 column (in-house packed with Reprosil C18-AQ Pur 3 µm resin, Dr. Maisch; Proxeon Biosystems, Odense, Denmark) equilibrated in 90% of buffer A (0.2% FA), with a gradient of 10 to 30% of buffer B (80% ACN, 0.2% FA) for 100 min then 30% to 45% for 20 min at a flow rate of 300 nL/min. The instrument was operated in data-dependent acquisition (DDA) mode using a top-speed approach (cycle time of 3 s). Survey scans MS were acquired in the Orbitrap over 350–1400 m/z with a resolution of 120,000 (at 200 m/z), an automatic gain control (AGC) target value of 4e5, and a maximum injection time of 60 ms. Most intense multiply charged ions (2+ to 6+) per survey scan were selected at 1.7 m/z with quadrupole and fragmented by Higher Energy Collisional Dissociation (HCD). The monoisotopic precursor selection was turned on, the intensity threshold for fragmentation was set to 25,000 and the normalized collision energy was set to 28%. The resulting fragments were analyzed in the Orbitrap with a resolution of 30,000 (at 200 m/z), an automatic gain control (AGC) target value of 5e4, and a maximum injection time of 54 ms. Dynamic exclusion was used within 60 s with a 10 ppm tolerance, to prevent repetitive selection of the same peptide. For internal calibration the 445.120025 ion was used as lock mass.

#### MS-based protein identification

Acquired MS and MS/MS data as raw MS files were converted to the mzDB format using the pwiz-mzdb converter (version 0.9.10, https://github.com/mzdb/pwiz-mzdb) executed with its default parameters (*64*). Generated mzDB files were processed with the mzdb-access library (version 0.7, https://github.com/mzdb/mzdb-access) to generate peaklists. Peak lists were searched against UniProtKB/Swiss-Prot protein database with homo sapiens taxonomy in Mascot search engine (version 2.6.2, Matrix Science, London, UK). Cysteine carbamidomethylation was set as a fixed modification and methionine oxidation as variable modification. Up to two missed trypsin/P cleavages were allowed. Mass tolerances in MS and MS/MS were set to 10 ppm and 0.6 Da, respectively. Validation of identifications was performed through a false-discovery rate set to 1% at protein and peptide-sequence match level, determined by target-decoy search using the in-house-developed software Proline software version 1.6 (*65*). The mass spectrometry proteomics data have been deposited to the ProteomeXchange Consortium via the PRIDE [1] partner repository with the dataset identifier PXD047905.

### Immunoblotting assays

Protein extraction was done in ice-cold lysis buffer (1% Triton, 50 mM Tris-HCl pH 7.4, 200 mM NaCl, 1 mM EDTA and complete protease inhibitor cocktail (Roche)). Protein concentrations of the extracts were determined using a Bio-Rad protein assay kit (Biorad, 5000006). Proteins were diluted in Invitrogen 2X sample buffer (NuPAGE™ LDS Sample Buffer (4X) (NP0007) and NuPAGE™ Sample Reducing Agent (10X) (NP0009)) and 20 µg/lane of protein were loaded on NuPAGE™ 4 to 12 %, Bis-Tris protein gels (Invitrogen, NP0321BOX) and transferred to nitrocellulose membrane using the Trans-Blot Turbo Transfer System from Biorad. Membranes were immunoblotted with a primary antibody, followed by incubation with a secondary antibody coupled with HRP. The blots were visualized using the Clarity Western ECL kit from Biorad. The following antibodies were used: p53 (Invitrogen, MA5-12557 (DO7)), p21 (CDKN1A/p21CIP1 AB clonal, A2691), SURF2 (Bethyl lab, A304-611A), Flag (MERCK, F3165), GAPDH (Genetex, GTX627408), RPL11 (Invitrogen, 37-3000), RPL5 (Invitrogen, PA5-102539), RPL17 (Gentech, GTX111934), Fibrilarin (collaborators), MDM2 (SMP14 Santa Cruz, SC965), Actin (Sigma, A4700).

### Cell cycle assays

The cell cycle was analyzed by flow cytometry. Cells were harvested by trypsinization, fixed in 70% ethanol, and stained with DAPI (1 µg/ml; Sigma, D9542) in PBS completed with RNase A (100 µg/ml, Thermo Scientific, EN0531) for 30 min at RT. Samples were then analyzed for their DNA content using CytoFLEX S Flow Cytometer and CytExpert software.

### Viability assays

Cells depleted for SURF2 were plated in 6 well culture dishes. After 12h, cells were treated with actinomycin D (10 ng/mL, Sigma-Aldrich A9415) then, at each time point, cells were incubated with 400 µL of 1% crystal violet staining solution (Sigma-Aldrich, V5265) for 20 min at RT. Cells were washed 2 times with PBS and 3 pictures were taken under microscope. Then, crystal violet was resolubilized with 33% acid acetic and diluted into 96 well plate to read optical density OD595 with a plate reader.

### Northern-blot assays

Total RNAs were extracted with Trizol from cell pellets containing 20×10^6^ cells. The aqueous phase was extracted with phenol-chloroform-isoamylic alcohol (25:24:1; Sigma), then with chloroform. Total RNAs were recovered after precipitation with 2-propanol. For northern blot analyses, 3µg/lane of total RNAs were separated on two types of gels. Long RNAs were separated on a 1.2% agarose gel containing 1.2 % formaldehyde and Tri/Tri buffer (30 mM triethanolamine, 30 mM tricine, pH 7.9). Small RNAs were separated on 6% polyacrylamide gel containing 7% urea and TBE buffer (90 mM Trizma base, 90mM Boric acid, 2mM EDTA). Then, RNAs were transferred to a Hybond N+ nylon membrane by a passive transfer overnight. Pre-hybridization was performed for 1 h at 45° C in a buffer containing 6× SSC, 5× Denhardt’s solution, 0.5 % SDS and 0.9 g/mL tRNAs). The 5′-radiolabeled oligonucleotide probe were incubated overnight. The sequences of the probes were: ITS1: CCT-CGC-CCT-CCG-GGC-TCC-GTT-AAT-GAT-C, ITS2: GCG-CGA-CGG-CGG-ACG-ACA-CCG-CGG-CGT, + CTG-CGA-GGG-AAC-CCC-CAG-CCG-CGC-A, 18S: CCG-GCC-GTC-CCT-CTT-AAT-CAT-GGC, 28S: CCC-GTT-CCC-TTG-GCT-GTG-GTT-TCG-CTG-GAT-A, 5.8S: GGG-GCG-ATT-GAT-CGG-CAA-GCG-ACG-CTC, 5S: CCU-CGC-CCU-CCG-GGC-UCC-GUU-AAU-GAU-C. Membranes were washed twice for 10 min in 2× SSC, 0.1% SDS and once in 1× SSC, 0.1% SDS, and then exposed. Signals were acquired with a Typhoon Trio PhosphoImager (GE Healthcare) and quantified using the MultiGauge software.

### Immunofluorescence assays

The expression of SURF2-GFP was induced to the same level of the endogenous SURF2 protein (tetracycline at 5 ng/mL for 24h). Cells were seeded in 12-well plates on microscope cover glasses and grown for 24h. Cells were fixed with 4% paraformaldehyde for 5 min, permeabilized with (0.1% Triton X-100 and 0.02% SDS in PBS). Fixed cells were incubated in blocking solution 2% BSA in PBS for 30 min and incubated overnight at 4°C with anti-fibrillarin antibodies at 1:200. Cells were washed 3 times for 5 min with (2% BSA in PBS), and subsequently incubated for 30 min with secondary antibodies (Alexa Fluor goat anti-mousse IgG (H+L)/Alexa Fluor 647 secondary antibody (Invitrogen, A21236) at 1:1000. After 3 washes, cells were incubated briefly in 0.1% Triton X-100, 0.02% SDS in PBS, and post-fixed with 4% PFA in PBS. Cells were incorporated with DAPI (1µg/mL, Sigma, D9542) for 10 minutes. After washes with PBS, coverslips were mounted in Mowiol. Imaging was performed on a Leica TCS SP8 MP Multiphoton microscope. Images were captured in confocal mode using x63 objective. Image analyses were performed using ImageJ software.

### Sucrose sedimentation profiling

Buffers contains cycloheximide (10 µg/mL, Merck, C7698) at each step of this protocol. 50×10^6^ cells were harvested and resuspended in a lysis buffer (10 mM Hepes KOH, pH 7.9, 1.5 mM MgCl2, 10 mM KCl, 1 mM DTT, 10 µg/mL cycloheximide). Then cells were homogenized with a Dounce tissue grinder on ice with a tight pestle and centrifuged at 1000g for 10 min at 4 °C. The top soluble phase was clarified through two centrifugations at 10 000g for 15 min at 4 °C. Supernatants were collected and protein concentrations were determined using a Bio-Rad protein assay kit (Biorad 5000006). 750 µg of extracts were loaded on a 10–50% sucrose gradient. Gradients were centrifuged at 36,000 rpm for 2 h at 4 °C in an Optima L-100XP ultracentrifuge (Beckman–Coulter). Following centrifugation, the fractions were collected using a Foxy Jr fraction collector (Teledyne ISCO) and the absorbance at 254 nm was measured with a UA-6 device (Teledyne ISCO). For protein analyses, 250 µL of each fraction were precipitated with TCA (Sigma-Aldrich, T9159) and glycogen (Thermo Scientific, R0551) and protein pellets were resuspended in Invitrogen 2X sample buffer (NuPAGE™ LDS Sample Buffer (4X) (NP0007) and NuPAGE™ Sample Reducing Agent (10X) (NP0009)).

### Cell fractionation following PSE method

Cells were harvested by scrapping with PBS and pelleted by centrifugation. Cells were gently resuspended in SN1 buffer (20 mM HEPES-NaOH (pH 7.5), 130 mM KCl, 10 mM MgCl2, 0.05% Igepal, 600U/mL RNasin ribonuclease inhibitor (Promega, N261B)) completed with cOmplete protease inhibitor (1/30) (Roche) and centrifugated (3,800 rpm, 3 minutes, 4°C). SN1 supernatants were collected and proteins were diluted in Invitrogen 2X sample buffer (NuPAGE™ LDS Sample Buffer (4X) (NP0007) and NuPAGE™ Sample Reducing Agent (10X) (NP0009)). SN1 was then store at −80°C. The previous pellets of cellular extracts were vortexed with SN1 buffer and spined down (3,800 rpm, 3 min, 4°C). The supernatants were eliminated and the pellets were resuspended in SN2 buffer (10 mM HEPES-NaOH (pH 7.5), 10 mM NaCl, 5 mM MgCl2, 0.1% Igepal, 0.5 mg/mL heparin, 600 U/ml RNasin ribonuclease inhibitor (Promega, N261B)) completed with Dnase I (Invitrogen, 18068015), then incubated 10 min at RT with gentle mixing. Extracts were centrifugated (11,500 rpm, 10 min, 4 °C). SN2 supernatants were collected and proteins were diluted in Invitrogen 2X sample buffer (NuPAGE™ LDS Sample Buffer (4X) (NP0007) and NuPAGE™ Sample Reducing Agent (10X) (NP0009)). SN2 was then store at −80°C. Finally, remaining pellets of cellular extracts were resuspended with SN3 buffer (20 mM HEPES-NaOH (pH 7.5), 200 mM NaCl, 4 mM EDTA, 0.1% Igepal, 0.04% sodium deoxycholate, 4 mM imidazole, 0.1 mg/ml heparin, 1mM DTT, cOmplete protease inhibitor (1/100), 600 U/ml RNasin ribonuclease inhibitor (Promega, N261B)) and mixed for 20 minutes at RT. Extracts were centrifugated (11,500 rpm, 10 min, 4°C). SN3 supernatants were collected and proteins were diluted in Invitrogen 2X sample buffer (NuPAGE™ LDS Sample Buffer (4X) (NP0007) and NuPAGE™ Sample Reducing Agent (10X) (NP0009)). SN3 was then store at −80°C. Proteins were analyzed by western blots.

### Proteins immunoprecipitation after sucrose cushion

Cells were harvested, washed with 1mM EDTA in PBS, resuspended in buffer E (20mM Tris-HCl pH 7.5, 200 mM NaCl, 5 mM MgCl2, 0.5 mM EDTA, 0.2% Triton, 1 mM DTT, complete protease inhibitor cocktail (Roche), RNasin ribonuclease inhibitor (Promega, N261B)) and disrupted with a Bioruptor Sonicator by sonication (2 min, 5 s/5 s on/off, 20% amplitude). Cell debris were removed by centrifugation (10 min, 14,000 g, 4 °C). Protein concentrations of the extracts were determined using a Bio-Rad protein assay kit (Biorad, 5000006). The same amounts of proteins were loaded on a double sucrose cushion (20% and 30% sucrose) and centrifuged at 190,000 g for 2 h at 4 °C in an Optima L-100XP ultracentrifuge (Beckman–Coulter). Extracts were incubated with pre-washed anti-Flag beads (Sigma, A2220) or with antibodies (anti-SURF2 or anti-MDM2 antibodies) coupled to protein G sepharose beads (Cytiva, 17061801) for 2 h at 4 °C. 10 % of inputs were conserved, precipitated with TCA (Sigma-Aldrich, T9159) and glycogen (Thermo Scientific, R0551), pellets were resuspended with Invitrogen 2X sample buffer (NuPAGE™ LDS Sample Buffer (4X) (NP0007) and NuPAGE™ Sample Reducing Agent (10X) (NP0009)). After immunoprecipitation, beads were washed tree times with Flag buffer (20 mM Tris-HCl pH 7.5, 200 mM NaCl, 5 mM MgCl2) and the associated proteins were eluted with Invitrogen 2X sample buffer (NuPAGE™ LDS Sample Buffer (4X) (NP0007) and NuPAGE™ Sample Reducing Agent (10X) (NP0009)).

### _[5’-32P]_pCp labelling of immunoprecipitated RNAs

Cells were harvested in TBS buffer (150 mM NaCl, 40 mM Tris-HCl pH 7.4), resuspended in lysis buffer (150 mM NaCl, 0.05% Igepal, 50 mM Tris-HCl pH 7.4, 5 mM MgCl2) completed with complete protease inhibitor cocktail (Roche) and disrupted with a Bioruptor Sonicator by sonication (2 min, 5 s/5 s on/off, 20% amplitude). Cell debris were removed by centrifugation at 16,000 g for 10 min and the clarified extracts were incubated with antibodies (anti-SURF2 or anthi-HEXIM1 antibodies) coupled to protein G sepharose beads (Cytiva, 17061801) for 2 h at 4 °C. Then, beads were washed 3 times and immunoprecipitated RNAs were extracted using phenol-chloroform protocol. RNAs were incubated with ^[5’-32P]^ pCp and T4 RNA ligase (Promega, M1051) O/N at 4 °C. Then RNAs were precipitated with ammonium acetate and ethanol for 10 min at -70 °C and pelleted by centrifugation (10 min, 13,000 g). RNA pellets were washed with 70 % ethanol and recovered with a formamide loading buffer. Then, a loading dye buffer was added and RNA were analyzed on a 12 % acrylamide gel.

### In vitro pull-down assays

The cDNAs of the SURF2 or SURF2(1–136) were cloned into pScodon plasmid (open biosystem) in translational fusion with HIS^6^ (His tag). RPL5 or RPL11 cDNAs were synthetized by Genscript and cloned into pGEX-6T. The expression and purification were essentially as described (*66*). Briefly, the proteins were expressed in the BL21 strain from *Escherichia coli* at 37°C in LB medium (Sigma) supplemented with 100 µg/mL ampicillin until OD_600_ between 0.4 and 0.5. Recombinant protein expression was induced by adding 1 mM isopropyl-β-D-1-thiogalactopyranoside, incubating overnight at 20°C, harvesting by centrifugation and cell pellets were frozen at -20°C. Cells pellets were resuspended in buffer A (300 mM NaCl, 20 mM Tris-HCl pH 8.0, 0.5 mM EDTA, 10 mM B-mercaptonethanol, 10% glycerol, tablet roche, 5mM Imidazole) supplemented with complete EDTA-free protease inhibitors (Roche). Cells were lysed by sonication, and lysate was centrifuged at 20,000 rpm for 30 min. The cleared lysates were mixed as indicated for 2 hours at 4 °C. The mix was then loaded on gluthatione-sepharose beads pre-equilibrated with buffer A (300 mM NaCl, 20 mM Tris-HCl pH 8.0, 0.5 mM EDTA, 10 mM B-mercaptonethanol, 10% glycerol, tablet roche, 5mM Imidazole) and mixed for 2 h at 4 °C. Beads were then washed 4 times with buffer A and resuspended in loading dye (1X LDS with denaturing reagent from Invitrogen) and loaded on Nu-PAGE 4-12% gels using 1X MOPS as running buffer.

### Pancancer analysis

Different databases were used to export data of interest, namely transcriptome quantification, clinical and genomic data.

GTEx: (https://gtexportal.org/home/) Genotype-Tissue Expression is a public platform containing molecular data of healthy tissues derived from people of all age gender and ethnicity.

TCGA: (https://www.cancer.gov/ccg/research/genome-sequencing/tcga) The Cancer Genome Atlas is a public databased compiled from the National Cancer Institute’s CDM portal. It includes 33 cancers from 11,000 patient samples over 12 years and contains annotated clinical data and molecular data. TARGET:(https://www.cancer.gov/ccg/research/genome-sequencing/target/about) therapeutically Applicable Research to Generate Effective Treatments is a public platform dedicated to molecular characterization of paediatric cancers with available clinical, genomic and transcriptomic data. cBioPortal: (https://www.cbioportal.org/) This public portal contains data from over 300 multidimensional studies in open-access. It includes genomic, transcriptomic, molecular and clinical data from multiple datasets. It allows exploratory analysis and corresponds to a global visualization webtool, which includes exportation of data from TCGA, TARGET, ICGC and other individual datasets. The raw data are not directly downloadable on the portal.

XENA UCSC: (https://xena.ucsc.edu/) The University of California Santa Cruz (UCSC) Xena browser. This public data portal contains over 1,500 datasets and 50 different types of cancers with clinical, genomic, transcriptomic and other type of data. It enables interactive exploratory analysis and exportation of accurate data of interest from TCGA, ICGC, GDC, TARGET, GTEx and other databases.

### Statistical Analysis

#### Label-free quantitative proteomics analysis

Proteins enriched over 2-fold with a p-value below 0.05 were considered significantly enriched. For label-free relative quantification across samples, raw MS signal extraction of identified peptides was performed using Proline (*65*). The cross-assignment of MS/MS information between runs was enabled, allowing to assign peptide sequences to detected but non-identified features. Each protein intensity was based on the sum of unique peptide intensities and was normalized across all samples by the median intensity. Missing values were independently replaced for each run by its 5% quantile. For each pairwise comparison, an unpaired two-tailed Student’s t-test was performed and proteins were considered significantly enriched when their absolute log2-transformed fold change was higher than 1 and their p-value lower than 0.05. To eliminate false-positive hits from quantitation of low intensity signals, two additional criteria were applied: only the proteins identified with a total number of averaged peptide spectrum match (PSM) counts>4 and quantified in a minimum of two biological replicates, before missing value replacement, for at least one of the two compared conditions were selected. The p-value and fold change were calculated between the IP groups from RPL5_Flag overexpressing cells and the control IP groups (IPs with the α-flag antibody). The same statistical analysis was performed on equivalent IP groups resulting from cells incubated with Actinomycin D during 24 h. Volcano plots were drawn to visualize significant protein abundance variations between conditions in the presence or absence of drogue. They represent -log10 (p-value) according to the log2 ratio. The complete list of proteins identified and quantified in immunopurified samples and analyzed according to this statistical procedure is described in Supplementary file (Table S1, S2 and S3).

#### General Data Analysis

Data are expressed as means ± SD. All statistical data (n≥3) were calculated using GraphPad Prism. Statistical details and significance reports can be found in the corresponding figure legends.

#### PanCancer analysis

Datasets were uploaded from XENA UCSC portal (https://xena.ucsc.edu/) that compiled normalized gene expression levels from normal (GTEX) and tumoral (TCGA) tissues, as well as related clinical and genomic data of tumoral tissues (Goldman et al, Nat Biotechnol 2020; GTEx consortium, Nat Genet 2013; Grossman et al, New England J Medecine 2016). After data export and transformation, tumor datasets were built by conserving only tumoral tissues collected at diagnosis from non-metastatic patients. A quality control was performed to verify normal distribution of gene expression dataset (Shapiro test) or cohort consistency (follow-up median, survival association with gold-standards such as metastatic relapse). Comparison of gene expression between two conditions was assessed using non-parametric test (Wilcoxon-test). Survival was investigated by plotting Kaplan-Meier survival curves and log-rank tests using median as cut-off of gene expression. Overall survival (OS) corresponds to the length of time from the date of cancer diagnosis to the date of death or censoring, while Progression-Free survival (PFS) corresponds to the length of time from the date of cancer diagnosis to the date of progression (i.e., death or local/distant relapse) or censoring. The statistical significance was based on p-value < 0.05 where the H0 hypothesis is rejected. Statistics and visualization were performed using R studio (version 4.2.2).

## Acknowledgments

We would like to thank the engineers and staff working on the CBI facilities for their great help, as well as Marion Aguirrebengoa for her help on statistical analysis.

## Funding

Ligue Régionale Midi-pyrénées contre le cancer 62339 (SL)

Ligue National Contre le Cancer, PhD grant (ST)

Institut du cancer (INCA), PLBIO022-065, INCA-RESICA (SL, DR, QP, CPC, SL, VM

French Ministry of Research (Investissements d’Avenir Program, Proteomics French Infrastructure, ANR-10-INBS-08).

INSERM The CNRS

The University of Toulouse-Paul Sabatier

## Author contributions

Conceptualization: SL, VM, CPC, JM, PEG, NW

Methodology: SL, VM, ST, JM, CF, CPC, PES

Investigation: ST, PES, JR, DR, CF, QP, SC, SL

Visualization: ST, PES, CF, SL

Supervision: SL, CPC, VM, JM

Writing—original draft: SL, VM, JM, CPC

## Data and materials availability

All data are available in the main text or the supplementary materials.

