## Supplementary figures and images for "SURF2 is a MDM2 antagonist in triggering the nucleolar stress response"

### Supplemental Figure S1

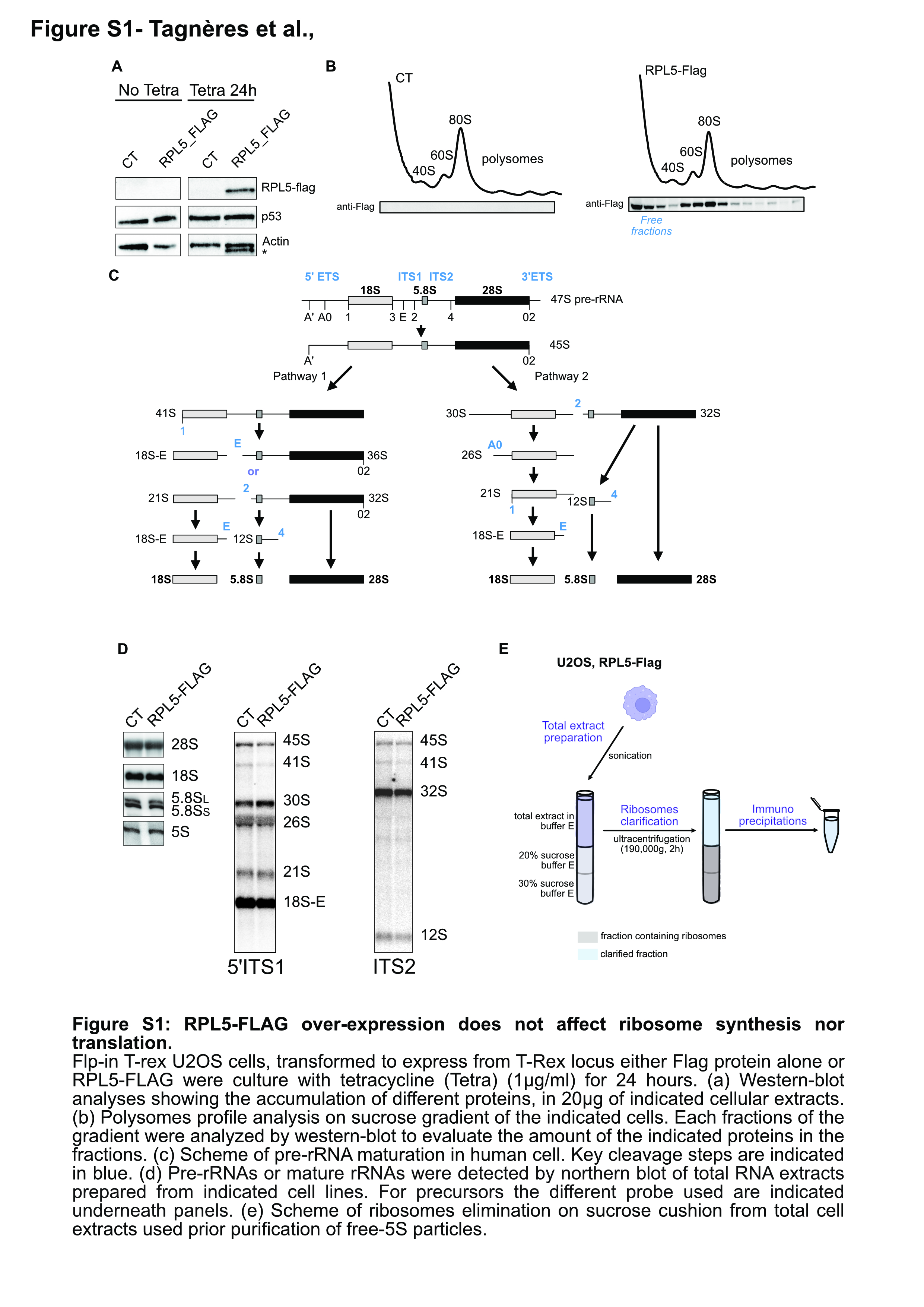

### Supplemental Figure S2

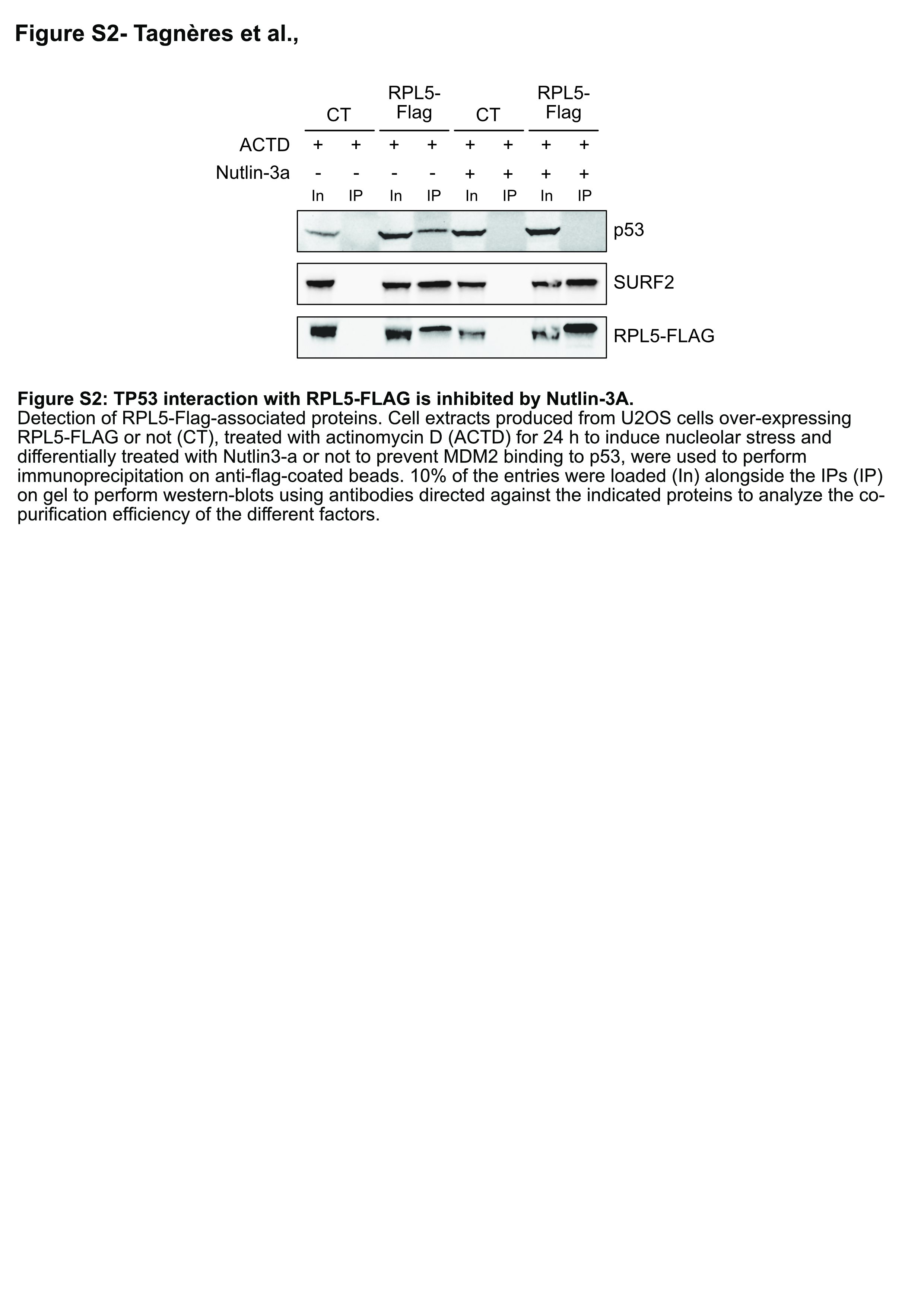

### Supplemental Figure S3

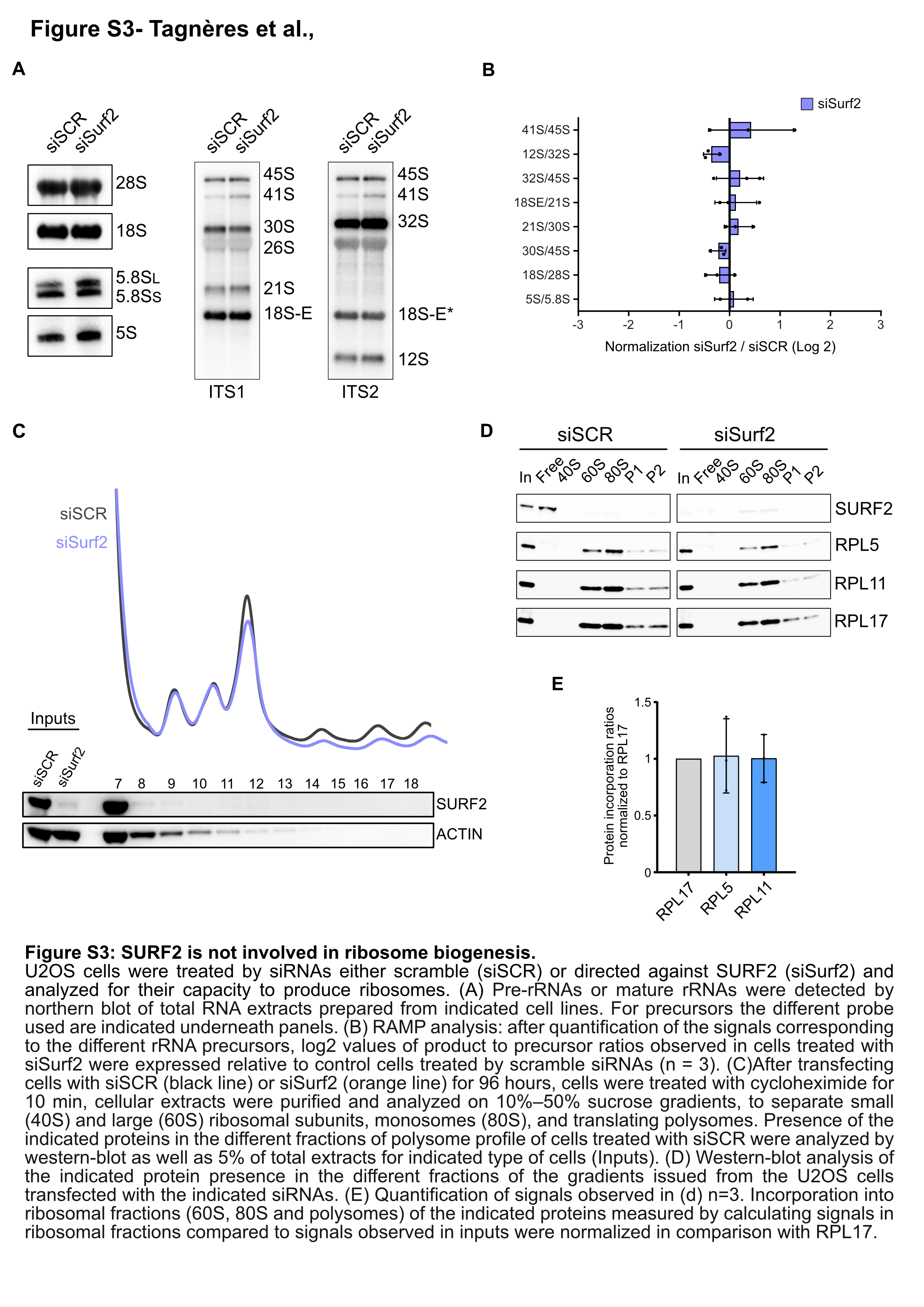

### Supplemental Figure S4

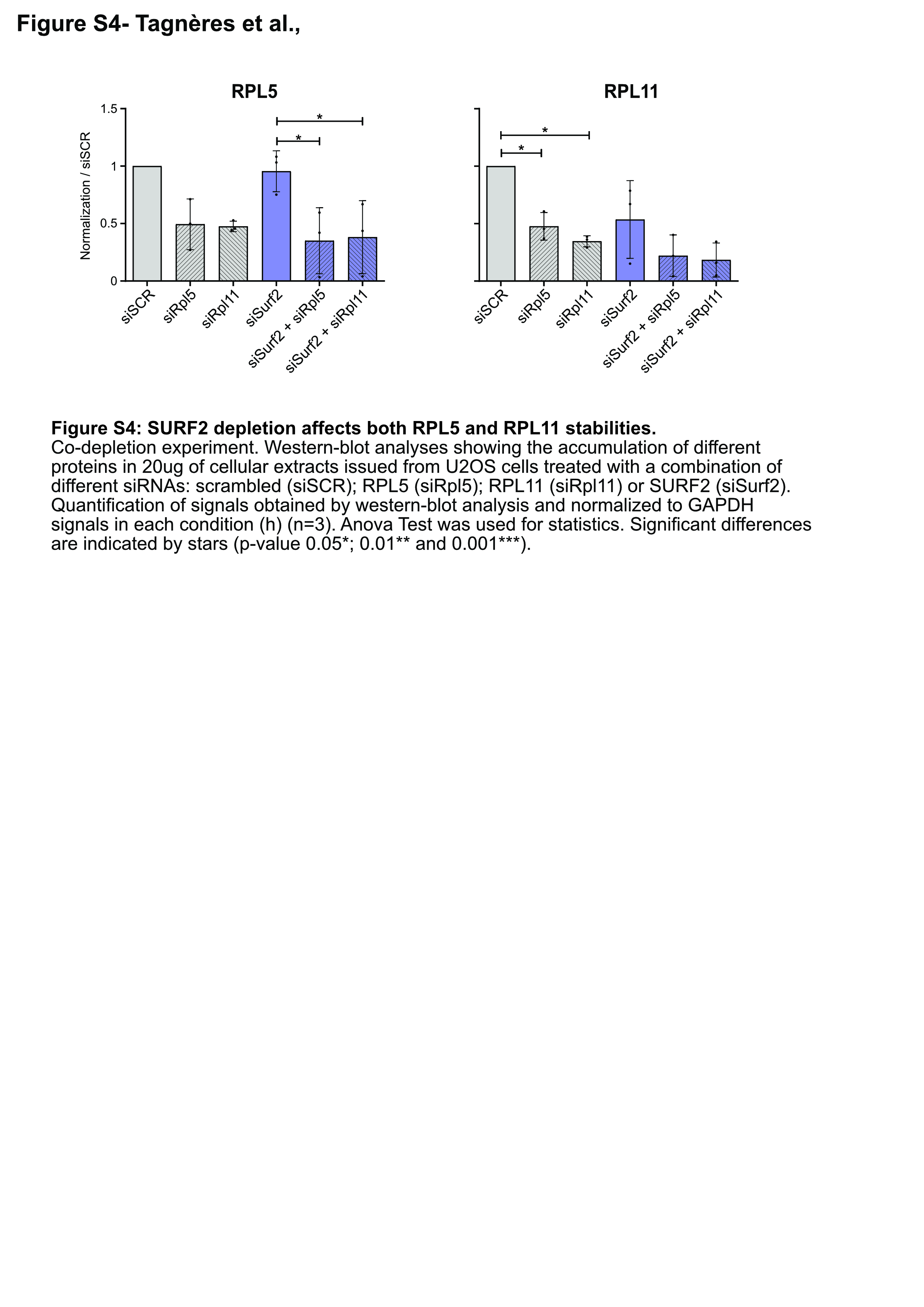

### Supplemental Figure S5

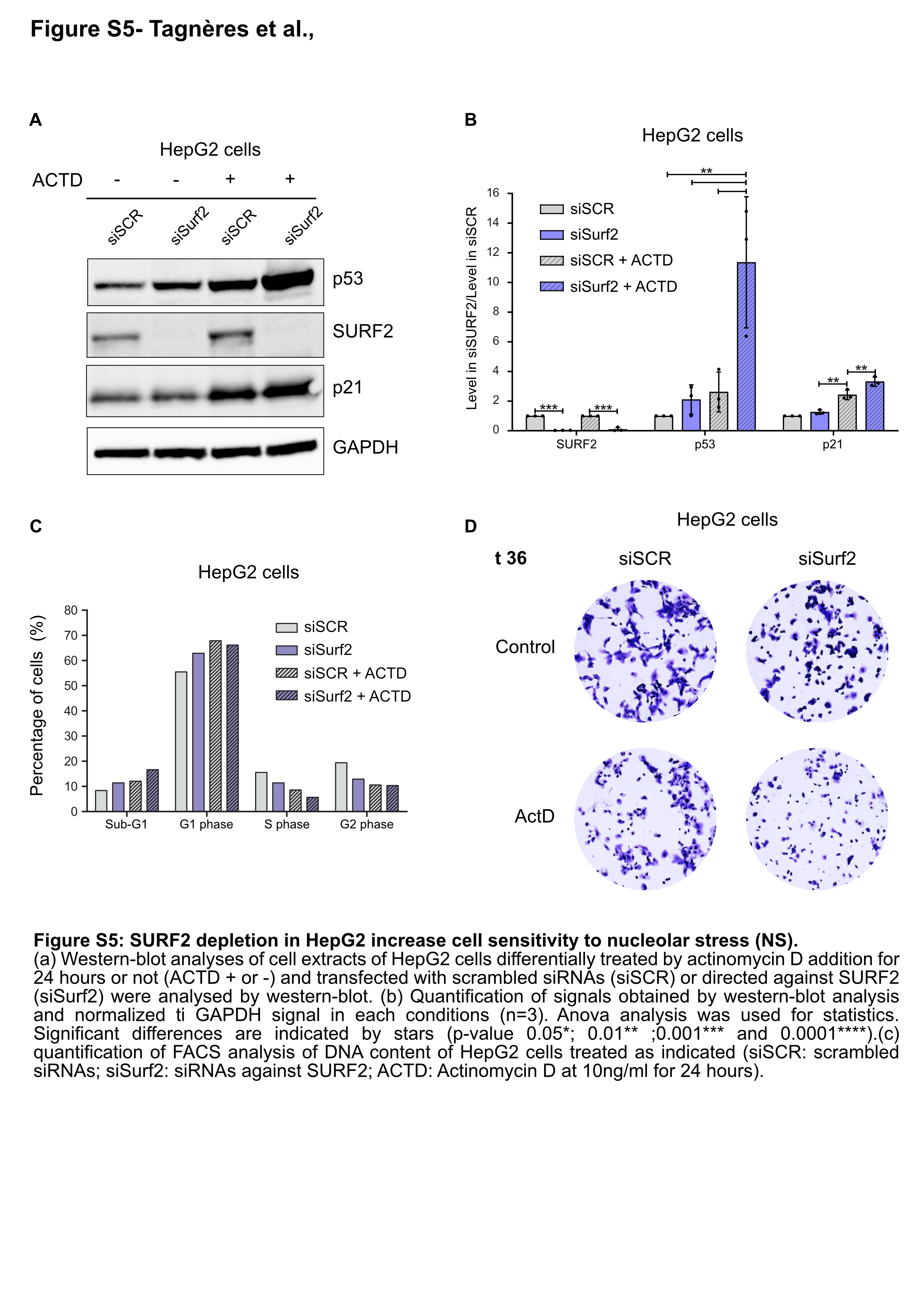

### Supplemental Figure S6

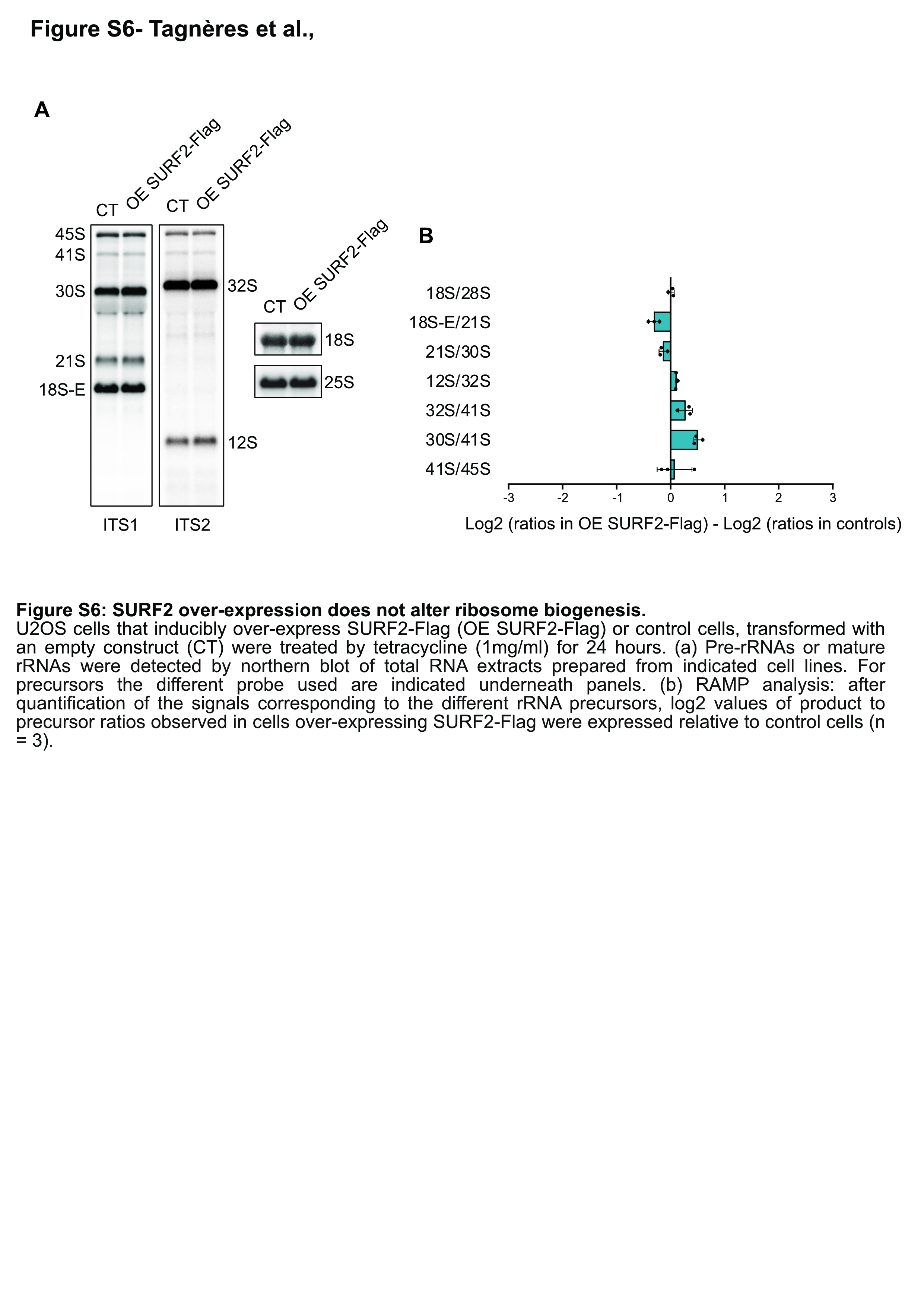
