## Supplemental Table S2 for "SURF2 is a MDM2 antagonist in triggering the nucleolar stress response"

**Table_S3 Mass spectrometry analysis of untreated U2OS extracts precipitated with either GAPDH or SURF2 antibodies**

| gene_name | iBAQ Surf2 | iBAQ GAPD | log2_fc_SURF2-GAPD | log10_pval_SURF2-GAPD | significant_SURF2-GAPD |
|---|---|---|---|---|---|
| DLST | 10295192,821 | 94627,423 | 6,952 | 1,340 | TRUE |
| SURF2 | 5528257,167 | 55182,745 | 6,813 | 2,365 | TRUE |
| NDUFC1 | 2203784,500 | 26348,768 | 6,589 | 2,977 | TRUE |
| IGKV1-17 | 1224701,417 | 25323,937 | 5,801 | 5,966 | TRUE |
| RNF220 | 482829,392 | 10167,162 | 5,780 | 5,377 | TRUE |
| TRIM21 | 9026163,022 | 190457,736 | 5,779 | 3,347 | TRUE |
| DLD | 1922278,222 | 57235,917 | 5,256 | 1,431 | TRUE |
| MRPS36 | 2885264,524 | 92701,366 | 5,113 | 1,499 | TRUE |
| OGDH | 3123475,200 | 110055,357 | 5,013 | 1,428 | TRUE |
| TUBG1 | 1049287,818 | 38397,906 | 4,947 | 3,182 | TRUE |
| ITPR3 | 129694,779 | 5277,062 | 4,805 | 1,453 | TRUE |
| PRRC2C | 932928,441 | 45420,855 | 4,554 | 1,704 | TRUE |
| TUBGCP2 | 342531,172 | 18363,398 | 4,433 | 3,986 | TRUE |
| PRDX2 | 2388616,952 | 128384,708 | 4,422 | 3,633 | TRUE |
| MZT2A | 222847,723 | 11592,247 | 4,417 | 1,404 | TRUE |
| PRDX1 | 13070579,911 | 707632,867 | 4,405 | 2,683 | TRUE |
| SPTAN1 | 1524853,613 | 96777,548 | 4,176 | 1,955 | TRUE |
| SPTBN1 | 1958640,818 | 128509,730 | 4,126 | 2,130 | TRUE |
| TUBGCP3 | 315306,901 | 21551,289 | 4,061 | 3,235 | TRUE |
| TUBGCP4 | 77216,079 | 5381,035 | 4,048 | 3,575 | TRUE |
| AHNAK2 | 130815,611 | 9634,860 | 3,973 | 5,450 | TRUE |
| PHLDB2 | 37366,813 | 2879,346 | 3,926 | 4,061 | TRUE |
| HSD17B10 | 248278,719 | 20107,394 | 3,889 | 3,593 | TRUE |
| KIF23 | 55447,948 | 4457,298 | 3,818 | 3,559 | TRUE |
| SOGA1 | 67528,503 | 5872,673 | 3,724 | 3,550 | TRUE |
| TRIM68 | 172581,503 | 17199,229 | 3,545 | 3,565 | TRUE |
| GPHN | 45821,365 | 5094,464 | 3,379 | 2,448 | TRUE |
| SOX9 | 198087,718 | 22291,078 | 3,343 | 2,839 | TRUE |
| ZNF346 | 278407,006 | 31303,680 | 3,317 | 2,233 | TRUE |
| CTR9 | 41174,649 | 4887,774 | 3,272 | 4,049 | TRUE |
| SOS1 | 15747,466 | 1921,571 | 3,244 | 4,132 | TRUE |
| KAT7 | 32001,248 | 4287,613 | 3,118 | 3,467 | TRUE |
| PARD3 | 12970,223 | 1787,873 | 3,058 | 4,220 | TRUE |
| PRRC2A | 26118,600 | 3830,605 | 2,974 | 3,322 | TRUE |
| PRRC2B | 1376947,196 | 213910,865 | 2,890 | 3,482 | TRUE |
| CDK11A | 17006,296 | 2798,795 | 2,845 | 1,737 | TRUE |
| JADE1 | 17002,495 | 2762,611 | 2,835 | 2,677 | TRUE |
| MRPS14 | 256708,709 | 44417,624 | 2,730 | 3,318 | TRUE |

| | | | | | |
|---|---|---|---|---|---|
| C4A | 8495,429 | 1583,662 | 2,627 | 2,941 | TRUE |
| LUZP1 | 9602,092 | 1829,723 | 2,626 | 2,257 | TRUE |
| PAF1 | 129473,400 | 24125,010 | 2,618 | 1,786 | TRUE |
| RACGAP1 | 21674,328 | 4106,584 | 2,598 | 2,327 | TRUE |
| LRFN4 | 24715,983 | 5426,558 | 2,385 | 2,112 | TRUE |
| TUBGCP5 | 21028,794 | 4821,891 | 2,346 | 2,651 | TRUE |
| SCAMP3 | 79529,128 | 18256,463 | 2,345 | 2,330 | TRUE |
| RCOR3 | 18385,839 | 4267,274 | 2,287 | 2,202 | TRUE |
| C9orf78 | 85972,679 | 20491,489 | 2,266 | 2,275 | TRUE |
| RPL5 | 9500375,843 | 2346453,098 | 2,221 | 3,901 | TRUE |
| FXR1 | 140116,793 | 36013,244 | 2,169 | 3,686 | TRUE |
| EXOSC10 | 7890,116 | 2705,502 | 1,767 | 1,801 | TRUE |
| RPL11 | 9006892,267 | 3288018,800 | 1,657 | 4,569 | TRUE |
| TUBGCP6 | 4467,206 | 1669,710 | 1,625 | 2,039 | TRUE |
| DNMBP | 4399,108 | 1705,531 | 1,580 | 1,712 | TRUE |
| DNAJC13 | 14767,273 | 5655,828 | 1,576 | 2,182 | TRUE |
| ZC3H14 | 8599,239 | 3747,036 | 1,411 | 1,972 | TRUE |
| BMP2K | 71465,720 | 31126,197 | 1,391 | 1,621 | TRUE |
| SQSTM1 | 52823,025 | 25894,060 | 1,256 | 2,482 | TRUE |
| HDAC2 | 32370,363 | 15739,581 | 1,255 | 1,531 | TRUE |
| RPF2 | 82940,131 | 42480,151 | 1,169 | 3,574 | TRUE |
| LDHA | 68772,875 | 163878,125 | -1,004 | 1,364 | TRUE |
| GAPDH | 957349,000 | 2310516,140 | -1,041 | 2,181 | TRUE |
| RGPD3 | 2400,673 | 5843,588 | -1,067 | 1,346 | TRUE |
| SFN | 15225,426 | 37544,196 | -1,117 | 1,459 | TRUE |
| RANBP2 | 7127,764 | 19878,304 | -1,266 | 2,058 | TRUE |
| ALDOA | 9512,108 | 27070,805 | -1,300 | 1,576 | TRUE |
| PGAM1 | 10434,777 | 30113,892 | -1,304 | 1,451 | TRUE |
| STMN2 | 10409,602 | 31284,037 | -1,368 | 1,442 | TRUE |
| CALCOCO2 | 7320,057 | 24663,461 | -1,538 | 1,433 | TRUE |
| STMN1 | 23913,009 | 84603,492 | -1,558 | 1,462 | TRUE |
| ENO3 | 9595,423 | 33993,227 | -1,600 | 1,771 | TRUE |
| CCT2 | 8982,101 | 34114,157 | -1,658 | 1,713 | TRUE |
| XPO1 | 8010,271 | 31691,846 | -1,736 | 1,977 | TRUE |
| CCT3 | 5523,354 | 22936,495 | -1,784 | 1,465 | TRUE |
| PSMB4 | 26897,383 | 104876,405 | -1,791 | 1,973 | TRUE |
| DPYSL2 | 2055,188 | 8464,786 | -1,810 | 1,308 | TRUE |
| FLII | 400812,374 | 1653597,758 | -1,867 | 1,591 | TRUE |
| EEF1A2 | 10063,418 | 47594,022 | -1,970 | 1,385 | TRUE |
| PGK1 | 7003,894 | 33237,950 | -1,973 | 1,567 | TRUE |
| IGKV2-30 | 23773,838 | 108028,667 | -1,974 | 1,883 | TRUE |
| MDH1 | 5367,063 | 26185,330 | -2,082 | 2,545 | TRUE |
| ACTR1A | 3406,466 | 17765,397 | -2,194 | 2,091 | TRUE |

| | | | | | |
|---|---|---|---|---|---|
| PARP9 | 1650,663 | 11071,256 | -2,528 | 2,344 | TRUE |
| CCT4 | 2826,150 | 19093,616 | -2,529 | 2,459 | TRUE |
| IGKV2D-24 | 23773,838 | 162276,843 | -2,552 | 1,661 | TRUE |
| ITGAV | 4030,530 | 34909,802 | -2,896 | 2,723 | TRUE |
| ARNT2 | 3602,097 | 31194,292 | -2,901 | 1,685 | TRUE |
| IGHV1-45 | 14858,649 | 136181,263 | -2,989 | 1,945 | TRUE |
| PNMA2 | 45171,183 | 483746,258 | -3,176 | 1,873 | TRUE |
| NFIB | 2904,498 | 87628,849 | -4,691 | 2,246 | TRUE |
| IGKV2-29 | 38461,933 | 3466143,056 | -6,359 | 2,428 | TRUE |
| OGDHL | 308804,205 | 10246,134 | 5,085 | 1,190 | FALSE |
| MRPL52 | 518301,771 | 119637,667 | 2,351 | 1,062 | FALSE |
| ZMYM3 | 8877,602 | 2192,775 | 2,194 | 1,267 | FALSE |
| MACF1 | 1218,005 | 318,540 | 2,167 | 0,925 | FALSE |
| WDR5 | 28332,756 | 10471,159 | 1,624 | 0,910 | FALSE |
| GABPA | 17234,649 | 6606,244 | 1,568 | 0,893 | FALSE |
| DDX20 | 11663,557 | 4524,137 | 1,562 | 0,981 | FALSE |
| ZNF608 | 105719,539 | 40485,988 | 1,560 | 0,572 | FALSE |
| BAZ1B | 41863,085 | 20042,774 | 1,429 | 0,671 | FALSE |
| PPP1CA | 313280,183 | 141014,992 | 1,354 | 1,004 | FALSE |
| CTCF | 43222,941 | 20604,224 | 1,322 | 0,607 | FALSE |
| PPP1R9B | 58093,384 | 29847,149 | 1,199 | 0,536 | FALSE |
| ITPR1 | 3647,969 | 1765,304 | 1,178 | 0,927 | FALSE |
| EXOSC6 | 14260,214 | 7425,177 | 1,134 | 0,715 | FALSE |
| FTSJ3 | 10228,928 | 5545,022 | 1,123 | 1,218 | FALSE |
| GNL3 | 22328,335 | 11842,981 | 1,114 | 0,800 | FALSE |
| MPHOSPH10 | 11347,100 | 6274,004 | 1,105 | 0,482 | FALSE |
| CSN3 | 860116,643 | 458435,274 | 1,031 | 0,432 | FALSE |
| PLEKHG3 | 19409,453 | 11448,123 | 0,997 | 0,556 | FALSE |
| RRS1 | 64297,968 | 37255,567 | 0,995 | 2,601 | FALSE |
| RPL22L1 | 57240,721 | 35311,306 | 0,961 | 0,901 | FALSE |
| GRWD1 | 70881,579 | 44722,731 | 0,948 | 0,792 | FALSE |
| GBE1 | 27056,405 | 16629,767 | 0,945 | 0,294 | FALSE |
| GEMIN4 | 5944,195 | 3589,346 | 0,930 | 1,169 | FALSE |
| ACTN1 | 2339886,172 | 1429880,368 | 0,926 | 0,584 | FALSE |
| ACIN1 | 11471,116 | 6901,092 | 0,901 | 0,430 | FALSE |
| ACTN4 | 70034,816 | 44587,019 | 0,887 | 0,587 | FALSE |
| KLHDC10 | 19656,831 | 12254,155 | 0,884 | 0,437 | FALSE |
| WDR12 | 54296,709 | 35028,237 | 0,833 | 1,166 | FALSE |
| CDKN2AIPNL | 185138,136 | 120300,211 | 0,828 | 0,410 | FALSE |
| HDAC1 | 13527,988 | 8606,627 | 0,828 | 0,669 | FALSE |
| NUDT1 | 101845,860 | 65429,647 | 0,816 | 0,363 | FALSE |
| PGAM5 | 298291,697 | 195325,883 | 0,805 | 1,091 | FALSE |
| RSL1D1 | 625674,962 | 414367,141 | 0,805 | 1,862 | FALSE |

| | | | | | |
|---|---|---|---|---|---|
| CSNK2A1 | 67046,035 | 45071,355 | 0,780 | 1,071 | FALSE |
| SPATS2L | 285580,579 | 192170,131 | 0,777 | 1,970 | FALSE |
| ACTC1 | 776777,797 | 521190,319 | 0,776 | 1,872 | FALSE |
| RIOX1 | 7437,251 | 5053,359 | 0,774 | 0,603 | FALSE |
| MRPL55 | 37702,722 | 25218,016 | 0,769 | 0,718 | FALSE |
| DBN1 | 98045,373 | 67549,381 | 0,763 | 0,483 | FALSE |
| ZC3HAV1 | 279193,419 | 190064,616 | 0,761 | 2,023 | FALSE |
| SYNPO | 240994,558 | 165843,356 | 0,749 | 0,463 | FALSE |
| SMN1 | 184554,292 | 129121,356 | 0,734 | 2,902 | FALSE |
| CAPZB | 11386,535 | 7785,437 | 0,730 | 0,652 | FALSE |
| NCBP3 | 8438,873 | 5950,345 | 0,726 | 0,649 | FALSE |
| CCNT1 | 5322,043 | 3669,061 | 0,719 | 0,442 | FALSE |
| LLPH | 58671,249 | 40823,907 | 0,701 | 0,732 | FALSE |
| ALB | 37739,065 | 27369,259 | 0,668 | 0,577 | FALSE |
| MROH8 | 24503,583 | 18521,289 | 0,649 | 0,699 | FALSE |
| YLPM1 | 1247232,604 | 911206,541 | 0,644 | 0,355 | FALSE |
| THRAP3 | 26873,094 | 19639,744 | 0,643 | 0,685 | FALSE |
| PPP1CB | 265750,636 | 195693,092 | 0,641 | 0,404 | FALSE |
| DYNLL1 | 438324,304 | 316208,417 | 0,632 | 0,318 | FALSE |
| CSN1S1 | 5989730,778 | 4321026,556 | 0,624 | 0,729 | FALSE |
| LGB | 193040,828 | 139392,247 | 0,614 | 0,260 | FALSE |
| ILK | 21123,818 | 15944,027 | 0,614 | 0,512 | FALSE |
| CALM1 | 333258,384 | 253954,607 | 0,613 | 0,941 | FALSE |
| MISP | 16433,769 | 12608,074 | 0,612 | 0,569 | FALSE |
| Tryspin | 111503639,111 | 81673994,667 | 0,611 | 0,809 | FALSE |
| BOP1 | 52980,853 | 40735,286 | 0,610 | 0,723 | FALSE |
| MRPL48 | 59485,293 | 44574,938 | 0,603 | 0,736 | FALSE |
| DDX27 | 68367,326 | 52976,332 | 0,599 | 0,654 | FALSE |
| UTP15 | 9069,249 | 6913,145 | 0,597 | 0,361 | FALSE |
| EXOSC9 | 20828,397 | 15839,422 | 0,595 | 0,795 | FALSE |
| SRSF4 | 37663,696 | 28542,391 | 0,594 | 1,125 | FALSE |
| BCLAF1 | 10551,559 | 7750,526 | 0,590 | 0,327 | FALSE |
| PKP4 | 10119,085 | 7433,446 | 0,586 | 0,350 | FALSE |
| MTREX | 55735,087 | 42432,683 | 0,579 | 0,649 | FALSE |
| NOL6 | 6389,104 | 4896,519 | 0,576 | 0,391 | FALSE |
| KRT6B | 101685,516 | 77419,171 | 0,572 | 0,413 | FALSE |
| EIF3E | 27933,282 | 21936,947 | 0,561 | 1,851 | FALSE |
| SCAF11 | 4784,510 | 3509,127 | 0,560 | 0,247 | FALSE |
| HMG20A | 97584,838 | 75057,838 | 0,553 | 0,225 | FALSE |
| MRPL16 | 29902,589 | 23397,204 | 0,547 | 0,518 | FALSE |
| LIMA1 | 8787,911 | 7191,576 | 0,544 | 0,217 | FALSE |
| ZCCHC8 | 91949,642 | 72292,804 | 0,544 | 0,860 | FALSE |
| GTF3C3 | 34879,695 | 25135,775 | 0,540 | 0,174 | FALSE |

| | | | | | |
|---|---|---|---|---|---|
| CSN1S2 | 941865,271 | 728613,229 | 0,539 | 0,532 | FALSE |
| MRPL41 | 3303282,222 | 2671965,333 | 0,538 | 0,983 | FALSE |
| NIFK | 116866,265 | 92713,637 | 0,531 | 0,852 | FALSE |
| PTMA | 442868,791 | 365115,923 | 0,531 | 0,717 | FALSE |
| PKP1 | 8241,235 | 6898,575 | 0,528 | 0,424 | FALSE |
| C1QBP | 553863,381 | 443853,619 | 0,527 | 1,951 | FALSE |
| CDKN2AIP | 249364,018 | 200074,369 | 0,523 | 0,989 | FALSE |
| Cont Bovin | 312201,099 | 244941,183 | 0,523 | 0,711 | FALSE |
| MRPL2 | 45170,797 | 36822,076 | 0,496 | 1,341 | FALSE |
| RNPS1 | 47554,132 | 38910,057 | 0,494 | 0,498 | FALSE |
| H2AC18 | 1002467,530 | 856133,190 | 0,483 | 0,204 | FALSE |
| NOP2 | 271539,350 | 224407,567 | 0,482 | 2,516 | FALSE |
| EIF3K | 28377,775 | 23640,684 | 0,472 | 0,566 | FALSE |
| TBL3 | 3805,001 | 3194,289 | 0,471 | 0,351 | FALSE |
| MYO1C | 106607,226 | 89805,490 | 0,467 | 0,517 | FALSE |
| TJP1 | 3253,817 | 2826,588 | 0,459 | 0,409 | FALSE |
| RPUSD4 | 30503,561 | 25617,706 | 0,454 | 0,452 | FALSE |
| LBR | 13893,742 | 11643,798 | 0,446 | 0,633 | FALSE |
| PPP1CC | 195024,176 | 165108,259 | 0,442 | 0,357 | FALSE |
| SNRPA | 6020031,795 | 5134806,051 | 0,429 | 0,286 | FALSE |
| H1-4 | 175827,806 | 150181,972 | 0,423 | 0,382 | FALSE |
| BCL7B | 364285,482 | 307612,489 | 0,423 | 0,133 | FALSE |
| CNIH4 | 147349,005 | 127932,738 | 0,417 | 0,403 | FALSE |
| MRPL4 | 447693,796 | 377795,519 | 0,415 | 0,436 | FALSE |
| HSPA6 | 560505,248 | 486469,205 | 0,414 | 0,541 | FALSE |
| RBM45 | 13481,043 | 11380,062 | 0,413 | 0,549 | FALSE |
| RPL10A | 1606301,821 | 1380221,410 | 0,413 | 1,284 | FALSE |
| WDR61 | 185697,650 | 162415,688 | 0,410 | 0,463 | FALSE |
| MAGOHB | 213755,530 | 182490,875 | 0,409 | 0,725 | FALSE |
| HEATR1 | 1218,310 | 1097,655 | 0,408 | 0,199 | FALSE |
| MRPL47 | 112986,360 | 96736,586 | 0,406 | 0,857 | FALSE |
| TERF2IP | 7964,670 | 6906,528 | 0,406 | 0,259 | FALSE |
| Cont Bovin | 829019,137 | 711965,667 | 0,403 | 0,566 | FALSE |
| NUP42 | 75740,547 | 66543,378 | 0,400 | 0,307 | FALSE |
| MRPL40 | 60054,314 | 52914,368 | 0,398 | 0,403 | FALSE |
| SKIV2L | 11095,700 | 9714,476 | 0,391 | 0,230 | FALSE |
| RPL14 | 1932586,778 | 1723041,278 | 0,387 | 1,355 | FALSE |
| SNRPB2 | 211485,602 | 185624,750 | 0,386 | 1,441 | FALSE |
| SSB | 187922,190 | 166544,747 | 0,384 | 1,469 | FALSE |
| KRT16 | 1203831,717 | 1073642,222 | 0,381 | 0,361 | FALSE |
| CDK9 | 56428,630 | 50355,357 | 0,381 | 0,421 | FALSE |
| DLAT | 18501,138 | 16371,335 | 0,379 | 0,547 | FALSE |
| MRPL18 | 62391,167 | 53786,708 | 0,376 | 0,352 | FALSE |

| | | | | | |
|---|---|---|---|---|---|
| RBM28 | 42044,026 | 37449,791 | 0,374 | 2,162 | FALSE |
| TFG | 112987611,022 | 98946409,244 | 0,374 | 1,416 | FALSE |
| GLUD1 | 10759,957 | 9507,542 | 0,373 | 0,491 | FALSE |
| RPL19 | 1315642,667 | 1172629,815 | 0,373 | 1,129 | FALSE |
| SART3 | 143656,391 | 126437,261 | 0,368 | 0,919 | FALSE |
| ATXN2 | 140444,676 | 125508,689 | 0,365 | 0,465 | FALSE |
| ASPH | 6360,986 | 5634,984 | 0,364 | 0,332 | FALSE |
| BAG4 | 4590807,083 | 4055851,583 | 0,364 | 1,387 | FALSE |
| LOXL1 | 24084,417 | 21241,059 | 0,364 | 0,557 | FALSE |
| TIAL1 | 19095901,193 | 16712574,596 | 0,363 | 0,716 | FALSE |
| AATF | 40385,299 | 36333,259 | 0,362 | 0,850 | FALSE |
| RBM34 | 42899,535 | 38225,148 | 0,361 | 0,517 | FALSE |
| FAM168A | 1945991,444 | 1723340,056 | 0,357 | 0,661 | FALSE |
| C7orf50 | 169959,299 | 154510,694 | 0,355 | 0,722 | FALSE |
| RCN2 | 128467,922 | 115943,775 | 0,354 | 1,441 | FALSE |
| MRPL23 | 234189,720 | 210762,432 | 0,353 | 1,597 | FALSE |
| NUP155 | 19370,295 | 17569,930 | 0,352 | 0,810 | FALSE |
| QARS1 | 13691,748 | 12517,135 | 0,352 | 0,312 | FALSE |
| FLNB | 4513,938 | 4530,025 | 0,348 | 0,115 | FALSE |
| RBM4B | 416776,051 | 378855,319 | 0,347 | 0,277 | FALSE |
| XRN1 | 9077,872 | 8142,192 | 0,346 | 0,843 | FALSE |
| EXOSC1 | 19516,828 | 17899,470 | 0,346 | 0,225 | FALSE |
| DAP3 | 2000532,222 | 1794652,278 | 0,343 | 0,496 | FALSE |
| RFC3 | 31623,387 | 28570,429 | 0,342 | 0,956 | FALSE |
| CPEB4 | 112730,542 | 100475,392 | 0,342 | 0,564 | FALSE |
| MAP1B | 1928,230 | 1777,933 | 0,341 | 0,206 | FALSE |
| MRPL28 | 47822,576 | 42829,025 | 0,341 | 0,411 | FALSE |
| LOXL4 | 12131,977 | 11030,762 | 0,339 | 0,624 | FALSE |
| MRPL14 | 89705,636 | 81631,277 | 0,337 | 0,959 | FALSE |
| MYO1B | 9189,364 | 8523,553 | 0,337 | 0,175 | FALSE |
| RRP8 | 16214,901 | 15276,687 | 0,334 | 0,148 | FALSE |
| DDX18 | 36261,683 | 33833,394 | 0,331 | 0,344 | FALSE |
| GANAB | 6694,361 | 6216,330 | 0,325 | 0,259 | FALSE |
| YTHDC2 | 2532,650 | 2244,486 | 0,324 | 0,166 | FALSE |
| AGO2 | 1495646,870 | 1372682,621 | 0,321 | 2,517 | FALSE |
| MRPS22 | 2186844,160 | 1997663,947 | 0,321 | 0,483 | FALSE |
| RBM4 | 150346,295 | 142198,188 | 0,320 | 0,172 | FALSE |
| TNRC6C | 608764,982 | 555463,263 | 0,317 | 0,509 | FALSE |
| ETF1 | 6030,110 | 5582,880 | 0,316 | 0,178 | FALSE |
| ARHGEF18 | 79872,242 | 74319,523 | 0,316 | 0,832 | FALSE |
| NAT10 | 24034,901 | 22500,253 | 0,315 | 0,781 | FALSE |
| DHX15 | 214778,993 | 198693,873 | 0,313 | 3,006 | FALSE |
| RPS25 | 3559201,889 | 3304309,778 | 0,309 | 1,146 | FALSE |

| | | | | | |
|---|---|---|---|---|---|
| TUBB2B | 121512,892 | 118162,037 | 0,309 | 0,179 | FALSE |
| BUB3 | 32021,358 | 30000,401 | 0,307 | 0,429 | FALSE |
| TNRC6B | 3227416,061 | 2970946,667 | 0,306 | 1,100 | FALSE |
| TNRC6A | 1076571,223 | 997060,190 | 0,305 | 1,790 | FALSE |
| Cont Bonvin | 25166,190 | 23275,672 | 0,302 | 0,093 | FALSE |
| CSN2 | 5400637,333 | 4919833,037 | 0,301 | 0,478 | FALSE |
| RPL27 | 2471400,611 | 2306908,500 | 0,301 | 0,761 | FALSE |
| NCBP2 | 237795,231 | 225377,587 | 0,300 | 0,643 | FALSE |
| RAMAC | 5500843,636 | 5128706,182 | 0,298 | 0,868 | FALSE |
| YWHAH | 8708,697 | 8286,665 | 0,296 | 0,170 | FALSE |
| IGF2BP3 | 84151,443 | 79415,094 | 0,294 | 0,237 | FALSE |
| PUM3 | 24831,507 | 23620,649 | 0,291 | 0,600 | FALSE |
| FUS | 109897281,255 | 102297070,431 | 0,291 | 0,679 | FALSE |
| DIMT1 | 35821,185 | 33938,937 | 0,288 | 0,451 | FALSE |
| EIF3C | 12816,501 | 12009,036 | 0,286 | 0,423 | FALSE |
| NOL10 | 57738,947 | 54843,122 | 0,286 | 0,316 | FALSE |
| CCAR1 | 2488,638 | 2400,141 | 0,285 | 0,138 | FALSE |
| PRDX4 | 65598,376 | 62344,876 | 0,281 | 0,539 | FALSE |
| MRPS18C | 579273,438 | 542025,792 | 0,281 | 0,303 | FALSE |
| RPL21 | 1088427,042 | 1039607,188 | 0,280 | 1,852 | FALSE |
| KRT6A | 219436,167 | 208840,329 | 0,280 | 0,205 | FALSE |
| SNRPD1 | 3895647,533 | 3802595,200 | 0,280 | 0,371 | FALSE |
| RPL4 | 1343832,469 | 1283926,494 | 0,276 | 0,997 | FALSE |
| RPS19 | 3199740,722 | 3038500,389 | 0,272 | 1,238 | FALSE |
| DDX21 | 477765,822 | 455774,938 | 0,271 | 0,481 | FALSE |
| RAI1 | 1406,517 | 1356,896 | 0,271 | 0,134 | FALSE |
| SEC16A | 137095,295 | 129219,309 | 0,271 | 0,302 | FALSE |
| HNRNPH2 | 25944369,641 | 24347933,538 | 0,271 | 0,569 | FALSE |
| SMC2 | 1778,702 | 1711,043 | 0,270 | 0,136 | FALSE |
| SNRPB | 1064337,510 | 1026110,471 | 0,270 | 0,411 | FALSE |
| PSMC3 | 10425,556 | 10098,047 | 0,269 | 0,179 | FALSE |
| NPM1 | 546052,559 | 536180,510 | 0,268 | 0,339 | FALSE |
| MAGED2 | 16340,745 | 15829,964 | 0,268 | 0,351 | FALSE |
| RPL12 | 2439018,424 | 2329025,333 | 0,268 | 1,141 | FALSE |
| ALDOC | 7861,250 | 7503,952 | 0,266 | 0,172 | FALSE |
| MAGED1 | 72081,811 | 68287,255 | 0,264 | 0,517 | FALSE |
| RPL7 | 2724289,917 | 2615731,583 | 0,263 | 1,723 | FALSE |
| MRPL46 | 18248,835 | 17471,043 | 0,263 | 0,242 | FALSE |
| RBM7 | 415593,810 | 395081,464 | 0,261 | 0,716 | FALSE |
| MRTO4 | 211106,224 | 201753,849 | 0,261 | 0,329 | FALSE |
| EIF6 | 130523,402 | 126760,115 | 0,261 | 0,565 | FALSE |
| MCRIP1 | 21804,934 | 20706,827 | 0,258 | 0,156 | FALSE |
| RPL10 | 1753611,333 | 1673550,389 | 0,257 | 0,685 | FALSE |

| | | | | | |
|---|---|---|---|---|---|
| SMARCA5 | 10390,649 | 10017,202 | 0,257 | 0,726 | FALSE |
| MRPS12 | 560574,719 | 542376,802 | 0,257 | 0,918 | FALSE |
| SRSF3 | 1790593,815 | 1738355,296 | 0,257 | 0,561 | FALSE |
| DUSP14 | 20984,880 | 20269,429 | 0,255 | 0,280 | FALSE |
| GADD45GIP1 | 241561,750 | 231941,295 | 0,253 | 0,314 | FALSE |
| XRN2 | 1906359,908 | 1834968,276 | 0,253 | 0,740 | FALSE |
| AKAP8 | 2929950,495 | 2783074,280 | 0,251 | 0,156 | FALSE |
| RTCB | 3194393,195 | 3083795,586 | 0,251 | 0,453 | FALSE |
| MRPS7 | 1821370,316 | 1739856,351 | 0,251 | 0,345 | FALSE |
| MACROH2A1 | 100126,868 | 96314,165 | 0,251 | 0,461 | FALSE |
| DNAJB2 | 29917,242 | 28958,191 | 0,250 | 0,472 | FALSE |
| RPL15 | 830711,929 | 812966,643 | 0,248 | 0,519 | FALSE |
| PHF14 | 134239,541 | 128360,912 | 0,244 | 0,089 | FALSE |
| SNRPD2 | 4728300,917 | 4641375,500 | 0,244 | 0,275 | FALSE |
| P4HA1 | 9891,423 | 9729,944 | 0,242 | 0,286 | FALSE |
| MRPL45 | 207553,614 | 201267,034 | 0,240 | 0,407 | FALSE |
| MSH6 | 22633,354 | 22198,429 | 0,238 | 0,412 | FALSE |
| KPNA7 | 65579,070 | 63163,280 | 0,237 | 0,158 | FALSE |
| RTRAF | 4337818,593 | 4219886,222 | 0,237 | 0,956 | FALSE |
| RPS13 | 2788875,394 | 2720603,394 | 0,235 | 1,000 | FALSE |
| MRPL38 | 185168,600 | 180013,400 | 0,234 | 0,356 | FALSE |
| PWP1 | 23356,909 | 22585,474 | 0,234 | 0,132 | FALSE |
| CORO2A | 133527,769 | 129662,488 | 0,233 | 0,289 | FALSE |
| XRCC6 | 43041,747 | 41903,054 | 0,231 | 0,517 | FALSE |
| IREB2 | 9519,814 | 9170,611 | 0,230 | 0,258 | FALSE |
| MRPL44 | 345304,400 | 334981,925 | 0,229 | 0,277 | FALSE |
| RBMS1 | 2621670,980 | 2559956,784 | 0,228 | 0,368 | FALSE |
| TOP2B | 3514,171 | 3555,240 | 0,227 | 0,104 | FALSE |
| SNRPC | 2031868,429 | 2033258,048 | 0,226 | 0,314 | FALSE |
| HNRNPH3 | 72409015,467 | 70408802,133 | 0,225 | 0,625 | FALSE |
| MRPL19 | 220428,875 | 214646,910 | 0,224 | 0,263 | FALSE |
| EIF3L | 29593,128 | 29753,825 | 0,224 | 0,289 | FALSE |
| MRPS35 | 104346,083 | 103315,237 | 0,224 | 0,878 | FALSE |
| RPL35A | 2207142,074 | 2170418,667 | 0,223 | 1,031 | FALSE |
| RPS8 | 2238018,121 | 2214276,364 | 0,222 | 2,859 | FALSE |
| TAX1BP1 | 12226,897 | 12075,194 | 0,220 | 0,219 | FALSE |
| H1-2 | 230711,927 | 226920,556 | 0,219 | 0,230 | FALSE |
| CASC3 | 43139,753 | 42355,836 | 0,218 | 0,557 | FALSE |
| SSBP2 | 35939,779 | 35027,745 | 0,218 | 0,083 | FALSE |
| KRT13 | 62898,671 | 62449,801 | 0,217 | 0,103 | FALSE |
| ATXN2L | 102064,593 | 101303,304 | 0,217 | 1,342 | FALSE |
| HNRNPUL1 | 16614821,901 | 16135914,213 | 0,215 | 0,350 | FALSE |
| ISY1 | 59169,601 | 58321,744 | 0,214 | 0,600 | FALSE |

| | | | | | |
|---|---|---|---|---|---|
| RPS27L | 1901444,467 | 1879015,133 | 0,212 | 0,985 | FALSE |
| PHB | 40900,594 | 40968,858 | 0,212 | 0,225 | FALSE |
| MRPS16 | 1597800,333 | 1580367,933 | 0,210 | 0,512 | FALSE |
| DHX57 | 3932,431 | 3971,117 | 0,209 | 0,205 | FALSE |
| PLAT | 6552,862 | 6468,693 | 0,209 | 0,181 | FALSE |
| PHB2 | 127422,817 | 129799,521 | 0,209 | 0,120 | FALSE |
| KPRP | 60548,332 | 61263,225 | 0,208 | 0,129 | FALSE |
| RPL13 | 1557805,564 | 1560524,769 | 0,207 | 0,494 | FALSE |
| IGK | 117255,853 | 114617,785 | 0,206 | 0,171 | FALSE |
| ZNF638 | 48504,723 | 48495,021 | 0,204 | 0,533 | FALSE |
| SSRP1 | 28396,402 | 28472,937 | 0,203 | 0,212 | FALSE |
| ZNF609 | 298828,995 | 294505,177 | 0,200 | 0,118 | FALSE |
| YTHDF1 | 4373866,087 | 4351365,333 | 0,199 | 0,533 | FALSE |
| CPSF7 | 18753,457 | 18949,414 | 0,198 | 0,296 | FALSE |
| GAR1 | 106369,218 | 107277,936 | 0,198 | 0,529 | FALSE |
| YWHAG | 21455,313 | 21395,921 | 0,198 | 0,149 | FALSE |
| DHX30 | 670166,730 | 671490,317 | 0,198 | 0,240 | FALSE |
| RPL7A | 2095548,769 | 2108467,333 | 0,197 | 1,292 | FALSE |
| RPS7 | 2312651,800 | 2330615,733 | 0,197 | 0,960 | FALSE |
| PPHLN1 | 51434,645 | 51855,103 | 0,196 | 0,911 | FALSE |
| MRPL9 | 229691,254 | 230133,439 | 0,196 | 0,335 | FALSE |
| KCTD6 | 24717,355 | 24668,839 | 0,195 | 0,217 | FALSE |
| H2BC3 | 379201,924 | 381790,409 | 0,192 | 0,435 | FALSE |
| H1-10 | 472899,931 | 477036,764 | 0,192 | 0,548 | FALSE |
| SMARCC2 | 1718016,842 | 1688724,468 | 0,191 | 0,040 | FALSE |
| DUSP11 | 145275,987 | 144128,641 | 0,189 | 0,375 | FALSE |
| MRPL1 | 186163,774 | 185302,885 | 0,187 | 0,416 | FALSE |
| MRPL17 | 73152,696 | 73812,601 | 0,186 | 0,256 | FALSE |
| TFB1M | 28763,177 | 29217,640 | 0,185 | 0,491 | FALSE |
| NOP56 | 352884,685 | 356938,759 | 0,184 | 0,284 | FALSE |
| SRSF6 | 189186,858 | 190367,908 | 0,182 | 0,331 | FALSE |
| KRT3 | 6112,871 | 6162,470 | 0,181 | 0,130 | FALSE |
| AKAP8L | 116874,848 | 117180,565 | 0,181 | 0,297 | FALSE |
| MRPS17 | 2485739,750 | 2481304,667 | 0,179 | 0,239 | FALSE |
| MRPL20 | 53990,924 | 54761,449 | 0,179 | 0,231 | FALSE |
| MRPL43 | 210039,115 | 213458,929 | 0,179 | 0,319 | FALSE |
| PRPF8 | 179672,824 | 182612,883 | 0,178 | 1,045 | FALSE |
| RALYL | 1371026,167 | 1399965,917 | 0,177 | 0,164 | FALSE |
| NIP7 | 68321,402 | 68979,325 | 0,177 | 0,184 | FALSE |
| GALC | 132997,360 | 133827,118 | 0,176 | 0,293 | FALSE |
| CHERP | 400665,171 | 407440,667 | 0,176 | 0,494 | FALSE |
| SNRPD3 | 3485736,444 | 3577643,926 | 0,176 | 0,247 | FALSE |
| H1-0 | 42474,516 | 43777,632 | 0,176 | 0,141 | FALSE |

| | | | | | |
|---|---|---|---|---|---|
| RBM39 | 112870,417 | 115903,080 | 0,174 | 0,475 | FALSE |
| MRPS31 | 75880,355 | 76999,548 | 0,174 | 0,172 | FALSE |
| UTP18 | 6859,762 | 7091,807 | 0,173 | 0,095 | FALSE |
| MRPL3 | 215979,819 | 219442,979 | 0,172 | 0,564 | FALSE |
| TUBA4B | 115735,288 | 117798,933 | 0,172 | 0,553 | FALSE |
| YWHAZ | 168902,118 | 175113,939 | 0,169 | 0,424 | FALSE |
| RBMS2 | 1404572,222 | 1443039,500 | 0,168 | 0,165 | FALSE |
| RPS6 | 1334201,462 | 1362839,462 | 0,167 | 0,625 | FALSE |
| AGO1 | 233737,393 | 238567,738 | 0,167 | 0,877 | FALSE |
| RPS3 | 4600257,825 | 4726639,298 | 0,164 | 1,142 | FALSE |
| ZCCHC3 | 35682,626 | 36866,067 | 0,163 | 0,386 | FALSE |
| BRIX1 | 35098,001 | 36839,561 | 0,160 | 0,315 | FALSE |
| PURA | 1142138,643 | 1174903,476 | 0,160 | 0,205 | FALSE |
| XRCC5 | 48473,518 | 50107,021 | 0,160 | 0,802 | FALSE |
| PARP1 | 4588,929 | 4708,834 | 0,160 | 0,121 | FALSE |
| EFTUD2 | 372650,642 | 381215,247 | 0,158 | 0,812 | FALSE |
| RBM5 | 8211,458 | 8552,136 | 0,158 | 0,146 | FALSE |
| KPNA3 | 83315,831 | 86456,688 | 0,158 | 0,426 | FALSE |
| KPNA2 | 1045236,213 | 1076468,747 | 0,156 | 0,286 | FALSE |
| RPSA | 5554321,244 | 5729518,222 | 0,156 | 1,168 | FALSE |
| HCFC1 | 4289,929 | 4500,767 | 0,155 | 0,157 | FALSE |
| RPS16 | 3816755,444 | 3933866,667 | 0,155 | 0,913 | FALSE |
| RCBTB1 | 119199,414 | 122065,064 | 0,154 | 0,101 | FALSE |
| TIA1 | 12505204,148 | 12784180,741 | 0,153 | 0,429 | FALSE |
| RPS4X | 2494931,238 | 2581714,222 | 0,150 | 0,760 | FALSE |
| AGO3 | 865352,000 | 893511,048 | 0,148 | 1,031 | FALSE |
| YWHAE | 40319,344 | 41657,940 | 0,147 | 0,098 | FALSE |
| KRI1 | 39545,357 | 41306,059 | 0,146 | 0,537 | FALSE |
| RPA3 | 217208,254 | 228540,121 | 0,144 | 0,110 | FALSE |
| RPL26 | 260919,849 | 277321,157 | 0,143 | 0,072 | FALSE |
| MRPS9 | 1725099,862 | 1783700,368 | 0,143 | 0,284 | FALSE |
| RPL30 | 2500041,708 | 2572410,583 | 0,143 | 0,227 | FALSE |
| NUP98 | 37226,437 | 38679,754 | 0,141 | 0,211 | FALSE |
| YBX1 | 13995070,095 | 14599751,619 | 0,141 | 0,243 | FALSE |
| H4C1 | 22593220,000 | 23255580,000 | 0,139 | 0,247 | FALSE |
| TCF20 | 610033,975 | 631115,673 | 0,139 | 0,151 | FALSE |
| DMD | 20512,380 | 21667,545 | 0,139 | 0,271 | FALSE |
| RPL23 | 2760599,333 | 2872643,926 | 0,139 | 0,387 | FALSE |
| MAGEB2 | 232034,300 | 246571,344 | 0,138 | 0,382 | FALSE |
| PLOD3 | 163758,746 | 172040,018 | 0,137 | 0,811 | FALSE |
| PRPF6 | 19004,083 | 20120,772 | 0,137 | 0,201 | FALSE |
| MRPS23 | 1888653,810 | 1966542,333 | 0,136 | 0,286 | FALSE |
| EWSR1 | 43840094,609 | 45536106,667 | 0,136 | 0,544 | FALSE |

| | | | | | |
|---|---|---|---|---|---|
| SELENOF | 21178,301 | 21911,974 | 0,136 | 0,086 | FALSE |
| RPLP0 | 3616876,593 | 3770759,333 | 0,136 | 0,641 | FALSE |
| ZNF326 | 2103902,745 | 2183878,941 | 0,135 | 0,282 | FALSE |
| HNRNPA2B1 | 207093734,400 | 215142980,267 | 0,135 | 0,462 | FALSE |
| EIF4A3 | 313403,494 | 328420,756 | 0,135 | 0,230 | FALSE |
| PHAX | 54308,967 | 57819,373 | 0,134 | 0,192 | FALSE |
| HSPA5 | 2182932,175 | 2258170,877 | 0,134 | 0,110 | FALSE |
| NOP58 | 234303,816 | 246027,621 | 0,133 | 0,393 | FALSE |
| SVIL | 6223,665 | 6676,582 | 0,132 | 0,063 | FALSE |
| FAU | 444140,784 | 476814,041 | 0,131 | 0,133 | FALSE |
| RPL38 | 716003,167 | 738553,250 | 0,130 | 0,169 | FALSE |
| DDX50 | 31095,563 | 32670,770 | 0,130 | 0,251 | FALSE |
| KLHL22 | 99586,245 | 103021,502 | 0,129 | 0,237 | FALSE |
| RBFOX3 | 69964,479 | 73488,473 | 0,127 | 0,248 | FALSE |
| RPL27A | 1166576,208 | 1247366,729 | 0,127 | 0,399 | FALSE |
| HELZ | 19854,228 | 20818,235 | 0,127 | 0,153 | FALSE |
| H2AC21 | 267334,437 | 278536,632 | 0,127 | 0,179 | FALSE |
| HSPA1A | 597215,231 | 625717,537 | 0,127 | 0,249 | FALSE |
| CDC40 | 105004,686 | 108912,000 | 0,126 | 0,312 | FALSE |
| DYNC1I2 | 12392,724 | 13288,817 | 0,125 | 0,077 | FALSE |
| RBM6 | 221709,160 | 234379,280 | 0,124 | 0,335 | FALSE |
| HNRNPDL | 35898820,638 | 37182366,609 | 0,124 | 0,136 | FALSE |
| USP9X | 67597,049 | 70985,901 | 0,121 | 0,098 | FALSE |
| DPF2 | 1786417,810 | 1844486,493 | 0,118 | 0,023 | FALSE |
| RPL13A | 703181,314 | 755223,745 | 0,118 | 0,379 | FALSE |
| PRPF3 | 7048,033 | 7613,980 | 0,115 | 0,088 | FALSE |
| KLHL12 | 13295,686 | 14387,035 | 0,115 | 0,117 | FALSE |
| MRPL58 | 116523,928 | 123018,242 | 0,114 | 0,120 | FALSE |
| DDOST | 18441,849 | 19395,543 | 0,113 | 0,185 | FALSE |
| LARP7 | 33784,681 | 35982,966 | 0,112 | 0,407 | FALSE |
| BAG2 | 64700,478 | 69725,205 | 0,111 | 0,193 | FALSE |
| MRPS27 | 1479190,713 | 1576618,437 | 0,110 | 0,220 | FALSE |
| SEC24B | 2365929,167 | 2499074,667 | 0,109 | 0,122 | FALSE |
| PPP1R9A | 2564,266 | 2723,889 | 0,108 | 0,048 | FALSE |
| KRR1 | 111117,246 | 120019,551 | 0,107 | 0,165 | FALSE |
| UBAP2L | 6709497,500 | 7138647,333 | 0,107 | 0,131 | FALSE |
| PUF60 | 106867,589 | 113735,667 | 0,106 | 0,394 | FALSE |
| CSTF1 | 582113,603 | 626335,698 | 0,106 | 0,137 | FALSE |
| ARID1B | 1025482,538 | 1077945,221 | 0,106 | 0,046 | FALSE |
| MRPS21 | 234248,284 | 253422,275 | 0,104 | 0,233 | FALSE |
| DDX56 | 5879,833 | 6323,846 | 0,104 | 0,055 | FALSE |
| P4HA2 | 24149,568 | 26769,637 | 0,103 | 0,045 | FALSE |
| POGZ | 5530,986 | 6012,760 | 0,103 | 0,051 | FALSE |

| | | | | | |
|---|---|---|---|---|---|
| N4BP2L2 | 3607,546 | 3920,479 | 0,102 | 0,050 | FALSE |
| MTMR3 | 27465,151 | 29123,502 | 0,100 | 0,133 | FALSE |
| MYEF2 | 42868,821 | 46050,795 | 0,099 | 0,162 | FALSE |
| HSPA8 | 4882737,586 | 5213354,378 | 0,097 | 0,165 | FALSE |
| PHF5A | 145013,525 | 154269,836 | 0,097 | 0,041 | FALSE |
| PDCD6 | 13686243,333 | 14606680,444 | 0,096 | 0,182 | FALSE |
| RBMS3 | 599378,294 | 647717,588 | 0,096 | 0,084 | FALSE |
| HNRNPH1 | 17851231,590 | 19050875,897 | 0,095 | 0,301 | FALSE |
| SF3B1 | 236612,813 | 251948,457 | 0,095 | 0,038 | FALSE |
| MRPL50 | 511327,068 | 542949,712 | 0,094 | 0,100 | FALSE |
| DDX1 | 2270564,870 | 2441070,841 | 0,094 | 0,238 | FALSE |
| FABP5 | 40858,106 | 43636,220 | 0,093 | 0,044 | FALSE |
| PABPC4 | 1828384,718 | 1962922,427 | 0,092 | 0,093 | FALSE |
| RPS18 | 4228537,556 | 4577053,333 | 0,092 | 0,331 | FALSE |
| SNU13 | 507410,675 | 552782,583 | 0,091 | 0,185 | FALSE |
| RPS23 | 1545402,750 | 1689660,208 | 0,089 | 0,225 | FALSE |
| PLCD3 | 21655,964 | 23489,813 | 0,089 | 0,116 | FALSE |
| MRPL11 | 579674,500 | 616970,692 | 0,088 | 0,178 | FALSE |
| ARID1A | 2256594,756 | 2398756,652 | 0,088 | 0,031 | FALSE |
| NUFIP2 | 103110,492 | 111878,272 | 0,085 | 0,128 | FALSE |
| SEC13 | 229090,346 | 245911,343 | 0,085 | 0,047 | FALSE |
| CRBN | 64454,515 | 69362,096 | 0,083 | 0,053 | FALSE |
| ARG1 | 16099,630 | 17602,938 | 0,083 | 0,075 | FALSE |
| SMCHD1 | 1787,403 | 1964,255 | 0,081 | 0,044 | FALSE |
| PTCD1 | 1076749,825 | 1153928,807 | 0,080 | 0,107 | FALSE |
| UBA1 | 12308,041 | 13633,274 | 0,079 | 0,155 | FALSE |
| IGG1 | 25290,934 | 26800,868 | 0,079 | 0,060 | FALSE |
| DCTN1 | 6268,834 | 6930,054 | 0,078 | 0,060 | FALSE |
| CAPRIN1 | 253120,420 | 277516,297 | 0,078 | 0,119 | FALSE |
| LARP1 | 1120865,393 | 1229768,438 | 0,077 | 1,039 | FALSE |
| DDX5 | 11377078,815 | 12470722,370 | 0,077 | 0,761 | FALSE |
| TAF15 | 8230271,158 | 8838883,649 | 0,075 | 0,134 | FALSE |
| TTC37 | 7088,161 | 7830,861 | 0,075 | 0,075 | FALSE |
| MRPL49 | 477667,600 | 517919,444 | 0,075 | 0,110 | FALSE |
| TRIM5 | 13414,842 | 14668,037 | 0,074 | 0,093 | FALSE |
| KCTD7 | 94784,114 | 103064,273 | 0,074 | 0,230 | FALSE |
| KPNA6 | 730846,613 | 798727,933 | 0,074 | 0,270 | FALSE |
| NHP2 | 472257,095 | 514157,833 | 0,073 | 0,112 | FALSE |
| SNRNP70 | 1824051,772 | 2037076,807 | 0,073 | 0,053 | FALSE |
| TRA2A | 202874,318 | 220694,193 | 0,072 | 0,136 | FALSE |
| MRPL15 | 502124,341 | 543272,079 | 0,071 | 0,115 | FALSE |
| HNRNPAB | 18053360,381 | 19613324,571 | 0,071 | 0,068 | FALSE |
| NOP10 | 467878,291 | 509883,364 | 0,069 | 0,112 | FALSE |

| | | | | | |
|---|---|---|---|---|---|
| SNRNP200 | 172045,835 | 189258,811 | 0,069 | 0,857 | FALSE |
| CT45A1 | 67064,475 | 73183,625 | 0,069 | 0,139 | FALSE |
| PRMT1 | 305615,587 | 333487,754 | 0,068 | 0,103 | FALSE |
| METTL17 | 13075,886 | 14343,208 | 0,068 | 0,040 | FALSE |
| ILF3 | 13310945,778 | 14486781,968 | 0,068 | 0,072 | FALSE |
| MRPS18B | 2335199,857 | 2546804,571 | 0,067 | 0,144 | FALSE |
| MTMR4 | 57956,173 | 63258,688 | 0,067 | 0,128 | FALSE |
| CHTF18 | 5472,261 | 6138,090 | 0,067 | 0,034 | FALSE |
| SMARCE1 | 2255105,609 | 2418953,674 | 0,067 | 0,013 | FALSE |
| SF1 | 3300126,386 | 3601539,228 | 0,066 | 0,145 | FALSE |
| CCAR2 | 72180,287 | 79896,603 | 0,066 | 0,088 | FALSE |
| DDX28 | 128002,963 | 138652,368 | 0,066 | 0,117 | FALSE |
| TOP2A | 13389,714 | 14907,513 | 0,065 | 0,025 | FALSE |
| HNRNPUL2 | 3393351,504 | 3685333,085 | 0,065 | 0,128 | FALSE |
| HP1BP3 | 79535,940 | 88561,978 | 0,063 | 0,079 | FALSE |
| TRNAU1AP | 1287210,167 | 1408579,972 | 0,063 | 0,073 | FALSE |
| RPL6 | 1071482,333 | 1194585,519 | 0,063 | 0,178 | FALSE |
| MRPS11 | 16152,945 | 17970,335 | 0,063 | 0,035 | FALSE |
| SRP9 | 302812,190 | 330980,250 | 0,062 | 0,146 | FALSE |
| SLC25A1 | 25528,426 | 29705,294 | 0,061 | 0,032 | FALSE |
| SRSF9 | 1186148,875 | 1300805,292 | 0,061 | 0,093 | FALSE |
| MRPS6 | 1353784,407 | 1490791,704 | 0,060 | 0,147 | FALSE |
| LOXL2 | 58537,812 | 64911,028 | 0,058 | 0,149 | FALSE |
| POLR2B | 579524,907 | 634028,204 | 0,057 | 0,045 | FALSE |
| TIMM50 | 65414,083 | 72474,077 | 0,057 | 0,073 | FALSE |
| CAPZA1 | 96161,042 | 107793,135 | 0,057 | 0,087 | FALSE |
| SF3B4 | 68530,409 | 75158,666 | 0,057 | 0,026 | FALSE |
| RPS12 | 4110539,905 | 4463035,905 | 0,056 | 0,054 | FALSE |
| ELL2 | 37594,510 | 42681,646 | 0,056 | 0,029 | FALSE |
| RPS15A | 1969570,333 | 2176728,933 | 0,056 | 0,510 | FALSE |
| FIGN | 29750,204 | 32869,564 | 0,056 | 0,157 | FALSE |
| SMNDC1 | 13784,713 | 15178,955 | 0,056 | 0,026 | FALSE |
| MYH14 | 1416,663 | 1593,979 | 0,055 | 0,029 | FALSE |
| PABPC1 | 6272176,124 | 6906284,403 | 0,054 | 0,044 | FALSE |
| POLR2D | 204852,957 | 221973,779 | 0,053 | 0,021 | FALSE |
| RRP9 | 374371,060 | 419409,548 | 0,053 | 0,083 | FALSE |
| RBM14 | 38645854,359 | 42443972,103 | 0,053 | 0,142 | FALSE |
| NKRF | 499047,489 | 548651,404 | 0,052 | 0,090 | FALSE |
| STAU1 | 682289,373 | 757075,216 | 0,051 | 0,050 | FALSE |
| RPL3 | 1290102,762 | 1444774,381 | 0,050 | 0,162 | FALSE |
| FAM98A | 1342827,389 | 1495447,444 | 0,050 | 0,081 | FALSE |
| RPS24 | 391762,262 | 434016,702 | 0,050 | 0,159 | FALSE |
| MRPL24 | 147992,317 | 161036,017 | 0,049 | 0,062 | FALSE |

| | | | | | |
|---|---|---|---|---|---|
| ZC3H18 | 49289,159 | 54829,478 | 0,047 | 0,113 | FALSE |
| RPL17 | 1218141,056 | 1355198,694 | 0,047 | 0,087 | FALSE |
| CRNKL1 | 142196,521 | 157477,333 | 0,047 | 0,196 | FALSE |
| RPL8 | 2183764,564 | 2449173,590 | 0,046 | 0,093 | FALSE |
| DPM1 | 73905,384 | 81929,296 | 0,045 | 0,079 | FALSE |
| PRPF39 | 1384260,610 | 1528664,990 | 0,045 | 0,056 | FALSE |
| PLOD1 | 2114851,007 | 2356572,085 | 0,045 | 0,280 | FALSE |
| RPS2 | 779012,076 | 878522,924 | 0,044 | 0,177 | FALSE |
| TSPAN10 | 1942454,353 | 2146274,824 | 0,044 | 0,077 | FALSE |
| BAZ2A | 6000,764 | 6693,201 | 0,043 | 0,037 | FALSE |
| MRPS26 | 1351279,378 | 1503395,600 | 0,042 | 0,105 | FALSE |
| CDC5L | 232062,255 | 258528,220 | 0,041 | 0,148 | FALSE |
| KPNA1 | 455082,377 | 505859,609 | 0,039 | 0,118 | FALSE |
| RAE1 | 209850,877 | 234704,969 | 0,039 | 0,076 | FALSE |
| ERH | 577660,000 | 634188,625 | 0,039 | 0,022 | FALSE |
| RPS14 | 2632543,944 | 2960298,444 | 0,038 | 0,183 | FALSE |
| MRPS5 | 725073,643 | 814100,857 | 0,036 | 0,128 | FALSE |
| RPL36 | 694727,900 | 791183,283 | 0,036 | 0,054 | FALSE |
| SNX9 | 1701248,571 | 1901643,857 | 0,036 | 0,056 | FALSE |
| NCBP1 | 358062,368 | 401923,291 | 0,036 | 0,041 | FALSE |
| NCL | 290803,681 | 334058,928 | 0,035 | 0,040 | FALSE |
| YBX3 | 5932048,356 | 6643812,978 | 0,035 | 0,056 | FALSE |
| KRT2 | 8152845,643 | 9323023,876 | 0,035 | 0,037 | FALSE |
| RPLP2 | 9434803,467 | 10419952,267 | 0,033 | 0,024 | FALSE |
| ERAL1 | 568191,760 | 637031,093 | 0,033 | 0,058 | FALSE |
| RBX1 | 148814,680 | 167219,688 | 0,032 | 0,060 | FALSE |
| SRSF1 | 750075,200 | 848879,022 | 0,031 | 0,125 | FALSE |
| RPS9 | 1987947,000 | 2246328,333 | 0,031 | 0,116 | FALSE |
| CNTNAP4 | 7301,647 | 8461,795 | 0,030 | 0,025 | FALSE |
| TRUB2 | 408737,205 | 455501,394 | 0,028 | 0,051 | FALSE |
| EIF3A | 53298,329 | 59997,436 | 0,027 | 0,043 | FALSE |
| FIBCD1 | 44044,915 | 49077,456 | 0,027 | 0,020 | FALSE |
| DHX9 | 7019652,468 | 7981430,342 | 0,026 | 0,021 | FALSE |
| MCAT | 47585,365 | 52797,302 | 0,025 | 0,027 | FALSE |
| SLC25A13 | 22614,666 | 25548,113 | 0,025 | 0,038 | FALSE |
| KHSRP | 861610,862 | 981102,098 | 0,024 | 0,083 | FALSE |
| DAZAP1 | 16038793,026 | 17893600,821 | 0,023 | 0,038 | FALSE |
| MSI2 | 86494,426 | 98635,518 | 0,022 | 0,026 | FALSE |
| NOA1 | 1618049,600 | 1829156,000 | 0,022 | 0,047 | FALSE |
| BCL7A | 123508,571 | 137403,893 | 0,021 | 0,005 | FALSE |
| ELOB | 201908,846 | 228646,017 | 0,021 | 0,039 | FALSE |
| ESCO2 | 6881,279 | 7391,291 | 0,019 | 0,008 | FALSE |
| ELAVL2 | 343707,092 | 390078,246 | 0,019 | 0,010 | FALSE |

| | | | | | |
|---|---|---|---|---|---|
| SEC61A1 | 65698,968 | 75573,685 | 0,019 | 0,023 | FALSE |
| PTCD3 | 84258,898 | 94847,343 | 0,018 | 0,017 | FALSE |
| DCD | 699759,387 | 793647,833 | 0,018 | 0,012 | FALSE |
| MRPL37 | 58822,797 | 67005,079 | 0,017 | 0,040 | FALSE |
| KRT19 | 11233,245 | 12855,260 | 0,017 | 0,006 | FALSE |
| MYL12A | 138021,678 | 156478,625 | 0,017 | 0,022 | FALSE |
| CCDC9B | 35630,352 | 40411,790 | 0,015 | 0,026 | FALSE |
| ILF2 | 11582742,349 | 13051774,222 | 0,015 | 0,019 | FALSE |
| PABPN1 | 430306,844 | 491979,133 | 0,014 | 0,082 | FALSE |
| WDR6 | 99854,954 | 113624,077 | 0,012 | 0,029 | FALSE |
| CAP1 | 8924,133 | 10761,779 | 0,011 | 0,003 | FALSE |
| ACTN2 | 2333,558 | 2804,008 | 0,011 | 0,005 | FALSE |
| RPS21 | 1895286,000 | 2156470,000 | 0,010 | 0,027 | FALSE |
| MRPL21 | 123232,914 | 139243,923 | 0,008 | 0,008 | FALSE |
| DSG1 | 131003,057 | 152203,973 | 0,008 | 0,010 | FALSE |
| WDCP | 193949,676 | 217200,462 | 0,008 | 0,004 | FALSE |
| USP15 | 92217,613 | 105360,065 | 0,008 | 0,013 | FALSE |
| BCAS2 | 242440,899 | 276267,375 | 0,007 | 0,025 | FALSE |
| POLR2M | 100276,353 | 115333,893 | 0,007 | 0,015 | FALSE |
| RBM19 | 1771,958 | 2070,345 | 0,006 | 0,002 | FALSE |
| RALY | 8115147,810 | 9249532,190 | 0,006 | 0,005 | FALSE |
| LUC7L2 | 39548,787 | 46417,591 | 0,005 | 0,005 | FALSE |
| SNRPA1 | 224874,792 | 258751,625 | 0,005 | 0,019 | FALSE |
| P3H1 | 100015,435 | 114793,484 | 0,005 | 0,002 | FALSE |
| MRPS15 | 928260,759 | 1063442,037 | 0,004 | 0,010 | FALSE |
| SLC25A5 | 532952,400 | 611369,017 | 0,003 | 0,007 | FALSE |
| SERBP1 | 124333,218 | 144945,149 | 0,003 | 0,004 | FALSE |
| LGALS3BP | 2144514,299 | 2435848,276 | 0,003 | 0,006 | FALSE |
| SMARCC1 | 1003987,118 | 1125097,101 | 0,003 | 0,001 | FALSE |
| PDS5A | 3049,715 | 3607,289 | 0,002 | 0,001 | FALSE |
| DKC1 | 244272,204 | 283064,477 | 0,002 | 0,002 | FALSE |
| SMARCB1 | 2417616,700 | 2712028,033 | 0,001 | 0,000 | FALSE |
| DYNC1H1 | 18550,217 | 21702,105 | 0,000 | 0,000 | FALSE |
| CYC1 | 7155,269 | 8399,257 | -0,001 | 0,000 | FALSE |
| RPLP1 | 784568,271 | 900286,354 | -0,004 | 0,006 | FALSE |
| FBL | 171965,848 | 199434,924 | -0,004 | 0,007 | FALSE |
| HNRNPA3 | 46848597,333 | 53573090,133 | -0,005 | 0,008 | FALSE |
| KARS1 | 19462,955 | 22725,221 | -0,006 | 0,005 | FALSE |
| SCYL1 | 86299,627 | 100162,685 | -0,006 | 0,009 | FALSE |
| MRPS28 | 1627811,644 | 1881198,044 | -0,007 | 0,019 | FALSE |
| G3BP1 | 144652,679 | 170905,949 | -0,007 | 0,008 | FALSE |
| LRBA | 835,597 | 967,794 | -0,007 | 0,004 | FALSE |
| RBMX | 1048117,255 | 1210725,490 | -0,008 | 0,009 | FALSE |

| | | | | | |
|---|---|---|---|---|---|
| HNRNPM | 1322348,703 | 1537583,176 | -0,008 | 0,016 | FALSE |
| NGRN | 60698,036 | 69513,800 | -0,009 | 0,010 | FALSE |
| IARS1 | 20025,161 | 23731,018 | -0,009 | 0,006 | FALSE |
| MATR3 | 1591760,609 | 1844903,304 | -0,009 | 0,011 | FALSE |
| RPS17 | 8158195,556 | 9494724,444 | -0,011 | 0,092 | FALSE |
| TARBP2 | 90886,325 | 105948,401 | -0,012 | 0,008 | FALSE |
| EIF3B | 22340,892 | 26344,761 | -0,012 | 0,010 | FALSE |
| RBBP4 | 101766,615 | 118641,613 | -0,013 | 0,016 | FALSE |
| ACTG1 | 145658,612 | 169069,366 | -0,014 | 0,008 | FALSE |
| NUP62 | 82958,890 | 93779,646 | -0,014 | 0,005 | FALSE |
| XPNPEP3 | 11348,462 | 13474,833 | -0,014 | 0,013 | FALSE |
| HNRNPK | 6759718,769 | 7889235,077 | -0,015 | 0,028 | FALSE |
| YTHDF3 | 2069051,826 | 2412558,087 | -0,015 | 0,038 | FALSE |
| DHRS2 | 448096,324 | 519972,000 | -0,016 | 0,026 | FALSE |
| SEC31A | 65771,716 | 74899,732 | -0,016 | 0,007 | FALSE |
| RBMXL1 | 54112,714 | 62868,084 | -0,017 | 0,019 | FALSE |
| HSPA9 | 18662079,008 | 21414396,527 | -0,018 | 0,024 | FALSE |
| RO60 | 69005,105 | 80619,120 | -0,018 | 0,038 | FALSE |
| SEC24C | 3957313,855 | 4603833,391 | -0,019 | 0,028 | FALSE |
| RPL32 | 677183,893 | 777718,714 | -0,019 | 0,024 | FALSE |
| DNAJC10 | 4933213,694 | 5722892,829 | -0,019 | 0,067 | FALSE |
| CRTAP | 52459,871 | 60983,598 | -0,022 | 0,012 | FALSE |
| MRPS2 | 1177579,206 | 1368638,317 | -0,022 | 0,029 | FALSE |
| KRT78 | 75316,076 | 88526,081 | -0,022 | 0,035 | FALSE |
| H1-1 | 42655,753 | 50116,806 | -0,024 | 0,026 | FALSE |
| DDX6 | 111217,740 | 130815,907 | -0,024 | 0,087 | FALSE |
| RNMT | 5194087,556 | 6057234,133 | -0,026 | 0,039 | FALSE |
| RPL22 | 2877698,722 | 3343480,000 | -0,026 | 0,055 | FALSE |
| CUL2 | 27601,135 | 32220,943 | -0,027 | 0,073 | FALSE |
| SLC25A10 | 33326,233 | 39118,326 | -0,028 | 0,039 | FALSE |
| MRPL57 | 45665,009 | 52274,734 | -0,028 | 0,019 | FALSE |
| MRPL39 | 172872,378 | 202590,392 | -0,028 | 0,085 | FALSE |
| RPL9 | 1648682,333 | 1956672,000 | -0,028 | 0,094 | FALSE |
| RPS28 | 5074585,000 | 5930628,444 | -0,029 | 0,072 | FALSE |
| AZGP1 | 47726,936 | 56670,167 | -0,029 | 0,049 | FALSE |
| PCBD1 | 212822,036 | 252320,985 | -0,029 | 0,069 | FALSE |
| RPS5 | 4017001,600 | 4708320,667 | -0,029 | 0,090 | FALSE |
| ATAD3A | 116083,888 | 137329,907 | -0,032 | 0,032 | FALSE |
| KHDRBS3 | 79145,567 | 93449,406 | -0,033 | 0,102 | FALSE |
| RCC1L | 234163,300 | 271076,840 | -0,034 | 0,051 | FALSE |
| RPS3A | 2497417,123 | 2943918,667 | -0,035 | 0,227 | FALSE |
| SSBP1 | 3751458,700 | 4458501,600 | -0,035 | 0,020 | FALSE |
| LMO7 | 1186,259 | 1401,395 | -0,035 | 0,018 | FALSE |

| | | | | | |
|---|---|---|---|---|---|
| HNRNPR | 6143100,748 | 7282890,927 | -0,036 | 0,025 | FALSE |
| UBAP2 | 1174313,556 | 1381103,919 | -0,036 | 0,045 | FALSE |
| CHTOP | 179425,637 | 213337,897 | -0,037 | 0,168 | FALSE |
| NES | 37208,275 | 43997,139 | -0,039 | 0,037 | FALSE |
| MMTAG2 | 12121,218 | 14648,539 | -0,040 | 0,017 | FALSE |
| MYCBP | 63638,297 | 73994,451 | -0,041 | 0,013 | FALSE |
| HNRNPD | 64936385,422 | 76193998,222 | -0,041 | 0,069 | FALSE |
| SAR1A | 23608,034 | 28037,488 | -0,042 | 0,038 | FALSE |
| PRPF19 | 844160,515 | 994857,152 | -0,042 | 0,230 | FALSE |
| CACYBP | 16458,038 | 19814,491 | -0,042 | 0,033 | FALSE |
| H2BC4 | 802819,333 | 958467,727 | -0,043 | 0,107 | FALSE |
| MEPCE | 27702,244 | 32839,770 | -0,044 | 0,103 | FALSE |
| BUD31 | 51770,596 | 61038,138 | -0,046 | 0,052 | FALSE |
| LACTB | 15618,025 | 18639,417 | -0,047 | 0,047 | FALSE |
| NDUFA4 | 185260,337 | 216973,559 | -0,048 | 0,050 | FALSE |
| RTN4 | 3678,183 | 4483,546 | -0,048 | 0,030 | FALSE |
| SF3B2 | 173398,864 | 203872,030 | -0,048 | 0,022 | FALSE |
| NUP93 | 37758,533 | 45044,592 | -0,050 | 0,127 | FALSE |
| KRT14 | 497260,216 | 602760,931 | -0,050 | 0,052 | FALSE |
| SMARCD2 | 1037230,356 | 1206226,429 | -0,050 | 0,010 | FALSE |
| PRKRA | 1560044,044 | 1859855,889 | -0,051 | 0,041 | FALSE |
| SF3B3 | 219788,728 | 258604,150 | -0,051 | 0,018 | FALSE |
| RPL35 | 939115,233 | 1135008,033 | -0,052 | 0,093 | FALSE |
| EEF1D | 65234,111 | 79571,149 | -0,053 | 0,075 | FALSE |
| RPA2 | 218546,851 | 259222,813 | -0,053 | 0,053 | FALSE |
| GIPC1 | 191491,931 | 228354,694 | -0,056 | 0,127 | FALSE |
| CLTC | 517656,163 | 623102,735 | -0,057 | 0,282 | FALSE |
| PPIH | 13560,350 | 16247,195 | -0,057 | 0,032 | FALSE |
| UGGT1 | 109410,733 | 131477,865 | -0,057 | 0,093 | FALSE |
| SEC23B | 3547713,400 | 4205736,133 | -0,057 | 0,072 | FALSE |
| EIF3D | 144199,889 | 174869,250 | -0,057 | 0,056 | FALSE |
| ALYREF | 96645,228 | 115968,200 | -0,058 | 0,109 | FALSE |
| POLR2C | 719633,833 | 858868,979 | -0,059 | 0,051 | FALSE |
| CNN2 | 10424,804 | 12556,350 | -0,059 | 0,032 | FALSE |
| EEF1B2 | 63301,361 | 77667,649 | -0,059 | 0,082 | FALSE |
| SON | 20824,577 | 24990,041 | -0,060 | 0,058 | FALSE |
| ATP1A1 | 31114,701 | 37136,007 | -0,060 | 0,061 | FALSE |
| MRPL13 | 114294,268 | 135081,885 | -0,062 | 0,076 | FALSE |
| RPN2 | 9243,491 | 11246,872 | -0,064 | 0,042 | FALSE |
| PSMB1 | 67723,688 | 81149,061 | -0,065 | 0,183 | FALSE |
| RPN1 | 44565,164 | 53682,491 | -0,069 | 0,181 | FALSE |
| SNRNP40 | 119722,748 | 144622,458 | -0,069 | 0,498 | FALSE |
| MVP | 341412,190 | 410566,435 | -0,070 | 0,157 | FALSE |

| | | | | | |
|---|---|---|---|---|---|
| RPL24 | 936163,938 | 1129881,729 | -0,070 | 0,093 | FALSE |
| UPF1 | 525430,021 | 629197,448 | -0,073 | 0,104 | FALSE |
| FAM120B | 30113,006 | 36449,245 | -0,073 | 0,089 | FALSE |
| ZFP36L1 | 92175,705 | 111799,240 | -0,074 | 0,047 | FALSE |
| POFUT2 | 25956,439 | 31606,784 | -0,074 | 0,168 | FALSE |
| RACK1 | 2369418,667 | 2864464,121 | -0,074 | 0,210 | FALSE |
| TUBB4B | 456696,818 | 557145,879 | -0,075 | 0,187 | FALSE |
| SRP14 | 494760,167 | 604477,524 | -0,075 | 0,172 | FALSE |
| TRIM28 | 19390,206 | 23479,786 | -0,075 | 0,138 | FALSE |
| MLF2 | 54272,581 | 66569,811 | -0,076 | 0,036 | FALSE |
| AGL | 106870,111 | 128827,020 | -0,077 | 0,177 | FALSE |
| RUVBL2 | 35366,373 | 43243,545 | -0,078 | 0,117 | FALSE |
| TFIP11 | 1748,247 | 2164,789 | -0,079 | 0,032 | FALSE |
| PURB | 705440,550 | 860663,678 | -0,079 | 0,042 | FALSE |
| LENG8 | 338327,488 | 405496,589 | -0,081 | 0,047 | FALSE |
| SAFB | 44414,377 | 54843,847 | -0,081 | 0,149 | FALSE |
| SMARCD1 | 242264,952 | 288164,543 | -0,082 | 0,017 | FALSE |
| IGL1 | 143961,633 | 175164,242 | -0,082 | 0,074 | FALSE |
| ATP5F1C | 64755,903 | 79189,990 | -0,082 | 0,090 | FALSE |
| YTHDF2 | 587040,884 | 713054,145 | -0,085 | 0,121 | FALSE |
| RPUSD3 | 76708,154 | 91967,422 | -0,086 | 0,117 | FALSE |
| PSMA3 | 11053,490 | 13670,449 | -0,087 | 0,053 | FALSE |
| EMD | 33741,297 | 41097,808 | -0,088 | 0,117 | FALSE |
| GCA | 207142,600 | 255549,504 | -0,088 | 0,169 | FALSE |
| MRPS34 | 1517853,667 | 1842146,333 | -0,088 | 0,141 | FALSE |
| ADNP | 22672,794 | 28016,326 | -0,093 | 0,122 | FALSE |
| ATP5MG | 62838,588 | 75917,297 | -0,093 | 0,082 | FALSE |
| MTERF3 | 236341,359 | 286495,327 | -0,093 | 0,182 | FALSE |
| AQR | 65108,498 | 79334,027 | -0,093 | 0,414 | FALSE |
| RPS20 | 2974389,389 | 3632475,222 | -0,095 | 0,172 | FALSE |
| RPL34 | 1351863,722 | 1673665,111 | -0,095 | 0,262 | FALSE |
| RBM22 | 129232,840 | 159142,327 | -0,095 | 0,323 | FALSE |
| HNRNPA0 | 16656903,238 | 20435043,048 | -0,095 | 0,160 | FALSE |
| FUBP1 | 469719,926 | 581621,389 | -0,096 | 0,247 | FALSE |
| XAB2 | 136283,582 | 166822,394 | -0,096 | 0,344 | FALSE |
| RPAP2 | 22129,069 | 27050,783 | -0,097 | 0,132 | FALSE |
| TARDBP | 1669474,549 | 2050510,510 | -0,097 | 0,124 | FALSE |
| IGF2BP2 | 380423,657 | 467690,304 | -0,098 | 0,137 | FALSE |
| LSM14B | 34922,780 | 43698,745 | -0,100 | 0,169 | FALSE |
| ZFR | 704454,824 | 869852,680 | -0,100 | 0,097 | FALSE |
| RPL37A | 1169558,083 | 1446052,111 | -0,101 | 0,345 | FALSE |
| PDCD6IP | 26173,438 | 32360,899 | -0,102 | 0,215 | FALSE |
| PDCD11 | 5971,439 | 7428,131 | -0,103 | 0,046 | FALSE |

| | | | | | |
|---|---|---|---|---|---|
| STK3 | 5991,998 | 7419,422 | -0,105 | 0,072 | FALSE |
| STRAP | 25220,031 | 31434,484 | -0,106 | 0,230 | FALSE |
| RUVBL1 | 22702,606 | 28054,179 | -0,108 | 0,073 | FALSE |
| SEC23A | 6417156,650 | 7934798,906 | -0,109 | 0,192 | FALSE |
| POLR2A | 658002,608 | 807001,369 | -0,110 | 0,117 | FALSE |
| SLC25A3 | 264799,246 | 327972,886 | -0,110 | 0,177 | FALSE |
| ELOC | 312214,297 | 386511,802 | -0,111 | 0,219 | FALSE |
| NCAPH | 12498,352 | 15288,744 | -0,112 | 0,060 | FALSE |
| POLR2E | 641536,179 | 794894,551 | -0,113 | 0,107 | FALSE |
| RPL28 | 1537814,630 | 1940676,259 | -0,113 | 0,324 | FALSE |
| DDX17 | 1985946,537 | 2486061,984 | -0,115 | 0,511 | FALSE |
| RPL23A | 1983009,519 | 2501200,889 | -0,116 | 0,269 | FALSE |
| SYNCRIP | 1365308,462 | 1718486,154 | -0,117 | 0,092 | FALSE |
| HNRNPU | 27515567,891 | 33889421,061 | -0,117 | 0,133 | FALSE |
| RPS11 | 2775131,333 | 3467003,267 | -0,118 | 1,515 | FALSE |
| SNRPE | 4831216,417 | 6014773,500 | -0,119 | 0,122 | FALSE |
| NUP88 | 104757,791 | 129328,750 | -0,121 | 0,049 | FALSE |
| ACTB | 232079,768 | 291244,471 | -0,122 | 0,119 | FALSE |
| SLC3A2 | 3467,754 | 4404,714 | -0,122 | 0,056 | FALSE |
| EPB41L2 | 3685,785 | 4778,992 | -0,124 | 0,054 | FALSE |
| FLNA | 39830,001 | 50815,916 | -0,124 | 0,269 | FALSE |
| PSMA4 | 11794,166 | 15226,582 | -0,126 | 0,078 | FALSE |
| EIF3I | 20497,195 | 26224,386 | -0,127 | 0,175 | FALSE |
| SEC22B | 36774,783 | 45740,488 | -0,128 | 0,170 | FALSE |
| RBFOX2 | 451919,848 | 569770,167 | -0,129 | 0,400 | FALSE |
| ARHGEF39 | 7740,208 | 9756,993 | -0,129 | 0,065 | FALSE |
| CHD4 | 32360,250 | 41416,920 | -0,131 | 0,088 | FALSE |
| TBC1D8B | 4869,624 | 6192,901 | -0,133 | 0,149 | FALSE |
| PCMT1 | 714103,028 | 901859,514 | -0,133 | 0,191 | FALSE |
| PALLD | 111245,879 | 140997,624 | -0,134 | 0,595 | FALSE |
| DDX3X | 11710493,659 | 14820493,008 | -0,134 | 1,198 | FALSE |
| KPNB1 | 73548,691 | 94631,477 | -0,135 | 0,179 | FALSE |
| HNRNPA1 | 48349792,561 | 61187962,386 | -0,136 | 0,218 | FALSE |
| RPA1 | 232341,303 | 293680,741 | -0,136 | 0,156 | FALSE |
| SFXN3 | 35095,703 | 44987,047 | -0,137 | 0,283 | FALSE |
| ERP29 | 13841,143 | 18037,297 | -0,137 | 0,103 | FALSE |
| GRPEL1 | 30613,881 | 38388,041 | -0,139 | 0,124 | FALSE |
| KHDRBS1 | 2434273,651 | 3108066,286 | -0,140 | 0,536 | FALSE |
| HNRNPC | 23582409,697 | 29876979,394 | -0,140 | 0,109 | FALSE |
| LMNB1 | 100679,594 | 129047,794 | -0,141 | 0,142 | FALSE |
| EPRS1 | 13422,836 | 17472,938 | -0,141 | 0,077 | FALSE |
| NUP214 | 222185,472 | 278193,642 | -0,142 | 0,071 | FALSE |
| ELAVL1 | 5740452,593 | 7292193,259 | -0,142 | 0,095 | FALSE |

| | | | | | |
|---|---|---|---|---|---|
| MRPS25 | 1035362,515 | 1317030,697 | -0,143 | 0,358 | FALSE |
| GTF2I | 156160,538 | 197370,492 | -0,144 | 0,278 | FALSE |
| RRP1B | 83558,008 | 106974,917 | -0,144 | 0,170 | FALSE |
| MRPL30 | 28085,229 | 35782,932 | -0,145 | 0,152 | FALSE |
| PCID2 | 231212,192 | 292435,683 | -0,145 | 0,071 | FALSE |
| TXNL4A | 26752,797 | 34570,152 | -0,147 | 0,130 | FALSE |
| RPL31 | 3219916,286 | 4134570,286 | -0,148 | 0,372 | FALSE |
| LDB1 | 65335,692 | 81730,761 | -0,148 | 0,178 | FALSE |
| YWHAB | 36544,402 | 47191,863 | -0,150 | 0,165 | FALSE |
| SNW1 | 122099,529 | 154654,948 | -0,150 | 0,380 | FALSE |
| GLYR1 | 3845,556 | 4940,113 | -0,151 | 0,079 | FALSE |
| FUBP3 | 113261,890 | 145768,711 | -0,151 | 0,330 | FALSE |
| TUBA3C | 112927,420 | 144788,696 | -0,152 | 0,284 | FALSE |
| HNRNPF | 22884274,087 | 29129676,986 | -0,153 | 0,565 | FALSE |
| FKBP10 | 44718,631 | 57424,285 | -0,156 | 0,658 | FALSE |
| IGL@ | 229484,958 | 290089,117 | -0,156 | 0,173 | FALSE |
| EIF4A1 | 18106,274 | 23496,320 | -0,157 | 0,156 | FALSE |
| STAU2 | 423762,024 | 539909,130 | -0,160 | 0,160 | FALSE |
| COPA | 21489,629 | 28273,643 | -0,161 | 0,068 | FALSE |
| POLR2G | 368172,371 | 467921,158 | -0,161 | 0,102 | FALSE |
| TNKS1BP1 | 18995,266 | 24767,118 | -0,162 | 0,250 | FALSE |
| ATP2A2 | 11065,752 | 14032,789 | -0,162 | 0,162 | FALSE |
| DNAJA3 | 26678,594 | 33869,114 | -0,164 | 0,218 | FALSE |
| PCBP1 | 237827,976 | 302949,405 | -0,166 | 0,181 | FALSE |
| TRIM27 | 4457,489 | 5784,525 | -0,167 | 0,082 | FALSE |
| SKP1 | 133631,149 | 171583,719 | -0,167 | 0,384 | FALSE |
| SF3A1 | 34825,768 | 44570,849 | -0,167 | 0,077 | FALSE |
| CUL3 | 1815,847 | 2353,904 | -0,168 | 0,062 | FALSE |
| TEP1 | 1028,242 | 1342,968 | -0,168 | 0,107 | FALSE |
| ACTL6A | 2003914,712 | 2538427,515 | -0,169 | 0,051 | FALSE |
| MIF | 157869,071 | 207008,841 | -0,169 | 0,288 | FALSE |
| PNN | 2941,384 | 3766,881 | -0,172 | 0,076 | FALSE |
| PEF1 | 5631151,111 | 7317997,778 | -0,172 | 0,608 | FALSE |
| FAM120A | 782966,633 | 1010785,533 | -0,173 | 0,144 | FALSE |
| ALDH9A1 | 7704,945 | 10256,044 | -0,175 | 0,151 | FALSE |
| HADHA | 28819,858 | 37888,668 | -0,176 | 0,397 | FALSE |
| CPSF4 | 30803,898 | 39460,958 | -0,176 | 0,175 | FALSE |
| CARM1 | 3134620,346 | 4038067,160 | -0,177 | 0,394 | FALSE |
| NUMA1 | 1263,850 | 1624,762 | -0,178 | 0,132 | FALSE |
| AHCY | 232888,056 | 307212,167 | -0,178 | 0,482 | FALSE |
| SMARCD3 | 106595,328 | 135953,735 | -0,180 | 0,042 | FALSE |
| RPS26 | 2193262,889 | 2875839,222 | -0,182 | 0,367 | FALSE |
| KRT10 | 20160340,190 | 26736754,286 | -0,184 | 0,399 | FALSE |

| | | | | | |
|---|---|---|---|---|---|
| POLR2K | 185870,153 | 242510,674 | -0,185 | 0,198 | FALSE |
| NUDC | 9851,288 | 13202,142 | -0,185 | 0,121 | FALSE |
| CBX2 | 5846,228 | 7700,091 | -0,189 | 0,110 | FALSE |
| LANCL1 | 44202,594 | 58243,742 | -0,190 | 0,470 | FALSE |
| EIF3G | 5403,145 | 7246,577 | -0,190 | 0,096 | FALSE |
| HSPH1 | 7424,058 | 9836,251 | -0,192 | 0,226 | FALSE |
| TRAP1 | 59167,884 | 80254,747 | -0,192 | 0,180 | FALSE |
| RFC1 | 2860,246 | 3749,958 | -0,192 | 0,089 | FALSE |
| PRKDC | 22199,754 | 29497,830 | -0,197 | 0,341 | FALSE |
| SS18 | 2636284,222 | 3429258,556 | -0,199 | 0,145 | FALSE |
| CDSN | 34666,710 | 46817,861 | -0,200 | 0,157 | FALSE |
| CTTN | 143044,469 | 191164,611 | -0,201 | 0,573 | FALSE |
| MYH9 | 70205,503 | 94606,419 | -0,202 | 0,144 | FALSE |
| OGFR | 15378,415 | 20642,296 | -0,204 | 0,210 | FALSE |
| RBM8A | 139213,401 | 184021,943 | -0,205 | 0,312 | FALSE |
| SRSF2 | 219335,860 | 295253,318 | -0,207 | 0,642 | FALSE |
| HNRNPL | 5021066,667 | 6744179,030 | -0,209 | 0,318 | FALSE |
| TUBB4A | 22780,680 | 30280,763 | -0,210 | 0,506 | FALSE |
| PUM1 | 75420,391 | 99549,800 | -0,210 | 0,774 | FALSE |
| EIF2AK2 | 6005,927 | 8117,307 | -0,211 | 0,114 | FALSE |
| TRA2B | 501200,896 | 658855,514 | -0,211 | 0,335 | FALSE |
| SLTM | 2391246,536 | 3191692,549 | -0,213 | 0,312 | FALSE |
| LARP4 | 194123,766 | 263459,984 | -0,213 | 0,219 | FALSE |
| TENT4B | 13829,844 | 17938,900 | -0,215 | 0,107 | FALSE |
| KRT80 | 35910,437 | 48639,722 | -0,216 | 0,293 | FALSE |
| SRI | 1025779,077 | 1378763,974 | -0,216 | 0,593 | FALSE |
| AHNAK | 8097,347 | 10841,795 | -0,218 | 0,225 | FALSE |
| SRSF7 | 782592,964 | 1041645,595 | -0,218 | 0,485 | FALSE |
| SMARCA2 | 585488,379 | 770274,393 | -0,218 | 0,076 | FALSE |
| MRPL12 | 138043,654 | 183664,167 | -0,219 | 0,142 | FALSE |
| SUPT5H | 12132,183 | 16491,610 | -0,219 | 0,599 | FALSE |
| PYCR1 | 16666,127 | 22567,157 | -0,222 | 0,260 | FALSE |
| CAVIN1 | 72739,433 | 99730,314 | -0,223 | 0,229 | FALSE |
| FASTKD2 | 108553,886 | 142575,031 | -0,223 | 0,342 | FALSE |
| SUPT16H | 29572,173 | 40677,641 | -0,223 | 0,128 | FALSE |
| PSMA1 | 36996,768 | 53068,650 | -0,228 | 0,140 | FALSE |
| RPL29 | 1041542,633 | 1414883,667 | -0,229 | 0,398 | FALSE |
| APOBEC3B | 1381542,639 | 1858684,444 | -0,231 | 0,301 | FALSE |
| TUBB | 2065923,758 | 2807757,515 | -0,232 | 0,422 | FALSE |
| KRT17 | 164807,907 | 220645,275 | -0,234 | 0,252 | FALSE |
| VCP | 42578,203 | 58119,979 | -0,234 | 0,669 | FALSE |
| MOV10 | 1852319,196 | 2511358,899 | -0,235 | 0,616 | FALSE |
| SS18L1 | 979084,607 | 1311621,119 | -0,236 | 0,152 | FALSE |

| | | | | | |
|---|---|---|---|---|---|
| PLRG1 | 40575,811 | 54508,422 | -0,237 | 0,314 | FALSE |
| DHX8 | 8750,449 | 11868,471 | -0,239 | 0,703 | FALSE |
| SUGP2 | 74739,144 | 100920,446 | -0,240 | 0,873 | FALSE |
| JUP | 100526,261 | 138832,626 | -0,241 | 0,279 | FALSE |
| SF3B6 | 275970,899 | 372032,156 | -0,243 | 0,112 | FALSE |
| DDB1 | 93926,991 | 127058,273 | -0,244 | 0,417 | FALSE |
| KRT1 | 26586368,000 | 36984811,886 | -0,245 | 0,337 | FALSE |
| ADAR | 1706174,788 | 2317191,654 | -0,245 | 0,234 | FALSE |
| KRT9 | 17600868,978 | 24661886,222 | -0,245 | 0,227 | FALSE |
| PUM2 | 98403,145 | 136169,514 | -0,248 | 0,215 | FALSE |
| ATP5F1B | 183941,586 | 251903,900 | -0,248 | 0,567 | FALSE |
| EZHIP | 19598,964 | 27044,785 | -0,248 | 0,405 | FALSE |
| DSP | 55923,677 | 77625,034 | -0,249 | 0,208 | FALSE |
| PEBP1 | 11248,394 | 15725,011 | -0,251 | 0,129 | FALSE |
| NUDT21 | 17207,701 | 23849,155 | -0,251 | 0,335 | FALSE |
| MAGEA4 | 72925,021 | 101247,474 | -0,251 | 0,382 | FALSE |
| CCT6A | 15140,459 | 21084,285 | -0,253 | 0,395 | FALSE |
| H3-2 | 70976,586 | 96755,534 | -0,259 | 0,496 | FALSE |
| RPL36A | 385398,069 | 536123,653 | -0,260 | 0,574 | FALSE |
| CSTA | 157311,284 | 214589,453 | -0,262 | 0,742 | FALSE |
| RCN1 | 49415,582 | 69350,697 | -0,263 | 0,367 | FALSE |
| CLTA | 355178,000 | 493228,133 | -0,263 | 0,812 | FALSE |
| RBM3 | 1011291,889 | 1434018,315 | -0,264 | 0,261 | FALSE |
| GRSF1 | 18809,886 | 26321,804 | -0,266 | 0,157 | FALSE |
| MRPL22 | 339458,233 | 465592,956 | -0,270 | 0,903 | FALSE |
| RPL18 | 1297170,222 | 1803223,000 | -0,273 | 1,619 | FALSE |
| SF3A3 | 14764,061 | 20530,321 | -0,273 | 0,114 | FALSE |
| H2AC4 | 48651,880 | 63931,480 | -0,274 | 0,126 | FALSE |
| GRN | 1688993,101 | 2300737,101 | -0,275 | 0,255 | FALSE |
| TUBB6 | 615473,522 | 854374,754 | -0,275 | 0,782 | FALSE |
| MAP7D1 | 10557,339 | 14869,503 | -0,275 | 0,293 | FALSE |
| PGRMC1 | 43785,648 | 62276,599 | -0,277 | 0,344 | FALSE |
| STRBP | 337051,043 | 468687,906 | -0,278 | 0,327 | FALSE |
| SLC7A5 | 16472,513 | 23023,409 | -0,278 | 0,407 | FALSE |
| TUBB3 | 60597,303 | 84059,174 | -0,278 | 0,403 | FALSE |
| KIAA0586 | 33399,469 | 47084,523 | -0,279 | 0,686 | FALSE |
| LAS1L | 15045,628 | 21040,553 | -0,279 | 0,283 | FALSE |
| NIPSNAP1 | 11221,286 | 15546,925 | -0,284 | 0,161 | FALSE |
| SRSF10 | 112355,000 | 159436,782 | -0,286 | 0,748 | FALSE |
| PDXK | 20585,681 | 28559,548 | -0,287 | 0,456 | FALSE |
| POLR2I | 206006,275 | 283753,880 | -0,289 | 0,174 | FALSE |
| FLNC | 112717,125 | 161113,599 | -0,290 | 0,567 | FALSE |
| CAD | 54382,683 | 77441,164 | -0,292 | 1,098 | FALSE |

| | | | | | |
|---|---|---|---|---|---|
| FASN | 45178,015 | 64921,327 | -0,292 | 0,361 | FALSE |
| CUL4B | 1790,434 | 2543,703 | -0,296 | 0,145 | FALSE |
| FHL2 | 51167,761 | 71899,807 | -0,297 | 0,599 | FALSE |
| ANXA7 | 1862341,707 | 2641180,960 | -0,297 | 0,586 | FALSE |
| LRPPRC | 5476,753 | 7880,183 | -0,299 | 0,379 | FALSE |
| MYL6 | 85508,246 | 119270,239 | -0,300 | 0,450 | FALSE |
| AHDC1 | 6797,726 | 9560,369 | -0,301 | 0,705 | FALSE |
| S100A8 | 236324,860 | 342540,151 | -0,301 | 0,388 | FALSE |
| NGDN | 14765,846 | 21722,303 | -0,301 | 0,125 | FALSE |
| PPP2R1A | 13750,772 | 19244,600 | -0,303 | 0,603 | FALSE |
| EEF1G | 234864,731 | 339500,872 | -0,303 | 0,366 | FALSE |
| ZNF316 | 5306,777 | 7609,776 | -0,303 | 0,142 | FALSE |
| ADARB1 | 5297,822 | 7429,055 | -0,306 | 0,179 | FALSE |
| ANXA11 | 216212,697 | 312063,144 | -0,306 | 0,675 | FALSE |
| SMARCA4 | 1212155,662 | 1699467,698 | -0,309 | 0,117 | FALSE |
| PLEC | 225352,846 | 322753,793 | -0,313 | 0,212 | FALSE |
| SNRPF | 934743,950 | 1319156,000 | -0,315 | 0,247 | FALSE |
| SNRPGP15 | 2602394,200 | 3780061,067 | -0,318 | 0,466 | FALSE |
| EIF2S1 | 18697,365 | 27023,266 | -0,321 | 0,376 | FALSE |
| MRPL54 | 11371963,048 | 16216828,571 | -0,322 | 0,563 | FALSE |
| EPB41L3 | 4246,717 | 6180,769 | -0,325 | 0,186 | FALSE |
| CTPS1 | 278404,687 | 402990,323 | -0,326 | 0,701 | FALSE |
| CUL4A | 2351,295 | 3399,238 | -0,328 | 0,189 | FALSE |
| SDHA | 86802,916 | 123778,197 | -0,328 | 0,231 | FALSE |
| PDIA6 | 149981,657 | 220154,873 | -0,329 | 1,148 | FALSE |
| VARS1 | 7013,115 | 10487,897 | -0,330 | 0,195 | FALSE |
| OTUD4 | 7511,676 | 10817,670 | -0,331 | 0,112 | FALSE |
| HSPB1 | 3400328,524 | 5021848,476 | -0,332 | 0,361 | FALSE |
| CNN3 | 10311,316 | 15156,321 | -0,334 | 0,223 | FALSE |
| VDAC1 | 48853,972 | 71642,662 | -0,334 | 0,829 | FALSE |
| CREBBP | 1386,835 | 2035,112 | -0,336 | 0,149 | FALSE |
| RPL39P5 | 229254,623 | 325322,387 | -0,336 | 0,228 | FALSE |
| SRBD1 | 7591,363 | 10970,683 | -0,336 | 0,688 | FALSE |
| MRPS30 | 17579,710 | 26718,649 | -0,339 | 0,179 | FALSE |
| MAN2C1 | 651615,422 | 968553,524 | -0,341 | 0,290 | FALSE |
| ANXA2 | 537813,920 | 800283,627 | -0,341 | 0,705 | FALSE |
| USP5 | 2991,798 | 4402,653 | -0,342 | 0,218 | FALSE |
| PPT1 | 9806,752 | 14326,545 | -0,343 | 0,206 | FALSE |
| MRPL32 | 13254,842 | 19406,595 | -0,343 | 0,181 | FALSE |
| HSP90AA1 | 121914,046 | 180860,561 | -0,344 | 0,912 | FALSE |
| S100A9 | 557933,593 | 836471,519 | -0,344 | 0,597 | FALSE |
| MCM7 | 13509,443 | 19669,672 | -0,346 | 0,801 | FALSE |
| POLR2H | 525116,608 | 763830,033 | -0,346 | 0,354 | FALSE |

| | | | | | |
|---|---|---|---|---|---|
| ZC3H11A | 18618,354 | 27679,056 | -0,347 | 0,736 | FALSE |
| GRB2 | 32561,560 | 48397,137 | -0,347 | 0,792 | FALSE |
| PTBP1 | 187509,905 | 275702,111 | -0,349 | 0,836 | FALSE |
| SFPQ | 184545,130 | 272104,042 | -0,351 | 1,254 | FALSE |
| DSC1 | 52362,218 | 78383,415 | -0,351 | 0,571 | FALSE |
| KRT5 | 1124443,238 | 1682888,305 | -0,353 | 0,893 | FALSE |
| COPB2 | 9531,852 | 14345,359 | -0,354 | 0,175 | FALSE |
| DTX2 | 5460,570 | 8095,438 | -0,354 | 0,230 | FALSE |
| LMNA | 240462,015 | 356334,705 | -0,355 | 1,003 | FALSE |
| GNAS | 2865,271 | 4161,789 | -0,355 | 0,183 | FALSE |
| MSN | 8378,054 | 12571,869 | -0,356 | 0,350 | FALSE |
| C3 | 1477,141 | 2192,276 | -0,357 | 0,280 | FALSE |
| HDAC6 | 5301,858 | 7741,493 | -0,365 | 0,335 | FALSE |
| PPIL1 | 183704,446 | 268944,717 | -0,366 | 0,788 | FALSE |
| H3C15 | 1653904,056 | 2438480,889 | -0,367 | 0,594 | FALSE |
| ESYT1 | 7830,859 | 12108,585 | -0,372 | 0,162 | FALSE |
| PKP2 | 20047,936 | 29329,219 | -0,373 | 0,647 | FALSE |
| SERPINB14 | 10063,883 | 15163,782 | -0,382 | 0,284 | FALSE |
| HSP90B1 | 19650,625 | 29945,966 | -0,382 | 0,924 | FALSE |
| PDIA3 | 18273,229 | 28162,611 | -0,385 | 0,565 | FALSE |
| RPS15 | 2315395,625 | 3461800,083 | -0,391 | 1,585 | FALSE |
| SUPV3L1 | 2845,694 | 4208,473 | -0,391 | 0,217 | FALSE |
| TGM3 | 14456,158 | 22283,966 | -0,395 | 0,697 | FALSE |
| SLC25A6 | 56936,010 | 87171,830 | -0,395 | 0,664 | FALSE |
| TYMS | 6906,288 | 10747,791 | -0,400 | 0,253 | FALSE |
| EEF2 | 389670,692 | 599591,421 | -0,405 | 0,677 | FALSE |
| IGHG3 | 7305,651 | 11199,850 | -0,407 | 0,260 | FALSE |
| P4HB | 20975,780 | 32374,976 | -0,409 | 0,538 | FALSE |
| YARS2 | 96244,802 | 145577,632 | -0,409 | 0,698 | FALSE |
| FASTK | 4111,086 | 6287,084 | -0,410 | 0,254 | FALSE |
| DARS1 | 8105,956 | 12645,289 | -0,418 | 0,518 | FALSE |
| CASP14 | 64855,711 | 99872,604 | -0,423 | 1,609 | FALSE |
| SEC24D | 72830,386 | 113616,884 | -0,424 | 1,418 | FALSE |
| SEC24A | 266364,158 | 409674,658 | -0,425 | 0,839 | FALSE |
| GCN1 | 3158,979 | 4903,005 | -0,427 | 1,089 | FALSE |
| APOBEC3F | 238024,022 | 368025,000 | -0,429 | 1,072 | FALSE |
| IQGAP3 | 2779,630 | 4412,254 | -0,429 | 0,305 | FALSE |
| PSMB6 | 15932,272 | 24796,788 | -0,433 | 0,359 | FALSE |
| RPL18A | 509678,731 | 791163,282 | -0,434 | 1,307 | FALSE |
| SAFB2 | 10380,674 | 16173,705 | -0,435 | 1,185 | FALSE |
| SYPL1 | 23126,252 | 35667,567 | -0,441 | 0,514 | FALSE |
| TXN | 59613,398 | 94431,412 | -0,445 | 0,762 | FALSE |
| PHF2 | 62387,248 | 97069,816 | -0,449 | 0,413 | FALSE |

| | | | | | |
|---|---|---|---|---|---|
| TFRC | 9425,410 | 15194,075 | -0,450 | 0,260 | FALSE |
| VIM | 3971586,514 | 6182693,486 | -0,451 | 0,137 | FALSE |
| PPIB | 65112,660 | 102508,664 | -0,457 | 0,501 | FALSE |
| TCP1 | 25680,739 | 41258,898 | -0,460 | 1,188 | FALSE |
| FLG2 | 28714,189 | 46470,450 | -0,460 | 0,618 | FALSE |
| MEMO1 | 40610,754 | 65091,114 | -0,467 | 0,856 | FALSE |
| ATP5F1A | 167833,856 | 269657,236 | -0,472 | 1,515 | FALSE |
| SAP18 | 39898,319 | 63489,872 | -0,472 | 0,276 | FALSE |
| KPNA4 | 5848,889 | 9344,997 | -0,472 | 0,276 | FALSE |
| U2AF2 | 159044,452 | 255451,904 | -0,476 | 2,140 | FALSE |
| YTHDC1 | 23029,290 | 37450,976 | -0,485 | 0,549 | FALSE |
| POLR2J | 585594,007 | 893489,236 | -0,485 | 0,269 | FALSE |
| MTHFD1 | 9262,957 | 15106,553 | -0,486 | 1,102 | FALSE |
| SPG21 | 16243,924 | 26536,211 | -0,493 | 0,561 | FALSE |
| KRT18 | 655743,300 | 1064278,356 | -0,501 | 0,191 | FALSE |
| IMPDH2 | 50585,712 | 84371,452 | -0,510 | 0,810 | FALSE |
| RPS10 | 1623347,222 | 2702068,056 | -0,525 | 2,849 | FALSE |
| HM13 | 27906,879 | 46012,292 | -0,525 | 0,967 | FALSE |
| L1RE1 | 74275,542 | 122645,060 | -0,525 | 2,060 | FALSE |
| KRT8 | 1194086,521 | 1975133,500 | -0,527 | 0,225 | FALSE |
| POLR2F | 103621,065 | 173716,810 | -0,528 | 0,399 | FALSE |
| FAM186A | 16892,470 | 27677,835 | -0,540 | 0,120 | FALSE |
| HSP90AB1 | 256077,399 | 438993,754 | -0,546 | 1,050 | FALSE |
| GGCT | 42115,620 | 72464,504 | -0,549 | 0,896 | FALSE |
| TUBA1B | 356810,482 | 595761,826 | -0,549 | 0,769 | FALSE |
| LARS1 | 7263,092 | 12964,621 | -0,553 | 0,208 | FALSE |
| GNB2 | 46068,596 | 77143,401 | -0,558 | 0,391 | FALSE |
| PCBP2 | 198570,677 | 334060,650 | -0,559 | 0,795 | FALSE |
| CCT8 | 32124,009 | 55190,806 | -0,562 | 1,188 | FALSE |
| S100A10 | 89105,779 | 151415,154 | -0,569 | 0,735 | FALSE |
| SND1 | 5160,432 | 9130,504 | -0,573 | 0,307 | FALSE |
| ATP5MK | 57034,192 | 97592,318 | -0,584 | 0,497 | FALSE |
| NAP1L1 | 77809,430 | 137173,351 | -0,588 | 1,313 | FALSE |
| H2AZ2 | 412629,882 | 714907,694 | -0,598 | 0,362 | FALSE |
| HSPD1 | 65535,482 | 116904,140 | -0,603 | 1,203 | FALSE |
| HSPE1 | 57411,155 | 101304,435 | -0,604 | 1,495 | FALSE |
| U2AF1 | 180491,292 | 318960,597 | -0,608 | 3,391 | FALSE |
| DNAJA1 | 67759,094 | 122012,087 | -0,632 | 1,257 | FALSE |
| U2SURP | 297060,288 | 532282,847 | -0,634 | 0,765 | FALSE |
| CIRBP | 20598,592 | 38170,090 | -0,636 | 0,475 | FALSE |
| SEC61B | 76915,138 | 140396,253 | -0,653 | 1,041 | FALSE |
| SRSF5 | 25447,866 | 45535,399 | -0,655 | 0,701 | FALSE |
| LMNB2 | 4662,616 | 8628,667 | -0,657 | 0,648 | FALSE |

| | | | | | |
|---|---|---|---|---|---|
| LDHB | 86812,736 | 162294,425 | -0,659 | 0,917 | FALSE |
| PFN1 | 213775,389 | 398527,935 | -0,661 | 1,063 | FALSE |
| CAT | 7096,040 | 12639,320 | -0,670 | 0,421 | FALSE |
| FKBP4 | 19562,756 | 36983,857 | -0,672 | 0,933 | FALSE |
| SDHB | 13012,931 | 23309,844 | -0,677 | 0,626 | FALSE |
| ATP5MF | 177957,863 | 325016,021 | -0,684 | 0,937 | FALSE |
| NME2 | 68321,778 | 128738,860 | -0,685 | 0,840 | FALSE |
| CTSD | 40047,810 | 75128,470 | -0,687 | 1,945 | FALSE |
| ARF1 | 18457,320 | 35016,555 | -0,689 | 0,700 | FALSE |
| GIGYF2 | 3303,102 | 6162,262 | -0,690 | 0,511 | FALSE |
| AIMP2 | 25548,162 | 48052,961 | -0,706 | 0,964 | FALSE |
| RCHY1 | 110813,789 | 205666,232 | -0,707 | 0,982 | FALSE |
| POLR2L | 75392,243 | 139376,967 | -0,707 | 0,856 | FALSE |
| FAM98B | 5397,683 | 10385,140 | -0,715 | 0,420 | FALSE |
| CCT7 | 4844,492 | 9593,170 | -0,731 | 0,634 | FALSE |
| S100A7 | 145558,741 | 276426,786 | -0,733 | 0,918 | FALSE |
| ENO1 | 212763,784 | 418782,920 | -0,735 | 1,108 | FALSE |
| TAGLN2 | 97047,624 | 189803,603 | -0,745 | 1,053 | FALSE |
| RANGAP1 | 5350,342 | 10491,055 | -0,748 | 0,575 | FALSE |
| RPS29 | 963754,542 | 1845194,292 | -0,751 | 1,199 | FALSE |
| STARD9 | 2267,493 | 4353,866 | -0,758 | 1,577 | FALSE |
| NONO | 168240,093 | 331234,519 | -0,758 | 2,325 | FALSE |
| HBB | 11485,676 | 22440,196 | -0,765 | 0,617 | FALSE |
| PSMB5 | 43272,396 | 85813,150 | -0,768 | 1,018 | FALSE |
| PHGDH | 41284,644 | 82112,543 | -0,775 | 1,962 | FALSE |
| TUBB8 | 216754,128 | 419404,992 | -0,780 | 0,724 | FALSE |
| CALU | 15008,080 | 30637,545 | -0,789 | 0,999 | FALSE |
| DHX36 | 7747,029 | 15859,401 | -0,800 | 0,717 | FALSE |
| SOD1 | 29165,116 | 59924,637 | -0,802 | 1,011 | FALSE |
| BTRC | 5956,140 | 11977,535 | -0,807 | 1,089 | FALSE |
| RAN | 49204,132 | 101296,344 | -0,808 | 1,514 | FALSE |
| PRPF4 | 18365,320 | 37957,586 | -0,813 | 0,558 | FALSE |
| TPI1 | 171591,505 | 356730,598 | -0,824 | 1,531 | FALSE |
| CFL1 | 149090,302 | 312419,819 | -0,825 | 1,123 | FALSE |
| MCM4 | 3385,050 | 6843,811 | -0,840 | 0,441 | FALSE |
| RPS27A | 131135,129 | 268251,083 | -0,842 | 1,023 | FALSE |
| EEF1A1 | 327246,800 | 696146,587 | -0,862 | 1,894 | FALSE |
| BCL7C | 861075,065 | 1775045,889 | -0,863 | 0,400 | FALSE |
| HSPA4 | 12557,952 | 27837,808 | -0,868 | 0,720 | FALSE |
| XYLT1 | 12176,166 | 26345,240 | -0,881 | 1,071 | FALSE |
| CTNNA2 | 34199,631 | 72233,573 | -0,885 | 0,620 | FALSE |
| MT1E | 72031,482 | 154300,372 | -0,909 | 1,000 | FALSE |
| ST13 | 13455,539 | 29792,461 | -0,912 | 1,051 | FALSE |

| | | | | | |
|---|---|---|---|---|---|
| FLG | 4799,360 | 10743,576 | -0,916 | 1,440 | FALSE |
| GET4 | 14053,997 | 30093,426 | -0,924 | 0,623 | FALSE |
| LGALS1 | 239695,429 | 534149,333 | -0,929 | 2,071 | FALSE |
| FSCN1 | 28056,775 | 63943,996 | -0,944 | 1,272 | FALSE |
| PKM | 135208,866 | 308931,605 | -0,946 | 1,454 | FALSE |
| GSTP1 | 471583,068 | 1072207,000 | -0,949 | 1,675 | FALSE |
| RAB1B | 17855,471 | 41488,996 | -0,951 | 0,710 | FALSE |
| RBM10 | 5004,133 | 11459,239 | -0,952 | 0,893 | FALSE |
| KRT73 | 32488,039 | 72695,954 | -0,955 | 0,407 | FALSE |
| NACA | 3730,280 | 8428,269 | -0,956 | 1,655 | FALSE |
| PPIA | 229091,512 | 537287,000 | -0,986 | 1,207 | FALSE |
| SERPINB12 | 41751,179 | 97292,319 | -1,016 | 1,136 | FALSE |
| ATP5PO | 12761,811 | 30167,225 | -1,019 | 1,182 | FALSE |
| NCOA5 | 81279,353 | 192422,702 | -1,028 | 0,718 | FALSE |
| NUP133 | 2005,397 | 4595,637 | -1,057 | 0,476 | FALSE |
| APOBEC3C | 168700,219 | 417666,042 | -1,083 | 0,634 | FALSE |
| INPP5K | 5963,379 | 15630,361 | -1,095 | 0,591 | FALSE |
| DNAJA2 | 8798,104 | 21822,155 | -1,096 | 1,036 | FALSE |
| STIP1 | 6043,718 | 15511,610 | -1,106 | 1,266 | FALSE |
| SLU7 | 2629,567 | 6168,737 | -1,145 | 0,375 | FALSE |
| MCM3 | 4774,292 | 12783,636 | -1,188 | 1,211 | FALSE |
| GTPBP4 | 2211,743 | 5520,452 | -1,195 | 0,441 | FALSE |
| ALB | 6407,517 | 16800,474 | -1,214 | 0,754 | FALSE |
| PFN2 | 287704,891 | 772650,115 | -1,226 | 0,635 | FALSE |
| IGKV4-1 | 19811,532 | 53889,204 | -1,230 | 1,038 | FALSE |
| AHSG | 8080,318 | 21939,244 | -1,233 | 0,838 | FALSE |
| SBSN | 5800,137 | 16272,911 | -1,245 | 1,114 | FALSE |
| GPN1 | 17256,866 | 47256,774 | -1,268 | 1,173 | FALSE |
| IGKC | 212641,423 | 556302,050 | -1,296 | 0,629 | FALSE |
| PA2G4 | 7140,630 | 21343,561 | -1,308 | 1,037 | FALSE |
| KRT77 | 69122,072 | 199361,255 | -1,324 | 0,540 | FALSE |
| DTX3L | 2377,384 | 7240,216 | -1,387 | 1,075 | FALSE |
| HRNR | 82052,697 | 261031,046 | -1,397 | 0,932 | FALSE |
| CTNND1 | 3123,135 | 9974,306 | -1,425 | 0,842 | FALSE |
| MDH2 | 9352,620 | 31001,088 | -1,455 | 1,125 | FALSE |
| LOC506828 | 1409,381 | 4852,890 | -1,577 | 1,137 | FALSE |
| ANXA1 | 5403,145 | 16665,088 | -1,582 | 0,344 | FALSE |
| ANP32B | 11512,779 | 53926,447 | -2,077 | 0,990 | FALSE |
| CNTNAP1 | 1149,301 | 8325,226 | -2,564 | 1,055 | FALSE |
| FKBP9 | 3941,840 | 71257,708 | -3,969 | 0,534 | FALSE |
