## Supplemental Table S2 for "SURF2 is a MDM2 antagonist in triggering the nucleolar stress response"

**Table_S2 Mass spectrometry analysis of treated cell (24 hours with Actinomycin D at 10nM ) U2OS expressing either flag only (control) or RPL5-FLAG**

| gene_name | iBAQ control | iBAQ RPL5 | log2_fc_Surf2-Control | log10_pval_Surf2-Control | significant_Surf2-Control |
|---|---|---|---|---|---|
| RPL11 | 46933,985 | 1644501,178 | 4,824 | 2,208 | TRUE |
| RPL5 | 103756,897 | 6594235,919 | 4,551 | 1,522 | TRUE |
| SURF2 | 7798,170 | 391984,769 | 3,858 | 2,019 | TRUE |
| MDM2 | 3241,447 | 303197,022 | 3,682 | 2,186 | TRUE |
| HEXIM1 | 3447,354 | 37083,969 | 3,387 | 2,853 | TRUE |
| SSB | 9484,518 | 266362,490 | 3,126 | 2,615 | TRUE |
| TP53 | 3019,414 | 54070,751 | 2,960 | 2,162 | TRUE |
| EIF2B1 | 4908,403 | 39299,389 | 2,545 | 3,525 | TRUE |
| UBE2O | 2040,542 | 9396,294 | 1,908 | 2,173 | TRUE |
| RPS26 | 68354,678 | 204531,378 | 1,655 | 1,929 | TRUE |
| PTCD3 | 507,245 | 2962,179 | 1,622 | 1,472 | TRUE |
| RPL23 | 44530,418 | 136129,886 | 1,568 | 1,510 | TRUE |
| MRPS25 | 3061,925 | 8400,060 | 1,457 | 1,390 | TRUE |
| MRPS27 | 3342,902 | 5906,837 | 1,127 | 2,139 | TRUE |
| MRPS22 | 7002,426 | 13629,732 | 1,074 | 1,411 | TRUE |
| RPF2 | 2972,031 | 8864,063 | 1,044 | 1,572 | TRUE |
| PRSS1 | 860997,244 | 39164,999 | -1,082 | 1,307 | TRUE |
| RPS27 | 15620,165 | 245488,018 | 3,184 | 0,533 | FALSE |
| KRT1 | 2742070,472 | 3212202,917 | 3,054 | 0,444 | FALSE |
| RPL13 | 16254,443 | 172357,933 | 2,890 | 0,520 | FALSE |
| KRT16 | 2202844,424 | 10298401,015 | 2,875 | 0,471 | FALSE |
| LGALS7 | 35876,138 | 438686,763 | 2,642 | 0,505 | FALSE |
| FABP5 | 34713,962 | 524056,841 | 2,562 | 0,502 | FALSE |
| KRT6B | 2252901,542 | 10213274,319 | 2,347 | 0,467 | FALSE |
| HEATR3 | 826,037 | 16020,655 | 2,345 | 0,730 | FALSE |
| KRT17 | 1323106,103 | 5117675,228 | 2,267 | 0,502 | FALSE |
| CALML3 | 30712,298 | 222619,957 | 2,181 | 0,444 | FALSE |
| RPL35 | 44269,350 | 84457,785 | 2,151 | 0,610 | FALSE |
| RPL18 | 9136,846 | 99617,279 | 2,090 | 0,548 | FALSE |
| H2AC18 | 237523,551 | 228372,462 | 2,042 | 0,490 | FALSE |
| RPL13A | 13394,906 | 49712,008 | 2,027 | 0,422 | FALSE |
| RPL24 | 8996,682 | 47924,255 | 2,013 | 0,901 | FALSE |
| RPL28 | 16583,319 | 92885,057 | 1,995 | 0,469 | FALSE |
| KRT6A | 1819869,056 | 10539546,813 | 1,988 | 0,534 | FALSE |
| RPL31 | 13355,616 | 88556,873 | 1,918 | 0,663 | FALSE |
| KRT4 | 510894,469 | 1588974,625 | 1,898 | 0,700 | FALSE |
| RPL6 | 13229,718 | 87297,082 | 1,847 | 0,603 | FALSE |

| | | | | | |
|---|---|---|---|---|---|
| RPL7 | 11750,481 | 70485,534 | 1,844 | 0,447 | FALSE |
| POLR2G | 68614,687 | 1425565,313 | 1,834 | 0,479 | FALSE |
| RPL4 | 6935,909 | 43819,564 | 1,820 | 0,554 | FALSE |
| KRT13 | 912619,516 | 3057125,441 | 1,753 | 0,665 | FALSE |
| RPL29 | 16035,239 | 94803,504 | 1,741 | 0,540 | FALSE |
| RPL34 | 19143,293 | 117445,588 | 1,649 | 0,350 | FALSE |
| RPL21 | 8457,865 | 28008,813 | 1,636 | 0,902 | FALSE |
| RPL8 | 13465,959 | 82919,054 | 1,609 | 0,434 | FALSE |
| CALML5 | 19161,248 | 115150,755 | 1,577 | 0,648 | FALSE |
| RPL27 | 29295,313 | 118635,318 | 1,576 | 0,438 | FALSE |
| PPIA | 3143,386 | 4547,002 | 1,571 | 0,830 | FALSE |
| HNRNPA2B1 | 4592,905 | 7858,152 | 1,559 | 0,545 | FALSE |
| KRT14 | 2327285,897 | 8774705,132 | 1,550 | 0,462 | FALSE |
| EPPK1 | 1331,585 | 2278,258 | 1,540 | 0,896 | FALSE |
| RPL9 | 14973,798 | 76618,581 | 1,528 | 0,523 | FALSE |
| PKP1 | 8507,346 | 71553,356 | 1,527 | 0,640 | FALSE |
| RPL3 | 6874,359 | 36922,400 | 1,502 | 0,551 | FALSE |
| RPLP2 | 89539,633 | 342695,802 | 1,470 | 0,401 | FALSE |
| KRT74 | 246064,025 | 659195,337 | 1,438 | 0,425 | FALSE |
| RPL14 | 7466,037 | 33712,434 | 1,433 | 0,542 | FALSE |
| SERBP1 | 6101,249 | 11042,133 | 1,431 | 0,558 | FALSE |
| SFN | 103307,383 | 145221,906 | 1,414 | 0,620 | FALSE |
| RPL10A | 20278,500 | 67107,121 | 1,401 | 0,710 | FALSE |
| KRT5 | 1801213,757 | 6885431,914 | 1,375 | 0,466 | FALSE |
| RPL26 | 41176,296 | 116570,423 | 1,369 | 0,548 | FALSE |
| H2AC4 | 261695,462 | 245997,197 | 1,346 | 0,395 | FALSE |
| H2AZ2 | 197552,508 | 177812,380 | 1,339 | 0,355 | FALSE |
| S100A11 | 39650,255 | 128319,997 | 1,336 | 0,724 | FALSE |
| MRPS14 | 3577,573 | 9088,764 | 1,314 | 1,199 | FALSE |
| FLG | 581,169 | 2830,156 | 1,306 | 0,719 | FALSE |
| S100A14 | 9885,779 | 28320,383 | 1,295 | 0,802 | FALSE |
| TPI1 | 11913,799 | 24035,865 | 1,294 | 1,112 | FALSE |
| TGM5 | 998,444 | 2116,375 | 1,287 | 0,728 | FALSE |
| CYB5B | 13798,149 | 38634,043 | 1,272 | 0,544 | FALSE |
| MRPS18B | 6489,086 | 12714,360 | 1,261 | 0,974 | FALSE |
| SERPINB3 | 21973,298 | 81969,517 | 1,260 | 0,616 | FALSE |
| TKT | 22058,704 | 15542,498 | 1,249 | 0,453 | FALSE |
| SLC2A1 | 2355,230 | 2488,819 | 1,238 | 0,276 | FALSE |
| RPL12 | 22595,369 | 69730,554 | 1,232 | 0,468 | FALSE |
| RPL23A | 20558,504 | 123662,997 | 1,218 | 0,302 | FALSE |
| NCCRP1 | 9627,591 | 17691,089 | 1,213 | 0,856 | FALSE |

| | | | | | |
|---|---|---|---|---|---|
| RPL7A | 14102,637 | 63381,612 | 1,197 | 0,591 | FALSE |
| RPL37A | 18423,750 | 158954,833 | 1,187 | 0,277 | FALSE |
| RPL22 | 36620,638 | 102867,956 | 1,185 | 0,511 | FALSE |
| H1-10 | 3570,991 | 23979,975 | 1,169 | 0,554 | FALSE |
| RPL30 | 17850,868 | 108455,798 | 1,163 | 0,376 | FALSE |
| WDR6 | 795,393 | 2080,083 | 1,158 | 0,875 | FALSE |
| PKM | 11774,281 | 31833,924 | 1,158 | 0,867 | FALSE |
| ASS1 | 2278,361 | 1476,555 | 1,151 | 0,272 | FALSE |
| DSP | 113813,044 | 272989,390 | 1,143 | 0,498 | FALSE |
| PFDN2 | 3451,001 | 6347,195 | 1,132 | 1,040 | FALSE |
| GSTP1 | 26403,835 | 75487,369 | 1,109 | 0,672 | FALSE |
| DSC3 | 895,121 | 1843,898 | 1,103 | 0,929 | FALSE |
| RPL27A | 17091,956 | 52340,137 | 1,049 | 0,410 | FALSE |
| JUP | 40693,905 | 114891,812 | 1,035 | 0,557 | FALSE |
| MRPS26 | 9105,917 | 23245,270 | 1,014 | 0,801 | FALSE |
| EEF2 | 12668,806 | 31133,756 | 0,999 | 0,552 | FALSE |
| RPS15 | 9071,902 | 17236,885 | 0,997 | 1,073 | FALSE |
| HNRNPUL2 | 9451,933 | 10353,768 | 0,985 | 0,694 | FALSE |
| CYCS | 14493,753 | 18608,929 | 0,975 | 0,313 | FALSE |
| RPS2 | 13904,998 | 35847,516 | 0,968 | 0,713 | FALSE |
| ENO1 | 13141,398 | 31207,341 | 0,959 | 0,716 | FALSE |
| LMO7 | 7185,115 | 10693,124 | 0,947 | 0,868 | FALSE |
| KRT19 | 1153991,826 | 3164564,242 | 0,946 | 0,422 | FALSE |
| HSPE1 | 4403,147 | 5388,144 | 0,945 | 0,408 | FALSE |
| ALDOA | 4738,044 | 11866,652 | 0,937 | 0,734 | FALSE |
| RPL35A | 29877,266 | 96464,518 | 0,937 | 0,357 | FALSE |
| COL1A2 | 2169,366 | 2227,941 | 0,921 | 0,253 | FALSE |
| MRPS15 | 3362,063 | 8990,982 | 0,920 | 0,800 | FALSE |
| RPS6 | 21733,835 | 50751,434 | 0,909 | 0,454 | FALSE |
| LDHA | 20230,181 | 35143,361 | 0,894 | 0,776 | FALSE |
| CTNNB1 | 3172,656 | 6764,953 | 0,893 | 0,466 | FALSE |
| POF1B | 4933,885 | 10476,148 | 0,876 | 0,803 | FALSE |
| Krt75 | 1333935,451 | 3681585,226 | 0,865 | 0,469 | FALSE |
| MRPS16 | 6486,686 | 10932,688 | 0,864 | 1,539 | FALSE |
| MRPS2 | 3688,343 | 9050,873 | 0,851 | 0,636 | FALSE |
| RPS23 | 41060,516 | 73258,391 | 0,825 | 1,056 | FALSE |
| HNRNPA1 | 12796,168 | 19498,924 | 0,815 | 0,552 | FALSE |
| MRPL22 | 2308,611 | 4716,670 | 0,810 | 1,223 | FALSE |
| GNG5 | 21828,505 | 18671,341 | 0,808 | 0,233 | FALSE |
| KRT3 | 617520,525 | 2331317,732 | 0,806 | 0,540 | FALSE |
| P4HB | 11596,016 | 15551,550 | 0,805 | 2,364 | FALSE |

| | | | | | |
|---|---|---|---|---|---|
| YBX1 | 92174,087 | 162003,739 | 0,802 | 0,805 | FALSE |
| LAS1L | 1993,377 | 1542,946 | 0,797 | 0,814 | FALSE |
| GAPDH | 122878,063 | 147579,155 | 0,789 | 0,433 | FALSE |
| MRPS9 | 4278,439 | 11050,638 | 0,781 | 0,575 | FALSE |
| MRPS28 | 8325,358 | 14441,165 | 0,756 | 1,268 | FALSE |
| RPL38 | 33702,318 | 101782,085 | 0,750 | 0,584 | FALSE |
| ATP5MK | 195176,187 | 72689,898 | 0,743 | 0,258 | FALSE |
| PRDX2 | 49002,449 | 77937,486 | 0,738 | 0,621 | FALSE |
| KCT2 | 3030,317 | 2486,553 | 0,729 | 0,431 | FALSE |
| LAMP1 | 17472,003 | 18124,420 | 0,705 | 0,461 | FALSE |
| ANXA1 | 15405,588 | 25912,269 | 0,693 | 0,437 | FALSE |
| HNRNPH3 | 20785,878 | 25230,344 | 0,691 | 0,350 | FALSE |
| DDX21 | 2002,038 | 2353,075 | 0,683 | 1,449 | FALSE |
| RPLP0 | 46027,631 | 87996,746 | 0,677 | 0,322 | FALSE |
| RPS24 | 29134,152 | 60611,357 | 0,673 | 0,792 | FALSE |
| ARG1 | 36654,181 | 70856,399 | 0,670 | 0,425 | FALSE |
| PSMB5 | 5961,664 | 7506,592 | 0,669 | 0,462 | FALSE |
| PSMA5 | 20652,572 | 22037,695 | 0,667 | 0,457 | FALSE |
| QPCTL | 6755,634 | 9038,389 | 0,665 | 0,465 | FALSE |
| B2M | 24752,301 | 25397,182 | 0,658 | 0,342 | FALSE |
| RPS19 | 74188,107 | 117879,813 | 0,657 | 0,664 | FALSE |
| MRPL39 | 1323,934 | 1895,310 | 0,651 | 1,518 | FALSE |
| CD44 | 5815,529 | 4858,864 | 0,649 | 0,379 | FALSE |
| EEF1E1 | 4370,178 | 4116,215 | 0,649 | 0,424 | FALSE |
| DAP3 | 7034,184 | 17061,266 | 0,645 | 0,430 | FALSE |
| YBX3 | 72050,095 | 115039,129 | 0,642 | 0,920 | FALSE |
| YWHAQ | 115399,424 | 100830,463 | 0,636 | 0,619 | FALSE |
| SEC61B | 89155,056 | 75508,383 | 0,633 | 0,343 | FALSE |
| HNRNPC | 66525,117 | 130491,921 | 0,629 | 0,356 | FALSE |
| TOP2A | 2335,949 | 2407,153 | 0,623 | 0,888 | FALSE |
| SLC1A5 | 2814,540 | 2608,828 | 0,621 | 0,450 | FALSE |
| HNRNPU | 36234,506 | 61865,539 | 0,613 | 0,709 | FALSE |
| NCL | 2070,281 | 2762,846 | 0,600 | 0,365 | FALSE |
| PSMB6 | 10189,399 | 12510,565 | 0,599 | 0,498 | FALSE |
| RAB5A | 9188,707 | 10724,446 | 0,591 | 0,552 | FALSE |
| ATP5PF | 29323,231 | 25312,921 | 0,585 | 0,251 | FALSE |
| PSMA6 | 8488,275 | 7747,819 | 0,578 | 0,351 | FALSE |
| RPS9 | 20857,978 | 38845,940 | 0,574 | 0,395 | FALSE |
| EIF4A1 | 13593,084 | 17418,772 | 0,573 | 0,829 | FALSE |
| CDSN | 13958,620 | 18700,378 | 0,568 | 0,455 | FALSE |
| DSG1 | 43122,655 | 89730,688 | 0,568 | 0,352 | FALSE |

| | | | | | |
|---|---|---|---|---|---|
| NDUFA4 | 54284,498 | 44383,410 | 0,565 | 0,307 | FALSE |
| RPS14 | 74460,686 | 128880,148 | 0,558 | 0,440 | FALSE |
| RBM4 | 3981,360 | 5154,088 | 0,557 | 0,492 | FALSE |
| S100A8 | 56983,450 | 53814,373 | 0,549 | 0,433 | FALSE |
| KRT80 | 29484,313 | 53848,917 | 0,541 | 0,648 | FALSE |
| RPS27A | 175160,518 | 258604,192 | 0,538 | 2,202 | FALSE |
| IGKV2-29 | 2148384,708 | 2835032,435 | 0,533 | 0,137 | FALSE |
| HNRNPL | 4246,417 | 4953,744 | 0,531 | 0,408 | FALSE |
| RPS12 | 47536,083 | 73829,071 | 0,530 | 0,346 | FALSE |
| RFC2 | 2458,424 | 2712,046 | 0,526 | 1,034 | FALSE |
| ILF3 | 6396,295 | 11077,542 | 0,519 | 0,263 | FALSE |
| S100A9 | 53247,597 | 52719,358 | 0,518 | 0,428 | FALSE |
| RPS7 | 51435,999 | 116861,033 | 0,516 | 0,366 | FALSE |
| RALY | 27676,976 | 41497,179 | 0,511 | 0,434 | FALSE |
| MBD3 | 3115,191 | 3760,759 | 0,511 | 0,408 | FALSE |
| ANKRD28 | 1106,973 | 1204,543 | 0,506 | 0,491 | FALSE |
| RPS21 | 70025,534 | 108904,434 | 0,505 | 0,345 | FALSE |
| RPSA | 84070,281 | 146037,915 | 0,505 | 0,308 | FALSE |
| COPRS | 256285,422 | 299930,318 | 0,505 | 0,497 | FALSE |
| TPM3 | 9826,946 | 12381,578 | 0,504 | 0,842 | FALSE |
| PSMB1 | 19581,892 | 18036,347 | 0,501 | 0,329 | FALSE |
| E2F7 | 17692,680 | 20372,752 | 0,498 | 0,640 | FALSE |
| RPS3A | 52643,221 | 78085,671 | 0,497 | 0,366 | FALSE |
| HSDL1 | 4920,318 | 4587,709 | 0,496 | 0,322 | FALSE |
| RPS8 | 44399,074 | 80265,000 | 0,494 | 0,271 | FALSE |
| HSP90B1 | 9026,674 | 7328,419 | 0,493 | 0,440 | FALSE |
| Krt79 | 1155673,617 | 3769812,922 | 0,493 | 0,429 | FALSE |
| PSMB4 | 9915,586 | 6977,837 | 0,490 | 0,216 | FALSE |
| G3BP1 | 3998,117 | 2499,208 | 0,489 | 0,678 | FALSE |
| CFL1 | 51226,913 | 49799,016 | 0,482 | 0,269 | FALSE |
| SLC25A6 | 117800,061 | 86389,876 | 0,481 | 0,210 | FALSE |
| MRPS23 | 10425,030 | 22635,250 | 0,478 | 0,329 | FALSE |
| TGM3 | 15416,035 | 17619,065 | 0,474 | 0,436 | FALSE |
| SLTM | 893,562 | 1996,283 | 0,472 | 0,495 | FALSE |
| RPS16 | 46722,306 | 67488,867 | 0,471 | 0,402 | FALSE |
| PODXL | 3212,088 | 2624,729 | 0,462 | 0,357 | FALSE |
| RPS20 | 82958,758 | 128056,110 | 0,461 | 0,390 | FALSE |
| MRPS7 | 7706,242 | 16087,360 | 0,456 | 0,365 | FALSE |
| CTSD | 4833,062 | 5463,576 | 0,454 | 0,378 | FALSE |
| LDHB | 16415,168 | 16950,292 | 0,453 | 0,665 | FALSE |
| RPS28 | 133149,433 | 198126,872 | 0,452 | 0,323 | FALSE |

| | | | | | |
|---|---|---|---|---|---|
| TPD52 | 32757,241 | 25874,191 | 0,450 | 0,345 | FALSE |
| RPS10 | 35262,469 | 50546,481 | 0,448 | 0,323 | FALSE |
| RPS11 | 71994,830 | 110561,504 | 0,447 | 0,296 | FALSE |
| RPS4X | 50636,253 | 75551,347 | 0,445 | 0,358 | FALSE |
| NME1 | 20787,184 | 28858,373 | 0,439 | 0,479 | FALSE |
| TRIM28 | 17379,592 | 19895,072 | 0,430 | 1,135 | FALSE |
| CPA4 | 2792,968 | 2744,621 | 0,428 | 0,354 | FALSE |
| RPS15A | 47315,353 | 74387,069 | 0,428 | 0,226 | FALSE |
| CSTA | 78436,251 | 109432,875 | 0,428 | 0,244 | FALSE |
| PRPS2 | 22755,822 | 17045,553 | 0,425 | 0,460 | FALSE |
| HNRNPD | 6611,820 | 9506,477 | 0,425 | 0,541 | FALSE |
| ATP5ME | 53915,231 | 47574,491 | 0,423 | 0,197 | FALSE |
| RPS13 | 67871,934 | 100121,881 | 0,423 | 0,270 | FALSE |
| PTS | 49347,047 | 47500,854 | 0,415 | 0,596 | FALSE |
| MAT2A | 20248,764 | 20430,391 | 0,410 | 0,375 | FALSE |
| PSMB3 | 5082,938 | 4895,735 | 0,409 | 0,199 | FALSE |
| BRAT1 | 1365,333 | 1171,999 | 0,407 | 0,161 | FALSE |
| L2HGDH | 4097,696 | 3496,172 | 0,405 | 0,752 | FALSE |
| ALCAM | 2932,032 | 2808,628 | 0,396 | 0,250 | FALSE |
| KTN1 | 5123,393 | 5168,354 | 0,396 | 0,662 | FALSE |
| GSTO1 | 11419,315 | 17023,681 | 0,395 | 0,884 | FALSE |
| DDX5 | 3450,024 | 3509,085 | 0,393 | 0,691 | FALSE |
| SETSIP | 6994,085 | 6787,652 | 0,391 | 0,511 | FALSE |
| RPS3 | 104618,736 | 149934,263 | 0,388 | 0,242 | FALSE |
| LMAN1 | 3035,418 | 5842,429 | 0,387 | 0,260 | FALSE |
| KRT18 | 631029,208 | 1127671,629 | 0,383 | 0,435 | FALSE |
| IGHG2 | 5006,002 | 6882,355 | 0,382 | 0,124 | FALSE |
| PGRMC1 | 142472,742 | 123345,391 | 0,372 | 0,262 | FALSE |
| RPS17 | 115245,598 | 169900,250 | 0,368 | 0,186 | FALSE |
| RPS25 | 153226,159 | 229773,817 | 0,364 | 0,258 | FALSE |
| DSC1 | 12379,819 | 20061,708 | 0,361 | 0,273 | FALSE |
| SMARCC1 | 1412,145 | 1384,849 | 0,359 | 0,493 | FALSE |
| KRT8 | 863691,367 | 2703141,352 | 0,359 | 0,342 | FALSE |
| PSMD5 | 2806,603 | 3333,512 | 0,357 | 0,264 | FALSE |
| FLNB | 22210,465 | 19516,178 | 0,355 | 1,201 | FALSE |
| UQCRC2 | 7299,984 | 7722,534 | 0,355 | 0,976 | FALSE |
| PTPN1 | 2518,933 | 1606,457 | 0,353 | 0,212 | FALSE |
| HRNR | 9018,703 | 12053,105 | 0,348 | 0,185 | FALSE |
| TPD52L2 | 3974,683 | 2961,120 | 0,348 | 0,217 | FALSE |
| CHD4 | 5559,659 | 5564,583 | 0,339 | 0,667 | FALSE |
| CAT | 13706,540 | 14306,534 | 0,329 | 0,223 | FALSE |

| | | | | | |
|---|---|---|---|---|---|
| PRSS3 | 50206,973 | 36687,218 | 0,328 | 0,134 | FALSE |
| RPS5 | 48510,818 | 65094,636 | 0,326 | 0,248 | FALSE |
| SLC25A11 | 3709,223 | 3593,777 | 0,326 | 0,213 | FALSE |
| KRT23 | 3080,813 | 24112,279 | 0,325 | 0,243 | FALSE |
| GSDMA | 14078,632 | 14256,258 | 0,323 | 0,231 | FALSE |
| PDHB | 4849,631 | 3951,411 | 0,322 | 0,440 | FALSE |
| TXN | 82477,205 | 101195,360 | 0,318 | 0,312 | FALSE |
| ATP5PO | 29041,699 | 29187,969 | 0,315 | 0,157 | FALSE |
| TGM1 | 5317,131 | 6282,378 | 0,314 | 0,339 | FALSE |
| CTPS1 | 5812,762 | 5353,778 | 0,312 | 0,394 | FALSE |
| ATP1B1 | 3057,931 | 12383,285 | 0,311 | 0,184 | FALSE |
| LGALS3 | 35410,828 | 44473,594 | 0,311 | 0,502 | FALSE |
| SDHA | 2514,203 | 3122,529 | 0,307 | 0,192 | FALSE |
| NUP107 | 3414,871 | 1890,878 | 0,305 | 0,309 | FALSE |
| CDC42 | 21396,439 | 14467,077 | 0,304 | 0,242 | FALSE |
| COL1A1 | 757,469 | 756,872 | 0,300 | 0,094 | FALSE |
| FLG2 | 14746,462 | 16033,317 | 0,296 | 0,201 | FALSE |
| AKAP8 | 7993,545 | 12441,179 | 0,294 | 0,125 | FALSE |
| SCARB2 | 11798,152 | 4007,891 | 0,290 | 0,107 | FALSE |
| HNRNPR | 5038,751 | 7394,826 | 0,283 | 0,154 | FALSE |
| PSMA4 | 37324,921 | 33057,999 | 0,282 | 0,255 | FALSE |
| SBSN | 1817,339 | 2184,651 | 0,282 | 0,244 | FALSE |
| IKBIP | 4217,028 | 3955,114 | 0,282 | 0,705 | FALSE |
| ARPC3 | 2749,934 | 7336,597 | 0,277 | 0,493 | FALSE |
| LMNA | 81640,740 | 97816,349 | 0,275 | 0,471 | FALSE |
| VAMP3 | 23338,640 | 18252,606 | 0,272 | 0,108 | FALSE |
| ATP6V1A | 1773,470 | 3818,025 | 0,271 | 0,153 | FALSE |
| SFPQ | 3661,448 | 2942,783 | 0,269 | 0,357 | FALSE |
| RALA | 16753,867 | 18083,787 | 0,267 | 0,208 | FALSE |
| PRDX6 | 15969,779 | 17710,950 | 0,265 | 0,481 | FALSE |
| MAX | 3496,335 | 2581,073 | 0,261 | 0,152 | FALSE |
| RACK1 | 41716,430 | 56501,021 | 0,258 | 0,201 | FALSE |
| PPT1 | 12237,530 | 11509,932 | 0,258 | 1,639 | FALSE |
| FLNA | 230465,160 | 229034,449 | 0,255 | 0,244 | FALSE |
| PSMD7 | 17226,625 | 15616,355 | 0,254 | 0,408 | FALSE |
| ALB | 8680,836 | 7542,742 | 0,254 | 0,147 | FALSE |
| PLEC | 65910,067 | 62055,864 | 0,253 | 0,350 | FALSE |
| HNRNPF | 140005,655 | 106024,667 | 0,251 | 0,483 | FALSE |
| MAEA | 6737,033 | 6524,156 | 0,248 | 0,697 | FALSE |
| RBBP7 | 9253,746 | 9775,954 | 0,243 | 0,410 | FALSE |
| CORO2A | 1370,421 | 1135,827 | 0,241 | 0,237 | FALSE |

| | | | | | |
|---|---|---|---|---|---|
| PRDX1 | 166375,697 | 196000,154 | 0,237 | 0,446 | FALSE |
| ZNF326 | 2493,323 | 3639,106 | 0,235 | 0,143 | FALSE |
| ATP2B1 | 495,306 | 1130,171 | 0,232 | 0,337 | FALSE |
| PLS3 | 3632,133 | 4581,876 | 0,219 | 0,119 | FALSE |
| FBL | 9371,584 | 9189,475 | 0,217 | 0,326 | FALSE |
| ATP5PD | 34653,444 | 27697,736 | 0,210 | 0,076 | FALSE |
| PSMD14 | 36900,139 | 33113,229 | 0,207 | 0,437 | FALSE |
| DLST | 3395,932 | 3378,040 | 0,207 | 0,195 | FALSE |
| TUBA4A | 632150,735 | 585791,800 | 0,205 | 0,099 | FALSE |
| SNAPIN | 10983,224 | 8432,436 | 0,204 | 0,125 | FALSE |
| VDAC2 | 30657,536 | 28471,301 | 0,204 | 0,328 | FALSE |
| AGL | 1544,232 | 1454,199 | 0,203 | 0,130 | FALSE |
| SERPINH1 | 2792,626 | 2805,974 | 0,202 | 0,201 | FALSE |
| EMD | 10436,295 | 11064,555 | 0,201 | 0,129 | FALSE |
| RBFOX1 | 10819,138 | 14766,300 | 0,200 | 0,121 | FALSE |
| HDX | 35615,292 | 41837,542 | 0,192 | 0,182 | FALSE |
| EIF3M | 8758,738 | 8270,618 | 0,190 | 0,171 | FALSE |
| RANBP10 | 37520,516 | 30429,792 | 0,190 | 0,172 | FALSE |
| ANXA2 | 149271,253 | 178860,085 | 0,183 | 0,265 | FALSE |
| LTF | 558,241 | 533,807 | 0,181 | 0,070 | FALSE |
| GNG12 | 61708,688 | 39131,292 | 0,176 | 0,068 | FALSE |
| SERPINB12 | 29789,685 | 24077,547 | 0,176 | 0,115 | FALSE |
| RPS18 | 68798,028 | 68953,818 | 0,175 | 0,276 | FALSE |
| OAT | 2381,244 | 2176,442 | 0,171 | 0,151 | FALSE |
| SLC3A2 | 18941,787 | 19632,953 | 0,170 | 0,101 | FALSE |
| VAPA | 19201,082 | 18228,048 | 0,169 | 0,159 | FALSE |
| MYO5B | 1922,570 | 1703,063 | 0,169 | 0,103 | FALSE |
| TUBB3 | 971755,685 | 849348,259 | 0,165 | 0,157 | FALSE |
| TET2 | 3542,370 | 2781,981 | 0,164 | 0,220 | FALSE |
| SRP68 | 2032,893 | 1887,283 | 0,163 | 0,125 | FALSE |
| TMX1 | 19674,925 | 18621,274 | 0,162 | 0,285 | FALSE |
| MTA1 | 1512,844 | 1550,260 | 0,161 | 0,207 | FALSE |
| DHX38 | 21242,945 | 20512,906 | 0,161 | 0,211 | FALSE |
| LAMTOR4 | 12677,077 | 11134,722 | 0,158 | 0,297 | FALSE |
| GANAB | 5298,247 | 4991,072 | 0,157 | 0,594 | FALSE |
| SMARCE1 | 4848,239 | 4467,938 | 0,157 | 0,090 | FALSE |
| ELAVL1 | 2412,370 | 2498,148 | 0,155 | 0,279 | FALSE |
| BMS1 | 6168,326 | 5336,291 | 0,154 | 0,130 | FALSE |
| HSPA1A | 196621,432 | 187781,432 | 0,153 | 0,230 | FALSE |
| HDAC1 | 23959,109 | 19632,123 | 0,153 | 0,087 | FALSE |
| EPRS1 | 2910,662 | 2332,322 | 0,151 | 0,256 | FALSE |

| | | | | | |
|---|---|---|---|---|---|
| SNRNP70 | 18732,496 | 16637,643 | 0,150 | 0,088 | FALSE |
| MTA2 | 6648,456 | 5664,232 | 0,143 | 0,210 | FALSE |
| STT3A | 2975,586 | 3253,129 | 0,141 | 0,073 | FALSE |
| EXOSC6 | 3108,908 | 2826,564 | 0,139 | 0,180 | FALSE |
| DNAJB2 | 18602,351 | 17183,584 | 0,138 | 0,340 | FALSE |
| MATR3 | 41364,075 | 40003,170 | 0,138 | 0,141 | FALSE |
| GID8 | 55388,505 | 45014,928 | 0,136 | 0,223 | FALSE |
| RBM39 | 18744,959 | 18177,472 | 0,135 | 0,103 | FALSE |
| TMED10 | 18398,806 | 17970,116 | 0,133 | 0,123 | FALSE |
| SSR1 | 7942,355 | 12242,587 | 0,132 | 0,098 | FALSE |
| COPB2 | 7954,635 | 6077,559 | 0,124 | 0,330 | FALSE |
| ODR4 | 3474,370 | 3140,085 | 0,123 | 0,096 | FALSE |
| HCFC1 | 857,913 | 741,186 | 0,121 | 0,092 | FALSE |
| PHF5A | 48796,147 | 36371,386 | 0,121 | 0,113 | FALSE |
| HNRNPM | 70260,814 | 65159,074 | 0,121 | 0,136 | FALSE |
| XRCC6 | 7348,164 | 6290,261 | 0,120 | 0,472 | FALSE |
| MTDH | 9807,335 | 12035,641 | 0,116 | 0,092 | FALSE |
| MRPS34 | 5338,326 | 7765,873 | 0,116 | 0,092 | FALSE |
| TOP2B | 2103,265 | 2082,521 | 0,112 | 0,145 | FALSE |
| PLP2 | 28934,787 | 36240,276 | 0,112 | 0,030 | FALSE |
| PCGF6 | 14337,850 | 13316,148 | 0,111 | 0,114 | FALSE |
| CSN1S2 | 49876,414 | 45145,016 | 0,110 | 0,071 | FALSE |
| LBR | 3525,732 | 2798,463 | 0,105 | 0,064 | FALSE |
| TFRC | 8942,953 | 8271,560 | 0,104 | 0,086 | FALSE |
| RANBP2 | 2400,455 | 1791,237 | 0,104 | 0,238 | FALSE |
| PPHLN1 | 7664,235 | 7644,705 | 0,104 | 0,104 | FALSE |
| EEF1A1 | 221886,805 | 193260,354 | 0,102 | 0,152 | FALSE |
| PGRMC2 | 29242,116 | 25625,364 | 0,102 | 0,083 | FALSE |
| RPN2 | 19130,535 | 19488,468 | 0,099 | 0,080 | FALSE |
| SF3B6 | 90446,520 | 60912,173 | 0,097 | 0,095 | FALSE |
| TUBB2B | 934168,040 | 799962,513 | 0,095 | 0,121 | FALSE |
| HSPH1 | 9410,281 | 7335,258 | 0,095 | 0,184 | FALSE |
| TUBB2A | 929944,858 | 797143,331 | 0,094 | 0,071 | FALSE |
| PRKCSH | 9854,329 | 8407,752 | 0,093 | 0,130 | FALSE |
| LMNB1 | 93070,604 | 79776,971 | 0,092 | 0,220 | FALSE |
| CCNT2 | 1636,223 | 1441,187 | 0,090 | 0,094 | FALSE |
| LUC7L3 | 36314,415 | 23568,961 | 0,089 | 0,038 | FALSE |
| CCAR2 | 45279,042 | 33346,140 | 0,089 | 0,117 | FALSE |
| VDAC3 | 9597,040 | 7839,837 | 0,087 | 0,131 | FALSE |
| EIF3H | 18670,084 | 15829,599 | 0,085 | 0,119 | FALSE |
| PSMD1 | 22481,620 | 18981,528 | 0,085 | 0,312 | FALSE |

| | | | | | |
|---|---|---|---|---|---|
| CSN2 | 124972,531 | 104922,629 | 0,083 | 0,048 | FALSE |
| BCAM | 2569,623 | 6707,101 | 0,081 | 0,032 | FALSE |
| TUBB6 | 454428,250 | 399041,764 | 0,081 | 0,076 | FALSE |
| HSP90AB1 | 84289,135 | 69656,265 | 0,080 | 0,220 | FALSE |
| SEC61G | 34704,950 | 20543,224 | 0,078 | 0,039 | FALSE |
| YIF1B | 4634,736 | 4419,010 | 0,076 | 0,069 | FALSE |
| FASN | 3273,388 | 3061,467 | 0,070 | 0,099 | FALSE |
| PFKP | 10955,990 | 6901,023 | 0,069 | 0,052 | FALSE |
| VIM | 268783,961 | 1130775,600 | 0,067 | 0,050 | FALSE |
| KPNA2 | 17984,231 | 15125,258 | 0,067 | 0,122 | FALSE |
| MCM7 | 13705,440 | 10852,759 | 0,066 | 0,090 | FALSE |
| BCL2L1 | 12326,675 | 6047,941 | 0,065 | 0,045 | FALSE |
| KRT78 | 203070,354 | 1097016,189 | 0,065 | 0,048 | FALSE |
| PCNA | 23451,714 | 16348,987 | 0,061 | 0,109 | FALSE |
| PSMC6 | 51488,134 | 47611,820 | 0,060 | 0,135 | FALSE |
| MYCBP | 1426926,411 | 1161190,071 | 0,057 | 0,042 | FALSE |
| SON | 6714,441 | 5570,751 | 0,055 | 0,043 | FALSE |
| CASP14 | 38291,630 | 32848,542 | 0,054 | 0,041 | FALSE |
| TUFM | 3432,613 | 2669,250 | 0,054 | 0,059 | FALSE |
| CAP1 | 25291,681 | 23122,974 | 0,052 | 0,074 | FALSE |
| H3-4 | 88382,809 | 90139,133 | 0,051 | 0,059 | FALSE |
| MAGEA4 | 18157,177 | 19747,626 | 0,051 | 0,087 | FALSE |
| AHNAK | 6095,635 | 5555,697 | 0,051 | 0,092 | FALSE |
| PSME3 | 5430,917 | 6610,314 | 0,050 | 0,025 | FALSE |
| CISD1 | 9994,653 | 11971,497 | 0,049 | 0,032 | FALSE |
| RBM17 | 7218,934 | 5082,178 | 0,046 | 0,026 | FALSE |
| RBM6 | 113257,826 | 92323,386 | 0,043 | 0,032 | FALSE |
| TUBG1 | 1965,880 | 1722,522 | 0,041 | 0,084 | FALSE |
| SAR1A | 2679,506 | 3299,403 | 0,040 | 0,025 | FALSE |
| HSPA6 | 143531,122 | 127817,718 | 0,038 | 0,041 | FALSE |
| ADRM1 | 13581,636 | 11211,180 | 0,037 | 0,069 | FALSE |
| ADNP | 1965,717 | 1544,385 | 0,036 | 0,020 | FALSE |
| SKP1 | 8151,810 | 6387,022 | 0,035 | 0,083 | FALSE |
| PSMD2 | 20816,504 | 18152,067 | 0,035 | 0,077 | FALSE |
| DDX3X | 9824,305 | 7647,026 | 0,034 | 0,064 | FALSE |
| LUC7L | 35621,992 | 21459,327 | 0,034 | 0,020 | FALSE |
| KRT1 | 8944184,400 | 8871237,086 | 0,031 | 0,024 | FALSE |
| CSNK2A1 | 5479,891 | 4007,672 | 0,028 | 0,022 | FALSE |
| SLC25A10 | 8470,818 | 8368,526 | 0,028 | 0,012 | FALSE |
| PURA | 7002,139 | 6154,055 | 0,027 | 0,020 | FALSE |
| PSMD4 | 22582,745 | 19532,114 | 0,024 | 0,033 | FALSE |

| | | | | | |
|---|---|---|---|---|---|
| VDAC1 | 39389,536 | 38171,222 | 0,022 | 0,026 | FALSE |
| TUBAL3 | 66987,797 | 79326,700 | 0,018 | 0,010 | FALSE |
| RAN | 33937,027 | 33910,826 | 0,018 | 0,021 | FALSE |
| RCN1 | 16194,935 | 13478,821 | 0,015 | 0,093 | FALSE |
| HSPA4 | 5167,568 | 3909,891 | 0,014 | 0,025 | FALSE |
| TWF1 | 3750,792 | 4101,552 | 0,014 | 0,008 | FALSE |
| ACAP2 | 26341,160 | 19727,258 | 0,013 | 0,012 | FALSE |
| ATP5F1B | 85000,790 | 83448,828 | 0,013 | 0,007 | FALSE |
| NAP1L1 | 21352,988 | 26626,105 | 0,013 | 0,009 | FALSE |
| PRPF19 | 10854,051 | 8552,718 | 0,013 | 0,034 | FALSE |
| HSPB1 | 441672,389 | 450428,195 | 0,012 | 0,010 | FALSE |
| PSMA7 | 7387,244 | 6663,359 | 0,009 | 0,009 | FALSE |
| WDR26 | 21272,785 | 15041,008 | 0,007 | 0,007 | FALSE |
| AIFM1 | 1616,585 | 1782,783 | 0,006 | 0,006 | FALSE |
| COPA | 4654,800 | 4031,251 | 0,006 | 0,012 | FALSE |
| PPP1CB | 8758,315 | 9986,713 | 0,002 | 0,002 | FALSE |
| PSMD8 | 10940,651 | 9502,091 | 0,001 | 0,001 | FALSE |
| SMC4 | 3550,354 | 2584,482 | -0,002 | 0,004 | FALSE |
| PSMC3 | 48611,304 | 46477,772 | -0,004 | 0,007 | FALSE |
| CSN3 | 40203,425 | 34427,051 | -0,005 | 0,005 | FALSE |
| RPN1 | 25238,858 | 24152,275 | -0,005 | 0,004 | FALSE |
| BSG | 10846,278 | 9752,703 | -0,006 | 0,006 | FALSE |
| CKAP4 | 11825,741 | 18666,167 | -0,006 | 0,004 | FALSE |
| VAPB | 17275,148 | 15439,871 | -0,009 | 0,008 | FALSE |
| FRYL | 2304,779 | 1703,406 | -0,010 | 0,012 | FALSE |
| IARS1 | 5808,186 | 4502,798 | -0,012 | 0,022 | FALSE |
| EIF3C | 14060,345 | 11806,893 | -0,013 | 0,020 | FALSE |
| ATP1A1 | 9557,145 | 7847,138 | -0,013 | 0,009 | FALSE |
| GATAD2B | 3559,476 | 4418,348 | -0,015 | 0,010 | FALSE |
| SLC25A3 | 49362,115 | 45525,532 | -0,016 | 0,008 | FALSE |
| PSMC5 | 50391,321 | 42093,233 | -0,016 | 0,040 | FALSE |
| RANBP9 | 11245,300 | 9329,905 | -0,017 | 0,017 | FALSE |
| GLG1 | 4964,481 | 1963,960 | -0,017 | 0,011 | FALSE |
| TAB2 | 45639,702 | 35629,281 | -0,019 | 0,019 | FALSE |
| SPCS2 | 6728,785 | 5490,874 | -0,021 | 0,021 | FALSE |
| RTN1 | 3119,070 | 3391,949 | -0,021 | 0,019 | FALSE |
| RAB7A | 34604,918 | 28731,975 | -0,022 | 0,024 | FALSE |
| PFKFB2 | 35919,554 | 25438,076 | -0,022 | 0,034 | FALSE |
| MT-ATP6 | 21084,264 | 49588,937 | -0,026 | 0,011 | FALSE |
| SNU13 | 7351,435 | 5627,166 | -0,027 | 0,025 | FALSE |
| MGA | 23043,455 | 17440,449 | -0,027 | 0,024 | FALSE |

| | | | | | |
|---|---|---|---|---|---|
| H2BC4 | 194489,037 | 255963,523 | -0,030 | 0,045 | FALSE |
| MAGED2 | 25651,821 | 21884,911 | -0,032 | 0,052 | FALSE |
| SLC7A5 | 19893,449 | 13034,350 | -0,038 | 0,016 | FALSE |
| TUBA3C | 929048,060 | 844179,739 | -0,042 | 0,040 | FALSE |
| SF3B4 | 26733,794 | 19920,979 | -0,043 | 0,050 | FALSE |
| TACC3 | 2645,341 | 3297,726 | -0,044 | 0,028 | FALSE |
| EIF3D | 7341,561 | 7557,218 | -0,045 | 0,024 | FALSE |
| TARDBP | 8525,389 | 11326,481 | -0,045 | 0,025 | FALSE |
| PCBP1 | 10494,730 | 10290,043 | -0,046 | 0,036 | FALSE |
| KCTD2 | 122202,137 | 91136,188 | -0,047 | 0,022 | FALSE |
| CCAR1 | 8729,134 | 5740,336 | -0,047 | 0,053 | FALSE |
| ATP5MG | 44431,076 | 39101,381 | -0,049 | 0,016 | FALSE |
| SUPT16H | 2634,255 | 1983,377 | -0,051 | 0,033 | FALSE |
| TOMM70 | 6652,852 | 2957,830 | -0,052 | 0,033 | FALSE |
| P4HA1 | 2628,805 | 3571,869 | -0,052 | 0,043 | FALSE |
| ERG28 | 11193,591 | 6915,600 | -0,055 | 0,046 | FALSE |
| SF3B2 | 32205,815 | 24485,618 | -0,055 | 0,059 | FALSE |
| DARS1 | 15627,049 | 12355,674 | -0,057 | 0,205 | FALSE |
| HADHB | 41163,032 | 32515,630 | -0,057 | 0,083 | FALSE |
| CLINT1 | 3660,205 | 4157,101 | -0,058 | 0,025 | FALSE |
| CDK1 | 19368,100 | 18458,620 | -0,059 | 0,068 | FALSE |
| PHGDH | 30491,578 | 29308,146 | -0,063 | 0,029 | FALSE |
| DHRS2 | 117612,113 | 97079,263 | -0,064 | 0,061 | FALSE |
| KARS1 | 5755,213 | 5278,594 | -0,064 | 0,040 | FALSE |
| UQCRC1 | 9074,642 | 4349,794 | -0,064 | 0,021 | FALSE |
| FDFT1 | 5020,099 | 4147,821 | -0,064 | 0,042 | FALSE |
| HSP90AA1 | 50704,470 | 40162,593 | -0,067 | 0,173 | FALSE |
| KIF11 | 834083,878 | 769855,891 | -0,067 | 0,051 | FALSE |
| MYL6 | 97289,648 | 66723,877 | -0,068 | 0,057 | FALSE |
| NOP56 | 6527,507 | 7266,244 | -0,069 | 0,051 | FALSE |
| EIF3F | 44866,788 | 38623,290 | -0,069 | 0,132 | FALSE |
| RCN2 | 32920,497 | 46503,886 | -0,069 | 0,042 | FALSE |
| SURF4 | 16309,886 | 14428,665 | -0,071 | 0,101 | FALSE |
| PFKL | 5919,990 | 4369,532 | -0,075 | 0,036 | FALSE |
| SCYL2 | 23758,298 | 17249,203 | -0,078 | 0,101 | FALSE |
| ATP5F1C | 17878,911 | 16132,620 | -0,082 | 0,068 | FALSE |
| SNRPA1 | 17311,041 | 10734,593 | -0,082 | 0,113 | FALSE |
| CANX | 33278,580 | 28435,370 | -0,083 | 0,084 | FALSE |
| PSMD3 | 28403,201 | 23495,515 | -0,083 | 0,224 | FALSE |
| ATP5PB | 25524,561 | 22950,278 | -0,084 | 0,029 | FALSE |
| SLC4A1AP | 10279,381 | 15765,922 | -0,084 | 0,091 | FALSE |

| | | | | | |
|---|---|---|---|---|---|
| KPRP | 23547,060 | 19322,325 | -0,086 | 0,053 | FALSE |
| MOB2 | 63006,585 | 55937,476 | -0,088 | 0,076 | FALSE |
| OGT | 5758,407 | 4368,048 | -0,088 | 0,107 | FALSE |
| NT5C2 | 11828,953 | 13951,763 | -0,089 | 0,054 | FALSE |
| PRKAR2A | 3511,368 | 3964,948 | -0,089 | 0,061 | FALSE |
| PSMC2 | 41509,652 | 33660,794 | -0,094 | 0,367 | FALSE |
| SEC13 | 3268,589 | 3126,929 | -0,095 | 0,081 | FALSE |
| SYNGR2 | 15865,143 | 15649,294 | -0,096 | 0,048 | FALSE |
| DYNLL1 | 81533,315 | 69721,175 | -0,096 | 0,088 | FALSE |
| PSME1 | 33532,277 | 28195,520 | -0,096 | 0,180 | FALSE |
| BLOC1S2 | 80523,222 | 55386,570 | -0,097 | 0,175 | FALSE |
| ILF2 | 14461,955 | 18926,517 | -0,097 | 0,064 | FALSE |
| LMNB2 | 34195,520 | 29032,024 | -0,099 | 0,281 | FALSE |
| ARFGAP1 | 7273,969 | 8624,584 | -0,099 | 0,067 | FALSE |
| CBX3 | 28084,342 | 23036,714 | -0,099 | 0,084 | FALSE |
| CD2BP2 | 9512,881 | 7011,443 | -0,100 | 0,074 | FALSE |
| HSPA8 | 573145,377 | 413640,981 | -0,102 | 0,132 | FALSE |
| EIF3A | 13563,730 | 9302,101 | -0,102 | 0,217 | FALSE |
| PSMC4 | 50102,205 | 41335,142 | -0,103 | 0,254 | FALSE |
| PPM1A | 26641,065 | 48328,707 | -0,103 | 0,081 | FALSE |
| H4C1 | 437708,185 | 329270,933 | -0,106 | 0,225 | FALSE |
| PSMC1 | 43644,929 | 37941,765 | -0,107 | 0,292 | FALSE |
| DCD | 329736,246 | 271107,466 | -0,107 | 0,048 | FALSE |
| ALG13 | 1043,283 | 708,448 | -0,108 | 0,070 | FALSE |
| HSPA9 | 126361,010 | 95025,093 | -0,109 | 0,164 | FALSE |
| SEC22B | 68028,778 | 54040,185 | -0,109 | 0,126 | FALSE |
| PSMD6 | 16795,021 | 13164,398 | -0,114 | 0,233 | FALSE |
| RAB14 | 36192,746 | 28628,519 | -0,114 | 0,162 | FALSE |
| PGAM5 | 9666,339 | 6247,417 | -0,115 | 0,125 | FALSE |
| EPHX1 | 8394,493 | 4106,587 | -0,116 | 0,087 | FALSE |
| PRKDC | 7980,777 | 6176,776 | -0,118 | 0,097 | FALSE |
| PPP2R1A | 5551,945 | 4633,344 | -0,118 | 0,073 | FALSE |
| HNRNPUL1 | 4964,829 | 3880,568 | -0,118 | 0,096 | FALSE |
| ZBTB24 | 8376,716 | 11124,746 | -0,119 | 0,092 | FALSE |
| KPNB1 | 38178,475 | 28867,254 | -0,119 | 0,190 | FALSE |
| RARS1 | 6876,336 | 5613,919 | -0,121 | 0,073 | FALSE |
| SF3B3 | 48075,963 | 30097,497 | -0,122 | 0,145 | FALSE |
| DDOST | 14808,987 | 21386,561 | -0,125 | 0,086 | FALSE |
| GBA | 2956,904 | 2680,466 | -0,127 | 0,099 | FALSE |
| SFXN3 | 4250,056 | 3039,575 | -0,127 | 0,076 | FALSE |
| RAB10 | 51886,432 | 36686,524 | -0,128 | 0,158 | FALSE |

| | | | | | |
|---|---|---|---|---|---|
| PSME2 | 39830,935 | 33092,290 | -0,131 | 0,187 | FALSE |
| TNPO1 | 10496,570 | 6502,629 | -0,135 | 0,252 | FALSE |
| ORMDL1 | 16309,045 | 13375,013 | -0,136 | 0,086 | FALSE |
| WARS1 | 12428,107 | 8643,883 | -0,136 | 0,219 | FALSE |
| TAF12 | 300397,368 | 200762,090 | -0,140 | 0,160 | FALSE |
| U2SURP | 5876,917 | 4991,047 | -0,141 | 0,198 | FALSE |
| SNRNP200 | 8410,163 | 5785,545 | -0,142 | 0,216 | FALSE |
| NUP93 | 5115,695 | 4020,706 | -0,143 | 0,086 | FALSE |
| BLMH | 7460,656 | 7756,030 | -0,144 | 0,134 | FALSE |
| RAB1A | 115224,060 | 80109,624 | -0,144 | 0,145 | FALSE |
| MLEC | 8319,399 | 9497,651 | -0,144 | 0,093 | FALSE |
| ZBED5 | 2797,138 | 1915,205 | -0,147 | 0,102 | FALSE |
| KRT9 | 3416016,900 | 3766429,200 | -0,148 | 0,146 | FALSE |
| CALU | 14381,813 | 11571,948 | -0,148 | 0,269 | FALSE |
| NSF | 10049,024 | 8169,859 | -0,149 | 0,082 | FALSE |
| ARCN1 | 7426,541 | 5499,732 | -0,150 | 0,098 | FALSE |
| DLG1 | 6019,417 | 4160,062 | -0,151 | 0,481 | FALSE |
| UQCRQ | 12317,307 | 7538,479 | -0,152 | 0,138 | FALSE |
| SFXN1 | 7398,580 | 9014,783 | -0,153 | 0,069 | FALSE |
| EPB41L3 | 28338,321 | 21191,982 | -0,153 | 0,183 | FALSE |
| HSPA5 | 181595,639 | 151366,922 | -0,154 | 0,209 | FALSE |
| EIF3E | 34375,252 | 23091,326 | -0,154 | 0,411 | FALSE |
| EIF3G | 20038,868 | 15020,148 | -0,155 | 0,217 | FALSE |
| NELFB | 3314,588 | 2478,210 | -0,158 | 0,123 | FALSE |
| TPR | 605,487 | 459,267 | -0,159 | 0,136 | FALSE |
| EIF3I | 48180,685 | 34880,505 | -0,160 | 0,205 | FALSE |
| SRP14 | 22900,116 | 20013,722 | -0,160 | 0,346 | FALSE |
| GCN1 | 5425,122 | 3496,805 | -0,163 | 0,270 | FALSE |
| LARS1 | 3152,058 | 1632,385 | -0,164 | 0,252 | FALSE |
| HNRNPK | 315759,692 | 268570,022 | -0,164 | 0,183 | FALSE |
| HADHA | 39001,034 | 29388,309 | -0,168 | 0,222 | FALSE |
| ARMC8 | 17417,178 | 12038,836 | -0,168 | 0,173 | FALSE |
| HSPD1 | 219799,404 | 169952,603 | -0,168 | 0,364 | FALSE |
| S100A7 | 51233,471 | 46537,648 | -0,169 | 0,178 | FALSE |
| MAP1B | 64366,555 | 46138,552 | -0,169 | 0,170 | FALSE |
| NCAPH | 1657,601 | 1999,977 | -0,171 | 0,224 | FALSE |
| PAF1 | 4474,049 | 3158,122 | -0,172 | 0,108 | FALSE |
| RHOA | 21551,893 | 15887,890 | -0,173 | 0,153 | FALSE |
| CAD | 9992,908 | 6446,199 | -0,175 | 0,285 | FALSE |
| SKA1 | 42802,509 | 29881,572 | -0,175 | 0,197 | FALSE |
| CTTN | 6632,788 | 7883,583 | -0,176 | 0,131 | FALSE |

| | | | | | |
|---|---|---|---|---|---|
| FLII | 590,298 | 416,177 | -0,176 | 0,369 | FALSE |
| MAP3K7 | 70196,625 | 53468,055 | -0,177 | 0,206 | FALSE |
| RAP1A | 17667,965 | 13760,664 | -0,179 | 0,151 | FALSE |
| RTN3 | 3862,490 | 3442,198 | -0,181 | 0,174 | FALSE |
| VPS16 | 1085,027 | 854,449 | -0,181 | 0,355 | FALSE |
| DDRGK1 | 5095,935 | 3695,697 | -0,181 | 0,212 | FALSE |
| APMAP | 21979,685 | 16448,720 | -0,182 | 0,457 | FALSE |
| PABPC4 | 24393,473 | 15749,424 | -0,184 | 0,157 | FALSE |
| ADAR | 1640,797 | 1395,501 | -0,184 | 0,122 | FALSE |
| DAD1 | 22718,177 | 36007,395 | -0,186 | 0,080 | FALSE |
| ROCK1 | 21084,438 | 14440,286 | -0,186 | 0,206 | FALSE |
| PPP1CA | 6782,344 | 7799,154 | -0,188 | 0,278 | FALSE |
| HNRNPH1 | 720155,329 | 488223,207 | -0,189 | 0,209 | FALSE |
| RCL1 | 30635,427 | 20724,062 | -0,191 | 0,252 | FALSE |
| NUP98 | 1119,112 | 811,962 | -0,191 | 0,182 | FALSE |
| SPINDOC | 384588,534 | 289607,178 | -0,193 | 0,183 | FALSE |
| H2BC3 | 175321,957 | 236372,666 | -0,194 | 0,339 | FALSE |
| SEL1L | 1039,724 | 1024,407 | -0,195 | 0,210 | FALSE |
| KIF5B | 5199,512 | 4139,605 | -0,196 | 0,109 | FALSE |
| NES | 12756,112 | 8112,625 | -0,198 | 0,359 | FALSE |
| REST | 18451,364 | 11727,760 | -0,199 | 0,174 | FALSE |
| ATP5F1A | 86889,441 | 75247,704 | -0,199 | 0,132 | FALSE |
| TMEM97 | 8707,059 | 7189,421 | -0,200 | 0,193 | FALSE |
| LUC7L2 | 50810,833 | 29034,219 | -0,200 | 0,118 | FALSE |
| CSNK2B | 6018,195 | 6123,966 | -0,201 | 0,110 | FALSE |
| RUVBL2 | 56030,665 | 41231,791 | -0,204 | 0,599 | FALSE |
| JAK1 | 43244,659 | 27383,829 | -0,210 | 0,270 | FALSE |
| S100A16 | 150400,988 | 99645,653 | -0,210 | 0,203 | FALSE |
| ESYT1 | 12257,429 | 8024,161 | -0,210 | 0,484 | FALSE |
| CAMSAP3 | 29716,888 | 22009,068 | -0,212 | 0,346 | FALSE |
| AIMP1 | 6235,093 | 4912,162 | -0,212 | 0,141 | FALSE |
| LRPPRC | 5867,647 | 4051,726 | -0,212 | 0,274 | FALSE |
| TAB3 | 40835,405 | 28344,773 | -0,215 | 0,210 | FALSE |
| L3MBTL2 | 1554,915 | 1657,903 | -0,215 | 0,152 | FALSE |
| NUP133 | 2046,518 | 1391,444 | -0,217 | 0,174 | FALSE |
| GSN | 8768,333 | 10843,796 | -0,218 | 0,140 | FALSE |
| PPM1B | 51214,545 | 63077,582 | -0,219 | 0,255 | FALSE |
| CHP1 | 7950,353 | 5152,352 | -0,220 | 0,290 | FALSE |
| VAT1 | 46717,249 | 35270,213 | -0,220 | 0,558 | FALSE |
| SART1 | 29128,303 | 20694,220 | -0,220 | 0,261 | FALSE |
| PCYOX1 | 11716,616 | 8832,856 | -0,220 | 0,412 | FALSE |

| | | | | | |
|---|---|---|---|---|---|
| SNRPB2 | 53191,503 | 34174,842 | -0,222 | 0,330 | FALSE |
| ITGA3 | 4324,203 | 4805,419 | -0,224 | 0,212 | FALSE |
| DHX9 | 4935,092 | 5863,126 | -0,226 | 0,166 | FALSE |
| PDHA1 | 2595,204 | 2086,129 | -0,228 | 0,218 | FALSE |
| IVNS1ABP | 88474,461 | 56581,870 | -0,228 | 0,121 | FALSE |
| NIPSNAP1 | 11288,915 | 7883,241 | -0,229 | 0,775 | FALSE |
| PSMD13 | 32667,492 | 21933,870 | -0,231 | 0,745 | FALSE |
| EIF3L | 25272,517 | 16895,492 | -0,231 | 0,795 | FALSE |
| SPIN1 | 604918,172 | 463676,389 | -0,232 | 0,212 | FALSE |
| PSMD12 | 26482,782 | 19318,604 | -0,232 | 0,770 | FALSE |
| EFTUD2 | 14175,873 | 12080,352 | -0,232 | 0,476 | FALSE |
| XRCC5 | 6575,193 | 4669,828 | -0,233 | 0,141 | FALSE |
| XP32 | 17937,541 | 10653,317 | -0,236 | 0,163 | FALSE |
| IMPDH2 | 7610,035 | 4576,745 | -0,237 | 0,578 | FALSE |
| CCDC88A | 20137,562 | 13633,431 | -0,238 | 0,289 | FALSE |
| EIF3K | 12673,364 | 8577,693 | -0,239 | 0,480 | FALSE |
| GLUD1 | 2395,567 | 2618,197 | -0,240 | 0,252 | FALSE |
| GNB2 | 44878,388 | 31320,518 | -0,240 | 0,217 | FALSE |
| SRP72 | 1472,068 | 950,851 | -0,241 | 0,227 | FALSE |
| SEC61A1 | 34563,606 | 24151,910 | -0,242 | 0,221 | FALSE |
| XPO1 | 10087,965 | 6182,039 | -0,243 | 0,378 | FALSE |
| TNRC6B | 17116,562 | 11167,903 | -0,243 | 0,263 | FALSE |
| RAB11B | 99369,130 | 78386,540 | -0,245 | 0,402 | FALSE |
| EIF3B | 42886,235 | 26636,548 | -0,245 | 0,409 | FALSE |
| CYB5R1 | 2064,347 | 2125,563 | -0,246 | 0,478 | FALSE |
| AZGP1 | 15924,178 | 13327,190 | -0,247 | 0,263 | FALSE |
| ANKFY1 | 51844,680 | 37691,587 | -0,247 | 0,310 | FALSE |
| COPB1 | 3665,648 | 3004,946 | -0,249 | 0,175 | FALSE |
| TUBB | 1358368,449 | 1166622,692 | -0,253 | 0,213 | FALSE |
| SPCS3 | 10367,829 | 11800,049 | -0,254 | 0,200 | FALSE |
| GNA12 | 5761,719 | 4419,737 | -0,255 | 0,140 | FALSE |
| WTAP | 5396,721 | 5443,184 | -0,255 | 0,204 | FALSE |
| SRSF2 | 169140,596 | 63025,953 | -0,258 | 0,152 | FALSE |
| ADM | 25920,560 | 14444,142 | -0,259 | 0,154 | FALSE |
| GTF2I | 1775,037 | 1183,713 | -0,260 | 0,241 | FALSE |
| CAP2 | 34244,147 | 24827,694 | -0,260 | 0,524 | FALSE |
| KRT2 | 4526199,070 | 4860574,186 | -0,265 | 0,205 | FALSE |
| SNAP29 | 3954,144 | 4134,635 | -0,266 | 0,263 | FALSE |
| C1QBP | 14577,756 | 17852,748 | -0,266 | 0,249 | FALSE |
| RUVBL1 | 94300,419 | 56153,721 | -0,266 | 0,394 | FALSE |
| ITGAV | 1469,851 | 1788,610 | -0,268 | 0,162 | FALSE |

| | | | | | |
|---|---|---|---|---|---|
| PSMD11 | 35484,621 | 25693,475 | -0,270 | 0,759 | FALSE |
| DYNC1H1 | 12019,434 | 7544,808 | -0,271 | 0,525 | FALSE |
| POLR2B | 1951,856 | 1279,315 | -0,271 | 0,144 | FALSE |
| TAF5 | 3390,125 | 2126,692 | -0,277 | 0,202 | FALSE |
| IPO4 | 827,015 | 659,520 | -0,278 | 0,379 | FALSE |
| ATP5MF | 31066,472 | 43539,046 | -0,279 | 0,106 | FALSE |
| YTHDF2 | 1945,308 | 1742,590 | -0,280 | 0,212 | FALSE |
| RAB32 | 7525,016 | 5037,644 | -0,280 | 0,197 | FALSE |
| AGO2 | 3726,772 | 3579,664 | -0,281 | 0,319 | FALSE |
| TAB1 | 268574,365 | 183779,656 | -0,282 | 0,293 | FALSE |
| MAPK6 | 1345,546 | 1038,627 | -0,284 | 0,194 | FALSE |
| PNN | 6566,633 | 4296,204 | -0,285 | 0,189 | FALSE |
| KRT77 | 1379687,206 | 1641899,801 | -0,286 | 0,205 | FALSE |
| ATP2A2 | 6284,240 | 5071,200 | -0,287 | 0,153 | FALSE |
| SMARCA4 | 2000,944 | 1273,622 | -0,287 | 0,207 | FALSE |
| ARL6IP5 | 47066,784 | 32231,024 | -0,288 | 0,752 | FALSE |
| CSE1L | 66203,488 | 41002,691 | -0,290 | 0,510 | FALSE |
| SPIN3 | 267800,000 | 199132,605 | -0,291 | 0,391 | FALSE |
| YPEL5 | 14416,260 | 9005,090 | -0,292 | 0,250 | FALSE |
| SNRPF | 212438,945 | 108868,313 | -0,294 | 0,448 | FALSE |
| PRPF8 | 4410,723 | 2496,614 | -0,297 | 0,576 | FALSE |
| MYH9 | 93532,518 | 64449,320 | -0,298 | 0,431 | FALSE |
| WDR1 | 4465,732 | 2321,011 | -0,299 | 0,298 | FALSE |
| ACTN1 | 40970,723 | 22948,497 | -0,300 | 0,332 | FALSE |
| TUBB4B | 1186256,438 | 1008830,151 | -0,300 | 0,133 | FALSE |
| BCAP31 | 53253,690 | 39936,237 | -0,302 | 0,681 | FALSE |
| SEC31A | 3141,229 | 2271,106 | -0,304 | 0,236 | FALSE |
| PRPSAP1 | 19016,855 | 17891,723 | -0,304 | 0,167 | FALSE |
| OTUD4 | 212157,390 | 137508,031 | -0,304 | 0,219 | FALSE |
| PCBP2 | 8546,840 | 7494,083 | -0,305 | 0,450 | FALSE |
| TUBA1C | 1367593,397 | 1173189,076 | -0,306 | 0,205 | FALSE |
| SYNRG | 14118,338 | 8671,439 | -0,308 | 0,393 | FALSE |
| EIF4A3 | 11230,691 | 8905,154 | -0,309 | 0,260 | FALSE |
| KATNAL2 | 5960,402 | 3929,534 | -0,309 | 0,437 | FALSE |
| ZW10 | 1054,297 | 940,616 | -0,310 | 0,545 | FALSE |
| RAB1B | 120397,163 | 82695,688 | -0,312 | 0,547 | FALSE |
| SPIN2B | 90704,578 | 62928,118 | -0,313 | 0,388 | FALSE |
| RTN4 | 10139,709 | 6475,996 | -0,313 | 0,304 | FALSE |
| SRPRB | 12741,398 | 8471,583 | -0,313 | 0,963 | FALSE |
| SPTAN1 | 77347,970 | 49933,623 | -0,314 | 0,438 | FALSE |
| KRT10 | 8390652,071 | 10029423,536 | -0,314 | 0,327 | FALSE |

| | | | | | |
|---|---|---|---|---|---|
| RBBP4 | 41253,827 | 33780,471 | -0,315 | 0,528 | FALSE |
| MSH6 | 7361,003 | 8319,750 | -0,317 | 0,295 | FALSE |
| HDAC2 | 29134,686 | 24464,833 | -0,318 | 0,274 | FALSE |
| SMC2 | 2803,568 | 1886,623 | -0,320 | 0,240 | FALSE |
| CTNNA1 | 4060,599 | 3166,978 | -0,321 | 0,167 | FALSE |
| UQCR10 | 26766,033 | 26585,257 | -0,322 | 0,131 | FALSE |
| SPTBN1 | 83084,474 | 51880,525 | -0,322 | 0,440 | FALSE |
| HLA-C | 12123,632 | 10832,576 | -0,323 | 0,376 | FALSE |
| TRIP13 | 6458,345 | 4615,799 | -0,326 | 0,225 | FALSE |
| DYNLRB1 | 30221,932 | 13957,329 | -0,327 | 0,396 | FALSE |
| FLNC | 118887,193 | 77319,817 | -0,332 | 0,426 | FALSE |
| CAVIN1 | 19437,901 | 18634,718 | -0,332 | 0,261 | FALSE |
| PPIG | 12222,902 | 6563,285 | -0,332 | 0,215 | FALSE |
| TOR1AIP1 | 2365,547 | 2602,557 | -0,333 | 0,448 | FALSE |
| STX7 | 7947,629 | 16764,068 | -0,334 | 0,135 | FALSE |
| PRPF6 | 6953,818 | 3446,205 | -0,336 | 0,835 | FALSE |
| VARS1 | 6709,821 | 4489,100 | -0,336 | 0,192 | FALSE |
| CEP70 | 7309,693 | 7935,695 | -0,338 | 0,265 | FALSE |
| SLC25A1 | 4739,589 | 5978,655 | -0,340 | 0,244 | FALSE |
| PPIL4 | 13691,048 | 13300,860 | -0,344 | 0,256 | FALSE |
| PHB2 | 24493,283 | 29869,136 | -0,348 | 0,230 | FALSE |
| CSN1S1 | 99372,175 | 68241,184 | -0,348 | 0,357 | FALSE |
| CCT6B | 47480,626 | 32441,268 | -0,349 | 0,327 | FALSE |
| CCT4 | 108956,805 | 80593,796 | -0,350 | 0,505 | FALSE |
| EEF1D | 87047,830 | 55631,570 | -0,350 | 1,092 | FALSE |
| USP15 | 182375,220 | 120923,643 | -0,351 | 0,431 | FALSE |
| PHB | 43692,679 | 50934,065 | -0,351 | 0,183 | FALSE |
| CDK9 | 15250,729 | 14865,495 | -0,351 | 0,329 | FALSE |
| PLIN3 | 41348,208 | 47958,479 | -0,353 | 0,478 | FALSE |
| YWHAZ | 183388,766 | 166500,848 | -0,353 | 0,344 | FALSE |
| MAP1A | 795,178 | 644,372 | -0,354 | 0,469 | FALSE |
| CCT6A | 155336,432 | 106897,064 | -0,354 | 0,664 | FALSE |
| EEF1B2 | 46092,290 | 36085,168 | -0,354 | 1,013 | FALSE |
| ACTA1 | 4629000,880 | 1427085,731 | -0,355 | 0,405 | FALSE |
| RRBP1 | 3058,689 | 2929,891 | -0,358 | 0,363 | FALSE |
| GET4 | 10459,219 | 6179,114 | -0,359 | 0,301 | FALSE |
| SF3B1 | 41212,554 | 25723,203 | -0,360 | 0,470 | FALSE |
| PRPF31 | 974444,500 | 725384,686 | -0,364 | 0,323 | FALSE |
| SSR4 | 32086,430 | 20881,352 | -0,364 | 0,394 | FALSE |
| STK38L | 946032,245 | 646248,501 | -0,365 | 0,438 | FALSE |
| DDB1 | 7154,715 | 7375,591 | -0,370 | 0,372 | FALSE |

| | | | | | |
|---|---|---|---|---|---|
| EEF1A2 | 189606,034 | 161032,191 | -0,371 | 0,785 | FALSE |
| RAP2B | 23638,540 | 20274,866 | -0,371 | 0,172 | FALSE |
| STK38 | 1477151,005 | 933791,458 | -0,373 | 0,246 | FALSE |
| LETM1 | 1626,528 | 1708,383 | -0,373 | 0,278 | FALSE |
| SF3A3 | 1932,319 | 1078,321 | -0,374 | 0,437 | FALSE |
| CMBL | 31971,829 | 18076,707 | -0,375 | 0,392 | FALSE |
| RIF1 | 11435,484 | 6205,441 | -0,376 | 0,365 | FALSE |
| CCT3 | 119295,430 | 85628,797 | -0,376 | 0,704 | FALSE |
| TRIM21 | 167646,901 | 99452,654 | -0,376 | 0,396 | FALSE |
| DCTN3 | 82471,234 | 40473,486 | -0,377 | 0,604 | FALSE |
| CIRBP | 5043,732 | 3854,542 | -0,378 | 0,307 | FALSE |
| MOGS | 3904,763 | 4346,346 | -0,379 | 0,231 | FALSE |
| TIMM50 | 10819,616 | 7482,131 | -0,380 | 0,268 | FALSE |
| ACTR1B | 76070,831 | 38773,933 | -0,380 | 0,558 | FALSE |
| KCTD17 | 60682,249 | 44056,979 | -0,381 | 0,534 | FALSE |
| PSAP | 10429,726 | 3776,023 | -0,381 | 0,210 | FALSE |
| TCP1 | 131422,521 | 88430,236 | -0,382 | 0,733 | FALSE |
| CLTC | 44700,033 | 31456,018 | -0,383 | 0,497 | FALSE |
| FOXP4 | 2020,044 | 1658,785 | -0,385 | 0,413 | FALSE |
| ACSL3 | 3378,155 | 2251,685 | -0,385 | 0,355 | FALSE |
| HSD17B11 | 4974,741 | 4788,229 | -0,386 | 0,375 | FALSE |
| PPP1CC | 7063,151 | 6887,119 | -0,389 | 0,499 | FALSE |
| USO1 | 4200,990 | 2809,784 | -0,389 | 0,306 | FALSE |
| IPO9 | 1564,101 | 2749,452 | -0,390 | 0,299 | FALSE |
| SCFD1 | 3320,505 | 3576,007 | -0,390 | 0,289 | FALSE |
| RAB6B | 31880,058 | 20171,378 | -0,392 | 0,640 | FALSE |
| GEMIN4 | 3382,387 | 3148,515 | -0,393 | 0,356 | FALSE |
| AFTPH | 4007,408 | 3141,401 | -0,397 | 0,216 | FALSE |
| TAF4 | 324258,051 | 220481,980 | -0,398 | 0,439 | FALSE |
| DNAJA2 | 6239,640 | 6690,768 | -0,399 | 0,394 | FALSE |
| CKAP5 | 95261,278 | 60745,271 | -0,401 | 0,385 | FALSE |
| RBM5 | 214183,221 | 130893,166 | -0,401 | 0,447 | FALSE |
| CSTF2 | 5025,818 | 5364,549 | -0,402 | 0,586 | FALSE |
| GNAI2 | 17506,961 | 11684,279 | -0,406 | 0,266 | FALSE |
| MOB1A | 62808,756 | 46853,156 | -0,408 | 0,918 | FALSE |
| TMEM33 | 17119,376 | 19820,110 | -0,410 | 0,326 | FALSE |
| TPP1 | 6289,508 | 2421,749 | -0,412 | 0,452 | FALSE |
| CYB5R3 | 32100,758 | 20294,951 | -0,413 | 0,625 | FALSE |
| SLC25A5 | 240103,064 | 164183,787 | -0,415 | 0,265 | FALSE |
| PUF60 | 17067,686 | 13309,440 | -0,415 | 0,293 | FALSE |
| SCD | 10091,268 | 9853,726 | -0,415 | 0,390 | FALSE |

| | | | | | |
|---|---|---|---|---|---|
| TMPO | 22507,454 | 20483,749 | -0,416 | 0,501 | FALSE |
| SNRPGP15 | 333874,318 | 188991,969 | -0,416 | 0,825 | FALSE |
| ZC3H18 | 19143,742 | 9408,348 | -0,418 | 0,233 | FALSE |
| ZYX | 40230,858 | 23445,839 | -0,418 | 0,518 | FALSE |
| AGO3 | 4533,474 | 4322,000 | -0,419 | 0,348 | FALSE |
| ARHGAP21 | 3148,505 | 2775,597 | -0,420 | 0,340 | FALSE |
| SLAIN2 | 173950,443 | 107824,437 | -0,421 | 0,702 | FALSE |
| AGPS | 6292,921 | 3880,745 | -0,422 | 0,282 | FALSE |
| PFKFB3 | 87296,462 | 57461,934 | -0,426 | 0,494 | FALSE |
| STAT3 | 25123,136 | 13027,222 | -0,427 | 0,517 | FALSE |
| NUP160 | 819,647 | 486,129 | -0,427 | 1,100 | FALSE |
| SMARCC2 | 2366,161 | 2297,294 | -0,427 | 0,673 | FALSE |
| RIOK1 | 67321,745 | 37417,375 | -0,428 | 0,624 | FALSE |
| CCT7 | 125829,867 | 83383,742 | -0,429 | 0,707 | FALSE |
| CCT8 | 146447,582 | 101089,324 | -0,429 | 0,604 | FALSE |
| ITGB1 | 9969,321 | 11734,835 | -0,430 | 0,250 | FALSE |
| MMS19 | 1796,697 | 1029,406 | -0,431 | 0,487 | FALSE |
| DCTN1 | 22960,142 | 12703,509 | -0,433 | 0,890 | FALSE |
| COPG2 | 2563,488 | 1780,006 | -0,434 | 0,410 | FALSE |
| GOLT1B | 11400,798 | 14157,103 | -0,437 | 0,382 | FALSE |
| MYL12A | 134437,697 | 77449,153 | -0,438 | 0,559 | FALSE |
| LGALS1 | 49037,169 | 36197,242 | -0,438 | 0,509 | FALSE |
| DNAJA1 | 16699,321 | 17201,676 | -0,440 | 0,451 | FALSE |
| SYMPK | 1064,954 | 951,493 | -0,443 | 0,496 | FALSE |
| NOP58 | 11082,565 | 10843,953 | -0,445 | 0,441 | FALSE |
| SRPRA | 1191,373 | 894,497 | -0,449 | 0,999 | FALSE |
| BZW1 | 11538,065 | 12023,871 | -0,451 | 0,451 | FALSE |
| RRP9 | 2224,024 | 848,436 | -0,454 | 0,520 | FALSE |
| MT1E | 50156,034 | 15109,620 | -0,454 | 0,297 | FALSE |
| RNPS1 | 26231,208 | 12395,369 | -0,457 | 0,250 | FALSE |
| CCT5 | 230940,512 | 142913,801 | -0,457 | 0,610 | FALSE |
| TAF6 | 4502,445 | 4298,606 | -0,465 | 0,528 | FALSE |
| LRRC59 | 68694,896 | 45995,796 | -0,470 | 0,701 | FALSE |
| VCP | 75674,826 | 45718,001 | -0,470 | 0,855 | FALSE |
| RBM10 | 3063772,043 | 1950061,217 | -0,473 | 0,406 | FALSE |
| MT-CO2 | 8592,262 | 8783,430 | -0,476 | 0,557 | FALSE |
| HLA-A | 12010,000 | 10189,328 | -0,479 | 0,307 | FALSE |
| EEF1G | 97205,294 | 58539,987 | -0,482 | 0,837 | FALSE |
| EIF4B | 7303889,500 | 4704441,701 | -0,483 | 0,530 | FALSE |
| SRRT | 37481,851 | 19750,806 | -0,483 | 0,328 | FALSE |
| NCAPG | 4151,226 | 2438,063 | -0,484 | 0,373 | FALSE |

| | | | | | |
|---|---|---|---|---|---|
| MCM3 | 1591,369 | 1401,466 | -0,485 | 0,786 | FALSE |
| DIABLO | 9258,916 | 4262,386 | -0,489 | 0,682 | FALSE |
| TUBA4B | 47598,305 | 41281,589 | -0,490 | 0,447 | FALSE |
| DHX15 | 42154,367 | 22201,135 | -0,491 | 0,348 | FALSE |
| MVP | 14723,722 | 7276,599 | -0,493 | 0,214 | FALSE |
| ZC3H13 | 3208,253 | 1226,194 | -0,493 | 0,245 | FALSE |
| TMEM109 | 17240,693 | 11274,307 | -0,496 | 1,102 | FALSE |
| ZC3HAV1 | 1128,501 | 1103,326 | -0,498 | 0,631 | FALSE |
| DCTN4 | 13504,887 | 5782,612 | -0,500 | 0,994 | FALSE |
| ATL3 | 6331,839 | 3645,206 | -0,502 | 0,348 | FALSE |
| CPSF6 | 8532,857 | 206017,278 | -0,504 | 0,250 | FALSE |
| EIF2B4 | 3038,298 | 2914,901 | -0,506 | 0,613 | FALSE |
| HEATR1 | 1287,651 | 1157,470 | -0,506 | 0,636 | FALSE |
| PTGES2 | 9101,841 | 5205,316 | -0,514 | 1,684 | FALSE |
| GPD2 | 3839,670 | 2278,233 | -0,514 | 0,353 | FALSE |
| IPO7 | 6199,918 | 4768,518 | -0,519 | 0,255 | FALSE |
| NAMPT | 37185,257 | 23038,471 | -0,521 | 0,776 | FALSE |
| WDR77 | 5872379,531 | 3103608,943 | -0,523 | 0,326 | FALSE |
| ACTR10 | 171598,543 | 143203,461 | -0,527 | 1,021 | FALSE |
| SNRPE | 387035,531 | 232965,773 | -0,529 | 0,567 | FALSE |
| CCT2 | 115830,946 | 75129,634 | -0,530 | 0,635 | FALSE |
| DCTN2 | 95605,477 | 51224,080 | -0,531 | 1,291 | FALSE |
| ACTR1A | 124919,169 | 64471,309 | -0,531 | 1,001 | FALSE |
| YWHAH | 105603,295 | 79496,326 | -0,532 | 0,793 | FALSE |
| TUBB4A | 798460,222 | 673618,759 | -0,533 | 0,813 | FALSE |
| NSDHL | 6801,328 | 6164,075 | -0,538 | 0,704 | FALSE |
| PRMT5 | 6279361,897 | 3630736,404 | -0,539 | 0,464 | FALSE |
| RAB15 | 22369,144 | 14827,507 | -0,542 | 0,508 | FALSE |
| RAB33B | 26842,973 | 17793,009 | -0,542 | 0,508 | FALSE |
| WDR5 | 6400,579 | 4890,032 | -0,543 | 0,542 | FALSE |
| LYZ | 31353,814 | 18191,183 | -0,544 | 0,309 | FALSE |
| PRMT1 | 94587,829 | 50068,694 | -0,555 | 0,397 | FALSE |
| CAPRIN2 | 22304,502 | 11070,924 | -0,555 | 0,650 | FALSE |
| BAG2 | 11633,169 | 9660,289 | -0,561 | 0,626 | FALSE |
| ZDBF2 | 9011,513 | 8627,083 | -0,565 | 0,485 | FALSE |
| SLC25A13 | 4597,186 | 4862,907 | -0,568 | 0,259 | FALSE |
| BZW2 | 3939,321 | 3449,593 | -0,569 | 0,646 | FALSE |
| SF3A1 | 2772,455 | 2507,850 | -0,570 | 0,663 | FALSE |
| MYOF | 1010,867 | 458,438 | -0,571 | 0,523 | FALSE |
| SNRPD2 | 2236521,750 | 1315065,113 | -0,576 | 1,820 | FALSE |
| ARGLU1 | 127588,094 | 54960,072 | -0,580 | 0,285 | FALSE |

| | | | | | |
|---|---|---|---|---|---|
| HM13 | 18582,485 | 17736,304 | -0,586 | 0,606 | FALSE |
| KCTD5 | 420826,327 | 252521,373 | -0,586 | 0,535 | FALSE |
| PRPSAP2 | 12225,120 | 9456,359 | -0,588 | 0,535 | FALSE |
| CAPZB | 58819,790 | 27718,936 | -0,589 | 0,682 | FALSE |
| MYH14 | 11311,429 | 7313,709 | -0,591 | 0,722 | FALSE |
| PABPC1 | 54500,690 | 32664,472 | -0,593 | 0,600 | FALSE |
| CALM1 | 83796,657 | 82023,225 | -0,597 | 0,226 | FALSE |
| RAD50 | 1889,545 | 1523,554 | -0,601 | 0,733 | FALSE |
| RAB2A | 19286,543 | 17734,896 | -0,606 | 0,574 | FALSE |
| PRPS1 | 30276,419 | 19126,387 | -0,614 | 0,452 | FALSE |
| STOML2 | 12277,264 | 11417,839 | -0,615 | 0,591 | FALSE |
| MOV10 | 5650,674 | 5237,161 | -0,617 | 0,390 | FALSE |
| BAX | 52576,116 | 28270,298 | -0,620 | 1,542 | FALSE |
| HMOX2 | 5950,391 | 10855,756 | -0,625 | 0,439 | FALSE |
| CAPZA1 | 111262,826 | 52458,549 | -0,626 | 0,742 | FALSE |
| OCIAD1 | 12370,831 | 12654,140 | -0,628 | 0,306 | FALSE |
| YWHAB | 145581,049 | 109393,712 | -0,631 | 0,975 | FALSE |
| MRE11 | 4385,550 | 3985,391 | -0,634 | 0,772 | FALSE |
| CHERP | 4138,814 | 3098,463 | -0,640 | 0,834 | FALSE |
| CRTAP | 2289,551 | 1176,915 | -0,641 | 0,980 | FALSE |
| Trypsin | ########### | 98239149,333 | -0,642 | 0,493 | FALSE |
| REEP5 | 8460,858 | 5431,099 | -0,649 | 1,076 | FALSE |
| SNRPA | 103932,411 | 62997,270 | -0,653 | 0,465 | FALSE |
| SRSF4 | 295910,517 | 93954,661 | -0,662 | 0,447 | FALSE |
| PRAF2 | 17190,866 | 15351,879 | -0,662 | 0,447 | FALSE |
| CADPS2 | 7714,404 | 6030,681 | -0,662 | 0,554 | FALSE |
| PTRH2 | 16235,119 | 9741,175 | -0,665 | 0,469 | FALSE |
| SNRPB | 82410,283 | 45364,323 | -0,666 | 0,528 | FALSE |
| RAB5C | 48012,841 | 29545,938 | -0,669 | 0,926 | FALSE |
| GOLGA3 | 3754,902 | 2638,470 | -0,669 | 0,648 | FALSE |
| ACADVL | 5193,106 | 5263,747 | -0,672 | 0,694 | FALSE |
| SNRPD1 | 549502,008 | 376124,921 | -0,676 | 0,698 | FALSE |
| RANGAP1 | 7923,409 | 6755,441 | -0,676 | 0,841 | FALSE |
| CLNS1A | 804523,477 | 463134,946 | -0,678 | 1,180 | FALSE |
| SRSF10 | 27336,904 | 13449,296 | -0,680 | 0,386 | FALSE |
| IPO8 | 53911,350 | 27622,825 | -0,690 | 0,660 | FALSE |
| FIP1L1 | 2462,945 | 1712,061 | -0,691 | 0,479 | FALSE |
| LGB | 71878,547 | 48547,114 | -0,695 | 0,798 | FALSE |
| YWHAE | 190027,348 | 122095,535 | -0,700 | 1,195 | FALSE |
| MAGOH | 10949,527 | 10662,656 | -0,702 | 0,566 | FALSE |
| CD9 | 48916,234 | 47307,750 | -0,705 | 0,272 | FALSE |

| | | | | | |
|---|---|---|---|---|---|
| YWHAG | 144389,497 | 99301,506 | -0,710 | 1,174 | FALSE |
| TMOD3 | 64579,361 | 27512,511 | -0,720 | 0,819 | FALSE |
| PDGFA | 19391,546 | 14067,747 | -0,726 | 0,725 | FALSE |
| RAB18 | 6061,268 | 5350,112 | -0,729 | 1,275 | FALSE |
| LIMA1 | 25527,504 | 8796,340 | -0,732 | 0,862 | FALSE |
| ACTN4 | 31893,758 | 18065,607 | -0,732 | 0,726 | FALSE |
| TTYH3 | 4144,452 | 2661,508 | -0,733 | 0,945 | FALSE |
| PRPF40A | 4727,455 | 1279,563 | -0,734 | 0,385 | FALSE |
| DPM1 | 17007,238 | 16421,856 | -0,739 | 0,874 | FALSE |
| DHRS7 | 10785,647 | 9444,721 | -0,761 | 0,551 | FALSE |
| RAB34 | 8288,282 | 7243,678 | -0,764 | 0,993 | FALSE |
| BET1 | 23449,223 | 15744,033 | -0,765 | 0,599 | FALSE |
| DDX23 | 2183,866 | 1855,031 | -0,771 | 1,319 | FALSE |
| RDH11 | 16352,060 | 8831,139 | -0,779 | 0,541 | FALSE |
| SUN2 | 86080,280 | 39100,184 | -0,779 | 0,533 | FALSE |
| SVIL | 32956,870 | 9219,023 | -0,807 | 0,612 | FALSE |
| DPP7 | 281888,930 | 141253,743 | -0,815 | 2,177 | FALSE |
| DCTN5 | 16100,491 | 9058,017 | -0,818 | 1,059 | FALSE |
| ERGIC3 | 9430,169 | 5023,255 | -0,821 | 1,263 | FALSE |
| CAPZA2 | 31151,172 | 20767,188 | -0,832 | 1,153 | FALSE |
| NCBP1 | 5692,388 | 3271,967 | -0,835 | 0,442 | FALSE |
| M6PR | 5733,187 | 5182,408 | -0,842 | 0,414 | FALSE |
| TRAP1 | 1911,579 | 934,929 | -0,847 | 0,910 | FALSE |
| SSR3 | 15070,883 | 12088,871 | -0,848 | 0,696 | FALSE |
| MYH10 | 20775,983 | 11912,098 | -0,854 | 0,726 | FALSE |
| TRA2A | 53357,334 | 16906,951 | -0,866 | 0,505 | FALSE |
| PLAA | 5141,800 | 1789,635 | -0,870 | 0,609 | FALSE |
| IGHA1 | 7807,831 | 5698,909 | -0,902 | 0,406 | FALSE |
| ABLIM1 | 455763,647 | 300296,116 | -0,904 | 0,737 | FALSE |
| NUDT21 | 19169,251 | 13026,215 | -0,906 | 0,808 | FALSE |
| ACTG1 | 6318905,554 | 2013970,837 | -0,907 | 0,896 | FALSE |
| DYNC1I2 | 14738,796 | 9411,285 | -0,910 | 0,905 | FALSE |
| ZRANB2 | 10775,846 | 6532,168 | -0,916 | 1,265 | FALSE |
| HNRNPH2 | 628959,969 | 605150,302 | -0,928 | 1,196 | FALSE |
| SNRPD3 | 82249,497 | 42250,165 | -0,948 | 0,793 | FALSE |
| SRRM1 | 7574,115 | 4400,746 | -0,956 | 0,547 | FALSE |
| NEXN | 13801,688 | 3321,620 | -0,967 | 1,083 | FALSE |
| CLU | 13539,332 | 13367,883 | -0,973 | 0,513 | FALSE |
| THRAP3 | 391846,621 | 146396,640 | -0,984 | 0,452 | FALSE |
| ACIN1 | 42572,754 | 14808,489 | -1,017 | 0,500 | FALSE |
| CORO1C | 71634,763 | 14787,500 | -1,020 | 1,284 | FALSE |

| | | | | | |
|---|---|---|---|---|---|
| SRSF5 | 219613,999 | 75823,719 | -1,029 | 0,520 | FALSE |
| SRRM2 | 35567,892 | 14319,495 | -1,033 | 0,554 | FALSE |
| NCBP2 | 15111,204 | 7585,215 | -1,064 | 0,572 | FALSE |
| DBN1 | 18726,445 | 4771,861 | -1,068 | 1,284 | FALSE |
| TRA2B | 133315,059 | 52915,948 | -1,073 | 0,605 | FALSE |
| SRSF9 | 122365,067 | 44179,237 | -1,077 | 0,544 | FALSE |
| SRSF3 | 491812,005 | 146531,699 | -1,083 | 0,545 | FALSE |
| SRSF7 | 337164,540 | 100855,228 | -1,146 | 0,533 | FALSE |
| MAP2 | 985,988 | 624,038 | -1,178 | 0,868 | FALSE |
| CD63 | 11439,511 | 6767,748 | -1,182 | 0,671 | FALSE |
| GPRC5A | 13846,159 | 5993,518 | -1,185 | 1,234 | FALSE |
| BCLAF1 | 323976,224 | 94608,929 | -1,200 | 0,639 | FALSE |
| SRSF6 | 505604,784 | 158707,126 | -1,215 | 0,573 | FALSE |
| MYO1B | 25718,586 | 4895,057 | -1,233 | 0,505 | FALSE |
| PARP2 | 3056,298 | #DIV/0! | -1,234 | 0,493 | FALSE |
| SRSF1 | 805163,727 | 233242,620 | -1,331 | 0,587 | FALSE |
| ERH | 6882206,958 | 2120219,750 | -1,452 | 0,783 | FALSE |
| SAP18 | 230306,189 | 59572,758 | -1,566 | 0,607 | FALSE |
| ACTB | 6350620,880 | 2017309,043 | -1,815 | 1,201 | FALSE |
| MYO1C | 69990,040 | 11195,077 | -1,850 | 0,508 | FALSE |
| APOD | 36787,992 | 16323,718 | -2,079 | 0,381 | FALSE |
| FMR1 | 9742,418 | 3092,136 | -2,126 | 1,029 | FALSE |
| PIP | 10038,677 | 3364,609 | -2,237 | 0,584 | FALSE |
| STX3 | 6525,241 | 2439,473 | -3,009 | 0,517 | FALSE |
