## Supplemental Table S1 for "SURF2 is a MDM2 antagonist in triggering the nucleolar stress response"

**TABLE_S1 Mass spectrometry analysis of untreated U2OS expressing either flag only (control) or RPL5-FLAG**

| gene_name | iBAQ control | iBAQ RPL5 | log2_fc_Surf2-Control | log10_pval_Surf2-Control | significant_Surf2-Control |
|---|---|---|---|---|---|
| RPL5 | 35327.800 | 9306328.706 | 5.886 | 2.643 | TRUE |
| MDM2 | 1117.400 | 310414.910 | 5.870 | 4.701 | TRUE |
| SURF2 | 9754.587 | 1353395.932 | 5.514 | 2.097 | TRUE |
| RPL11 | 55403.464 | 2499797.533 | 5.060 | 2.857 | TRUE |
| SSB | 20231.409 | 461173.537 | 3.492 | 5.497 | TRUE |
| HEXIM1 | 3075.896 | 51933.554 | 3.132 | 2.964 | TRUE |
| UBE2O | 1379.214 | 15661.651 | 2.593 | 2.297 | TRUE |
| TP53 | 931.828 | 12426.932 | 2.384 | 2.361 | TRUE |
| EIF2B1 | 8382.039 | 68248.268 | 2.299 | 2.431 | TRUE |
| PTCD3 | 1549.516 | 4834.738 | 1.425 | 2.414 | TRUE |
| MRPS15 | 5193.185 | 13895.296 | 1.375 | 1.951 | TRUE |
| MRPS14 | 3685.028 | 10385.069 | 1.352 | 2.587 | TRUE |
| MRPS18B | 10612.217 | 24579.522 | 1.301 | 2.498 | TRUE |
| LMO7 | 18070.205 | 39177.274 | 1.150 | 2.182 | TRUE |
| MRPS26 | 8370.225 | 21386.454 | 1.145 | 1.992 | TRUE |
| MRPS25 | 6433.330 | 14073.665 | 1.128 | 1.617 | TRUE |
| MRPS22 | 10711.903 | 23326.339 | 1.082 | 1.777 | TRUE |
| PIP | 11551.391 | 29158.631 | 1.030 | 1.376 | TRUE |
| MRPS16 | 9215.140 | 19087.059 | 1.003 | 1.798 | TRUE |
| MAP2 | 2398.497 | 1173.095 | -1.634 | 1.401 | TRUE |
| HEATR3 | 5055.964 | 88810.255 | 3.378 | 0.974 | FALSE |
| PRSS3 | 37388.156 | 81269.672 | 2.776 | 0.401 | FALSE |
| H2AC18 | 353211.077 | 443707.238 | 2.281 | 0.509 | FALSE |
| MRPL11 | 2954.034 | 39726.702 | 2.068 | 0.409 | FALSE |
| TUBB8 | 904635.394 | 1047709.576 | 1.556 | 0.791 | FALSE |
| RBFOX1 | 16904.113 | 23078.427 | 1.384 | 0.851 | FALSE |
| H2AC7 | 392106.488 | 463259.774 | 1.316 | 0.436 | FALSE |
| FLG | 612.111 | 862.069 | 1.233 | 0.588 | FALSE |
| XP32 | 19621.343 | 32157.799 | 1.145 | 0.611 | FALSE |
| TUBAL3 | 122410.843 | 176546.625 | 1.089 | 0.620 | FALSE |
| AKAP8 | 14679.453 | 27851.558 | 1.080 | 0.501 | FALSE |
| RPF2 | 6906.860 | 16830.365 | 1.057 | 0.929 | FALSE |
| MRPS2 | 4569.227 | 10084.017 | 0.975 | 1.111 | FALSE |
| PFDN2 | 12081.859 | 20975.454 | 0.969 | 1.619 | FALSE |
| KPRP | 33971.565 | 39713.228 | 0.969 | 0.533 | FALSE |
| MRPS9 | 7958.585 | 17454.814 | 0.965 | 1.350 | FALSE |
| MRPS23 | 18438.468 | 40478.727 | 0.959 | 1.037 | FALSE |
| ERGIC2 | 5546.275 | 6997.554 | 0.927 | 0.964 | FALSE |
| H2AZ2 | 315783.111 | 306375.448 | 0.927 | 0.300 | FALSE |

| | | | | | |
|---|---|---|---|---|---|
| STX3 | 3540.580 | 3781.703 | 0.893 | 0.310 | FALSE |
| MRPS28 | 15012.256 | 31132.718 | 0.878 | 1.132 | FALSE |
| PDIA3 | 5513.305 | 9874.159 | 0.877 | 0.697 | FALSE |
| MRPS34 | 8519.682 | 20214.063 | 0.871 | 1.444 | FALSE |
| MRPS27 | 6386.110 | 12385.688 | 0.860 | 1.385 | FALSE |
| PLS3 | 2875.815 | 2761.662 | 0.818 | 0.276 | FALSE |
| TOP2A | 6471.237 | 10149.359 | 0.797 | 0.842 | FALSE |
| SNRPD3 | 184752.924 | 192261.066 | 0.791 | 0.841 | FALSE |
| RHOA | 32141.084 | 38732.390 | 0.787 | 0.504 | FALSE |
| ALB | 16712.569 | 35862.262 | 0.777 | 0.355 | FALSE |
| PSMA5 | 12096.162 | 15721.608 | 0.768 | 0.549 | FALSE |
| DAP3 | 14967.021 | 26310.012 | 0.758 | 0.936 | FALSE |
| RBM6 | 134715.831 | 170514.465 | 0.756 | 1.327 | FALSE |
| CLINT1 | 7398.793 | 10168.424 | 0.744 | 0.362 | FALSE |
| PRPSAP1 | 129822.458 | 223523.921 | 0.717 | 0.362 | FALSE |
| IVNS1ABP | 274817.682 | 516838.557 | 0.714 | 0.473 | FALSE |
| PSMB5 | 3910.942 | 5270.382 | 0.704 | 0.902 | FALSE |
| FLNA | 350204.265 | 620846.852 | 0.702 | 0.881 | FALSE |
| PRPS2 | 154024.592 | 279622.353 | 0.701 | 0.359 | FALSE |
| MT1E | 65120.300 | 60800.871 | 0.685 | 0.519 | FALSE |
| SAR1A | 7508.386 | 10883.270 | 0.682 | 0.694 | FALSE |
| MRPS7 | 18371.889 | 32496.269 | 0.671 | 0.895 | FALSE |
| HDX | 27491.774 | 76538.974 | 0.661 | 0.638 | FALSE |
| PFDN5 | 5274.346 | 8741.933 | 0.650 | 1.058 | FALSE |
| NACA | 487.159 | 676.412 | 0.644 | 1.254 | FALSE |
| TUFM | 4533.863 | 6162.917 | 0.644 | 1.168 | FALSE |
| MRPS21 | 6890.800 | 19195.493 | 0.643 | 0.758 | FALSE |
| HACD3 | 7770.383 | 11499.519 | 0.639 | 0.544 | FALSE |
| HDAC1 | 41931.787 | 58236.090 | 0.618 | 0.809 | FALSE |
| SERPINH1 | 4858.618 | 7472.390 | 0.604 | 1.486 | FALSE |
| PSMA3 | 7906.937 | 10276.407 | 0.594 | 0.699 | FALSE |
| NAP1L1 | 59734.363 | 91580.613 | 0.589 | 1.349 | FALSE |
| MRPL19 | 1305.241 | 1504.651 | 0.589 | 0.396 | FALSE |
| DCD | 323557.732 | 348884.113 | 0.582 | 0.288 | FALSE |
| SPIN4 | 66437.975 | 75665.329 | 0.573 | 1.134 | FALSE |
| GPI | 4195.693 | 3729.468 | 0.573 | 0.181 | FALSE |
| LTBP1 | 4247.059 | 6275.347 | 0.572 | 0.548 | FALSE |
| TUBA3C | 1811343.797 | 2567649.536 | 0.564 | 0.666 | FALSE |
| DDX21 | 6316.696 | 11714.338 | 0.557 | 1.656 | FALSE |
| DNAJB6 | 12815.340 | 15270.645 | 0.547 | 0.478 | FALSE |
| EIF2B4 | 4218.025 | 5893.067 | 0.545 | 0.627 | FALSE |
| YBX3 | 157723.006 | 203679.164 | 0.544 | 1.211 | FALSE |
| CORO1C | 56496.100 | 83504.101 | 0.540 | 0.498 | FALSE |

| | | | | | |
|---|---|---|---|---|---|
| P4HB | 24609.928 | 36295.968 | 0.537 | 1.358 | FALSE |
| AFTPH | 5637.043 | 7951.258 | 0.537 | 0.359 | FALSE |
| ACAP2 | 61208.761 | 367463.608 | 0.536 | 0.627 | FALSE |
| TUBB3 | 1729922.030 | 2264203.439 | 0.536 | 0.436 | FALSE |
| EPB41L2 | 5628.558 | 7087.403 | 0.529 | 2.105 | FALSE |
| PLEC | 104560.694 | 155870.278 | 0.528 | 2.096 | FALSE |
| RCN2 | 63042.772 | 112813.420 | 0.518 | 0.533 | FALSE |
| SMARCB1 | 3562.293 | 5948.527 | 0.517 | 0.550 | FALSE |
| PRPS1 | 220161.333 | 355400.469 | 0.511 | 0.252 | FALSE |
| TUBB6 | 818769.000 | 1142818.678 | 0.511 | 0.872 | FALSE |
| TACC3 | 10150.742 | 11901.264 | 0.503 | 0.616 | FALSE |
| S100A14 | 10413.176 | 11565.696 | 0.502 | 0.308 | FALSE |
| TUBA1C | 2516807.710 | 3666756.986 | 0.501 | 0.438 | FALSE |
| TUBA4B | 90999.045 | 182128.585 | 0.499 | 0.330 | FALSE |
| RANBP10 | 281310.986 | 230904.778 | 0.499 | 0.557 | FALSE |
| SET | 54478.609 | 41815.537 | 0.499 | 0.506 | FALSE |
| PTS | 130079.721 | 168849.324 | 0.498 | 1.014 | FALSE |
| HDAC2 | 58385.121 | 74232.512 | 0.497 | 1.504 | FALSE |
| MISP | 5009.370 | 4586.602 | 0.493 | 0.239 | FALSE |
| HSP90B1 | 24302.443 | 30876.419 | 0.489 | 0.605 | FALSE |
| CTPS1 | 20411.340 | 29253.574 | 0.489 | 0.719 | FALSE |
| P4HA2 | 2384.393 | 3141.297 | 0.486 | 1.125 | FALSE |
| HNRNPM | 114529.112 | 161365.720 | 0.476 | 0.403 | FALSE |
| KTN1 | 13862.498 | 21163.539 | 0.461 | 0.877 | FALSE |
| PPIA | 17172.489 | 16949.372 | 0.460 | 0.477 | FALSE |
| YPEL5 | 31607.437 | 37713.692 | 0.456 | 0.815 | FALSE |
| GSDMA | 13616.416 | 12012.999 | 0.455 | 0.316 | FALSE |
| CEP43 | 8360.955 | 12112.798 | 0.454 | 0.667 | FALSE |
| TARDBP | 10389.885 | 13643.734 | 0.452 | 0.309 | FALSE |
| PSMB3 | 10453.900 | 11245.078 | 0.451 | 0.337 | FALSE |
| E2F7 | 42131.297 | 55372.650 | 0.445 | 1.157 | FALSE |
| SPIN1 | 1097790.817 | 1355242.375 | 0.445 | 0.722 | FALSE |
| AKAP2 | 2807.944 | 3034.268 | 0.444 | 0.399 | FALSE |
| SUCLG2 | 1134.766 | 2588.127 | 0.444 | 0.500 | FALSE |
| MTHFD1L | 3513.366 | 4357.072 | 0.443 | 1.013 | FALSE |
| MAT2A | 87796.854 | 105571.266 | 0.443 | 0.875 | FALSE |
| MKLN1 | 6024.494 | 11000.747 | 0.440 | 0.360 | FALSE |
| MAP4 | 2099.809 | 2522.008 | 0.438 | 0.716 | FALSE |
| CIT | 242.417 | 502.031 | 0.429 | 0.765 | FALSE |
| RANBP9 | 23835.603 | 33712.783 | 0.426 | 0.577 | FALSE |
| PPP1CA | 20710.413 | 22410.867 | 0.424 | 0.763 | FALSE |
| PRKAR2A | 12428.074 | 15101.252 | 0.424 | 0.582 | FALSE |
| RBBP4 | 86247.625 | 107132.157 | 0.420 | 1.077 | FALSE |

| | | | | | |
|---|---|---|---|---|---|
| PCNA | 40626.490 | 50215.606 | 0.420 | 0.941 | FALSE |
| PPHLN1 | 7131.251 | 8286.206 | 0.418 | 0.498 | FALSE |
| KRT8 | 1283427.542 | 1228890.771 | 0.418 | 1.404 | FALSE |
| PPOX | 1874.903 | 3400.291 | 0.411 | 0.461 | FALSE |
| MAPK6 | 3081.791 | 3940.586 | 0.408 | 0.414 | FALSE |
| AHNAK | 10756.084 | 14428.745 | 0.408 | 1.155 | FALSE |
| ALYREF | 6365.276 | 6562.241 | 0.402 | 0.321 | FALSE |
| U2AF2 | 16125.958 | 23878.234 | 0.398 | 0.205 | FALSE |
| L2HGDH | 11970.648 | 18110.324 | 0.397 | 0.988 | FALSE |
| ALDOA | 4811.323 | 5302.218 | 0.394 | 0.464 | FALSE |
| HNRNPH1 | 1024645.333 | 1160294.038 | 0.394 | 0.908 | FALSE |
| DHX38 | 24668.370 | 33824.853 | 0.393 | 1.514 | FALSE |
| WDR5 | 20841.865 | 22473.416 | 0.391 | 0.527 | FALSE |
| KYNU | 4083.117 | 3124.573 | 0.388 | 0.170 | FALSE |
| CBX1 | 36000.243 | 43094.547 | 0.387 | 0.570 | FALSE |
| P4HA1 | 3939.018 | 4807.790 | 0.386 | 0.666 | FALSE |
| PRPSAP2 | 71780.824 | 104530.579 | 0.385 | 0.173 | FALSE |
| LDHB | 44043.930 | 46981.868 | 0.379 | 0.443 | FALSE |
| WDR1 | 3235.366 | 3282.635 | 0.378 | 0.275 | FALSE |
| SEC13 | 4737.775 | 9541.068 | 0.371 | 0.207 | FALSE |
| MAGED1 | 9347.208 | 10581.066 | 0.370 | 0.465 | FALSE |
| PAICS | 2902.550 | 3639.507 | 0.367 | 0.862 | FALSE |
| RPL23 | 81525.748 | 125865.458 | 0.364 | 0.304 | FALSE |
| RFC5 | 3855.422 | 5401.973 | 0.363 | 0.750 | FALSE |
| NT5C2 | 26895.680 | 30841.337 | 0.362 | 0.508 | FALSE |
| ACTC1 | 5643734.957 | 3921114.870 | 0.362 | 0.187 | FALSE |
| TUBB2A | 1891885.273 | 2409138.856 | 0.360 | 1.191 | FALSE |
| TRIM28 | 38483.999 | 48615.004 | 0.358 | 0.824 | FALSE |
| SLC25A6 | 194287.494 | 181919.340 | 0.354 | 0.156 | FALSE |
| SART1 | 40199.602 | 46236.645 | 0.353 | 0.991 | FALSE |
| TNRC6B | 37530.569 | 44480.065 | 0.351 | 0.677 | FALSE |
| TPM1 | 10805.227 | 10640.582 | 0.348 | 0.393 | FALSE |
| CTSD | 19907.253 | 23321.338 | 0.345 | 0.249 | FALSE |
| SPINDOC | 712024.924 | 954000.186 | 0.344 | 0.591 | FALSE |
| ARL8B | 10857.216 | 12563.441 | 0.341 | 0.220 | FALSE |
| LPCAT1 | 3513.670 | 4728.456 | 0.340 | 0.825 | FALSE |
| RBM17 | 19478.930 | 22036.624 | 0.330 | 0.289 | FALSE |
| GAPDH | 209080.724 | 200122.250 | 0.329 | 0.250 | FALSE |
| CEP350 | 1674.421 | 1676.401 | 0.326 | 0.369 | FALSE |
| TPD52L2 | 25358.184 | 45915.910 | 0.326 | 1.105 | FALSE |
| MGA | 49499.444 | 59065.384 | 0.324 | 0.717 | FALSE |
| PCBP1 | 23027.146 | 27413.627 | 0.324 | 1.139 | FALSE |
| PPM1A | 44924.167 | 54744.050 | 0.318 | 0.803 | FALSE |

| | | | | | |
|---|---|---|---|---|---|
| KARS1 | 18422.819 | 23658.032 | 0.318 | 1.280 | FALSE |
| TPM3 | 14903.433 | 14099.137 | 0.316 | 0.387 | FALSE |
| SPTLC2 | 1119.828 | 2746.169 | 0.315 | 0.381 | FALSE |
| PSMA7 | 12832.426 | 14051.901 | 0.315 | 0.595 | FALSE |
| KRT2 | 5943467.783 | 5890286.078 | 0.314 | 0.230 | FALSE |
| RPS27A | 319360.091 | 384932.778 | 0.311 | 1.455 | FALSE |
| DPF2 | 3278.559 | 4003.251 | 0.309 | 0.570 | FALSE |
| PDIA6 | 4786.921 | 6679.400 | 0.309 | 0.326 | FALSE |
| TUBB2B | 1899124.424 | 2420075.871 | 0.308 | 1.406 | FALSE |
| MRPL22 | 2695.690 | 3299.347 | 0.308 | 0.604 | FALSE |
| PSMA6 | 12110.894 | 13841.008 | 0.307 | 0.375 | FALSE |
| SNAPIN | 5253.103 | 5512.163 | 0.305 | 0.276 | FALSE |
| FLG2 | 16078.376 | 18419.646 | 0.303 | 0.206 | FALSE |
| EIF4B | ########### | 11850597.481 | 0.303 | 0.431 | FALSE |
| PSMB6 | 18081.132 | 19020.042 | 0.301 | 0.226 | FALSE |
| PTBP1 | 2239.934 | 1515.706 | 0.297 | 0.125 | FALSE |
| PSMC6 | 82848.082 | 109656.810 | 0.293 | 0.692 | FALSE |
| DDX5 | 7853.729 | 9361.668 | 0.292 | 0.594 | FALSE |
| PRDX6 | 43759.116 | 52251.724 | 0.291 | 0.416 | FALSE |
| CLU | 14071.236 | 19520.299 | 0.290 | 0.431 | FALSE |
| PSMC3 | 82864.258 | 112333.832 | 0.289 | 0.611 | FALSE |
| YBX1 | 188378.339 | 242212.393 | 0.286 | 0.635 | FALSE |
| VBP1 | 2489.572 | 4958.825 | 0.284 | 0.228 | FALSE |
| PFN1 | 19759.734 | 18544.350 | 0.283 | 0.220 | FALSE |
| ZBTB24 | 10226.978 | 11962.208 | 0.283 | 1.436 | FALSE |
| PSAP | 27670.204 | 26118.488 | 0.281 | 0.129 | FALSE |
| RMND5A | 2098.676 | 2626.887 | 0.281 | 0.267 | FALSE |
| WDR26 | 30530.041 | 38656.262 | 0.281 | 0.266 | FALSE |
| CTTN | 15002.385 | 16829.436 | 0.280 | 0.269 | FALSE |
| GLUD1 | 6431.845 | 7275.529 | 0.278 | 0.400 | FALSE |
| PSMD4 | 44392.435 | 58998.391 | 0.277 | 1.336 | FALSE |
| PA2G4 | 4396.915 | 2534.204 | 0.275 | 0.103 | FALSE |
| TAB2 | 72142.877 | 81772.957 | 0.275 | 0.507 | FALSE |
| Trypsin | ########### | ########### | 0.267 | 0.239 | FALSE |
| BRI3BP | 6399.899 | 7565.508 | 0.265 | 0.291 | FALSE |
| SPIN2B | 134713.717 | 195877.023 | 0.263 | 0.477 | FALSE |
| MCAM | 1877.228 | 4488.899 | 0.262 | 0.120 | FALSE |
| LDHAL6B | 13582.734 | 9601.789 | 0.262 | 0.118 | FALSE |
| HSPE1 | 10797.451 | 8462.812 | 0.262 | 0.127 | FALSE |
| E2F6 | 11331.496 | 12058.593 | 0.261 | 0.650 | FALSE |
| HTATSF1 | 3356.925 | 3844.859 | 0.258 | 0.472 | FALSE |
| RPL24 | 18248.823 | 26030.570 | 0.257 | 0.138 | FALSE |
| RBM5 | 690843.351 | 639806.895 | 0.251 | 0.433 | FALSE |

| | | | | | |
|---|---|---|---|---|---|
| HSPA1A | 466695.926 | 526088.481 | 0.250 | 0.597 | FALSE |
| KRT18 | 913120.511 | 887146.689 | 0.245 | 0.420 | FALSE |
| RAB5A | 53622.329 | 44897.568 | 0.240 | 0.205 | FALSE |
| MAX | 11186.658 | 11896.201 | 0.240 | 0.282 | FALSE |
| HNRNPK | 502923.885 | 575969.622 | 0.239 | 0.395 | FALSE |
| RING1 | 3969.768 | 4420.414 | 0.237 | 0.517 | FALSE |
| ERC1 | 2925.068 | 3578.977 | 0.232 | 0.954 | FALSE |
| PSMD3 | 40101.214 | 51670.713 | 0.230 | 1.118 | FALSE |
| ERAL1 | 2364.377 | 5226.784 | 0.229 | 0.144 | FALSE |
| HSP90AB1 | 176228.504 | 186241.625 | 0.228 | 0.859 | FALSE |
| ARFGAP1 | 18346.402 | 18540.313 | 0.227 | 0.321 | FALSE |
| HSPD1 | 382452.162 | 438868.129 | 0.227 | 1.506 | FALSE |
| MYL1 | 7013.404 | 8617.849 | 0.227 | 0.234 | FALSE |
| AZGP1 | 25074.028 | 32143.177 | 0.226 | 0.212 | FALSE |
| MAP3K7 | 173656.616 | 162836.737 | 0.226 | 0.386 | FALSE |
| HNRNPF | 234545.623 | 289802.670 | 0.225 | 0.746 | FALSE |
| U2AF1 | 33859.392 | 42247.509 | 0.224 | 0.170 | FALSE |
| PSMD5 | 3551.468 | 8815.889 | 0.224 | 0.126 | FALSE |
| TAB1 | 416770.453 | 493939.433 | 0.223 | 0.363 | FALSE |
| ODR4 | 6739.443 | 8217.462 | 0.220 | 0.885 | FALSE |
| PSMC5 | 82176.854 | 97452.620 | 0.218 | 1.349 | FALSE |
| AP1M1 | 2106.111 | 1985.398 | 0.218 | 0.309 | FALSE |
| SNRNP40 | 9832.620 | 10791.631 | 0.218 | 0.490 | FALSE |
| OSTC | 3957.821 | 6950.153 | 0.218 | 0.287 | FALSE |
| ARGLU1 | 145915.431 | 132864.656 | 0.217 | 0.110 | FALSE |
| AGO2 | 23728.719 | 28557.771 | 0.213 | 0.363 | FALSE |
| EZR | 3615.666 | 3759.493 | 0.213 | 0.599 | FALSE |
| BRCA2 | 400.900 | 443.958 | 0.212 | 0.463 | FALSE |
| HSPA5 | 386822.149 | 447275.371 | 0.210 | 0.726 | FALSE |
| MATR3 | 75837.889 | 91974.198 | 0.209 | 0.158 | FALSE |
| ALG13 | 4128.323 | 4600.446 | 0.206 | 0.178 | FALSE |
| PKM | 33584.031 | 29532.402 | 0.204 | 0.230 | FALSE |
| MSH6 | 15009.967 | 17850.193 | 0.204 | 0.441 | FALSE |
| DNAJA3 | 7204.186 | 8304.352 | 0.204 | 0.604 | FALSE |
| VDAC3 | 16551.108 | 19057.668 | 0.204 | 0.129 | FALSE |
| SNRPC | 79473.498 | 72678.639 | 0.204 | 0.105 | FALSE |
| DYNLL2 | 26539.728 | 35631.556 | 0.204 | 0.618 | FALSE |
| SUPT16H | 5649.896 | 6735.863 | 0.204 | 0.447 | FALSE |
| SYNRG | 24647.751 | 28843.824 | 0.201 | 0.516 | FALSE |
| APOD | 15648.424 | 16983.465 | 0.201 | 0.115 | FALSE |
| ARHGAP21 | 4235.105 | 4449.843 | 0.200 | 0.303 | FALSE |
| PCGF6 | 42481.029 | 45262.189 | 0.200 | 0.323 | FALSE |
| FDFT1 | 20052.267 | 26324.607 | 0.200 | 0.160 | FALSE |

| | | | | | |
|---|---|---|---|---|---|
| LGALS7 | 33595.687 | 40176.079 | 0.200 | 0.195 | FALSE |
| FLNB | 37832.300 | 58580.181 | 0.199 | 0.251 | FALSE |
| KRT17 | 1948534.078 | 1747188.725 | 0.198 | 0.349 | FALSE |
| NNMT | 17813.218 | 19157.589 | 0.193 | 0.143 | FALSE |
| VIM | 439270.933 | 472917.190 | 0.191 | 0.552 | FALSE |
| RCL1 | 22027.701 | 25078.077 | 0.189 | 0.281 | FALSE |
| EIF2B3 | 3004.243 | 3225.315 | 0.189 | 0.238 | FALSE |
| PSMB1 | 34969.810 | 34123.752 | 0.186 | 0.242 | FALSE |
| ARAF | 1310.470 | 1349.388 | 0.184 | 0.304 | FALSE |
| FABP5 | 35225.739 | 35945.662 | 0.183 | 0.134 | FALSE |
| CSNK2B | 6720.568 | 7207.797 | 0.183 | 0.232 | FALSE |
| MBD2 | 1516.453 | 2328.714 | 0.181 | 0.226 | FALSE |
| DHX9 | 18729.456 | 23577.866 | 0.179 | 0.761 | FALSE |
| PSMC2 | 69446.484 | 86511.986 | 0.179 | 0.389 | FALSE |
| RBBP7 | 33497.147 | 40342.578 | 0.179 | 0.291 | FALSE |
| EIF3D | 21868.779 | 23965.028 | 0.178 | 0.544 | FALSE |
| ERG28 | 6400.094 | 8376.264 | 0.178 | 0.130 | FALSE |
| ECM1 | 895.853 | 977.596 | 0.174 | 0.147 | FALSE |
| G3BP1 | 10518.532 | 11900.114 | 0.173 | 0.466 | FALSE |
| PSMD2 | 42408.734 | 54083.628 | 0.172 | 0.305 | FALSE |
| EIF2B2 | 13189.575 | 16993.606 | 0.171 | 0.125 | FALSE |
| EIF4A1 | 35500.002 | 23941.568 | 0.171 | 0.179 | FALSE |
| EIF3H | 42243.195 | 50048.169 | 0.170 | 0.557 | FALSE |
| PAF1 | 6776.478 | 6839.850 | 0.169 | 0.319 | FALSE |
| KNSTRN | 4586.382 | 4143.074 | 0.169 | 0.141 | FALSE |
| NCL | 7658.038 | 6073.505 | 0.169 | 0.088 | FALSE |
| S100A16 | 232825.507 | 213725.382 | 0.167 | 0.104 | FALSE |
| HSPA6 | 334723.393 | 362858.784 | 0.166 | 0.653 | FALSE |
| OTUD4 | 353367.253 | 344305.856 | 0.165 | 0.246 | FALSE |
| DLST | 3063.139 | 3355.951 | 0.164 | 0.162 | FALSE |
| PRPF31 | 1572647.639 | 1721573.847 | 0.164 | 0.200 | FALSE |
| SMARCE1 | 8286.307 | 8516.847 | 0.164 | 0.391 | FALSE |
| MAGED2 | 47983.150 | 57557.013 | 0.164 | 0.513 | FALSE |
| REST | 15337.238 | 16524.942 | 0.164 | 0.223 | FALSE |
| BMS1 | 28217.977 | 27937.832 | 0.163 | 0.247 | FALSE |
| LDHA | 53652.925 | 50232.928 | 0.162 | 0.138 | FALSE |
| KRT80 | 42492.794 | 48757.903 | 0.162 | 0.117 | FALSE |
| TWF1 | 2445.137 | 2714.512 | 0.161 | 0.298 | FALSE |
| MAEA | 11735.919 | 12701.657 | 0.158 | 0.274 | FALSE |
| AGO3 | 10604.275 | 11758.754 | 0.155 | 0.272 | FALSE |
| BLOC1S2 | 46585.827 | 47241.599 | 0.155 | 0.197 | FALSE |
| KPNA2 | 35007.991 | 42314.324 | 0.155 | 0.246 | FALSE |
| CDSN | 24128.837 | 25660.681 | 0.154 | 0.091 | FALSE |

| | | | | | |
|---|---|---|---|---|---|
| MTREX | 1171.475 | 1703.324 | 0.153 | 0.123 | FALSE |
| HSPH1 | 22704.613 | 25211.027 | 0.151 | 0.463 | FALSE |
| PSMD8 | 27224.546 | 30085.172 | 0.150 | 1.320 | FALSE |
| TRO | 7099.162 | 7482.310 | 0.149 | 0.374 | FALSE |
| CCT5 | 424796.906 | 569682.534 | 0.149 | 0.122 | FALSE |
| ACTL6A | 8497.313 | 8120.148 | 0.149 | 0.293 | FALSE |
| EIF3F | 91961.589 | 95724.582 | 0.148 | 0.510 | FALSE |
| NONO | 20204.176 | 23043.615 | 0.148 | 0.435 | FALSE |
| GEMIN5 | 1455.954 | 1528.271 | 0.147 | 0.246 | FALSE |
| EIF3A | 41223.815 | 45527.861 | 0.146 | 0.211 | FALSE |
| PPP1CC | 27437.326 | 29177.189 | 0.146 | 0.249 | FALSE |
| DHRS2 | 177633.042 | 174556.542 | 0.145 | 0.222 | FALSE |
| DNAJB2 | 36731.674 | 39513.010 | 0.144 | 0.232 | FALSE |
| TIMM44 | 8157.807 | 9347.645 | 0.144 | 0.556 | FALSE |
| MAP1B | 117001.238 | 128151.688 | 0.144 | 0.333 | FALSE |
| DDX3X | 24216.166 | 26501.256 | 0.143 | 0.316 | FALSE |
| CHD4 | 12028.839 | 14025.119 | 0.141 | 0.288 | FALSE |
| GTF3C5 | 1502.682 | 2698.182 | 0.140 | 0.108 | FALSE |
| GPRC5A | 20677.876 | 15176.504 | 0.138 | 0.080 | FALSE |
| PSMC1 | 92660.910 | 103805.927 | 0.136 | 0.331 | FALSE |
| RAB6A | 74880.609 | 60685.308 | 0.135 | 0.049 | FALSE |
| ROCK1 | 46193.951 | 50963.220 | 0.135 | 0.213 | FALSE |
| CFL1 | 111847.208 | 101110.460 | 0.135 | 0.136 | FALSE |
| EPRS1 | 7903.587 | 9239.913 | 0.131 | 0.358 | FALSE |
| RHOB | 10727.331 | 15833.235 | 0.131 | 0.055 | FALSE |
| CNOT1 | 1635.518 | 1669.839 | 0.128 | 0.534 | FALSE |
| SNRPB2 | 126495.009 | 114031.588 | 0.127 | 0.164 | FALSE |
| GLG1 | 13874.622 | 16505.586 | 0.126 | 0.090 | FALSE |
| MCM7 | 42022.337 | 50305.668 | 0.125 | 0.157 | FALSE |
| HSDL1 | 8108.494 | 7257.749 | 0.125 | 0.154 | FALSE |
| BAIAP2 | 1812.261 | 1859.250 | 0.125 | 0.147 | FALSE |
| IQGAP1 | 1039.288 | 895.996 | 0.123 | 0.099 | FALSE |
| USP7 | 1459.345 | 1146.250 | 0.123 | 0.130 | FALSE |
| EIF3G | 55318.167 | 63799.325 | 0.123 | 0.603 | FALSE |
| MTA2 | 13995.298 | 15123.644 | 0.122 | 0.223 | FALSE |
| HLA-A | 5071.077 | 6870.866 | 0.120 | 0.065 | FALSE |
| TRIM32 | 3193.826 | 3492.882 | 0.119 | 0.191 | FALSE |
| HNRNPU | 58100.930 | 70742.170 | 0.119 | 0.168 | FALSE |
| LAS1L | 3947.015 | 4479.168 | 0.119 | 0.362 | FALSE |
| TET2 | 9051.591 | 9415.427 | 0.118 | 0.195 | FALSE |
| TUBB4A | 1474538.545 | 1888883.538 | 0.117 | 0.545 | FALSE |
| SKA1 | 90765.005 | 90184.631 | 0.117 | 0.281 | FALSE |
| PHF5A | 77361.023 | 81773.789 | 0.116 | 0.125 | FALSE |

| | | | | | |
|---|---|---|---|---|---|
| KRT10 | ########## | 11068132.381 | 0.116 | 0.091 | FALSE |
| HCFC1 | 1131.141 | 1101.213 | 0.113 | 0.218 | FALSE |
| CAVIN1 | 27708.239 | 31911.348 | 0.113 | 0.375 | FALSE |
| SSRP1 | 4192.189 | 4438.235 | 0.112 | 0.152 | FALSE |
| IGF2R | 992.363 | 1028.582 | 0.111 | 0.105 | FALSE |
| DDB1 | 20766.195 | 26842.658 | 0.111 | 0.219 | FALSE |
| NPM1 | 14093.440 | 15185.093 | 0.110 | 0.154 | FALSE |
| PCBP2 | 16366.347 | 17002.481 | 0.109 | 0.158 | FALSE |
| RPL10A | 27688.995 | 35172.613 | 0.107 | 0.040 | FALSE |
| DNAJA1 | 50996.195 | 66209.201 | 0.106 | 0.186 | FALSE |
| SNAP29 | 15157.329 | 13473.075 | 0.105 | 0.076 | FALSE |
| LGALS3 | 15932.394 | 17289.838 | 0.104 | 0.144 | FALSE |
| CBX3 | 58254.794 | 64573.639 | 0.103 | 0.243 | FALSE |
| SCFD1 | 3708.999 | 3960.290 | 0.099 | 0.232 | FALSE |
| HNRNPD | 7589.271 | 7776.944 | 0.099 | 0.086 | FALSE |
| SERPINB12 | 50661.141 | 46737.290 | 0.097 | 0.060 | FALSE |
| NUP93 | 18950.599 | 23531.129 | 0.095 | 0.105 | FALSE |
| KCTD17 | 89328.117 | 113319.486 | 0.095 | 0.145 | FALSE |
| TGM3 | 17360.484 | 17141.710 | 0.095 | 0.066 | FALSE |
| EXOSC6 | 4791.487 | 4797.378 | 0.094 | 0.185 | FALSE |
| GSTP1 | 21885.097 | 23094.925 | 0.094 | 0.179 | FALSE |
| PPP1CB | 25806.474 | 26351.405 | 0.090 | 0.177 | FALSE |
| COPG1 | 4849.241 | 5184.178 | 0.089 | 0.127 | FALSE |
| GOLT1B | 15798.566 | 16555.959 | 0.088 | 0.097 | FALSE |
| TUBB | 2654364.303 | 3313336.576 | 0.088 | 0.062 | FALSE |
| VDAC1 | 52106.900 | 55653.766 | 0.088 | 0.054 | FALSE |
| XRCC5 | 12937.372 | 14993.225 | 0.085 | 0.105 | FALSE |
| WDR77 | 9013548.889 | 9734761.611 | 0.085 | 0.128 | FALSE |
| L3MBTL2 | 9460.643 | 9587.822 | 0.083 | 0.151 | FALSE |
| SLC4A1AP | 9292.306 | 11365.968 | 0.082 | 0.133 | FALSE |
| DNAJA2 | 11375.531 | 16815.481 | 0.082 | 0.117 | FALSE |
| PPP6R1 | 2265.722 | 2288.127 | 0.081 | 0.237 | FALSE |
| FLII | 2249.020 | 3314.698 | 0.081 | 0.167 | FALSE |
| CALU | 19414.115 | 18356.060 | 0.076 | 0.254 | FALSE |
| EIF3I | 90231.640 | 97329.049 | 0.074 | 0.194 | FALSE |
| BLMH | 4702.646 | 4816.934 | 0.074 | 0.046 | FALSE |
| FIP1L1 | 3088.825 | 2682.293 | 0.074 | 0.051 | FALSE |
| SMARCC1 | 5434.994 | 5454.533 | 0.074 | 0.117 | FALSE |
| SF3B2 | 61254.867 | 63834.044 | 0.072 | 0.092 | FALSE |
| RBM4 | 7418.780 | 7737.782 | 0.069 | 0.069 | FALSE |
| CAMSAP3 | 30876.708 | 32494.918 | 0.069 | 0.145 | FALSE |
| ZRANB2 | 14265.473 | 14702.919 | 0.067 | 0.048 | FALSE |
| FASN | 8303.762 | 8747.547 | 0.067 | 0.098 | FALSE |

| | | | | | |
|---|---|---|---|---|---|
| RBM10 | 4351179.234 | 4571332.766 | 0.067 | 0.105 | FALSE |
| VDAC2 | 44704.441 | 44959.475 | 0.066 | 0.034 | FALSE |
| CD2BP2 | 5536.984 | 5453.514 | 0.066 | 0.139 | FALSE |
| SBSN | 2397.327 | 2085.958 | 0.065 | 0.031 | FALSE |
| DSG1 | 53269.887 | 50581.185 | 0.065 | 0.038 | FALSE |
| PSMD14 | 68105.308 | 77017.419 | 0.065 | 0.279 | FALSE |
| POLR2B | 2306.367 | 2432.229 | 0.063 | 0.109 | FALSE |
| SPAG5 | 3774.508 | 3614.339 | 0.062 | 0.056 | FALSE |
| SFPQ | 12964.560 | 12396.340 | 0.061 | 0.138 | FALSE |
| SF3B4 | 61126.871 | 69449.948 | 0.060 | 0.096 | FALSE |
| SLAIN2 | 299722.435 | 296144.738 | 0.060 | 0.163 | FALSE |
| TOP2B | 4678.836 | 6277.067 | 0.060 | 0.071 | FALSE |
| EEF1E1 | 13365.130 | 12792.617 | 0.059 | 0.045 | FALSE |
| ABCB7 | 1067.493 | 2525.707 | 0.059 | 0.059 | FALSE |
| SF3A1 | 19583.440 | 20058.976 | 0.057 | 0.059 | FALSE |
| SF3B1 | 101804.269 | 112084.273 | 0.057 | 0.072 | FALSE |
| RPL38 | 79271.634 | 95538.672 | 0.057 | 0.060 | FALSE |
| ANKRD28 | 6252.455 | 7221.806 | 0.057 | 0.074 | FALSE |
| VAPA | 30782.990 | 32157.134 | 0.057 | 0.058 | FALSE |
| CDK1 | 42893.101 | 43129.297 | 0.056 | 0.051 | FALSE |
| FOXP4 | 32404.286 | 32278.463 | 0.054 | 0.057 | FALSE |
| SPCS2 | 3073.209 | 2954.742 | 0.054 | 0.054 | FALSE |
| AGL | 3935.759 | 4446.356 | 0.053 | 0.057 | FALSE |
| RUVBL1 | 158554.694 | 164207.078 | 0.053 | 0.104 | FALSE |
| ARMC8 | 35342.532 | 37267.559 | 0.052 | 0.080 | FALSE |
| MANSC1 | 6882.944 | 7673.566 | 0.051 | 0.096 | FALSE |
| EIF5A | 33678.615 | 11559.853 | 0.050 | 0.015 | FALSE |
| GID8 | 105926.293 | 134625.883 | 0.050 | 0.065 | FALSE |
| HSP90AA1 | 125701.524 | 125991.849 | 0.050 | 0.096 | FALSE |
| RPLP1 | 13666.435 | 46367.068 | 0.049 | 0.018 | FALSE |
| FLNC | 191698.976 | 216670.091 | 0.048 | 0.095 | FALSE |
| SFN | 206828.236 | 119119.163 | 0.047 | 0.029 | FALSE |
| HSPA4 | 19097.825 | 20224.180 | 0.047 | 0.365 | FALSE |
| TGM1 | 12074.124 | 6376.780 | 0.046 | 0.026 | FALSE |
| CSNK2A2 | 7524.805 | 8675.370 | 0.045 | 0.034 | FALSE |
| RFC2 | 5213.219 | 5674.395 | 0.045 | 0.083 | FALSE |
| TAB3 | 58874.175 | 59670.066 | 0.044 | 0.056 | FALSE |
| EEF1A2 | 321422.687 | 314734.720 | 0.043 | 0.043 | FALSE |
| CAPZB | 81192.294 | 76250.548 | 0.038 | 0.068 | FALSE |
| TAF9B | 11656.552 | 10771.654 | 0.038 | 0.032 | FALSE |
| LMNA | 106463.625 | 109683.462 | 0.036 | 0.067 | FALSE |
| SKP1 | 14629.738 | 14782.814 | 0.036 | 0.060 | FALSE |
| PSMD7 | 45920.337 | 53532.250 | 0.035 | 0.071 | FALSE |

| | | | | | |
|---|---|---|---|---|---|
| GATAD2B | 6170.570 | 6228.806 | 0.035 | 0.046 | FALSE |
| KPNA3 | 7495.777 | 8509.163 | 0.034 | 0.058 | FALSE |
| SRRT | 36679.254 | 35648.893 | 0.033 | 0.053 | FALSE |
| STK38 | 2266660.000 | 2321935.864 | 0.031 | 0.036 | FALSE |
| CSNK2A1 | 15200.889 | 14389.261 | 0.031 | 0.023 | FALSE |
| SPIN3 | 476951.439 | 588890.816 | 0.031 | 0.033 | FALSE |
| EEF1A1 | 385624.613 | 378572.468 | 0.030 | 0.032 | FALSE |
| PYCR1 | 9112.295 | 9101.621 | 0.029 | 0.051 | FALSE |
| SMARCD2 | 3043.965 | 3105.722 | 0.028 | 0.099 | FALSE |
| CDK9 | 21916.737 | 21525.774 | 0.027 | 0.048 | FALSE |
| KRT14 | 3241382.196 | 2915777.020 | 0.026 | 0.022 | FALSE |
| AIFM1 | 3068.302 | 3147.033 | 0.024 | 0.026 | FALSE |
| NCCRP1 | 8104.020 | 6070.532 | 0.024 | 0.011 | FALSE |
| KRT74 | 301153.838 | 313349.238 | 0.024 | 0.013 | FALSE |
| PON2 | 6514.415 | 6430.561 | 0.023 | 0.019 | FALSE |
| TCP1 | 183441.314 | 196884.615 | 0.023 | 0.048 | FALSE |
| SNU13 | 8213.058 | 8303.660 | 0.022 | 0.020 | FALSE |
| NCAPH | 5229.783 | 5715.039 | 0.021 | 0.047 | FALSE |
| MT-CO2 | 9621.469 | 7971.993 | 0.021 | 0.013 | FALSE |
| BZW1 | 32015.430 | 36064.541 | 0.020 | 0.040 | FALSE |
| PSMD12 | 48605.402 | 56929.791 | 0.019 | 0.036 | FALSE |
| ZDBF2 | 6140.333 | 6754.557 | 0.018 | 0.032 | FALSE |
| KRT5 | 2638193.295 | 2437926.933 | 0.016 | 0.010 | FALSE |
| PLIN3 | 71690.868 | 76804.242 | 0.014 | 0.020 | FALSE |
| POF1B | 3366.835 | 2793.508 | 0.012 | 0.006 | FALSE |
| SEPTIN7 | 2931.425 | 2472.344 | 0.012 | 0.010 | FALSE |
| PPM1B | 87617.726 | 88519.139 | 0.009 | 0.011 | FALSE |
| EIF3B | 91579.959 | 96893.811 | 0.008 | 0.011 | FALSE |
| PSMD11 | 71986.387 | 81256.648 | 0.008 | 0.016 | FALSE |
| H4C1 | 720757.917 | 673996.986 | 0.006 | 0.009 | FALSE |
| PSME2 | 93295.370 | 109410.142 | 0.006 | 0.004 | FALSE |
| SF3B3 | 113692.958 | 120223.133 | 0.006 | 0.006 | FALSE |
| WARS1 | 22918.291 | 22451.810 | 0.005 | 0.010 | FALSE |
| MARS1 | 5657.539 | 6878.481 | 0.004 | 0.006 | FALSE |
| ATP6V1A | 2297.885 | 2367.957 | 0.004 | 0.002 | FALSE |
| NOP56 | 4332.480 | 4136.119 | 0.004 | 0.005 | FALSE |
| PRDX1 | 263571.583 | 251854.047 | -0.001 | 0.002 | FALSE |
| TIMM23B | 6181.101 | 12075.653 | -0.003 | 0.003 | FALSE |
| TPR | 3239.494 | 3174.418 | -0.003 | 0.006 | FALSE |
| RB1CC1 | 1141.451 | 1374.567 | -0.005 | 0.009 | FALSE |
| NCAPD2 | 6197.065 | 6163.322 | -0.006 | 0.009 | FALSE |
| RAD50 | 3195.399 | 3439.150 | -0.007 | 0.011 | FALSE |
| MCM3 | 3744.523 | 3917.526 | -0.008 | 0.025 | FALSE |

| | | | | | |
|---|---|---|---|---|---|
| APOL2 | 12139.653 | 10997.416 | -0.009 | 0.015 | FALSE |
| PSME1 | 86434.804 | 90167.009 | -0.010 | 0.008 | FALSE |
| SERPINB3 | 28125.971 | 26245.230 | -0.012 | 0.011 | FALSE |
| KRT77 | 1976186.745 | 1722945.059 | -0.013 | 0.011 | FALSE |
| NSDHL | 16558.232 | 16388.345 | -0.013 | 0.011 | FALSE |
| PFKM | 13904.257 | 13550.926 | -0.014 | 0.021 | FALSE |
| YTHDF2 | 3625.214 | 3288.290 | -0.014 | 0.020 | FALSE |
| RRBP1 | 6306.668 | 6689.240 | -0.015 | 0.030 | FALSE |
| XRCC6 | 13691.308 | 14486.809 | -0.016 | 0.030 | FALSE |
| RNF213 | 227.231 | 206.951 | -0.019 | 0.027 | FALSE |
| PSMC4 | 95567.788 | 108354.322 | -0.019 | 0.027 | FALSE |
| ZBED5 | 9449.570 | 8591.571 | -0.020 | 0.041 | FALSE |
| PSMD1 | 48853.854 | 49994.047 | -0.021 | 0.037 | FALSE |
| RUVBL2 | 104089.106 | 110228.506 | -0.021 | 0.043 | FALSE |
| EIF3C | 29951.346 | 32164.189 | -0.023 | 0.047 | FALSE |
| RPS24 | 45519.802 | 38407.562 | -0.023 | 0.043 | FALSE |
| CRTAP | 3832.671 | 6215.862 | -0.024 | 0.012 | FALSE |
| PRKCSH | 16131.310 | 15142.595 | -0.024 | 0.034 | FALSE |
| LBR | 4989.105 | 4695.729 | -0.026 | 0.010 | FALSE |
| LMNB1 | 180133.211 | 171858.322 | -0.026 | 0.055 | FALSE |
| GANAB | 13367.391 | 13543.566 | -0.027 | 0.076 | FALSE |
| RIF1 | 34440.733 | 32957.101 | -0.028 | 0.042 | FALSE |
| TMX1 | 43274.896 | 49184.047 | -0.031 | 0.025 | FALSE |
| CCAR1 | 16051.187 | 15192.795 | -0.032 | 0.059 | FALSE |
| OAT | 4665.420 | 4628.892 | -0.033 | 0.183 | FALSE |
| IARS1 | 17739.754 | 19316.678 | -0.034 | 0.046 | FALSE |
| ANKFY1 | 116781.137 | 121720.934 | -0.035 | 0.048 | FALSE |
| ADAR | 1475.483 | 1601.995 | -0.036 | 0.039 | FALSE |
| TMPO | 20942.592 | 20636.728 | -0.037 | 0.077 | FALSE |
| PPIL4 | 30178.238 | 35072.631 | -0.040 | 0.102 | FALSE |
| UNC45A | 1953.179 | 1845.079 | -0.040 | 0.083 | FALSE |
| RPL9 | 32724.179 | 32792.015 | -0.041 | 0.014 | FALSE |
| EPHB2 | 2755.857 | 2536.544 | -0.042 | 0.046 | FALSE |
| COX4I1 | 46278.123 | 47020.568 | -0.043 | 0.030 | FALSE |
| PSMD6 | 33168.639 | 35830.313 | -0.043 | 0.052 | FALSE |
| SNRPB | 208743.336 | 187658.449 | -0.043 | 0.030 | FALSE |
| CKAP5 | 178528.645 | 182228.179 | -0.044 | 0.053 | FALSE |
| TKT | 28899.785 | 28655.809 | -0.044 | 0.015 | FALSE |
| CEP70 | 51009.454 | 40605.553 | -0.045 | 0.042 | FALSE |
| KCTD5 | 415523.013 | 454970.394 | -0.046 | 0.066 | FALSE |
| MBD3 | 6034.760 | 10776.810 | -0.046 | 0.030 | FALSE |
| SNRPA1 | 64008.329 | 67180.762 | -0.048 | 0.053 | FALSE |
| RAB11FIP1 | 612.690 | 1129.824 | -0.050 | 0.114 | FALSE |

| | | | | | |
|---|---|---|---|---|---|
| EDC4 | 999.025 | 1660.245 | -0.050 | 0.044 | FALSE |
| PRDX2 | 63969.241 | 57709.491 | -0.051 | 0.039 | FALSE |
| EMD | 20249.948 | 21504.096 | -0.054 | 0.080 | FALSE |
| AIMP1 | 14668.556 | 13691.415 | -0.054 | 0.069 | FALSE |
| NES | 20631.844 | 21179.342 | -0.056 | 0.108 | FALSE |
| SRSF2 | 307423.356 | 257626.570 | -0.056 | 0.044 | FALSE |
| OGT | 12517.665 | 13009.063 | -0.057 | 0.064 | FALSE |
| HSPA8 | 1125758.072 | 1139872.883 | -0.057 | 0.080 | FALSE |
| CAP1 | 67821.264 | 68806.979 | -0.059 | 0.626 | FALSE |
| GOLGA3 | 16189.615 | 15960.880 | -0.060 | 0.100 | FALSE |
| USP15 | 271826.792 | 269948.564 | -0.061 | 0.166 | FALSE |
| CMBL | 23297.613 | 22190.734 | -0.061 | 0.067 | FALSE |
| ASPH | 6648.046 | 6702.314 | -0.061 | 0.074 | FALSE |
| EPB41L3 | 50030.204 | 55349.061 | -0.062 | 0.053 | FALSE |
| SMC2 | 10806.508 | 10757.846 | -0.062 | 0.127 | FALSE |
| MTHFD1 | 1904.068 | 1860.535 | -0.062 | 0.133 | FALSE |
| NUP133 | 5804.126 | 5229.542 | -0.062 | 0.121 | FALSE |
| JAK1 | 123055.587 | 139140.852 | -0.063 | 0.059 | FALSE |
| TUBGCP3 | 2115.702 | 1765.492 | -0.064 | 0.121 | FALSE |
| SF3B6 | 164987.789 | 154096.786 | -0.068 | 0.256 | FALSE |
| EIF3K | 40684.876 | 39710.256 | -0.069 | 0.113 | FALSE |
| RCN1 | 24699.280 | 18895.643 | -0.069 | 0.205 | FALSE |
| GTF2I | 2615.179 | 2453.460 | -0.069 | 0.079 | FALSE |
| EIF3M | 38322.547 | 44825.221 | -0.070 | 0.117 | FALSE |
| TAF4 | 546721.659 | 526272.156 | -0.070 | 0.135 | FALSE |
| EPPK1 | 1555.651 | 2112.671 | -0.070 | 0.026 | FALSE |
| HSPA9 | 222041.209 | 224204.176 | -0.071 | 0.384 | FALSE |
| DSC1 | 13202.291 | 11256.875 | -0.072 | 0.040 | FALSE |
| GEMIN4 | 6758.216 | 6165.218 | -0.074 | 0.131 | FALSE |
| ARCN1 | 10239.635 | 10518.080 | -0.075 | 0.200 | FALSE |
| C1QBP | 39085.425 | 25793.123 | -0.075 | 0.072 | FALSE |
| ENO1 | 50629.268 | 36471.138 | -0.075 | 0.066 | FALSE |
| ASS1 | 13121.839 | 6923.726 | -0.075 | 0.026 | FALSE |
| ARFGEF2 | 2508.692 | 2410.147 | -0.076 | 0.061 | FALSE |
| PFKL | 19361.305 | 17065.069 | -0.077 | 0.117 | FALSE |
| RBM25 | 4106.098 | 3230.849 | -0.077 | 0.056 | FALSE |
| PRMT1 | 190906.595 | 174470.643 | -0.077 | 0.057 | FALSE |
| PGK1 | 4514.540 | 3636.327 | -0.080 | 0.095 | FALSE |
| COPRS | 442574.802 | 372964.344 | -0.081 | 0.051 | FALSE |
| POGLUT3 | 6910.191 | 7467.518 | -0.082 | 0.113 | FALSE |
| EIF3L | 67862.836 | 64545.413 | -0.083 | 0.194 | FALSE |
| LAMTOR4 | 15111.178 | 22113.602 | -0.085 | 0.186 | FALSE |
| SON | 15317.929 | 14650.928 | -0.086 | 0.048 | FALSE |

| | | | | | |
|---|---|---|---|---|---|
| TMEM109 | 44231.391 | 40678.047 | -0.087 | 0.088 | FALSE |
| MBOAT7 | 2134.739 | 1974.241 | -0.090 | 0.043 | FALSE |
| RANBP2 | 6246.419 | 6536.561 | -0.090 | 0.199 | FALSE |
| TAF12 | 322559.375 | 276382.572 | -0.091 | 0.156 | FALSE |
| PSMD13 | 56484.966 | 57488.581 | -0.092 | 0.252 | FALSE |
| CIRBP | 9589.849 | 9262.545 | -0.093 | 0.105 | FALSE |
| DCTN2 | 136999.353 | 127790.726 | -0.093 | 0.923 | FALSE |
| KIF11 | 2108261.694 | 2159440.973 | -0.093 | 0.086 | FALSE |
| VARS1 | 7393.185 | 7296.596 | -0.093 | 0.143 | FALSE |
| FRYL | 8948.351 | 8688.697 | -0.093 | 0.176 | FALSE |
| CCT7 | 188474.786 | 187997.611 | -0.094 | 0.251 | FALSE |
| QARS1 | 9245.274 | 8884.896 | -0.094 | 0.463 | FALSE |
| TXN | 140325.328 | 119208.912 | -0.094 | 0.087 | FALSE |
| PTPN1 | 10022.814 | 10588.698 | -0.094 | 0.132 | FALSE |
| COPB2 | 18097.835 | 20939.666 | -0.096 | 0.287 | FALSE |
| SCARB2 | 14638.054 | 4691.691 | -0.097 | 0.028 | FALSE |
| NOP58 | 9936.819 | 9690.075 | -0.098 | 0.142 | FALSE |
| MYO5B | 3329.633 | 3121.893 | -0.098 | 0.192 | FALSE |
| CCDC88A | 52344.975 | 45821.916 | -0.099 | 0.129 | FALSE |
| ARG1 | 50479.651 | 48759.986 | -0.100 | 0.091 | FALSE |
| CCAR2 | 7679.256 | 7826.828 | -0.101 | 0.116 | FALSE |
| TMED10 | 20963.855 | 20869.260 | -0.103 | 0.071 | FALSE |
| CHERP | 21702.341 | 19741.260 | -0.103 | 0.128 | FALSE |
| SMC4 | 10197.014 | 10676.264 | -0.103 | 0.237 | FALSE |
| ACTR1A | 155152.660 | 145860.464 | -0.104 | 0.276 | FALSE |
| MRE11 | 7901.885 | 7186.664 | -0.105 | 0.158 | FALSE |
| PFKFB2 | 35302.173 | 29559.262 | -0.106 | 0.191 | FALSE |
| PDGFA | 28481.677 | 23742.703 | -0.106 | 0.170 | FALSE |
| MOGS | 7881.052 | 7845.945 | -0.106 | 0.281 | FALSE |
| FAM114A1 | 9748.017 | 7613.296 | -0.107 | 0.076 | FALSE |
| ZYX | 53415.106 | 45968.212 | -0.109 | 0.437 | FALSE |
| CSTA | 142048.521 | 99063.474 | -0.111 | 0.046 | FALSE |
| TRIP13 | 10054.323 | 10431.998 | -0.112 | 0.073 | FALSE |
| JUP | 50926.363 | 41535.473 | -0.115 | 0.077 | FALSE |
| SNRNP200 | 16167.067 | 16675.443 | -0.115 | 0.250 | FALSE |
| NUDC | 3761.290 | 6752.015 | -0.116 | 0.060 | FALSE |
| KRT23 | 10728.562 | 9208.940 | -0.118 | 0.083 | FALSE |
| BCS1L | 1584.062 | 1460.924 | -0.118 | 0.286 | FALSE |
| NAMPT | 71007.099 | 66498.960 | -0.119 | 0.107 | FALSE |
| DARS1 | 30739.979 | 27313.105 | -0.119 | 0.593 | FALSE |
| SF3A3 | 7317.169 | 7368.981 | -0.120 | 0.133 | FALSE |
| SNRNP70 | 49320.987 | 36389.648 | -0.120 | 0.052 | FALSE |
| NME1 | 32772.773 | 26953.848 | -0.122 | 0.172 | FALSE |

| | | | | | |
|---|---|---|---|---|---|
| ERGIC3 | 13760.195 | 12186.345 | -0.122 | 0.084 | FALSE |
| ZC3H18 | 21464.164 | 18571.500 | -0.124 | 0.109 | FALSE |
| OS9 | 2700.254 | 4428.002 | -0.126 | 0.074 | FALSE |
| SEC31A | 10976.750 | 10297.187 | -0.127 | 0.544 | FALSE |
| SFXN1 | 15391.655 | 15984.067 | -0.127 | 0.268 | FALSE |
| DSP | 33771.310 | 27079.882 | -0.129 | 0.080 | FALSE |
| RPL26L1 | 20894.973 | 24036.901 | -0.130 | 0.055 | FALSE |
| TFRC | 42343.672 | 45370.747 | -0.131 | 0.058 | FALSE |
| RPL30 | 14038.010 | 13030.208 | -0.131 | 0.046 | FALSE |
| EEF1G | 150064.635 | 138771.543 | -0.133 | 0.432 | FALSE |
| CCT6A | 253238.964 | 249818.347 | -0.133 | 0.385 | FALSE |
| FAF2 | 4178.212 | 3854.398 | -0.134 | 0.398 | FALSE |
| ZC3H13 | 4300.756 | 3669.145 | -0.134 | 0.057 | FALSE |
| HNRNPUL1 | 7621.620 | 7021.978 | -0.136 | 0.129 | FALSE |
| UBQLN2 | 11899.408 | 10004.817 | -0.136 | 0.103 | FALSE |
| RPS18 | 131636.309 | 124670.825 | -0.136 | 0.124 | FALSE |
| TUBA4A | 1212085.681 | 1793697.029 | -0.137 | 0.274 | FALSE |
| PRPF19 | 16274.034 | 14918.286 | -0.137 | 0.553 | FALSE |
| EEF1D | 150350.286 | 135419.502 | -0.137 | 0.244 | FALSE |
| ANXA2 | 268529.313 | 236891.110 | -0.137 | 0.408 | FALSE |
| RBM39 | 68581.319 | 65655.473 | -0.141 | 0.074 | FALSE |
| CNP | 3424.607 | 2847.948 | -0.141 | 0.287 | FALSE |
| TPD52 | 61099.170 | 60109.808 | -0.143 | 0.359 | FALSE |
| SMC3 | 5299.267 | 4647.260 | -0.143 | 0.597 | FALSE |
| KRT78 | 286038.152 | 276048.838 | -0.144 | 0.103 | FALSE |
| ITGA3 | 4633.771 | 3734.417 | -0.145 | 0.110 | FALSE |
| CASP14 | 49959.190 | 37456.361 | -0.145 | 0.128 | FALSE |
| CLIP1 | 421.478 | 567.501 | -0.147 | 0.450 | FALSE |
| SMARCA4 | 2929.905 | 2892.900 | -0.148 | 0.220 | FALSE |
| DCTN1 | 37176.628 | 36801.444 | -0.148 | 0.688 | FALSE |
| SRP68 | 6451.466 | 6846.228 | -0.149 | 0.194 | FALSE |
| CAD | 23110.165 | 21564.612 | -0.149 | 0.388 | FALSE |
| HSD17B12 | 16335.213 | 17042.543 | -0.154 | 0.194 | FALSE |
| CCT4 | 174982.623 | 170076.202 | -0.154 | 0.323 | FALSE |
| CAP2 | 75450.678 | 67169.293 | -0.155 | 0.285 | FALSE |
| PSPC1 | 5888.958 | 4720.919 | -0.155 | 0.187 | FALSE |
| MAGEA4 | 29655.697 | 28755.949 | -0.156 | 0.278 | FALSE |
| DCTN4 | 25894.902 | 21548.474 | -0.158 | 0.591 | FALSE |
| KRT6A | 2714459.704 | 2454359.630 | -0.158 | 0.098 | FALSE |
| NUP107 | 7834.022 | 7061.649 | -0.158 | 0.367 | FALSE |
| VPS13B | 625.311 | 1046.717 | -0.159 | 0.103 | FALSE |
| CCT3 | 213877.883 | 204492.935 | -0.162 | 0.284 | FALSE |
| WIZ | 3696.877 | 3187.500 | -0.162 | 0.160 | FALSE |

| | | | | | |
|---|---|---|---|---|---|
| NUP85 | 15788.151 | 13814.386 | -0.164 | 0.166 | FALSE |
| AIMP2 | 11519.364 | 19811.724 | -0.165 | 0.093 | FALSE |
| HNRNPR | 6501.746 | 5849.805 | -0.165 | 0.105 | FALSE |
| EIF3E | 81604.176 | 77139.316 | -0.168 | 0.308 | FALSE |
| TRIM21 | 222929.183 | 207844.075 | -0.170 | 0.166 | FALSE |
| ILF2 | 21143.958 | 21835.139 | -0.171 | 0.244 | FALSE |
| BRAT1 | 7393.469 | 5970.853 | -0.171 | 0.134 | FALSE |
| CCT8 | 208339.744 | 188836.643 | -0.175 | 0.400 | FALSE |
| COPB1 | 9022.208 | 9144.113 | -0.176 | 0.422 | FALSE |
| QPCTL | 9560.959 | 9293.533 | -0.176 | 0.110 | FALSE |
| SLC1A5 | 6927.006 | 7470.181 | -0.180 | 0.109 | FALSE |
| BAG6 | 7069.276 | 7095.021 | -0.181 | 0.283 | FALSE |
| CRYBG3 | 1092.068 | 1003.637 | -0.184 | 0.277 | FALSE |
| MYH9 | 139230.171 | 142484.596 | -0.184 | 0.402 | FALSE |
| EFTUD2 | 33788.173 | 30910.129 | -0.187 | 0.528 | FALSE |
| PRMT5 | 9360061.812 | 8535058.803 | -0.188 | 0.307 | FALSE |
| STK38L | 1234560.692 | 1125866.846 | -0.188 | 0.411 | FALSE |
| SNRPA | 256468.446 | 213829.091 | -0.189 | 0.124 | FALSE |
| TAF5 | 9481.356 | 7785.815 | -0.190 | 1.098 | FALSE |
| U2SURP | 28468.638 | 26365.872 | -0.191 | 0.419 | FALSE |
| H2BC4 | 260349.261 | 328092.508 | -0.192 | 0.440 | FALSE |
| HRNR | 19312.600 | 13504.953 | -0.194 | 0.112 | FALSE |
| LMNB2 | 68592.088 | 63044.781 | -0.194 | 0.317 | FALSE |
| SPTBN1 | 178292.933 | 164356.616 | -0.197 | 0.564 | FALSE |
| FAM210A | 2910.671 | 4216.344 | -0.200 | 0.263 | FALSE |
| ATP5F1A | 115834.657 | 114681.879 | -0.201 | 0.199 | FALSE |
| KLC2 | 3758.938 | 3258.324 | -0.201 | 0.531 | FALSE |
| CPSF6 | 11870.862 | 11401.882 | -0.201 | 0.130 | FALSE |
| MCM2 | 2170.248 | 1959.774 | -0.201 | 0.417 | FALSE |
| VAPB | 13760.586 | 13150.601 | -0.202 | 0.231 | FALSE |
| CCT6B | 67675.817 | 63654.132 | -0.202 | 0.529 | FALSE |
| RPL13A | 9798.750 | 9477.542 | -0.203 | 0.085 | FALSE |
| HNRNPL | 8340.939 | 8696.680 | -0.203 | 0.098 | FALSE |
| RARS1 | 13973.372 | 12745.520 | -0.204 | 0.565 | FALSE |
| ATP5MC1 | 18140.163 | 7564.850 | -0.205 | 0.062 | FALSE |
| DYNLRB1 | 59353.979 | 49102.247 | -0.206 | 2.555 | FALSE |
| MLEC | 9149.356 | 8509.384 | -0.207 | 0.174 | FALSE |
| RPL7 | 22237.110 | 24657.748 | -0.207 | 0.060 | FALSE |
| CAT | 13410.778 | 12583.921 | -0.208 | 0.145 | FALSE |
| CAAP1 | 9139.483 | 8618.363 | -0.209 | 0.108 | FALSE |
| HNRNPH3 | 5856.249 | 4943.051 | -0.209 | 0.119 | FALSE |
| RGPD3 | 4594.671 | 4641.849 | -0.209 | 0.811 | FALSE |
| CADPS2 | 29105.318 | 26145.872 | -0.209 | 0.418 | FALSE |

| | | | | | |
|---|---|---|---|---|---|
| IKBIP | 7125.679 | 5688.429 | -0.210 | 0.221 | FALSE |
| H2BC3 | 232360.769 | 303927.856 | -0.210 | 0.344 | FALSE |
| TAGLN2 | 11378.601 | 6867.489 | -0.210 | 0.089 | FALSE |
| LYZ | 30701.464 | 20088.285 | -0.211 | 0.134 | FALSE |
| TMPO | 37591.087 | 37149.476 | -0.211 | 0.385 | FALSE |
| SMC1A | 3410.487 | 3472.432 | -0.212 | 0.900 | FALSE |
| DHX15 | 76379.082 | 72492.867 | -0.214 | 0.205 | FALSE |
| NIPSNAP1 | 19192.722 | 16577.379 | -0.214 | 0.437 | FALSE |
| SCYL2 | 48607.696 | 42335.083 | -0.214 | 0.238 | FALSE |
| LMAN2 | 14048.086 | 16291.639 | -0.214 | 0.169 | FALSE |
| XPO1 | 21404.861 | 20838.818 | -0.215 | 0.228 | FALSE |
| GSTO1 | 16580.240 | 14306.290 | -0.215 | 0.452 | FALSE |
| DAGLB | 4763.890 | 3578.451 | -0.216 | 0.160 | FALSE |
| DYNC1LI1 | 1809.534 | 1497.054 | -0.217 | 0.322 | FALSE |
| MYCBP | 2959203.095 | 2246716.095 | -0.218 | 0.214 | FALSE |
| PSMA4 | 66531.348 | 52584.090 | -0.218 | 0.323 | FALSE |
| CKAP4 | 15102.713 | 13937.755 | -0.218 | 0.259 | FALSE |
| CD44 | 5741.236 | 6185.730 | -0.219 | 0.179 | FALSE |
| ESYT1 | 26973.086 | 25649.614 | -0.221 | 0.316 | FALSE |
| ORMDL1 | 20154.945 | 15792.891 | -0.221 | 0.363 | FALSE |
| ITGA5 | 1520.782 | 1115.049 | -0.222 | 0.220 | FALSE |
| CAPZA2 | 36606.050 | 35070.863 | -0.224 | 0.656 | FALSE |
| CSE1L | 147855.048 | 135677.213 | -0.225 | 0.221 | FALSE |
| DHCR24 | 2926.111 | 3999.904 | -0.227 | 0.205 | FALSE |
| DDRGK1 | 6379.057 | 5535.686 | -0.227 | 0.585 | FALSE |
| DDOST | 20516.066 | 19017.562 | -0.229 | 0.285 | FALSE |
| CYB5B | 48149.345 | 50659.330 | -0.234 | 0.248 | FALSE |
| PFKP | 35338.732 | 31706.241 | -0.234 | 0.450 | FALSE |
| RFC3 | 10755.778 | 16274.894 | -0.234 | 0.358 | FALSE |
| P3H1 | 6099.957 | 5174.580 | -0.235 | 0.370 | FALSE |
| RIOK1 | 160951.701 | 126222.722 | -0.235 | 0.674 | FALSE |
| CLTC | 75541.560 | 79892.319 | -0.236 | 0.419 | FALSE |
| H3-4 | 155033.817 | 128721.697 | -0.237 | 0.291 | FALSE |
| UNC93B1 | 4187.494 | 4014.885 | -0.237 | 0.207 | FALSE |
| S100A8 | 225687.339 | 140429.778 | -0.237 | 0.133 | FALSE |
| COPA | 16938.342 | 15192.760 | -0.238 | 0.437 | FALSE |
| PTRH2 | 23848.839 | 23908.003 | -0.240 | 0.163 | FALSE |
| PKP1 | 4733.430 | 4046.129 | -0.240 | 0.145 | FALSE |
| SNRPD2 | 3806836.500 | 3142502.979 | -0.244 | 0.797 | FALSE |
| RAB1A | 278426.889 | 241510.093 | -0.244 | 0.233 | FALSE |
| WTAP | 5872.030 | 4735.878 | -0.245 | 0.292 | FALSE |
| RPA1 | 1951.494 | 2786.586 | -0.246 | 0.297 | FALSE |
| PPP2R1A | 13375.635 | 12660.782 | -0.246 | 0.369 | FALSE |

| | | | | | |
|---|---|---|---|---|---|
| GLMN | 1314.672 | 1401.967 | -0.247 | 0.201 | FALSE |
| TAF6 | 16980.473 | 14104.200 | -0.248 | 0.770 | FALSE |
| IPO5 | 2841.223 | 4903.539 | -0.249 | 0.150 | FALSE |
| PRPF6 | 13328.382 | 11747.559 | -0.249 | 0.947 | FALSE |
| SNRPF | 258830.592 | 248666.087 | -0.252 | 0.615 | FALSE |
| DNAJB12 | 4025.061 | 5801.145 | -0.254 | 0.270 | FALSE |
| RPS26 | 142493.684 | 154532.097 | -0.254 | 0.142 | FALSE |
| CTNNA1 | 10568.333 | 44957.301 | -0.254 | 0.205 | FALSE |
| PRPF4 | 1472.022 | 1874.099 | -0.255 | 0.573 | FALSE |
| NUP160 | 6151.151 | 5282.987 | -0.256 | 0.227 | FALSE |
| AHNAK2 | 336.473 | 237.355 | -0.256 | 0.311 | FALSE |
| RPS5 | 126688.419 | 118281.137 | -0.257 | 0.205 | FALSE |
| KIF5B | 5493.467 | 5133.040 | -0.259 | 0.357 | FALSE |
| VCP | 119817.188 | 109614.218 | -0.259 | 1.065 | FALSE |
| SORT1 | 1007.423 | 1685.388 | -0.263 | 0.197 | FALSE |
| ARL6IP5 | 76193.459 | 71521.070 | -0.264 | 0.220 | FALSE |
| ACSL3 | 15170.483 | 12434.613 | -0.269 | 0.538 | FALSE |
| FAM3C | 2803.087 | 4179.169 | -0.269 | 1.602 | FALSE |
| ATP2A2 | 11193.201 | 12073.145 | -0.271 | 0.296 | FALSE |
| PRPF8 | 10308.867 | 10082.819 | -0.271 | 0.555 | FALSE |
| MMS19 | 2217.693 | 1928.619 | -0.271 | 0.232 | FALSE |
| SNRPD1 | 1620016.467 | 2573572.933 | -0.272 | 0.374 | FALSE |
| CNOT9 | 3704.891 | 4732.543 | -0.272 | 0.302 | FALSE |
| SURF4 | 26614.228 | 22934.867 | -0.273 | 0.346 | FALSE |
| KRT7 | 462564.167 | 418284.544 | -0.275 | 0.250 | FALSE |
| HADHB | 91366.753 | 85556.187 | -0.276 | 0.392 | FALSE |
| KRT6B | 3411163.185 | 3000278.407 | -0.277 | 0.153 | FALSE |
| SYNCRIP | 2739.669 | 2443.232 | -0.278 | 0.379 | FALSE |
| LRPPRC | 15070.716 | 12893.138 | -0.282 | 0.347 | FALSE |
| RAB34 | 33302.139 | 29427.092 | -0.282 | 0.353 | FALSE |
| HMOX2 | 21177.299 | 19447.174 | -0.282 | 0.283 | FALSE |
| SMARCC2 | 8300.684 | 7392.608 | -0.284 | 0.641 | FALSE |
| RTN4 | 24674.790 | 20005.405 | -0.284 | 0.195 | FALSE |
| DOCK7 | 647.210 | 938.919 | -0.285 | 0.319 | FALSE |
| NSF | 16115.970 | 15335.696 | -0.287 | 0.405 | FALSE |
| SERPINB4 | 21025.705 | 18561.996 | -0.288 | 0.158 | FALSE |
| GPX8 | 19410.679 | 19859.065 | -0.291 | 0.648 | FALSE |
| MYL6 | 174903.508 | 151434.590 | -0.291 | 0.347 | FALSE |
| DHRS7 | 17724.407 | 15321.287 | -0.291 | 0.497 | FALSE |
| TSPAN10 | 4080.679 | 3473.350 | -0.292 | 0.244 | FALSE |
| BCAP31 | 99647.943 | 86323.845 | -0.292 | 0.691 | FALSE |
| SNRPG | 1026543.722 | 821418.083 | -0.293 | 0.459 | FALSE |
| ACTN1 | 52811.200 | 47814.091 | -0.293 | 0.575 | FALSE |

| | | | | | |
|---|---|---|---|---|---|
| SPTAN1 | 174599.745 | 155414.750 | -0.293 | 0.829 | FALSE |
| ACAD9 | 2913.224 | 3624.211 | -0.293 | 0.171 | FALSE |
| KRT1 | ########### | 12004625.829 | -0.294 | 0.234 | FALSE |
| PSMB7 | 6725.174 | 4601.648 | -0.294 | 0.259 | FALSE |
| ACTR10 | 337189.373 | 217311.913 | -0.295 | 0.722 | FALSE |
| ACTN4 | 37665.303 | 31438.216 | -0.295 | 0.429 | FALSE |
| LSS | 11373.251 | 9117.836 | -0.296 | 1.424 | FALSE |
| GCN1 | 20586.036 | 17345.933 | -0.298 | 0.403 | FALSE |
| ZW10 | 3634.057 | 5479.300 | -0.299 | 0.219 | FALSE |
| TNPO1 | 21805.977 | 18858.545 | -0.300 | 0.327 | FALSE |
| CIP2A | 1613.942 | 2179.782 | -0.301 | 0.216 | FALSE |
| CORO2A | 4894.808 | 3617.584 | -0.301 | 0.496 | FALSE |
| TBRG4 | 2671.164 | 3825.330 | -0.301 | 0.276 | FALSE |
| RPLP0 | 46755.462 | 48790.724 | -0.302 | 0.178 | FALSE |
| RPN2 | 26887.248 | 23032.167 | -0.303 | 0.434 | FALSE |
| LAMP1 | 29864.825 | 26677.894 | -0.305 | 0.247 | FALSE |
| NEXN | 10929.813 | 7106.318 | -0.306 | 0.385 | FALSE |
| EXOC4 | 1703.275 | 1196.603 | -0.306 | 0.397 | FALSE |
| PUF60 | 50144.269 | 44354.622 | -0.307 | 0.447 | FALSE |
| RAB11B | 220300.919 | 202253.143 | -0.307 | 0.236 | FALSE |
| TRAP1 | 21943.344 | 23718.189 | -0.309 | 0.234 | FALSE |
| SSR3 | 21024.739 | 21757.011 | -0.310 | 0.240 | FALSE |
| EEF1B2 | 98492.103 | 81915.878 | -0.310 | 0.556 | FALSE |
| CTNNB1 | 10094.833 | 8751.947 | -0.311 | 0.343 | FALSE |
| HNRNPA1 | 33383.083 | 20317.801 | -0.312 | 0.178 | FALSE |
| HNRNPH2 | 758267.962 | 705323.506 | -0.312 | 0.648 | FALSE |
| ALB | 22578.537 | 24798.686 | -0.312 | 0.267 | FALSE |
| LARS1 | 6886.770 | 5582.810 | -0.313 | 0.719 | FALSE |
| ATP5F1C | 30893.804 | 27819.940 | -0.314 | 0.393 | FALSE |
| APMAP | 57391.229 | 52432.205 | -0.317 | 0.402 | FALSE |
| HNRNPA2B1 | 30576.450 | 20769.269 | -0.320 | 0.235 | FALSE |
| RPS7 | 101786.169 | 98520.081 | -0.320 | 0.259 | FALSE |
| EEF2 | 31132.049 | 19317.461 | -0.323 | 0.254 | FALSE |
| LUC7L | 122897.781 | 82013.898 | -0.324 | 0.200 | FALSE |
| CLNS1A | 1614208.848 | 1319836.780 | -0.325 | 1.048 | FALSE |
| CAPZA1 | 112429.891 | 96930.198 | -0.326 | 1.265 | FALSE |
| HRAS | 6967.356 | 6350.572 | -0.327 | 0.506 | FALSE |
| RPS28 | 239812.639 | 217409.336 | -0.328 | 0.229 | FALSE |
| SLC3A2 | 42797.888 | 41136.911 | -0.330 | 0.159 | FALSE |
| SCD | 25209.139 | 19804.835 | -0.331 | 0.434 | FALSE |
| RAB10 | 117654.610 | 104287.632 | -0.331 | 0.503 | FALSE |
| PRKAR1A | 4879.208 | 3697.242 | -0.331 | 1.234 | FALSE |
| RPS20 | 134440.500 | 121486.734 | -0.333 | 0.264 | FALSE |

| | | | | | |
|---|---|---|---|---|---|
| RALY | 28222.406 | 23520.345 | -0.333 | 0.310 | FALSE |
| ACTB | 8068552.580 | 5519277.536 | -0.334 | 0.161 | FALSE |
| MTDH | 22105.786 | 13724.736 | -0.336 | 0.405 | FALSE |
| ACTG1 | 8041112.348 | 5500540.203 | -0.336 | 0.142 | FALSE |
| CYB5A | 12799.249 | 11351.668 | -0.336 | 0.318 | FALSE |
| PDHB | 3898.241 | 6261.666 | -0.337 | 0.219 | FALSE |
| NELFB | 10364.912 | 8972.468 | -0.338 | 1.430 | FALSE |
| NPTN | 2068.225 | 3407.975 | -0.338 | 0.513 | FALSE |
| PNN | 15538.068 | 13112.828 | -0.339 | 0.237 | FALSE |
| KATNAL2 | 9800.779 | 7876.219 | -0.340 | 1.350 | FALSE |
| S100A7L2 | 21756.241 | 15216.114 | -0.342 | 0.372 | FALSE |
| LRRC59 | 149910.939 | 137325.966 | -0.342 | 0.198 | FALSE |
| ATP5F1B | 107421.424 | 102821.083 | -0.343 | 0.282 | FALSE |
| RPS3 | 207211.377 | 183205.078 | -0.344 | 0.262 | FALSE |
| RPN1 | 45878.567 | 41186.943 | -0.344 | 0.467 | FALSE |
| RAB5C | 143439.717 | 114522.922 | -0.345 | 0.253 | FALSE |
| PHB2 | 33013.915 | 27886.771 | -0.346 | 1.294 | FALSE |
| HSPB1 | 777399.548 | 678232.488 | -0.346 | 0.579 | FALSE |
| HSD17B11 | 18124.938 | 14231.289 | -0.347 | 1.371 | FALSE |
| RAN | 67705.639 | 51216.599 | -0.350 | 0.382 | FALSE |
| CSTF2 | 9721.242 | 7657.707 | -0.351 | 2.191 | FALSE |
| AFDN | 1422.739 | 2034.334 | -0.353 | 0.255 | FALSE |
| KRT16 | 3115522.828 | 2790420.606 | -0.354 | 0.333 | FALSE |
| DCTN5 | 23632.818 | 19849.033 | -0.355 | 0.725 | FALSE |
| BAG2 | 32232.396 | 27814.825 | -0.358 | 0.829 | FALSE |
| ATL2 | 3152.846 | 3940.415 | -0.359 | 0.238 | FALSE |
| RAB7A | 82845.733 | 72366.093 | -0.360 | 0.334 | FALSE |
| RPL6 | 10694.886 | 9268.683 | -0.360 | 0.125 | FALSE |
| SNRPE | 726554.292 | 574795.229 | -0.360 | 0.670 | FALSE |
| RAB2A | 60661.469 | 54477.712 | -0.361 | 0.425 | FALSE |
| GSN | 8519.074 | 6320.226 | -0.363 | 0.376 | FALSE |
| DNAAF5 | 3491.058 | 2514.087 | -0.363 | 0.530 | FALSE |
| HADHA | 83186.295 | 83564.190 | -0.364 | 0.386 | FALSE |
| NDUFS3 | 3765.527 | 4990.108 | -0.365 | 0.602 | FALSE |
| RPL34 | 20083.373 | 23138.754 | -0.365 | 0.106 | FALSE |
| RACK1 | 131875.496 | 140630.023 | -0.365 | 0.274 | FALSE |
| RPL12 | 29047.746 | 28391.881 | -0.366 | 0.106 | FALSE |
| MYH14 | 16662.042 | 15487.117 | -0.366 | 0.975 | FALSE |
| SEC61G | 80866.643 | 69093.036 | -0.366 | 0.227 | FALSE |
| DIABLO | 5463.083 | 4400.431 | -0.367 | 0.631 | FALSE |
| KPNB1 | 91427.828 | 83271.331 | -0.368 | 0.374 | FALSE |
| STAT3 | 49590.933 | 44918.669 | -0.370 | 0.295 | FALSE |
| RPS8 | 87970.377 | 82651.578 | -0.371 | 0.339 | FALSE |

| | | | | | |
|---|---|---|---|---|---|
| TLN1 | 277.222 | 296.260 | -0.371 | 1.669 | FALSE |
| PCYOX1 | 28690.898 | 22592.236 | -0.371 | 0.839 | FALSE |
| DLG1 | 29171.081 | 22696.570 | -0.372 | 0.514 | FALSE |
| NCBP2 | 60933.200 | 41124.129 | -0.372 | 0.242 | FALSE |
| LIMA1 | 27622.668 | 20533.709 | -0.372 | 0.605 | FALSE |
| XPO7 | 3236.023 | 2429.111 | -0.374 | 1.942 | FALSE |
| HNRNPC | 93493.339 | 80051.195 | -0.375 | 0.301 | FALSE |
| PI4KA | 580.236 | 694.189 | -0.378 | 0.593 | FALSE |
| KRT3 | 837400.971 | 809834.171 | -0.379 | 0.287 | FALSE |
| S100A7 | 110750.821 | 82738.941 | -0.380 | 0.700 | FALSE |
| RPL18 | 11836.037 | 13252.312 | -0.384 | 0.141 | FALSE |
| NCBP1 | 12666.348 | 9271.232 | -0.384 | 0.225 | FALSE |
| RPS10 | 52627.803 | 51824.185 | -0.384 | 0.291 | FALSE |
| PSME3 | 9606.919 | 16851.078 | -0.384 | 0.156 | FALSE |
| SEC22B | 132634.304 | 115816.897 | -0.387 | 0.570 | FALSE |
| IPO4 | 3411.832 | 4505.210 | -0.388 | 0.199 | FALSE |
| GNB2 | 83979.735 | 62454.572 | -0.388 | 0.316 | FALSE |
| PHB | 42911.009 | 34584.819 | -0.390 | 0.818 | FALSE |
| MAGOHB | 15849.026 | 11298.078 | -0.391 | 0.374 | FALSE |
| TRAPPC3 | 9675.090 | 7883.923 | -0.392 | 0.279 | FALSE |
| BCL2L1 | 15719.313 | 12238.891 | -0.392 | 0.498 | FALSE |
| GOSR1 | 5786.765 | 7976.427 | -0.393 | 0.248 | FALSE |
| PLAA | 9136.839 | 6989.039 | -0.398 | 0.355 | FALSE |
| BSG | 17551.034 | 14747.526 | -0.401 | 0.504 | FALSE |
| LGALS1 | 86027.230 | 69116.408 | -0.402 | 0.258 | FALSE |
| NCAPG | 5827.385 | 8386.239 | -0.407 | 0.265 | FALSE |
| PDS5A | 1902.283 | 2833.096 | -0.410 | 0.358 | FALSE |
| PRKDC | 18644.061 | 16016.520 | -0.410 | 0.296 | FALSE |
| CISD2 | 9232.424 | 12285.291 | -0.412 | 0.570 | FALSE |
| RRAS2 | 5999.488 | 7752.665 | -0.413 | 0.252 | FALSE |
| TBC1D8B | 1832.911 | 1316.085 | -0.413 | 1.604 | FALSE |
| PFKFB3 | 94940.006 | 79260.502 | -0.413 | 0.476 | FALSE |
| MYL12A | 249472.008 | 197835.737 | -0.414 | 0.614 | FALSE |
| ATP5MF | 28619.345 | 44272.144 | -0.415 | 0.222 | FALSE |
| RAB1B | 274063.435 | 233485.443 | -0.415 | 0.382 | FALSE |
| LIN7A | 3830.129 | 2653.559 | -0.417 | 0.499 | FALSE |
| PRSS1 | 1256873.253 | 61838.781 | -0.420 | 0.139 | FALSE |
| CDC42 | 28811.703 | 25970.514 | -0.420 | 0.286 | FALSE |
| SRPRB | 25057.892 | 20597.234 | -0.420 | 0.403 | FALSE |
| RPS19 | 151630.597 | 117258.240 | -0.421 | 0.397 | FALSE |
| PGRMC2 | 47222.116 | 40259.888 | -0.421 | 0.997 | FALSE |
| TIMM50 | 20803.075 | 17134.018 | -0.421 | 0.318 | FALSE |
| TMX3 | 1983.254 | 2107.736 | -0.421 | 0.633 | FALSE |

| | | | | | |
|---|---|---|---|---|---|
| PI4K2A | 2102.251 | 2313.531 | -0.422 | 0.448 | FALSE |
| REEP5 | 11723.300 | 14254.258 | -0.423 | 0.328 | FALSE |
| SLIRP | 18108.538 | 14816.699 | -0.424 | 0.468 | FALSE |
| CLTA | 46729.928 | 34683.180 | -0.425 | 1.825 | FALSE |
| LUC7L3 | 115791.494 | 83430.907 | -0.427 | 0.273 | FALSE |
| TMEM33 | 30204.002 | 28273.981 | -0.428 | 0.602 | FALSE |
| S100A9 | 157311.190 | 97408.129 | -0.428 | 0.242 | FALSE |
| CCT2 | 164450.632 | 125085.227 | -0.431 | 0.751 | FALSE |
| RAB5B | 52396.026 | 41342.928 | -0.432 | 0.670 | FALSE |
| MYOF | 3910.760 | 2642.897 | -0.432 | 0.364 | FALSE |
| DCTN3 | 75559.992 | 54326.437 | -0.433 | 1.879 | FALSE |
| DYNC1H1 | 23513.463 | 19153.855 | -0.434 | 1.258 | FALSE |
| HTATIP2 | 9184.753 | 7306.615 | -0.435 | 1.262 | FALSE |
| SLC25A10 | 6986.056 | 5006.569 | -0.435 | 0.403 | FALSE |
| NDUFA4 | 89810.134 | 64722.203 | -0.435 | 0.445 | FALSE |
| FADS2 | 5701.445 | 4211.243 | -0.436 | 0.888 | FALSE |
| DPM1 | 32655.785 | 24782.506 | -0.436 | 0.956 | FALSE |
| LMAN1 | 8435.738 | 5280.840 | -0.440 | 0.360 | FALSE |
| RAB18 | 39365.851 | 31596.758 | -0.443 | 0.631 | FALSE |
| AUP1 | 12838.896 | 9208.949 | -0.443 | 0.683 | FALSE |
| PABPC4 | 52102.143 | 50044.071 | -0.443 | 0.305 | FALSE |
| ATL3 | 36390.739 | 33471.502 | -0.444 | 0.513 | FALSE |
| RPS12 | 122453.911 | 113244.953 | -0.444 | 0.278 | FALSE |
| CANX | 71597.284 | 63234.729 | -0.445 | 0.531 | FALSE |
| FKBP8 | 5404.535 | 7140.346 | -0.446 | 0.538 | FALSE |
| TMOD3 | 87143.920 | 68171.353 | -0.446 | 0.680 | FALSE |
| POR | 4224.382 | 3580.580 | -0.447 | 0.781 | FALSE |
| SYMPK | 1357.698 | 1335.921 | -0.448 | 0.335 | FALSE |
| EIF4A3 | 18912.701 | 12885.799 | -0.448 | 0.510 | FALSE |
| SSR1 | 28193.008 | 21946.130 | -0.448 | 0.961 | FALSE |
| SRSF6 | 575921.579 | 402527.521 | -0.449 | 0.247 | FALSE |
| SRSF5 | 313091.458 | 214434.679 | -0.450 | 0.340 | FALSE |
| RAB14 | 85747.307 | 70410.386 | -0.451 | 0.492 | FALSE |
| ALDH3A2 | 7923.837 | 5243.917 | -0.456 | 0.613 | FALSE |
| COX6C | 10331.959 | 7484.044 | -0.461 | 0.599 | FALSE |
| GBA | 3551.436 | 4698.615 | -0.462 | 0.418 | FALSE |
| SRP14 | 48192.325 | 42208.439 | -0.462 | 0.784 | FALSE |
| SND1 | 812.283 | 781.311 | -0.465 | 0.774 | FALSE |
| GNG12 | 43017.663 | 34951.151 | -0.466 | 0.312 | FALSE |
| SGPL1 | 12613.329 | 9868.199 | -0.467 | 0.614 | FALSE |
| RAP1A | 37697.065 | 26919.155 | -0.467 | 0.496 | FALSE |
| RPS9 | 31535.026 | 22962.670 | -0.469 | 0.357 | FALSE |
| IMPDH2 | 10256.356 | 7523.065 | -0.470 | 0.670 | FALSE |

| | | | | | |
|---|---|---|---|---|---|
| RPS16 | 104122.344 | 91637.311 | -0.471 | 0.454 | FALSE |
| MOB1A | 108567.577 | 78078.461 | -0.471 | 1.097 | FALSE |
| BZW2 | 18090.206 | 15678.687 | -0.472 | 1.117 | FALSE |
| DDX39A | 2147.427 | 1198.512 | -0.473 | 0.451 | FALSE |
| TFIP11 | 3514.850 | 2485.703 | -0.473 | 0.283 | FALSE |
| ATP1A1 | 13255.228 | 11601.901 | -0.475 | 0.717 | FALSE |
| RPS14 | 205962.143 | 160470.754 | -0.475 | 0.408 | FALSE |
| LARP1 | 7130.586 | 4843.974 | -0.478 | 0.504 | FALSE |
| USO1 | 2561.501 | 2037.672 | -0.478 | 0.531 | FALSE |
| ILF3 | 5920.135 | 4031.174 | -0.479 | 0.358 | FALSE |
| SLC25A13 | 4160.137 | 5872.517 | -0.481 | 0.343 | FALSE |
| RAB15 | 47468.074 | 38623.152 | -0.485 | 0.409 | FALSE |
| RAB33B | 56961.689 | 46347.782 | -0.485 | 0.409 | FALSE |
| PGAM5 | 15219.335 | 10568.015 | -0.487 | 0.650 | FALSE |
| RPS21 | 138855.709 | 106799.218 | -0.488 | 0.346 | FALSE |
| B2M | 38835.015 | 34923.764 | -0.489 | 0.336 | FALSE |
| ACADVL | 13026.877 | 10124.522 | -0.491 | 0.541 | FALSE |
| RPS2 | 45111.167 | 35228.627 | -0.493 | 0.702 | FALSE |
| KRT9 | 7343786.756 | 5269582.178 | -0.494 | 0.372 | FALSE |
| DYNC1I2 | 20916.744 | 16696.121 | -0.495 | 0.714 | FALSE |
| NUP205 | 3370.625 | 2788.428 | -0.497 | 0.675 | FALSE |
| RPL14 | 13514.400 | 10870.969 | -0.497 | 0.253 | FALSE |
| KCTD2 | 168617.295 | 160540.551 | -0.498 | 0.526 | FALSE |
| RPSA | 171945.289 | 144981.777 | -0.498 | 0.400 | FALSE |
| RPS4X | 97064.381 | 77209.007 | -0.498 | 0.323 | FALSE |
| IPO8 | 123254.228 | 111176.708 | -0.499 | 0.193 | FALSE |
| RPL4 | 8485.512 | 5842.838 | -0.502 | 0.150 | FALSE |
| IGHG2 | 12211.636 | 5756.033 | -0.503 | 0.261 | FALSE |
| NPM3 | 7934.798 | 7671.446 | -0.504 | 0.414 | FALSE |
| MOB2 | 108051.239 | 73482.422 | -0.505 | 0.409 | FALSE |
| SNX8 | 5009.439 | 6381.466 | -0.506 | 0.502 | FALSE |
| SUN2 | 139236.532 | 119299.202 | -0.507 | 0.321 | FALSE |
| RPS25 | 275086.653 | 232772.517 | -0.507 | 0.441 | FALSE |
| RPL7A | 9659.571 | 9156.639 | -0.509 | 0.142 | FALSE |
| PGRMC1 | 240138.592 | 195486.150 | -0.510 | 1.694 | FALSE |
| RPL13 | 11836.164 | 25559.265 | -0.510 | 0.131 | FALSE |
| GNAI3 | 25527.857 | 19941.456 | -0.511 | 0.456 | FALSE |
| GNG5 | 18809.804 | 13415.886 | -0.512 | 0.256 | FALSE |
| GET4 | 17633.388 | 12989.866 | -0.513 | 1.062 | FALSE |
| DDX23 | 2996.635 | 2164.409 | -0.514 | 1.233 | FALSE |
| DPP7 | 421789.590 | 258234.330 | -0.516 | 1.487 | FALSE |
| RPL27 | 47864.536 | 44485.689 | -0.517 | 0.170 | FALSE |
| RPS3A | 116043.024 | 94881.428 | -0.517 | 0.311 | FALSE |

| | | | | | |
|---|---|---|---|---|---|
| GOSR2 | 6751.965 | 4970.058 | -0.520 | 0.527 | FALSE |
| MYH10 | 38832.952 | 36181.170 | -0.520 | 0.376 | FALSE |
| SLC25A5 | 400803.779 | 329590.207 | -0.522 | 0.334 | FALSE |
| RDH11 | 32674.340 | 25246.711 | -0.524 | 0.466 | FALSE |
| RPL19 | 18745.161 | 15799.342 | -0.533 | 0.182 | FALSE |
| BAX | 67554.960 | 56552.823 | -0.533 | 0.401 | FALSE |
| ACIN1 | 79415.030 | 52815.140 | -0.538 | 0.226 | FALSE |
| SLC25A3 | 87247.781 | 74053.648 | -0.543 | 0.405 | FALSE |
| CFAP20 | 11979.137 | 16067.323 | -0.544 | 0.465 | FALSE |
| THRAP3 | 736133.603 | 468516.069 | -0.545 | 0.211 | FALSE |
| RAP2B | 11322.090 | 15014.086 | -0.547 | 0.433 | FALSE |
| NUDT21 | 37266.639 | 27730.309 | -0.547 | 0.594 | FALSE |
| PSMD9 | 8028.300 | 10639.608 | -0.548 | 0.360 | FALSE |
| SLC7A5 | 18544.703 | 26048.360 | -0.548 | 0.250 | FALSE |
| SLC35A4 | 35098.704 | 23102.058 | -0.553 | 0.287 | FALSE |
| VAT1 | 92222.148 | 71422.113 | -0.553 | 1.166 | FALSE |
| POLR2G | 126621.558 | 90662.164 | -0.554 | 0.705 | FALSE |
| DNAJC19 | 9794.503 | 11121.848 | -0.555 | 0.457 | FALSE |
| EXOC7 | 2416.675 | 3290.591 | -0.555 | 0.531 | FALSE |
| RANGAP1 | 22666.413 | 14576.274 | -0.558 | 0.632 | FALSE |
| PABPC1 | 94846.690 | 74995.842 | -0.559 | 0.534 | FALSE |
| ELAVL1 | 3362.284 | 3719.772 | -0.563 | 0.730 | FALSE |
| TMEM214 | 3367.939 | 3415.158 | -0.563 | 0.316 | FALSE |
| IGKV2-29 | 6422413.278 | 4073450.028 | -0.563 | 0.166 | FALSE |
| MOV10 | 13610.697 | 9781.424 | -0.564 | 0.365 | FALSE |
| HM13 | 29328.914 | 19393.257 | -0.564 | 0.719 | FALSE |
| IPO9 | 2850.083 | 8336.880 | -0.565 | 0.236 | FALSE |
| SREK1 | 5105.235 | 4423.789 | -0.573 | 0.355 | FALSE |
| SSR4 | 67422.597 | 53708.302 | -0.575 | 0.824 | FALSE |
| SEC61B | 169283.507 | 137797.840 | -0.576 | 0.501 | FALSE |
| CHP1 | 17962.803 | 20982.616 | -0.577 | 0.933 | FALSE |
| RPL23A | 28162.801 | 27513.702 | -0.580 | 0.166 | FALSE |
| SLC25A11 | 3863.823 | 4209.566 | -0.581 | 0.467 | FALSE |
| RPS17 | 212114.576 | 172958.533 | -0.588 | 0.449 | FALSE |
| RPS11 | 139495.513 | 101949.372 | -0.588 | 0.359 | FALSE |
| PDHA1 | 3357.072 | 4087.916 | -0.589 | 0.438 | FALSE |
| GAS6 | 4462.157 | 3689.091 | -0.590 | 0.411 | FALSE |
| SEC61A1 | 96655.425 | 70576.446 | -0.599 | 0.422 | FALSE |
| RPL31 | 24982.478 | 20121.074 | -0.603 | 0.178 | FALSE |
| RPL8 | 27517.750 | 24208.134 | -0.605 | 0.180 | FALSE |
| CYP51A1 | 18879.339 | 14233.279 | -0.607 | 0.822 | FALSE |
| M6PR | 25059.355 | 23337.763 | -0.607 | 0.526 | FALSE |
| RALA | 37435.147 | 26423.661 | -0.607 | 0.667 | FALSE |

| | | | | | |
|---|---|---|---|---|---|
| SH3GLB1 | 6905.528 | 2977.204 | -0.609 | 0.725 | FALSE |
| TMEM97 | 12404.413 | 8832.747 | -0.610 | 1.972 | FALSE |
| PRPF40A | 5002.366 | 3158.964 | -0.612 | 1.693 | FALSE |
| ALCAM | 2555.116 | 3077.772 | -0.617 | 0.428 | FALSE |
| CLCC1 | 11471.951 | 8812.243 | -0.621 | 0.702 | FALSE |
| ITGAV | 1081.711 | 1142.265 | -0.624 | 0.533 | FALSE |
| RPS6 | 45522.354 | 32365.915 | -0.628 | 0.415 | FALSE |
| FAR1 | 5630.500 | 3283.622 | -0.628 | 1.174 | FALSE |
| RPS13 | 115982.025 | 88846.871 | -0.630 | 0.481 | FALSE |
| COPG2 | 4354.412 | 3541.152 | -0.632 | 0.920 | FALSE |
| TMEM65 | 15977.918 | 18896.212 | -0.637 | 0.415 | FALSE |
| ABLIM1 | 831869.567 | 606382.860 | -0.639 | 0.764 | FALSE |
| SVIL | 91609.047 | 45998.098 | -0.641 | 0.925 | FALSE |
| PHGDH | 70449.520 | 56557.876 | -0.641 | 0.282 | FALSE |
| ANXA1 | 15672.893 | 10031.020 | -0.642 | 0.645 | FALSE |
| SAP18 | 323914.296 | 207970.716 | -0.644 | 0.275 | FALSE |
| RPS15A | 89347.479 | 67541.524 | -0.645 | 0.597 | FALSE |
| CAPRIN2 | 4544.949 | 2823.284 | -0.652 | 0.805 | FALSE |
| STT3A | 6449.245 | 4872.277 | -0.653 | 0.757 | FALSE |
| SRSF3 | 751591.731 | 484815.033 | -0.654 | 0.466 | FALSE |
| YIF1B | 6619.569 | 4589.100 | -0.658 | 1.280 | FALSE |
| ITGB1 | 31848.301 | 24568.628 | -0.658 | 0.732 | FALSE |
| CYB5R3 | 71431.392 | 52296.096 | -0.660 | 1.149 | FALSE |
| SRSF9 | 204316.443 | 122055.461 | -0.660 | 0.276 | FALSE |
| PRAF2 | 17162.186 | 20228.276 | -0.666 | 0.498 | FALSE |
| SRSF1 | 1911363.178 | 1242590.811 | -0.666 | 0.342 | FALSE |
| LUC7L2 | 198190.173 | 126441.889 | -0.667 | 0.393 | FALSE |
| ABLIM2 | 6713.246 | 4619.743 | -0.668 | 0.434 | FALSE |
| DAD1 | 21467.818 | 13048.930 | -0.670 | 0.428 | FALSE |
| RPS23 | 100145.725 | 66183.528 | -0.670 | 0.622 | FALSE |
| TUBB4B | 2329109.939 | 2884476.773 | -0.671 | 0.286 | FALSE |
| MAP1A | 2102.909 | 1922.297 | -0.673 | 0.872 | FALSE |
| CD9 | 72339.198 | 56777.315 | -0.675 | 0.403 | FALSE |
| GNA12 | 9371.392 | 7423.574 | -0.676 | 0.600 | FALSE |
| TOR1AIP1 | 4709.120 | 5042.241 | -0.681 | 0.775 | FALSE |
| NUP98 | 3992.225 | 4687.590 | -0.681 | 0.525 | FALSE |
| TRA2B | 179848.976 | 129032.469 | -0.683 | 0.484 | FALSE |
| SRSF7 | 499376.312 | 303649.963 | -0.683 | 0.391 | FALSE |
| MAVS | 5300.216 | 6516.446 | -0.689 | 0.712 | FALSE |
| CADM4 | 6870.122 | 13459.409 | -0.703 | 0.453 | FALSE |
| ZFPL1 | 5453.084 | 5235.476 | -0.704 | 0.780 | FALSE |
| ERH | ########### | 6134603.333 | -0.710 | 0.385 | FALSE |
| BCLAF1 | 492579.067 | 273263.807 | -0.714 | 0.260 | FALSE |

| | | | | | |
|---|---|---|---|---|---|
| CALM1 | 93215.483 | 73149.323 | -0.723 | 0.221 | FALSE |
| SRPRA | 4641.357 | 5390.734 | -0.727 | 0.594 | FALSE |
| XPO5 | 4965.910 | 6325.968 | -0.727 | 0.496 | FALSE |
| DBN1 | 6893.003 | 3717.847 | -0.728 | 1.139 | FALSE |
| SRSF4 | 502807.617 | 334038.667 | -0.734 | 0.455 | FALSE |
| GNAI2 | 30128.500 | 23998.063 | -0.736 | 0.577 | FALSE |
| STOML2 | 17898.519 | 12761.798 | -0.741 | 0.832 | FALSE |
| TTYH3 | 5699.498 | 5904.315 | -0.746 | 0.813 | FALSE |
| COPE | 4221.966 | 2733.438 | -0.746 | 0.372 | FALSE |
| RPL22 | 50507.409 | 33124.107 | -0.747 | 0.253 | FALSE |
| EIF5 | 6071.136 | 7722.329 | -0.750 | 0.393 | FALSE |
| SEL1L | 4700.732 | 5677.328 | -0.761 | 0.520 | FALSE |
| AGPS | 20019.501 | 14405.927 | -0.762 | 0.825 | FALSE |
| EPHX1 | 11129.814 | 7001.620 | -0.766 | 1.634 | FALSE |
| SSBP1 | 12251.685 | 6412.611 | -0.768 | 1.939 | FALSE |
| TOMM70 | 2280.994 | 2117.545 | -0.771 | 1.140 | FALSE |
| CCDC47 | 5137.307 | 5435.871 | -0.776 | 1.159 | FALSE |
| RTN3 | 8811.247 | 5876.395 | -0.780 | 1.233 | FALSE |
| RPL28 | 26985.783 | 21626.640 | -0.784 | 0.215 | FALSE |
| YWHAH | 256987.278 | 145364.476 | -0.793 | 0.923 | FALSE |
| YWHAB | 365037.412 | 209835.518 | -0.803 | 1.171 | FALSE |
| IPO7 | 10829.399 | 13490.998 | -0.816 | 0.386 | FALSE |
| RNPS1 | 88660.073 | 51269.911 | -0.828 | 0.323 | FALSE |
| SCAMP3 | 16371.209 | 18716.338 | -0.830 | 0.633 | FALSE |
| XPOT | 5078.011 | 4881.157 | -0.834 | 0.372 | FALSE |
| PTGES2 | 18024.025 | 10561.261 | -0.837 | 2.831 | FALSE |
| PODXL | 3238.960 | 1737.557 | -0.843 | 0.753 | FALSE |
| SYNGR2 | 38960.008 | 27849.938 | -0.844 | 0.834 | FALSE |
| RPL37A | 18709.655 | 9729.916 | -0.845 | 0.364 | FALSE |
| PPIG | 25982.248 | 23549.928 | -0.847 | 0.294 | FALSE |
| YWHAG | 357785.926 | 190410.249 | -0.848 | 1.373 | FALSE |
| NOMO1 | 1436.858 | 1200.720 | -0.849 | 1.291 | FALSE |
| RPLP2 | 92507.580 | 71541.665 | -0.853 | 0.233 | FALSE |
| IPO11 | 2161.991 | 1740.814 | -0.853 | 0.662 | FALSE |
| DPM3 | 33297.732 | 38898.290 | -0.856 | 0.910 | FALSE |
| YWHAE | 422466.713 | 238655.630 | -0.858 | 1.143 | FALSE |
| ITGB6 | 4839.169 | 3146.130 | -0.868 | 1.216 | FALSE |
| IQGAP3 | 1527.233 | 1421.344 | -0.893 | 0.820 | FALSE |
| ATP5PO | 31608.961 | 18414.261 | -0.910 | 0.650 | FALSE |
| OCIAD1 | 16847.336 | 17036.573 | -0.910 | 0.641 | FALSE |
| LETM1 | 5848.959 | 5976.144 | -0.911 | 0.831 | FALSE |
| GPD2 | 9947.099 | 11519.330 | -0.916 | 0.595 | FALSE |
| RAB32 | 28613.408 | 9317.692 | -0.919 | 1.312 | FALSE |

| | | | | | |
|---|---|---|---|---|---|
| SYPL1 | 12885.835 | 8294.074 | -0.920 | 1.240 | FALSE |
| SRSF10 | 63595.265 | 39825.194 | -0.936 | 0.473 | FALSE |
| VAMP2 | 37202.983 | 21065.384 | -0.945 | 0.702 | FALSE |
| YWHAZ | 438255.711 | 242911.270 | -0.965 | 1.317 | FALSE |
| AAAS | 5426.106 | 5286.524 | -0.975 | 0.658 | FALSE |
| TRA2A | 63778.851 | 32695.969 | -0.979 | 0.603 | FALSE |
| ADM | 26120.232 | 13642.921 | -0.985 | 0.558 | FALSE |
| ATP5PB | 24570.518 | 13897.129 | -0.998 | 0.611 | FALSE |
| VTI1B | 9176.179 | 19652.917 | -1.020 | 0.491 | FALSE |
| TMEM123 | 24904.967 | 14906.765 | -1.047 | 0.906 | FALSE |
| YWHAQ | 272893.806 | 156426.708 | -1.067 | 1.154 | FALSE |
| ATXN10 | 9057.449 | 7783.316 | -1.087 | 0.835 | FALSE |
| CD63 | 16838.656 | 9924.287 | -1.088 | 0.652 | FALSE |
| ATP5PD | 16546.243 | 15334.351 | -1.167 | 0.533 | FALSE |
| CSN1S1 | 480474.028 | 256501.188 | -1.235 | 0.336 | FALSE |
| MYO1B | 19193.963 | 9298.649 | -1.255 | 0.406 | FALSE |
| SRRM2 | 92126.943 | 37350.569 | -1.344 | 0.339 | FALSE |
| CSN3 | 259589.711 | 102892.380 | -1.378 | 0.338 | FALSE |
| SRRM1 | 28423.142 | 9995.036 | -1.401 | 0.363 | FALSE |
| CSN2 | 582589.963 | 233195.299 | -1.471 | 0.352 | FALSE |
| LGB | 440855.526 | 151494.048 | -1.676 | 0.379 | FALSE |
| CSN1S2 | 221209.464 | 83898.973 | -1.760 | 0.372 | FALSE |
| MYO1C | 78404.807 | 25539.479 | -1.876 | 0.524 | FALSE |
| KRT13 | 1291293.583 | 1149992.146 | -2.095 | 0.684 | FALSE |
| KRT4 | 712426.188 | 631604.208 | -2.103 | 1.294 | FALSE |
